# A climate and population dependent diffusion model forecasts the spread of *Aedes Albopictus* mosquitoes in Europe

**DOI:** 10.1101/2024.09.20.614113

**Authors:** Sandra Barman, Jan C. Semenza, Pratik Singh, Henrik Sjödin, Joacim Rocklöv, Jonas Wallin

**Author notes:** **Corresponding author:** Sandra Barman. These authors contributed equally (Joacim Rocklöv, Jonas Wallin).

## Abstract

Vectors of Dengue, Chikungunya, Zika, and Yellow Fever are emerging in new areas, posing increasing public health risks. *Aedes albopictus*, a key vector for these diseases, is expanding its range beyond its tropical and subtropical origins, driven by suitable climate, population mobility, trade, and urbanization. Since its introduction to Europe, *Ae. albopictus* has rapidly spread and triggered recurrent outbreaks.

Past model attempts have handled vector suitability and vector introduction as independent drivers. Here we develop a novel, highly predictive spatio-temporal vector diffusion model based on minimum temperature, median temperature, relative humidity and human population as predictors. The model predicts areas of presence or absence with an accuracy of 99% and 79% for new established vector populations. The model explains how short- and long-range spread of *Ae. albopictus* interacts with vector suitability.

These results show that the expansion of *Ae. albopictus* in Europe is predictable and closely linked to climate suitability, and human population. The new model integrates in one simultaneous model framework the climate and mobility drivers, providing a better basis for anticipating future outbreaks in situations of dependent interacting co-drivers.

**Significance statement:** The Tiger mosquito, *Aedes albopictus*, which spreads diseases like Dengue, Chikungunya, Zika and Yellow fever, has accelerated its presence in Europe over the past decades. This poses a public health risk as Europe face an upsurge of autochthonous arbovirus transmission events. The change is partly due to conducive environments, climate change and human mobility and trade. We develop a novel model explaining the spread of the mosquito over the years 2010-2023, observing a change in recorded mosquito presence areas from 138 regions in 2010 to 537 regions in 2023. The model incorporates patterns in both space and time, capturing with high accuracy how the mosquito spreads from region to region based on climate suitability, geographical diffusion and human population density.

## Introduction

Dengue, chikungunya, zika or yellow fever are caused by arthropod-borne viruses, transmitted by *Aedes* mosquitoes. Currently, almost 4 billion people, in over 129 countries, are at risk for *Aedes*-borne infections.^1^ Europe has also experienced recurrent dengue and chikungunya outbreaks, due to autochthonous transmission by invasive *Aedes* mosquitoes. Since 2010, there have been 18 dengue and 5 chikungunya outbreaks on mainland Europe. While the disease burden cannot be compared to tropical and subtropical countries, the epidemic potential of these diseases is of concern especially in the light of climate change.^2,3^ The introduction, establishment and spread of invasive vector species has been assisted by global environmental change, specifically travel, trade, urbanization, and climate change. Moreover, viraemic travellers returning from disease endemic countries increasingly introduce dengue and chikungunya viruses into areas with climatically suitable conditions, where competent mosquito vectors are established or re-established. These circumstances have resulted in an upsurge of local outbreaks in Europe.^4^

The mosquito vector responsible for these European outbreaks is *Aedes albopictus*^*5*^. It originated in tropical forests of South-East Asia and has since invaded all continents.^6^ This rapid and widespread expansion has been propelled by its remarkable ecological and physiological plasticity from zoophily to anthropophily, domestic container-breeding, diurnal biting habits, cold-acclimatization, and desiccation-resistant eggs.^7^ The passive transport of its eggs, aided by globalized travel and trade in e.g. used tires or lucky bamboo, enabled *Ae. albopictus* to invade novel habitats.^8,9,10^ In Europe, *Ae. albopictus* spread along transportation corridors from heavily infested to naïve areas, through passive transport on ground vehicles.^11,12,13,14^

*Ae. albopictus* was first detected in Albania in 1979 but might have been present there already since 1976.^15^ It was not until 1990, when it was discovered in another European country, close to a playground in Genova, Italy, where children played with discarded tires.^16^ Despite an attempted eradication campaign, *Ae. albopictus* became established throughout Italy, in areas below 600 meter above sea level, with particularly high densities in urban centres. Subsequently, *Ae. albopictus* was detected in France in 1999, and has since expanded to a number of other European countries.^17^ Suitable climatic conditions favoured by climate change, urbanization, and human population mobility, seems to have facilitated the expansion of this invasive mosquito species into novel habitats.^18,19^ However, based on the current introductions and subsequent establishment of *Ae. albopictus* it is not clear what processes and drivers contribute to a successful colonization of naïve areas and to what extent it is predictable.

The WHO global arbovirus initiative, launched in March 2022, calls for risk monitoring and the mapping of areas vulnerable for arbovirus transmission.^20^ However, defining and predicting which *Ae. Albopictus*-free areas will become colonised has proven challenging.^21^ Establishment of mosquitoes in an uncolonized area, depends in part on the origin of the imported mosquito species, but also on climatic and environmental conditions, population mobility and demographic factors.^6,22,23^ Here we present a novel dynamic, spatio-temporal diffusion model with human population network structures that in combination with the influences of climatic predictors and process-based model predictions of mosquito dynamics, forecasts the newly invaded areas of *Ae. albopictus* in Europe. The model reliably separates and integrates the influence of local vector suitability with vector introduction and predicts the continuing European invasion by first time (non-recurrent) establishment of *Ae. albopictus*. The novel model framework developed lends itself to be used for public health purposes in Europe and beyond to prepare for the introduction of *Ae. albopictus*, or *other similar challenges where the processes of suitability and introduction need to be considered simultaneously*. Here we describe the drivers and discuss how insights from such predictions can guide public health efforts and reduce the epidemic risk in unaffected areas where the local population has no immunity to *Aedes*-borne diseases. Predictions can help target awareness and prevention messages to susceptible populations and guide vector control efforts to counteract establishment of vector populations and local outbreaks. Moreover, it can help prepare the healthcare system for epidemics and target novel interventions to areas at risk.

## Results

We have developed a spatio-temporal model with a diffusion process that models the spread of *Ae. albopictus* from regions with presence to nearby absence regions. The model predicts the introduction of *Ae. albopictus* into naïve regions of Europe, during the period 2010-2023 (Fig. 1). We integrate the diffusion model with covariates determining the suitability of *Ae. albopictus* establishments using climate and human population data.

**Figure 1:**
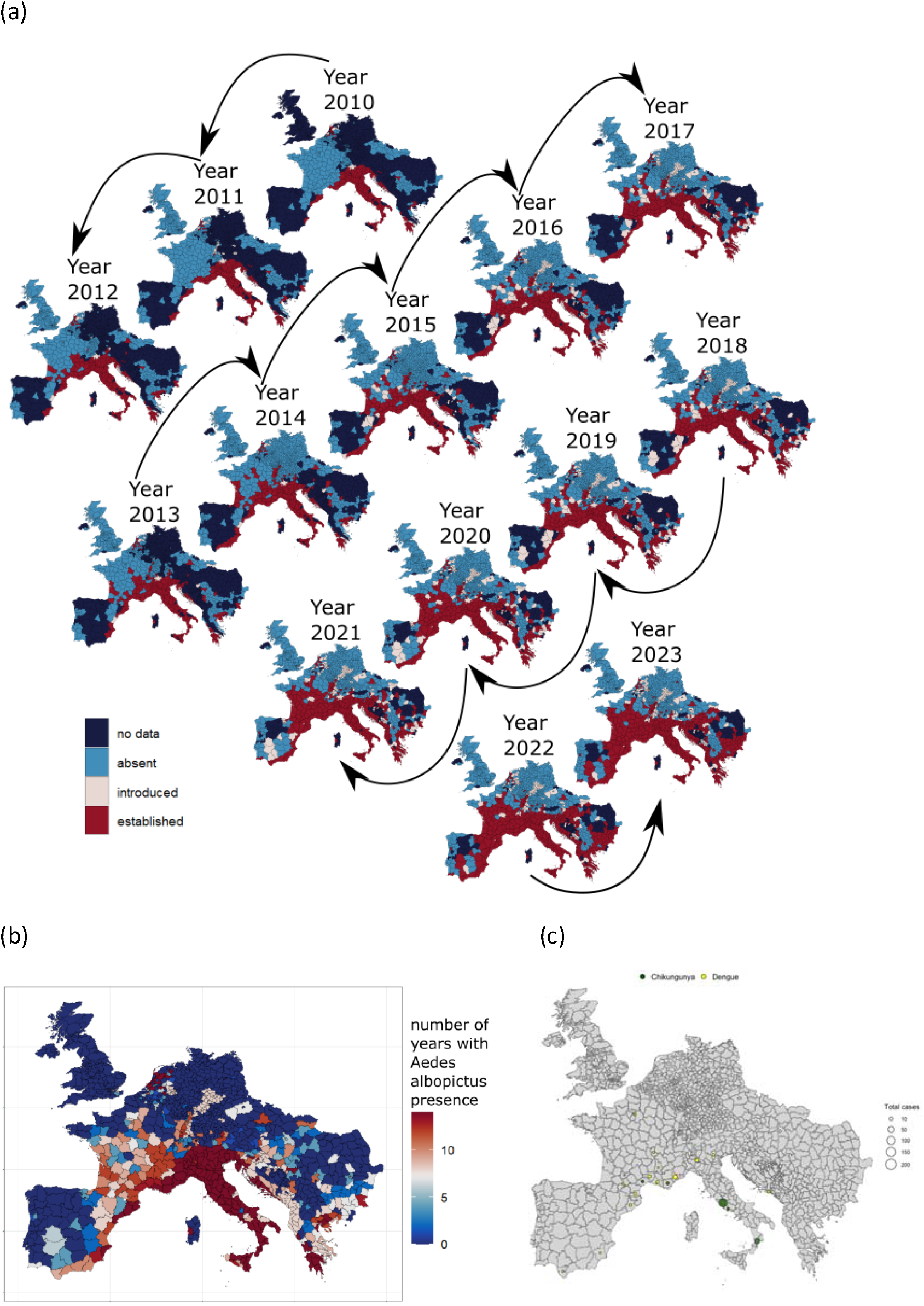
(a) VectorNet dataset of observations of *Ae. albopictus*, from 2010-2023, coloured by albopictus status: no data; absent; introduced; and established, (b) accumulated *Ae. albopictus* establishment over the years 2010-2023 and (c) local outbreaks of Dengue and Chikungunya.

The model is a generalized additive mixed (GAM) model with non-linear random effects, which is fitted within the approximated Bayesian spatio-temporal framework INLA based on annual data by NUTS3 areas. We predict a year ahead using a model trained on previous years data. The overall model performance is good, with high area under the receiving operating characteristic curve (AUC) values of 0.99 when evaluated on all observations independent of time of establishment of the vector. It is, however, easier to model the probability of *Ae. albopictus* presence re-occurring in a region as the recurrence rate in the same region is 99.8%. Thus, it is relevant to show model performance statistics only for new observations in a specific year, i.e., region-year locations where *Ae. albopictus* was not present the previous year. We show that the accuracy of our model is high with an AUC of around 0.80 (Table 1). The ROC curve used to estimate AUC values are presented in Fig. 2. (For more details about the model fit see Table S8 in the Supplement).

**Table 1.**
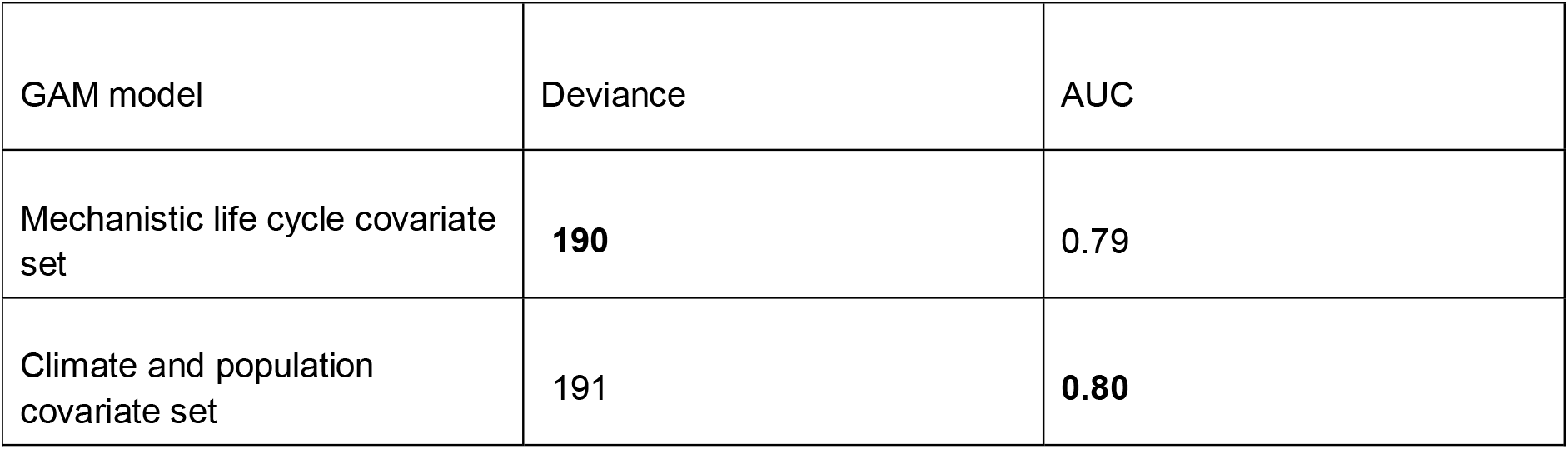
Comparison of the GAM model fit for the two sets of covariates. Both covariate sets include the proximity, presence previous year and first year covariate, see Section Methods. Deviance and AUC are computed on test data over seven cross-validation test sets, withholding one year ahead at a time for each of the years 2017-2023. Bold font indicates the value with the best fit.

**Figure 2.**
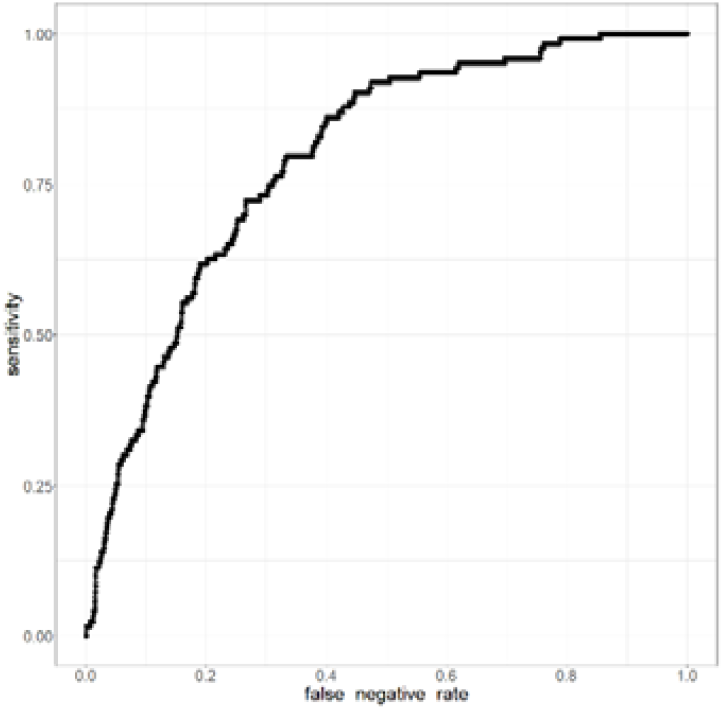
ROC curve for the mechanistic life cycle GAM-model, with region-year locations being left out one year ahead, with each of the years 2017 to 2023 being left out in turn. Only regions where *Aedes albopictus* was not present the previous year were included when computing the ROC curve.

The GAM model is fitted with a spatio-temporal SPDE term which accounts for extraneous variables not included in the model. A short-range SPDE is used, which (1) is conservative in that a large portion of the fitted values are essentially zero, (2) has a low correlation with the data and (3) has the effect that more weight is put on the covariates which improves the model fit on out-of-sample data (see Supplement, Controlling the SPDE range to improve model predictions).

We calibrate two different GAM models with climate and human population covariates considered in different ways. In one of the models the raw climate (median temperature, minimum temperature, and median relative humidity) and human population covariates are taken into account as non-linear random effects. In the other model these covariates are transformed by a mechanistic mosquito life cycle-model predicting adult *Ae. albopictus* abundance. The predicted adult abundance is then entered into the GAM model as a covariate to which a non-linear random effect. The diffusion process modelling the spread of *Ae. albopictus* from regions with presence to nearby regions, termed the proximity covariate, is included in both versions of the model. Summary statistics for the fit from each of these two models (termed the climate and population- and the mechanistic life cycle-GAM model, respectively) are presented in Table 1, and fitted non-linear random effects of covariates are shown in Fig. 3-4.

**Figure 3.**
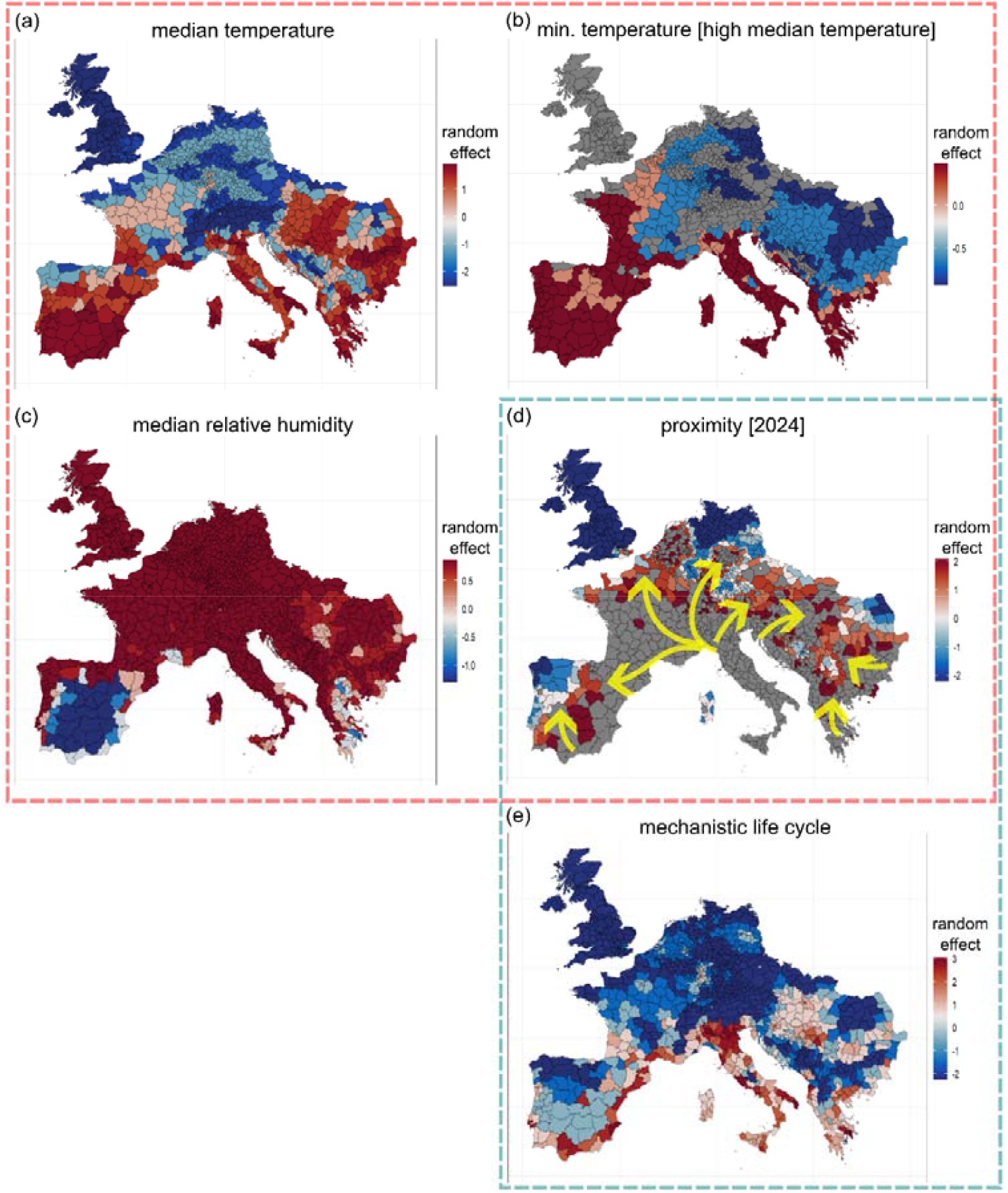
Random effect shown geographically, for (a) the median temperature; (b) the minimum temperature, conditioned on high median temperature; (c) the median relative humidity; (d) the proximity; and (e) the mechanistic life cycle covariate. The proximity covariate is different from year to year, it is here shown at year 2024. Regions in grey in the proximity map are regions that have recorded *Ae. albopictus* presence in 2023 and yellow arrows illustrate visually how the proximity covariate models risk of *Ae. albopictus* spreading from regions with presence to nearby regions with no presence. The maps indicated by the red and green dashed lines show covariates included in the climate and population GAM-model ((a)-(d)) and the mechanistic life cycle GAM-model-((d)-(e)), respectively.

The predictive performance of the model is further broken down by year, by distance to regions with recorded *Ae. albopictus* presence, and by geographical region with slightly reduced predictive performance for some of the subsets, for example, with longer distance from established areas (Table 2). When broken down by geographical region, the performance is more distinctly reduced for the south-west geographical region - The Iberian peninsula - which has a relatively high number of regions with status “no data” (Fig. 1(a)).

**Table 2.**
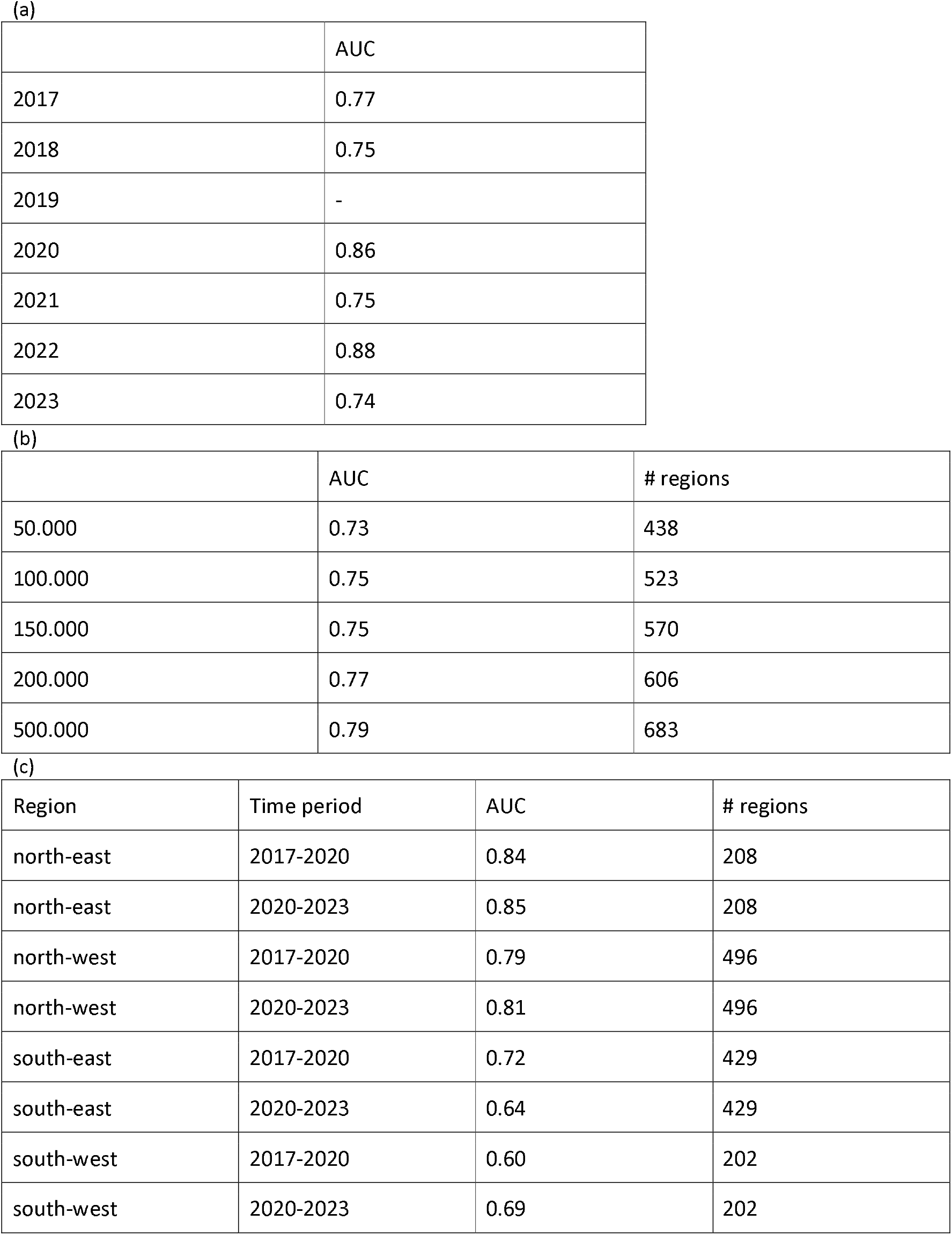
AUC reported for subdivisions of the data into year / geographical region for the mechanistic life cycle GAM-model. AUC was computed on test sets withholding one year ahead for each of the years 2017 to 2023. Only regions with confirmed *Ae. albopictus* presence or absence, but no recorded presence previous years are counted. (a) AUC for each of the years 2017-2023. 2019 had no new observations of *Ae. albopictus* presence. (b) AUC for the years 2017-2023, divided by distance to regions with recorded presence the previous year. (c) AUC by geographical region and by time period, with north: latitude >= 50, south: latitude < 50, east: longitude >= 10, west: longitude < 10. Only regions with confirmed *Ae. albopictus* presence or absence are counted.

|  | AUC |
| --- | --- |
| 2017 | 0.77 |
| 2018 | 0.75 |
| 2019 | - |
| 2020 | 0.86 |
| 2021 | 0.75 |
| 2022 | 0.88 |
| 2023 | 0.74 |

(b)
|  | AUC | # regions |
| --- | --- | --- |
| 50.000 | 0.73 | 438 |
| 100.000 | 0.75 | 523 |
| 150.000 | 0.75 | 570 |
| 200.000 | 0.77 | 606 |
| 500.000 | 0.79 | 683 |

Table 2. AUC reported for subdivisions of the data into year / geographical region for the mechanistic life cycle GAM-model.
| Region | Time period | AUC | # regions |
| --- | --- | --- | --- |
| north-east | 2017-2020 | 0.84 | 208 |
| north-east | 2020-2023 | 0.85 | 208 |
| north-west | 2017-2020 | 0.79 | 496 |
| north-west | 2020-2023 | 0.81 | 496 |
| south-east | 2017-2020 | 0.72 | 429 |
| south-east | 2020-2023 | 0.64 | 429 |
| south-west | 2017-2020 | 0.60 | 202 |
| south-west | 2020-2023 | 0.69 | 202 |

The fitted random effects can be compared with empirical relationships between covariates and *Ae. albopictus* presence in Fig. 4 and 5. See also Tables S1-S7 and Fig. S4 of the Supplementary for more details related to Fig. 4 and 5.

**Figure 4.**
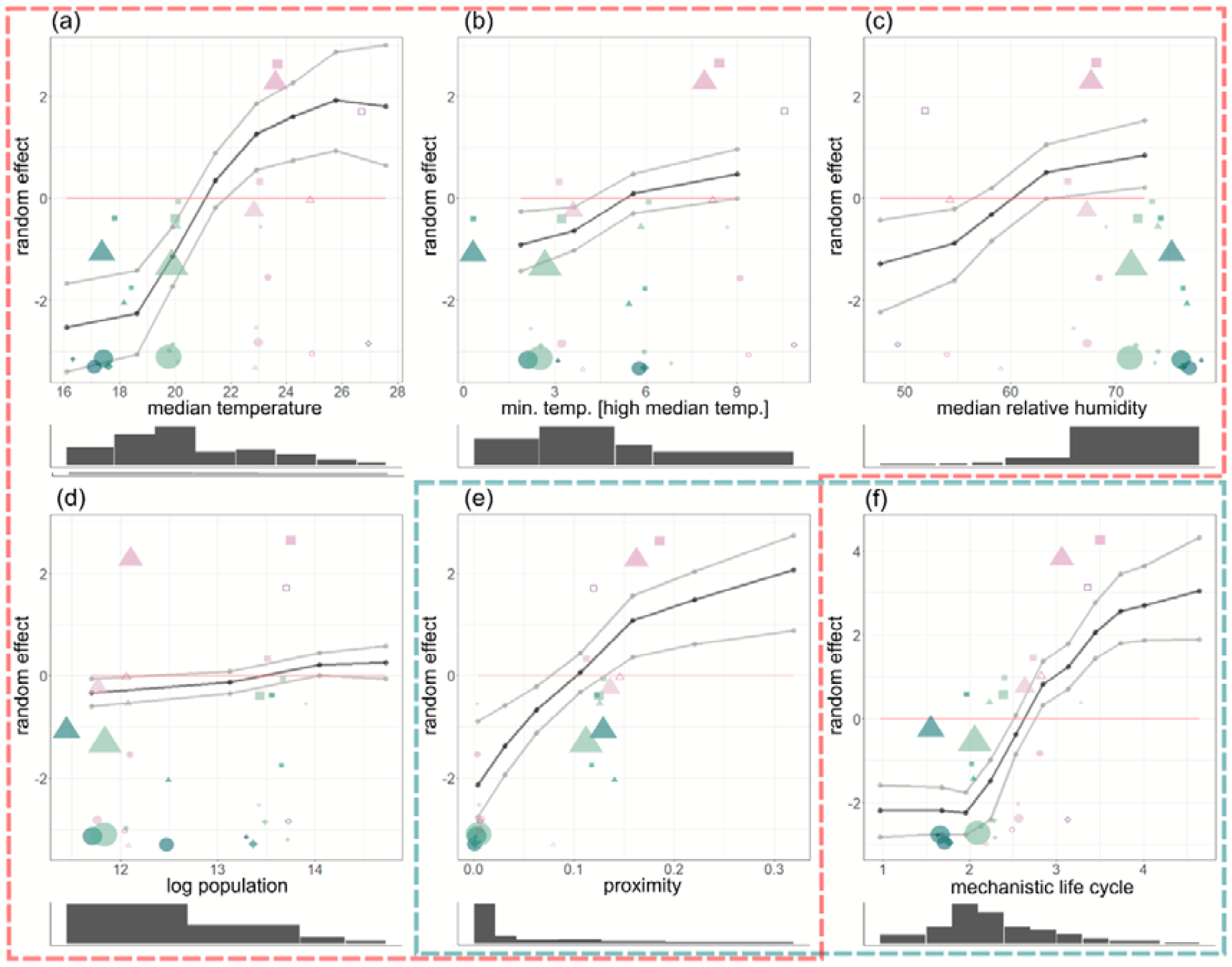
Random effect shown on functional form, for (a) the median temperature; (b) the minimum temperature, conditioned on high median temperature; (c) the median relative humidity; (d) the log human population; (e) the proximity; and (f) the mechanistic life cycle covariate. Black lines indicate the random effect, and gray lines indicate the lower and upper limits of 95% credible intervals. Scatter plots of the data, by a five-way grouping of covariates given in Table S7 of the Supplementary, are super-imposed on the plots. Refer to Fig. S4 in the Supplementary for an explanation of the symbols and colouring. The figures indicated by the red and green dashed lines show covariates included in the climate and population GAM-model ((a)-(e)) and the mechanistic life cycle GAM-model ((e)-(f)), respectively.

**Figure 5.**
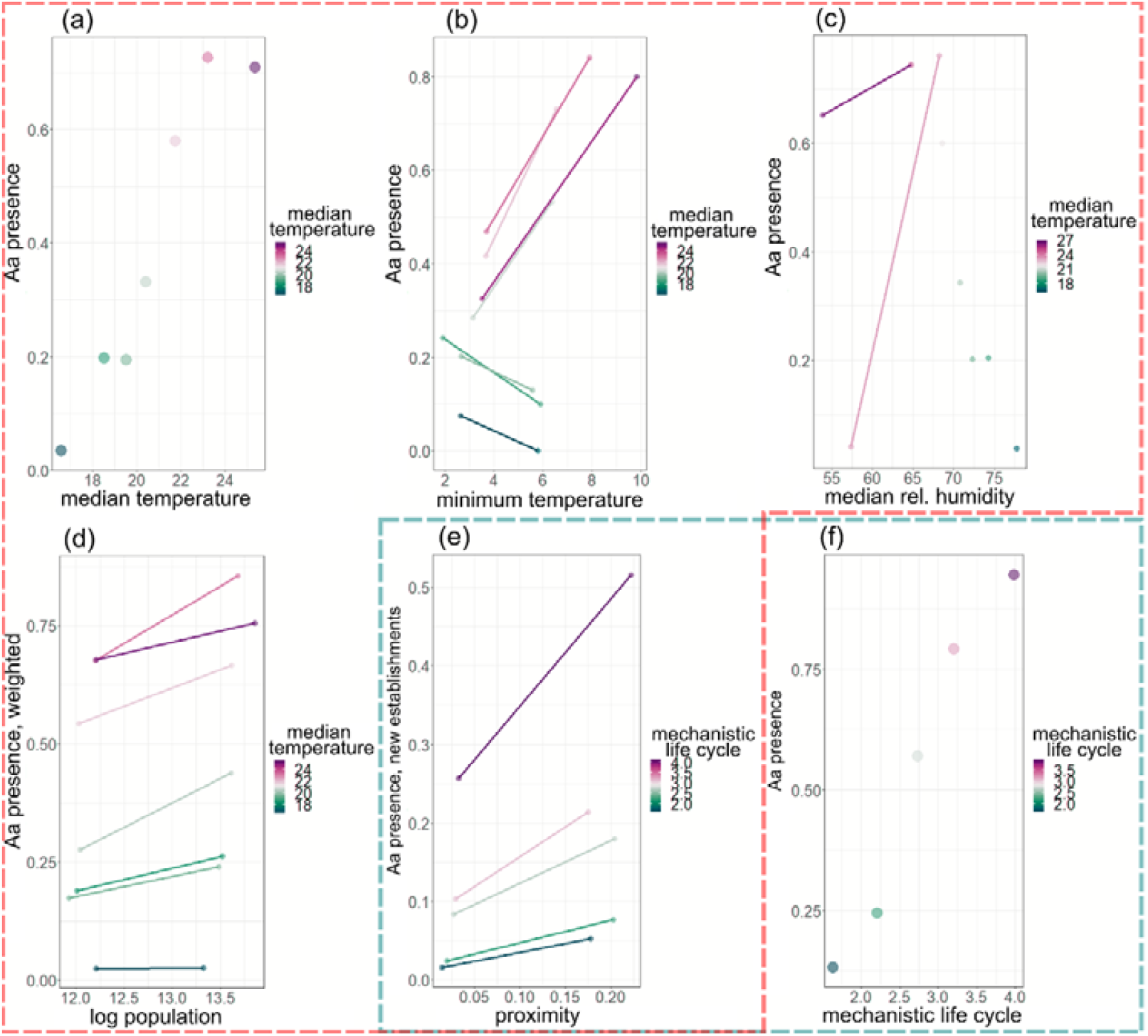
The empirical ratio of *Ae. albopictus* presence is shown for different sub-groups of the data. (a) The median temperature; (b) the median temperature crossed with minimum temperature; and (c) the median temperature crossed with the median relative humidity; (d) the log human population crossed with the median temperature; (e) the mechanistic life cycle covariate crossed with the proximity; and (f) the mechanistic life cycle covariate. The same median temperature groups have been used in panels (a)-(c), and the same mechanistic life cycle covariate groups have been used in panels (e)-(d), and pairs within the same median temperature or mechanistic life cycle covariate group are linked with lines. The sub-groups in (a)-(d) are shown for the whole dataset, whereas the sub-groups in (e)-(f) are shown for only new establishments. The data and groupings used to create these plots can be found in Tables S1-S6. The figures indicated by the red and green rectangles show covariates included in the climate and population GAM-model ((a)-(e)) and the mechanistic life cycle GAM-model ((e)-(f)), respectively.

We find that the mechanistic life cycle GAM-model performs similarly on out-of-sample data compared to with the climate and population GAM-model (Table 1). The combined random effects of the climate and human population covariates (Fig. 3(a)-(c), Fig. 4(a)-(d)) can be compared to the random effect of the mechanistic life cycle covariate (Fig. 3(e), Fig. 4(f)), as the underlying mechanistic mosquito life cycle-model takes both human population and climate data as input.^24^

### Climate covariates

The raw climate covariates (median temperature, minimum temperature, and median relative humidity) were binned into relatively few bins to reduce overfitting and to reduce the dependence between the covariates. All but the minimum temperature covariate was computed over the warmer months April-October, and minimum/median values were taken over 30-day averages. The non-linear random effects show the strongest effects overall for median temperature (Fig. 3(a), Fig. 4(a)). The fitted random effect increases at median temperatures up to 24°C, to then level out and start to decrease at temperatures above 25°C (Fig. 4(a)).

We included a random effect of minimum temperature conditioned on high median temperature (Fig. 3(b), Fig. 4(b)). This since the random effect for minimum temperature shows a positive correlation with *Ae. albopictus* presence (Aa presence) when conditioned on high median temperatures (median temperature >= 19°), and a negative correlation when conditioned on low median temperatures (Fig. 4(b)). The empirical Aa presence supports this conditioned relationship (Fig. 5(b)). The distribution of *Ae. albopictus* can be seen to follow a pattern that correlates with both median and minimum temperature, having been present for the longest along coastal areas around the Mediterranean (Fig. 1(a)) which have both high median and minimum temperature.

*Ae. albopictus* presence is negatively correlated with low levels of relative humidity (Fig. 3(c), Fig. 4(c)). Regions in Spain have relatively low rates of presence, given the risk posed by high temperature and proximity to regions with presence.

### Proximity and human population covariates

We hypothesized that *Ae. albopictus* spreads to regions that are geographically close and through human mobility to highly populated regions farther away. We used a spatio-temporal diffusion process to estimate the rate of spread to regions that are geographically close (termed the proximity covariate), and the logarithm of human population as a covariate to capture mobility to highly populated regions further away. The fitted random effects show a significant impact of geographical proximity through the spatio-temporal diffusion process (Fig. 3(e), Fig. 4(e)) and there is a clear effect of *Ae albopictus* presence being higher in regions with higher population (Fig. 4(d)), supporting the hypothesis. Although there is a clear effect of the population covariate, the size of the effect is relatively small compared to the fitted random effects of other covariates. The result that the random effect is relatively small for the population covariate, is also seen when comparing the raw population data with empirical *Ae. albopictus* presence (Fig. 5). There is a clear trend related to population here (Fig. 5(d)), but the trend is relatively small in size compared to that found for the other covariates (Fig. 5(a)-(c),(e)-(f)).

We additionally tried a GAM model including human mobility using a radiation model in the proximity diffusion process.This did not result in any significant improvement in model fit though, and so this GAM model is not included in the results presented here. Instead, the local diffusion and human population covariate was used to efficiently predict the new establishments.

### Mechanistic life cycle covariate

The fitted random effect for the mechanistic life cycle covariate (Fig. 3(e)) shows similar effects for regions close to the Atlantic and around the Mediterranean Sea as discussed in relation to the climate covariates. Further, there is a relatively high risk of *Ae. albopictus* presence in the northern part of Europe, namely in the Low Countries (Belgium, Luxembourg and the Netherlands), and in parts of Germany and northern France. Comparing the fitted random effect from the mechanistic life cycle covariate (Fig. 3(e)) and the accumulated *Ae. albopictus* presence over the full dataset (Fig. 1(b)), we see some striking similarities. Note the *Ae. albopictus* presence in Berlin in year 2023, which is the first noted presence in the northern part of Germany.

## Discussion

The introduction, establishment, and expansion of *Ae. albopictus* in Europe are the primary causes of recurrent dengue and chikungunya outbreaks. Monitoring this expansion, through labour- and time-intensive vector surveillance has proven to be cumbersome and costly.^25^ However, our study demonstrates that this expansion is predictable, based on a vector population spatial diffusion process, with climate and human population covariates depicting suitability. This innovative model performs well, also on data withheld from model fitting, and will prove to be a valuable tool for preparedness and response to *Aedes*-borne infections. Notably, model evaluation reveals that new introduction of *Ae. albopictus* into naïve areas, are very well predicted, which has not been achieved before. In contrary to Kraemer et al.,^6^

We find that temperature, relative humidity, long-distance mobility and short-distance diffusion is enough to predict presence/absence and more interestingly the spatio-temporal patterns of new establishment of *Ae. albopictus* in Europe. In fact, for the first time, the models document an ability not only to predict presence/absence, but the changing risk landscape by predicting areas at risk for new establishment which is a more relevant outcome for timely early warnings and adaptation priorities. We calibrated two different models using the spatio-temporal Bayesian INLA modelling framework describing the patterns of the new establishments in Europe. The first model uses simply climate covariates, but the second model uses a climate-driven prediction from a mechanistic mosquito life cycle model as covariate. Both models include a diffusion process capturing how new establishments spread from regions with *Ae. albopictus* presence. We find both models provide reliable predictions on data withheld from model fitting with a classification error of around 20% when applied to naïve regions not previously colonized by *Ae. albopictus*.

In detail, we incorporate climate and demographic components in the model, such as relative humidity, temperature, rainfall, and human population data. We introduce a continuous spatio-temporal diffusion process for vector introductions from regions with *Ae. albopictus* presence, mainly contributing to the spread at shorter distances. Human mobility, which can be a key factor in the spread at both short and long distances, is included implicitly with a human population covariate. These covariates perform well together and show that all of these factors are important for the establishment of vectors in new areas, as is the mechanistic mosquito life cycle covariate which captures all of these processes in one covariate.

The finding that the mobility diffusion did not add significantly to the model performance fits with previous observations^6^, where including mobility was found to only give slight improvements in model fit. In our analysis we therefore chose the simpler model where the proximity-diffusion process and human population are included as separate covariates. Visual inspection of the accumulated *Ae. albopictus* presence indicates a clear effect of human mobility when comparing the presence in large cities (Berlin, London, Madrid, Porto, Prague and Thessalonique) with their surrounding regions (Fig. 1(b), Fig. S2-S3). Thus, there seems to be a clear effect of human population size and mobility in the raw data, though with the metrics we are using the measured effect is relatively small.

Interestingly the mechanistic mosquito life cycle covariate, with the underlying process-based life cycle model driven by climate and human population, does capture the patterns as well as the more flexible climate covariates. The same process-based model used is frequently applied to larger climate assessments in the *Lancet* countdown on climate change and health.^26,27^ These findings provide insights into the reliability of such more complex process-based models and provide proof of its validity.

The approach presented here is using the most recent data in a statistical framework that quantifies the contribution of known factors to the spread of *Ae. albopictus*. The work further shows that human population and climate along with a spatio-temporal diffusion can be integrated in a simultaneous modelling framework and that it can robustly predict the establishment through accounting for vector suitability and vector introduction processes. Compared to previous prediction models, the new model is reducing the model complexity essentially. Although a previous study in predicting presence-absence of vectors, the past modelling attempts of *Ae. albopictus* in Europe used climate and human population as well as a larger dataset of environmental and social data^6^, the nature of the previous model didn’t allow statistical inference in terms of the estimated explicit relationships between model components and risk of new *Ae. albopictus* establishments. Especially, there is a risk of covariate bias as it lacked a simultaneous estimation of the key drivers of vector suitability and vector introduction in one model framework. We additionally explore and describe both empirical and fitted non-linear relationships between each model component and *Ae. albopictus* presence, providing insight into the dynamics of new establishments. This can help to better understand and predict emergence and by so guide future surveillance efforts.

Additionally, to the insights, the model framework and model may lend itself for early warning systems predicting the emergence of Ae. Albopictus, and other threats, in naïve areas. It could also be deployed as a climate-service using long-term projections or inter-annual early warnings to enhance public health preparedness within Europe.^28^

Limitations of inferences drawn from the model come partly from *Ae. albopictus* surveillance, where we have regional differences in surveillance efforts across countries, and where we expect more effort close to regions with recorded presence as well as in more densely populated regions. From the modelling perspective, we have rather coarse data with presence / absence on a yearly basis and on the NUTS3 / GAUL spatial resolution, which limits the inference. On the other hand, the study does not make use of pseudo-absence observations which may introduce bias.

In conclusion, our approach, using the most recent data and a streamlined statistical framework, offers several advantages: updated vector surveillance data, a simpler model structure, integration into a coherent simultaneously modelling framework capturing both vector suitability and vector introduction processes, statistical inference capabilities, and a focus on predicting new establishments. This model can serve as an early warning system, predicting the emergence of *Ae. albopictus* in new areas and enhancing public health preparedness in Europe without having to rely on vector surveillance data.^29^ Additionally, this framework could be adapted for use in other regions and for other vector-borne diseases.^30^

## Materials and methods

### Dataset

#### Aedes albopictus

The data comes from the project VectorNet, funded by the European Food Safety Authority and the European Centre for Disease Prevention and Control. The project collects data from different sources, and records confirmed observations of presence (established or introduced), absence and unknown/no data status of Aedes albopictus within administrative units in European countries and within some countries outside Europe. For countries belonging to the European Union (EU), the administrative units are NUTS (Nomenclature of Territorial Units for Statistics) regions, and for the countries outside the EU the administrative units are GAUL (Global Administrative Unit Layers) regions. We have chosen to work with those countries in Europe where there has been at least one observation of *Aedes albopictus* presence.

When cleaning the data, some observations had no associated geometry. This was solved by downloading maps of regions from NUTS version 2010. Some regions in the joined map were overlapping. Most often, this was with one set of regions having unknown/no data status and the other set having observations of presence or absence of *Aedes albopictus*. In Greece, however, some regions were located on top of each other with contradicting data of presence and absence. In this case, we choose the observations with presence over absence. The full dataset is shown in Fig. 1(a),(b).

#### Climate

Historical climate data for the year 2010 – 2023 were obtained from the Copernicus Climate Data Store (https://cds.climate.copernicus.eu/). Near-surface air temperature(°C) and total precipitation (mm) are obtained from the fifth generation of the European Centre for Medium-Range Weather Forecasts atmospheric reanalyses (ERA5). The grided ERA5-land hourly climate dataset (https://cds.climate.copernicus.eu/cdsapp#!/dataset/reanalysis-era5-land?tab=overview) is converted to daily estimates at a resolution 0.25 × 0.25 degree which approximately covers an area of 25 *km*^2^. Daily mean, minimum, maximum temperature and daily total rainfall were extracted from hourly ERA5-land dataset. All temporal and spatial data aggregation processes were conducted by utilizing the Climate Data Operators (CDO) software.^31^ The human population density data and population count is obtained from GPWv4 data set for the year 2010, 2015 and 2020.^32^ The population density of the rest of the years in between 2010 – 2020 and 2021 – 2023 is retrieved using linear interpolation and extrapolation respectively.

#### Human population

The human population data that was used as a covariate comes from Eurostat (https://ec.europa.eu/eurostat), complemented with population for GAUL regions in the dataset which do not have population data available from Eurostat.

### Model

We use a generalized additive mixed model with a spatio-temporal component^33^ and covariates presented below. The model captures both non-linear and linear effects related to the covariates, and a spatio-temporal Matérn field formulated as an SPDE^34^ is used to control for extraneous variables. The model is constructed as follows: For region *i* and year *t* we denote Aedes albopictus presence / absence as *y* _*i*,*t*_ = 1for presence (introduced or established status) and *y*_*it*_ = 0 for absence. Any locations (*i*,*t*) with no observation (denoted no data in Fig. S1) were excluded from the model fitting. The model can be written as

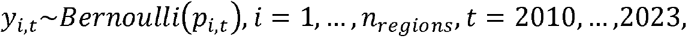

with *p*_*it*_ the probability of Aedes albopictus being recorded present. The log-odds are modelled as a function of the covariates, as

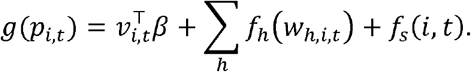

Here g is the logistic link function, v are covariates that enter the model as linear effects, *w* are covariates that enter the model as non-linear random effects. The non-linear random effects *f*_*h*,_ and the SPDE-based process *fs*, are described in the Supplementary. Here follows a description of the covariates.

#### Mechanistic mosquito life cycle-model

We estimated the mosquito abundance of adult Aedes albopictus using a stage-structured, climate-data driven dynamic mechanistic model based on the life cycle of dengue vector’s life stages (see Supplementary, Mechanistic life cycle model details).^35,36,37^ The mechanistic mosquito life cycle-model compute the population-density (number of mosquitoes per hectare) of mosquitoes independently in each grid cell with the assumption of well-mixed mosquito populations within 0.25 × 0.25 latitude longitude grid-cells.^38^ Daily time steps simulations are used for solving the model and then we aggregated the output to yearly values. To account for diurnal temperature variation, simulation for a single day is divided into 100-time steps according to numerical solver (deSolve4 package in R) as 0.14, 0.19 24.00 and temperature at each of these hours is used to simulate the development and mortality rates for one day. For more details on simulation of model and a simple example model simulation code (in octave v4.2.1), see Metelmann et al.^39^ Lastly, spatial aggregation at the NUTS 3 (Europe) level was executed in R version 4.1 utilizing the rasterR6 package.^40^

#### Climate

The climate data, in addition to being used to compute the mechanistic mosquito life cycle-covariate, used in the mosquito life cycle GAM-model, was also used directly as covariates in the climate and population GAM-model. The covariates are constructed from 30-day moving average of precipitation and temperature, and from which we compute the yearly median precipitation, median temperature and minimum temperature denoted simply *median precipitation, median temperature* and *minimum temperature*). These climate summaries are aggregated to the NUTS/GAUL regions by taking the mean value over each region.

#### Human population

Human population plays an important role in the spread and habitat suitability of Aedes albopictus.^41^ We use the total population of each NUTS/GAUL region, transformed to a logarithmic scale as covariate.

#### Presence previous year

If *Aedes albopictus* was recorded as present in a region at year *t*, then in 99.8% of the times it was also recorded as present at year *t* + 1. Thus, it is simple to model the probability of Aedes albopictus re-occuring in a region for this dataset. Re-occurrence is modelled here by adding a covariate *presence previous year*, which at year *t* > 1for region *i* equals one if Aedes albopictus was present in region *i* the previous year, and zero otherwise. It equals zero for all regions at the first year *t* = 2010.

#### First year

The first year, 2010, is different from the later years in the dataset in that it includes an accumulated presence of *Aedes albopictus* from previous years. To account for this difference, we include the first year as a covariate.

#### Proximity

To model how *Aedes albopictus* spreads from regions with presence to neighbouring regions, we developed a spatio-temporal covariate that has the property of diffusing out from regions with *Aedes albopictus* presence. Specifically, we define a Gaussian random field over a network and set the proximity covariate to the conditional expectation of the random field given observations of *Ae. albopictus* presence the previous year. The chosen network-structure then controls how the risk of *Ae. albopictus* presence is diffusing out from regions with presence.

#### Proximity with mobility

We used a radiation model based on diffusion dynamics^42^, to account for how human mobility impacts the spread of *Aedes albopictus*. In particular, we used a version of the radiation model which corrects for finite systems^43^. The radiation model takes as input the distance between regions and their total human population and gives the commuting flow between all pairs of regions. We extracted a mobility network by only keeping the flow that is higher than a specified threshold. Mobility links corresponding to high commuting flow were added to the proximity network (Fig. S5, right), which was then used to create a proximity with mobility-covariate. More details are given in Supplementary, Constructing the proximity covariate.

#### Model setup

We use two combinations of covariates. The first set consists only of the mechanistic covariate. The second set, termed the climate covariates, consists of the median temperature, the minimum temperature (conditioned on high median temperature), the median relative humidity and the log human population covariate. These are combined to form two GAM models, with both models including the proximity-, presence previous year- and first year-covariate.

### Model fitting

The GAM-models were fitted in a Bayesian setting using the R package R-INLA^44^, available at www.r-inla.org. The range- and precision-parameters of the SPDE were explored in terms of the deviance on test sets (see next section Assessing model fit).

#### Assessing model fit

Cross-validation was used to assess model fit. Rolling window cross-validation with one year ahead in time being left out, from 2017 until 2023. Summary statistics for the model fits were calculated as the average over the training/test sets in the case of DIC and deviance. A receiver operating curve (ROC) and area under the curve (AUC) statistics were computed on the joint set of regions left out over the test sets, encompassing seven test sets for the years 2017-2023 (Fig. 2). Summaries are listed in Table 1 and Table S7.

## Data sharing

The data on *Ae. albopictus* presence is freely available from the ECDC upon request, but it cannot be made available elsewhere. The climate, mechanistic life cycle and human population covariates are included in the Supplementary at bioRxiv, doi 10.1101/2024.09.20.614113.

Code used to fit the spatio-temporal GAM and create figures is included in the Supplementary at bioRxiv, doi 10.1101/2024.09.20.614113. Since the data on *Ae. albopictus* presence cannot be made available, the code runs on a simulated dataset.

## Author contribution statement

Conceptualization: SB, JCS, JR, JW

Data curation: SB, PS

Formal Analysis: SB, PS,

Funding acquisition: JR, JCS, JW

Methodology: SB, JR, JW

Resources: SB, JCS, PS

Software: SB, PS, JW, HS

Supervision: JCS, JR, JW

Validation: SB, PS

Visualization: SB, PS

Writing – original draft: SB, JCS, JR

Writing – review & editing: all authors

