## Supplementary info for "A climate and population dependent diffusion model forecasts the spread of *Aedes Albopictus* mosquitoes in Europe"

### ***Supplement Information***

**Raw data**

The relationship between the total number of years with *Ae. albopictus* presence and human population is shown in Figure S1-S2, with regions with large human population highlighted. Notice that the total number of years with Ae presence is high in regions with high human population compared to in neighbouring regions, as discussed in the main text. Table S1-S6 show empirical *Ae. albopictus* presence in sub-groups of the data, with groups defined based on two-way groupings of covariates. The information contained within these tables are shown in Fig. 6 (main text). This highlighting of interactions between covariates can be used to validate graphically the fitted random effects of the generalized additive model. Similarly, groups based on a five-way grouping of covariates are presented in Table S7. The information contained in this table is displayed in scatter plots in Fig. 5 (main text) and Figure S3.

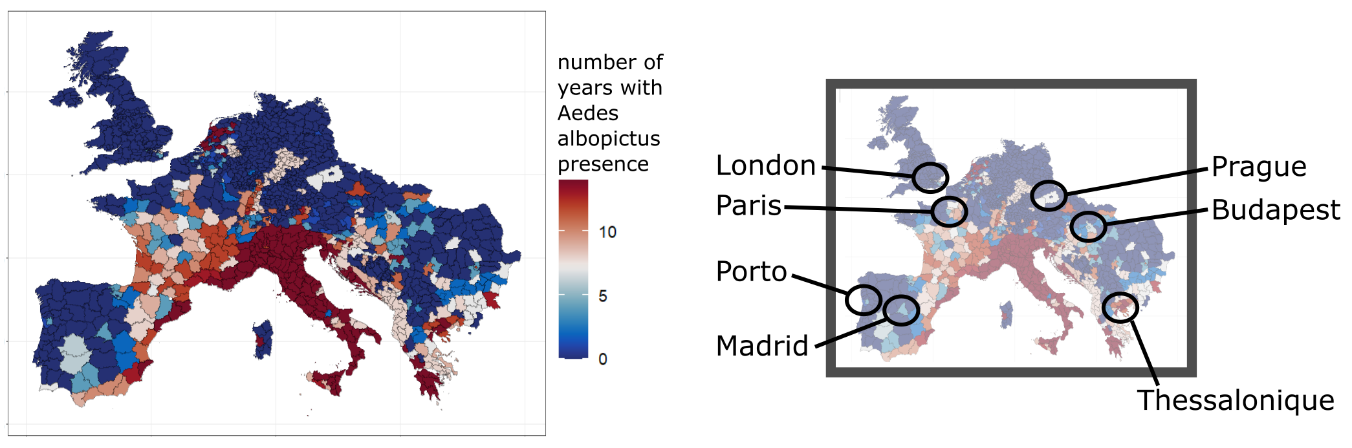

Figure S1: Summary of Ae. albopictus distribution over the years 2010-2020, showing for each region the total number of years with Ae. albopictus presence, i.e., total number of years when the Ae. albopictus status in the region was either “established” or “introduced”. The right figure highlights seven regions around large cities where the total number of years with Ae. albopictus presence was relatively high compared to the region’s neighbours.

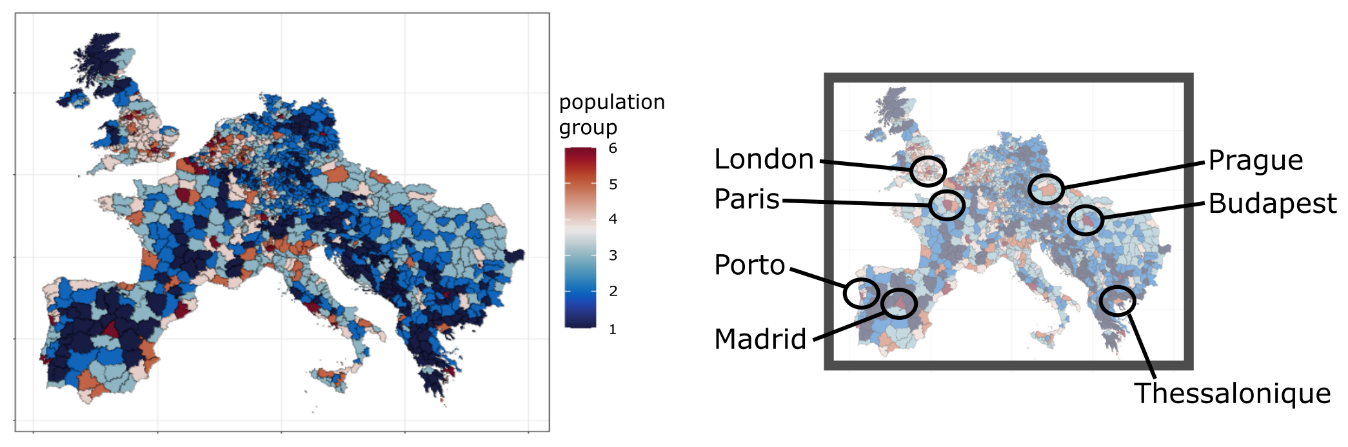

Figure S2: Regions classified into six groups based on their weighted human population: group 1, population between the 0% and 25% quantiles; group 2, population between the 25% and 50% quantiles; group 3, population between the 50% and 75% quantiles; group 4, population between the 75% and 90% quantiles; group 5, population between the 90% and 97.5% quantiles; and group 6, population between the 97.5% and 100% quantiles. The right figure highlights the same regions as in Figure S1.

| Group  index | Group limits,  median temperature (min, max) | Number of locations $(i,t)$ within group | Ratio,  *Ae. albopictus* presence | Group mean,  median temperature |
| --- | --- | --- | --- | --- |
| 1 | (13, 18) | 2180 | 0.037 | 16.6 |
| 2 | (18, 19) | 2281 | 0.20 | 18.5 |
| 3 | (19, 20) | 3007 | 0.20 | 19.5 |
| 4 | (20, 21) | 1807 | 0.34 | 20.4 |
| 5 | (21, 22.5) | 1184 | 0.60 | 21.7 |
| 6 | (22.5, 24) | 1401 | 0.74 | 23.2 |
| 7 | (24, 30) | 1489 | 0.72 | 25.4 |

Table S1: Two-way groups of region-year locations based on median temperature, with group index; the limits that define the group; the total number of locations within the group; the ratio of locations with Ae. albopictus presence in the group; and the group mean of median temperature. All region-year locations that have known Ae. albopictus status (i.e., excluding status “no data”) are included in the Ae. albopictus presence ratio.

| Group index. | Group limits,  median temperature (min, max) | Group limits, minimum temperature | Number of locations $(i,t)$ within group | Ratio,  *Ae. albopictus* presence | Group mean,  median temperature | Group mean,  minimum temperature |
| --- | --- | --- | --- | --- | --- | --- |
| 1-1 | (13, 18) | below 5 | 983 | 0.08 | 16.2 | 2.6 |
| 1-2 | (13, 18) | above 5 | 1197 | 0.0008 | 17.2 | 5.8 |
| 2-1 | (18, 19) | below 5 | 1577 | 0.25 | 18.6 | 1.9 |
| 2-2 | (18, 19) | above 5 | 704 | 0.10 | 18.4 | 5.9 |
| 3-1 | (19, 20) | below 5 | 2664 | 0.21 | 19.5 | 2.7 |
| 3-2 | (19, 20) | above 5 | 343 | 0.15 | 19.5 | 5.6 |
| 4-1 | (20, 21) | below 5 | 1467 | 0.30 | 20.4 | 3.1 |
| 4-2 | (20, 21) | above 5 | 340 | 0.55 | 20.5 | 6.4 |
| 5-1 | (21, 22.5) | below 5 | 573 | 0.44 | 21.7 | 3.6 |
| 5-2 | (21, 22.5) | above 5 | 611 | 0.75 | 21.8 | 6.5 |
| 6-1 | (22.5, 24) | below 5 | 437 | 0.50 | 23.3 | 3.7 |
| 6-2 | (22.5, 24) | above 5 | 964 | 0.84 | 23.2 | 7.9 |
| 7-1 | (24, 30) | below 5 | 284 | 0.36 | 24.6 | 3.5 |
| 7-2 | (24, 30) | above 5 | 1205 | 0.80 | 25.5 | 9.8 |

Table S2: Two-way groups of region-year locations based on median temperature and minimum temperature, with group index; the limits that define the group (median temperature and minimum temperature); the total number of locations within the group; the ratio of locations with Ae. albopictus presence in the group; and group means of median temperature and minimum temperature. All region-year locations that have known Ae. albopictus status (i.e., excluding status “no data”) are included in the Ae. albopictus presence ratio.

| Group index. | Group limits,  median temperature (min, max) | Group limits, median rel. humidity | Number of locations $(i,t)$ within group | Ratio,  *Ae. albopictus* presence | Group mean,  median temperature | Group mean,  median rel. humidity |
| --- | --- | --- | --- | --- | --- | --- |
| 1-1 | (13, 18) | below 59 | 0 | - | - | - |
| 1-2 | (13, 18) | above 59 | 2180 | 0.037 | 16.6 | 78 |
| 2-1 | (18, 19) | below 59 | 0 | - | - | - |
| 2-2 | (18, 19) | above 59 | 2281 | 0.20 | 18.5 | 74 |
| 3-1 | (19, 20) | below 59 | 0 | - | - | - |
| 3-2 | (19, 20) | above 59 | 3007 | 0.20 | 19.5 | 72 |
| 4-1 | (20, 21) | below 59 | 0 | - | - | - |
| 4-2 | (20, 21) | above 59 | 1807 | 0.34 | 20.4 | 71 |
| 5-1 | (21, 22.5) | below 59 | less than 30 | - | - | - |
| 5-2 | (21, 22.5) | above 59 | 1170 | 0.6 | 21.7 | 69 |
| 6-1 | (22.5, 24) | below 59 | 50 | 0.04 | 23.3 | 57 |
| 6-2 | (22.5, 24) | above 59 | 1351 | 0.76 | 23.2 | 68 |
| 7-1 | (24, 30) | below 59 | 379 | 0.65 | 26.4 | 54 |
| 7-2 | (24, 30) | above 59 | 1214 | 0.75 | 25.0 | 65 |

Table S3: Two-way groups of region-year locations based on median temperature and median relative humidity, with group index; the limits that define the group (median temperature and median relative humidity); the total number of locations within the group; the ratio of locations with Ae. albopictus presence in the group; and group means of median temperature and median relative humidity. Groups that have fewer than 30 locations are excluded. All region-year locations that have known Ae. albopictus status (i.e., excluding status “no data”) are included in the Ae. albopictus presence ratio.

| Group index. | Group limits,  median temperature (min, max) | Group limits,  log human population | Number of locations $(i,t)$ within group | Ratio,  *Ae. albopictus* presence | Group mean,  median temperature | Group mean,  log human population |
| --- | --- | --- | --- | --- | --- | --- |
| 1-1 | (13, 18) | below 13 | 1592 | 0.043 | 16.6 | 12.2 |
| 1-2 | (13, 18) | above 13 | 588 | 0.020 | 16.7 | 13.2 |
| 2-1 | (18, 19) | below 13 | 1834 | 0.21 | 18.5 | 12.0 |
| 2-2 | (18, 19) | above 13 | 447 | 0.16 | 18.6 | 13.5 |
| 3-1 | (19, 20) | below 13 | 2415 | 0.19 | 19.5 | 11.9 |
| 3-2 | (19, 20) | above 13 | 592 | 0.26 | 19.6 | 13.5 |
| 4-1 | (20, 21) | below 13 | 1243 | 0.28 | 20.4 | 12.0 |
| 4-2 | (20, 21) | above 13 | 564 | 0.49 | 20.5 | 13.6 |
| 5-1 | (21, 22.5) | below 13 | 873 | 0.56 | 21.8 | 12.0 |
| 5-2 | (21, 22.5) | above 13 | 311 | 0.71 | 21.7 | 13.6 |
| 6-1 | (22.5, 24) | below 13 | 1000 | 0.70 | 23.2 | 12.2 |
| 6-2 | (22.5, 24) | above 13 | 401 | 0.82 | 23.3 | 13.7 |
| 7-1 | (24, 30) | below 13 | 943 | 0.67 | 25.2 | 12.2 |
| 7-2 | (24, 30) | above 13 | 546 | 0.82 | 25.6 | 13.9 |

Table S4: Two-way groups of region-year locations based on median temperature and log human population, with group index; the limits that define the group (median temperature and log human population); the total number of locations within the group; the ratio of locations with Ae. albopictus presence in the group, weighted by human population; and group means of median temperature and log human population. All region-year locations that have known Ae. albopictus status (i.e., excluding status “no data”) are included in the Ae. albopictus presence ratio.

| Group  index | Group limits,  mechanistic life cycle (min, max) | Number of locations $(i,t)$ within group | Ratio,  *Ae. albopictus* presence | Group mean,  mechanistic life cycle |
| --- | --- | --- | --- | --- |
| 1 | (0, 2.0) | 5395 | 0.022 | 1.64 |
| 2 | (2.0, 2.5) | 4981 | 0.038 | 2.20 |
| 3 | (2.5, 3.0) | 1711 | 0.12 | 2.71 |
| 4 | (3.0, 3.5) | 534 | 0.16 | 3.14 |
| 5 | (3.5, 5.4) | 153 | 0.37 | 4.13 |

Table S5: Two-way groups of region-year locations based on the mechanistic life cycle covariate, with group index; the limits that define the group; the total number of locations within the group; the ratio of locations with Ae. albopictus presence in the group; and the group mean of the mechanistic life cycle covariate. Only region-year locations that have known Ae. albopictus status (i.e., excluding status “no data”), and for which the region did not have Aa recorded as present the previous year, are included in the Ae. albopictus presence ratio.

| Group index. | Group limits,  mechanistic life cycle (min, max) | Group limits, proximity | Number of locations $(i,t)$ within group | Ratio,  *Ae. albopictus* presence | Group mean,  mechanistic life cycle | Group mean,  proximity |
| --- | --- | --- | --- | --- | --- | --- |
| 1-1 | (0, 2.0) | below 0.1 | 4468 | 0.015 | 1.64 | 0.01 |
| 1-2 | (0, 2.0) | above 0.1 | 927 | 0.053 | 1.61 | 0.18 |
| 2-1 | (2.0, 2.5) | below 0.1 | 3781 | 0.024 | 2.19 | 0.02 |
| 2-2 | (2.0, 2.5) | above 0.1 | 1200 | 0.076 | 2.24 | 0.20 |
| 3-1 | (2.5, 3.0) | below 0.1 | 1126 | 0.083 | 2.71 | 0.03 |
| 3-2 | (2.5, 3.0) | above 0.1 | 585 | 0.18 | 2.72 | 0.20 |
| 4-1 | (3.0, 3.5) | below 0.1 | 275 | 0.10 | 3.13 | 0.03 |
| 4-2 | (3.0, 3.5) | above 0.1 | 259 | 0.21 | 3.15 | 0.17 |
| 5-1 | (3.5, 5.4) | below 0.1 | 86 | 0.26 | 4.15 | 0.03 |
| 5-2 | (3.5, 5.4) | above 0.1 | 67 | 0.52 | 4.11 | 0.22 |

Table S6: Two-way groups of region-year locations based on the mechanistic life cycle and proximity covariates, with group index; the limits that define the group (mechanistic life cycle and proximity); the total number of locations within the group; the ratio of locations with Ae. albopictus presence in the group; and group means of mechanistic life cycle and proximity covariates. Only region-year locations that have known Ae. albopictus status (i.e., excluding status “no data”), and for which the region did not have Aa recorded as present the previous year, are included in Ae. albopictus presence ratio.

| Group index | Category,  median temperature | Category, minimum temperature | Category, median relative humidity | Category, log human population | Category, proximity | Number of locations $(i,t)$ within group | Ratio,  *Ae. albopictus* presence |
| --- | --- | --- | --- | --- | --- | --- | --- |
| 1 | Low | Low | High | Low | Low | 1723 | 0.03 |
| 2 | Low | Low | High | Low | High | 1833 | 0.36 |
| 3 | Low | Low | High | High | Low | 226 | 0.03 |
| 4 | Low | Low | High | High | High | 222 | 0.47 |
| 5 | Low | High | High | Low | Low | 1230 | 0.003 |
| 6 | Low | High | High | Low | High | 184 | 0.21 |
| 7 | Low | High | High | High | Low | 642 | 0.005 |
| 8 | Low | High | High | High | High | 114 | 0.25 |
| 9 | Mid | Low | High | Low | Low | 2401 | 0.04 |
| 10 | Mid | Low | High | Low | High | 2331 | 0.32 |
| 11 | Mid | Low | High | High | Low | 321 | 0.07 |
| 12 | Mid | Low | High | High | High | 631 | 0.47 |
| 13 | Mid | High | High | Low | Low | 167 | 0.06 |
| 14 | Mid | High | High | Low | High | 267 | 0.45 |
| 15 | Mid | High | High | High | Low | 145 | 0.02 |
| 16 | Mid | High | High | High | High | 247 | 0.53 |
| *17* | *High* | *Low* | *Low* | *Low* | *Low* | *3* | *0* |
| *18* | *High* | *Low* | *Low* | *Low* | *High* | *11* | *0* |
| *19* | *High* | *High* | *Low* | *Low* | *Low* | *120* | *0.05* |
| *20* | *High* | *High* | *Low* | *Low* | *High* | *258* | *0.53* |
| *21* | *High* | *High* | *Low* | *High* | *Low* | *56* | *0.08* |
| *22* | *High* | *High* | *Low* | *High* | *High* | *266* | *0.81* |
| 23 | High | Low | High | Low | Low | 648 | 0.08 |
| 24 | High | Low | High | Low | High | 1312 | 0.50 |
| 25 | High | Low | High | High | Low | 170 | 0.13 |
| 26 | High | Low | High | High | High | 348 | 0.59 |
| 27 | High | High | High | Low | Low | 372 | 0.29 |
| 28 | High | High | High | Low | High | 1630 | 0.91 |
| 29 | High | High | High | High | Low | 116 | 0.45 |
| 30 | High | High | High | High | High | 696 | 0.97 |

Table S7: Five-way groups of region-year locations based on the median temperature, minimum temperature, median relative humidity, log human population and proximity covariates, with group index; the categories that define the group; the total number of locations within the group; and the ratio of locations with Ae. albopictus presence in the group. The limits that define the categories are: median temperature – low [below 19℃], mid [between 19℃ and 21℃], high [above 21℃]; minimum temperature – low [below 5℃], high [above 5℃]; median relative humidity – low [below 59], high [above 59]; log human population – low [below 13], high [above 13]; and proximity – low [below 0.02], high [above 0.02]. The row shading corresponds to the median temperature categories. Groups for which the median relative humidity is low are marked with cursive text. Note that not all combinations of categories are present in the table, due to that the relative humidity is only low in regions with high median temperature. Due to the low number of region-year locations in categories 17-18 (white text), those groups should be disregarded.

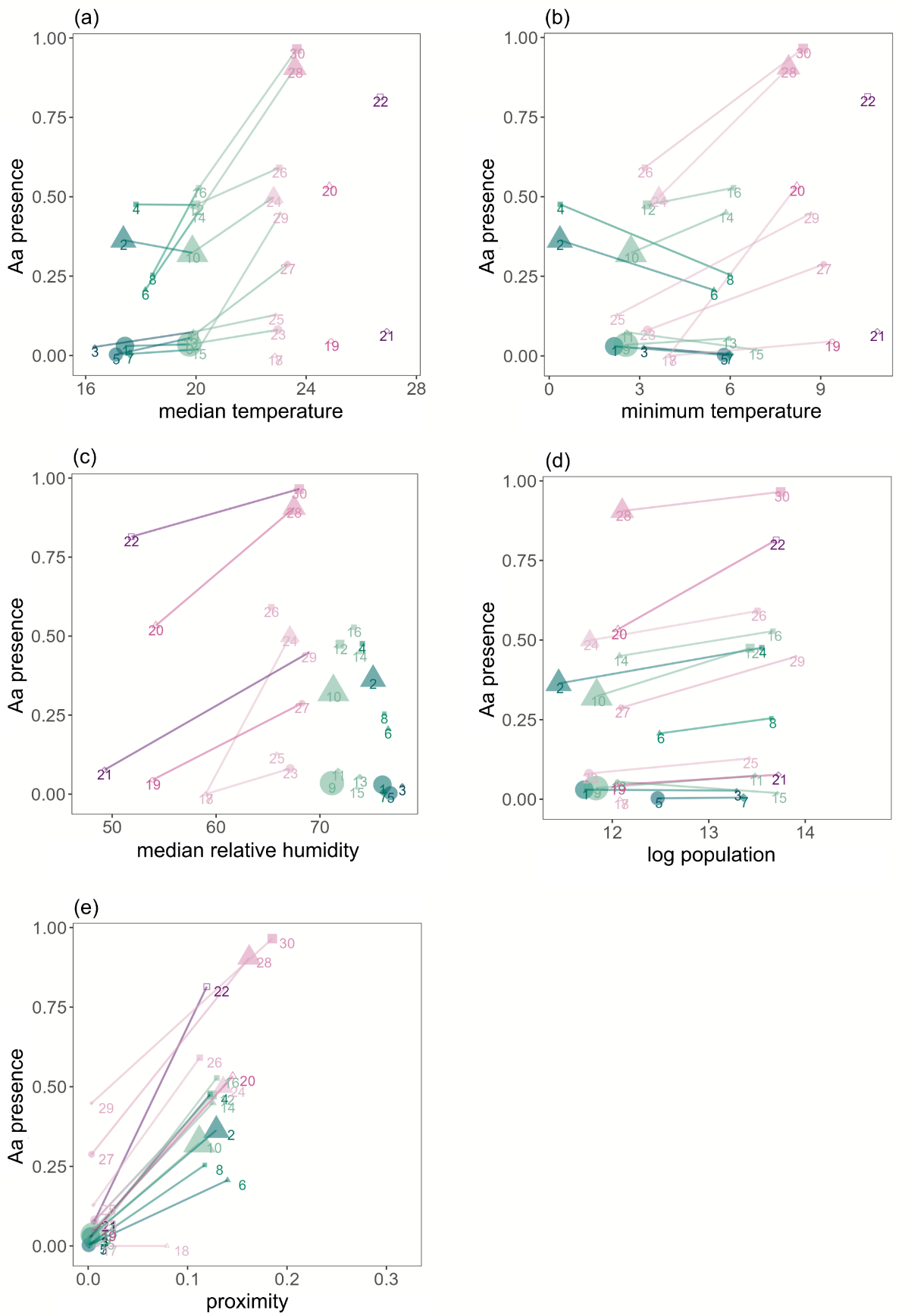

Figure S3: Same plots as in Fig. 5 of the main text showing the five-way grouping in Table S7, with the difference that here links between groups are shown. Links connect groups for which all categories are the same except for (a) median temperature, (b) minimum temperature, (c) median relative humidity, (d) log human population, and (e) proximity. The group coordinates in the plots are the group average of the covariate (x-axis) and the group average Ae. albopictus presence (y-axis). The group index from Table S7 is shown next to each group location. The median temperature category for each group is indicated with colour. Symbols indicate which combination of the log human population and proximity categories a group belong to, with symbols (low human population, low proximity) a circle, ●, (low human population, high proximity) a triangle, ▲, (high human population, low proximity) a diamond ♦, and (high human population, high proximity) a square ■. Groups with median relative humidity are indicated with hollow symbols. For each group, the size of the symbol shown in the scatter plot is proportional to the number of region-year locations that belong to the group.

**Constructing the proximity covariate**

As mentioned in the main text, the proximity covariate is constructed to capture how *Ae. Albopictus* spreads to nearby regions. For the regions with absence, we let the covariate be the conditional expectation of a Gaussian random field where one has observed 1 over all regions with presence at a specific year $t$. In more detail, the method is done as follows: Let $w^{t}$ be a Gaussian field over all regions, i.e., $w^{t}\sim N\left( 0,Q^{-1} \right)$, where $Q$ is the precision matrix (inverse covariance matrix). We split $w^{t}$ into regions with absence $w_{A}^{t}$ and regions with presence $w_{P}^{t}$. Then the precision matrix of $\left( w_{A}^{t},w_{P}^{t} \right)$ is

$$Q=Q=\left( \begin{matrix} Q_{AA} & Q_{AP} \\ Q_{PA} & Q_{PP} \end{matrix} \right).$$

The conditional expectation is given by

$$E\left[ w_{A}^{t} \mid w_{P}^{t}=1 \right]=Q_{AA}^{-1}Q_{AP}1,$$

which we set to bet the proximity covariate for year $t+1.$ This conditional expectation has the same dimension as $w_{A}^{t}$ and corresponds to those regions where *Ae. albopictus* was not present at year $t$.

The normal model is only used to define the conditional expectation, where we fix the values of the GMRF in regions that have recorded *Ae. albopictus* presence to 1. This creates a risk from proximity to regions with *Ae. albopictus* presence, where the risk is diffusing out from regions with *Ae. albopictus* presence.

We chose the precision matrix $Q$ that defines $w^{t}$ as a combination of an intristinc conditional autoregressiv (ICAR) precision matrix $Q_{ICAR}$ and an identity matrix $I$, $Q=0.5Q_{ICAR}+0.5I.$ The ICAR precision matrix is defined on a network $G = (H; E)$, with nodes $H=\left( 1,\ldots,n \right)$, and edges $E$ connecting the nodes. The matrix is written

$$\left( Q_{ICAR} \right)_{ij}=\left\{ \begin{aligned} \sum_{j:i\sim j} \alpha_{ij}, \text{ if} i=j, \\ -\alpha_{ij}, \text{if} i \sim j, \\ 0, \text{otherwise.} \end{aligned} \right.$$

Here $i\sim j$ means that the nodes $i$ and $j$ in the network are joined by an edge in the network. Each region is represented by a node, and we consider two cases for the edges. For the first case, edges connect regions that share a boundary, and each edge $\left( i,j \right)\in E$ has weight $\alpha_{ij}=1$ (Figure S4, left). The second case, which incorporates mobility edges, is described further down.

The proximity precision matrix $Q$ is taken to powers $\theta$, defined as $Q^{\theta}=UD^{\theta}U^{\top}$, where $UDU^{\top}$ is an eigenvalue decomposition of $Q$, with $D$ the diagonal matrix with the eigenvalues. $D^{\theta}$ is obtained by taking the $\theta$-power of the diagonal elements of $D$.

The proximity computed for $t=2020$, with different powers $\theta$, are shown in Figure S5. The diffusing of the risk from proximity to regions with *Ae. albopictus* presence is defined by the graph structure of $G$. One property of this kind of graph diffusion process, is that the proximity is higher in regions that are surrounded by regions with *Ae. albopictus* presence than the proximity in regions that have neighbours with *Ae. albopictus* presence on only one front.

Note that the graph $G$ we use here does not encode geographical distance between regions. Geographical distance could be incorporated in a number of ways, e.g., by letting the weights $\alpha_{ij}$ depend on geographical distance, or by using a precision matrix corresponding to other types of random fields such as an SPDE representation of a Matérn field.

To incorporate mobility, we extend the network between regions that share a boundary to also include links between regions with high mobility (Figure S4, right), computed from the commuting flow of the radiation model as described in the main text. The ICAR precision matrix was defined with weights $\alpha_{proximity}$ for links from proximity (geographical neighbours), and weights $\alpha_{mobility}$ for links from mobility. Different values of $\alpha_{proximity}$ and $\alpha_{mobility}$ were explored.

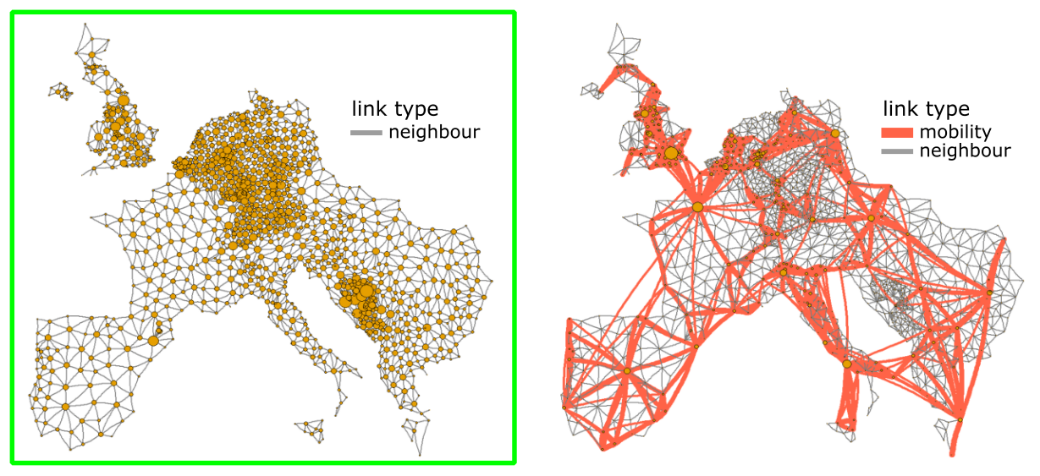

Figure S4: Graphical representation of precision matrices $Q$ corresponding to (left) the CAR model and (right) a combination of the CAR model and a mobility graph. The size of node i (yellow circle) represents the size of the diagonal element $Q_{ii}$. Non-zero diagonal elements $Q_{ij}$ are represented with a link between node i and j, and the width of the link represents the size of $Q_{ij}$. The green box indicates the $Q$ used to create the proximity covariate.

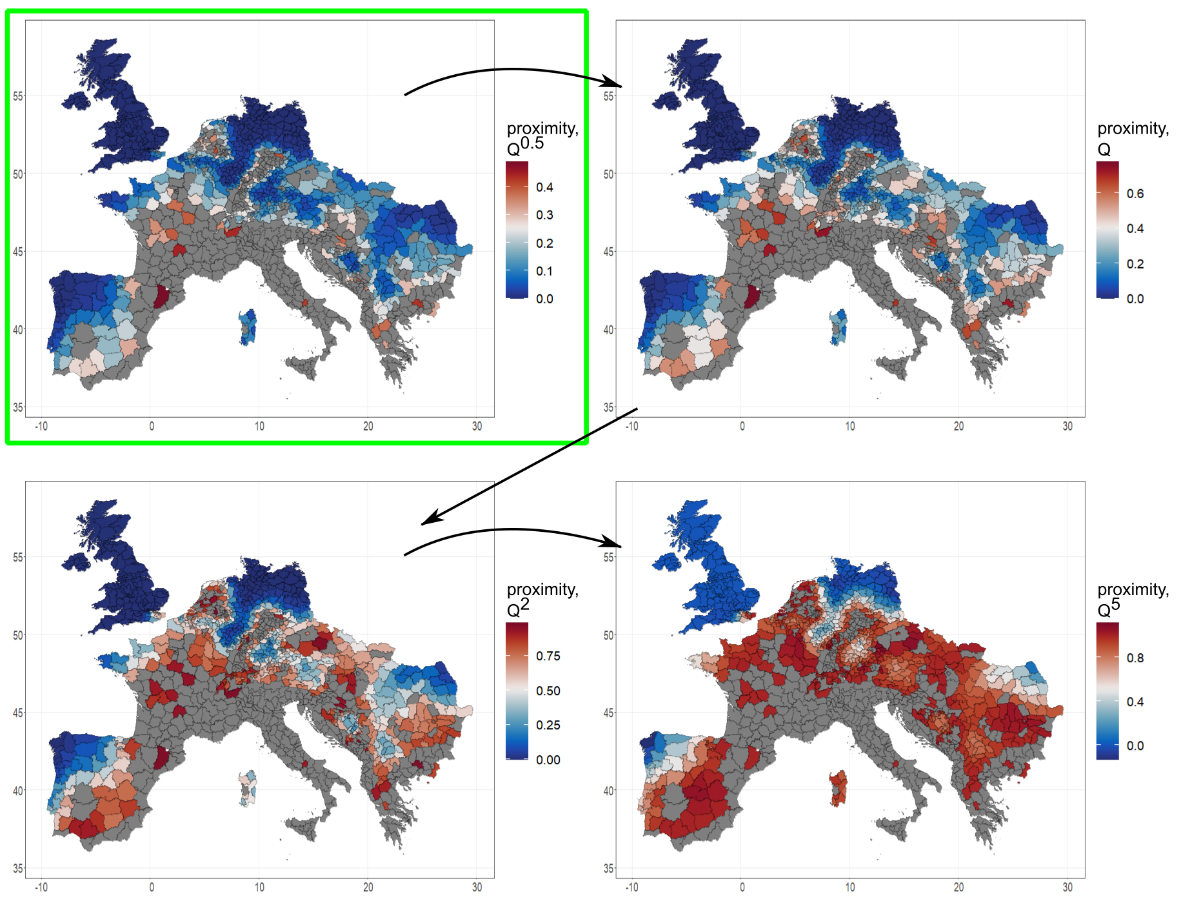

Figure S5: The proximity, computed for year 2020, with regions that had Ae. albopictus presence in 2020 shown in gray, and the proximity shown in color for (top left) $Q^{0.5}$, (top right) $Q$, (bottom left) $Q^{2}$ and (bottom right) $Q^{5}$. The green frame indicates the proximity that was used as a covariate in the GAM model.

#### **Modelling procedure**

**GAM model**

The random effects $f_{h}$ in the GAM model are constructed as follows: The covariate $w_{h,i,t}$ takes a finite number of values, represented by $(1,\ldots,n)$, for which we have a graph $G = (H; E)$ with nodes $H=\left( 1,\ldots,n \right)$, and edges E connecting the nodes. The non-linear random effect is defined as a Gaussian vector $f_{h}\sim N\left( 0,\Sigma\right)$ with a sparse inverse covariance matrix, a Gaussian Markov random field (GMRF), where the covariance matrix $\Sigma$ is determined by the graph G. We use either an intrinsic conditional autoregression (ICAR) or random walk covariance structure on the graph. See Rue and Held [2005] for details about these covariance models. The graph representation gives flexibility in modelling dependence within and between covariates. The sparsity of the inverse covariance matrix in turn allows for efficient model fitting. $f_{s}$ is an SPDE approximation of a continuous Matérn Gaussian random field. When there is a time-dependence, time and space is modelled as a separable (Kronecker) covariance with an IID dependence in time.

Continuous covariates are binned by first dividing the covariate into evenly spaced intervals. Bins with few locations $(i,t)$ were merged in a second step. In cases where it is reasonable to expect the effect to be increasing with increasing values of the covariate, the number of bins were reduced in a final step until the model fit produced a mainly increasing random effect. The bins limits are shown in histograms beneath each fitted random effect in Fig. 5 (main text).

Covariates need to be defined for all observations. The proximity covariate, however, is only defined for locations $(i,t)$ for which $y_{i,t-1}=0$, with $t>2010$. For locations that have no defined proximity, we set the proximity to dummy values, one dummy value for locations with $t = 2010$, and one dummy value for locations such that $y_{i,t-1}=1$, $t>2010$. When creating the proximity random effect, we give the dummy values each a separate bin. The nodes corresponding to the dummy proximity values are then disconnected from the nodes corresponding to the binned non-dummy proximity values in the graph G defining the random effect.

For the minimum temperature covariate, *Ae. albopictus* presence increases with increasing minimum temperature, but only when the median temperature is relatively high (Fig. 6(b), main text; Table S2). We include the minimum temperature conditioned on median temperature, with median temperature corresponding to the lowest two bins of the median temperature covariate excluded (median temperature < 19.3°C, see Fig. 5(a), main text). The minimum temperature is set to a dummy value, for regions $i$ for which the median temperature < 19.3°C, and the dummy value is disconnected from the non-dummy values when creating the random effect graph.

The proximity-, previous year- and first year-covariates are included in both GAM models (the mechanistic life cycle GAM model and the climate and population GAM model). They are the only covariates that differ between years. The previous year- and first year-covariates are binary variables. Comparing the fitted random effects for the more complex proximity-covariate, we see that they are almost identical for the two GAM models (Figure S6). Similarly, the fitted random effects for other covariates are robust to the model choice, i.e., to the choice of other covariates included in the model.

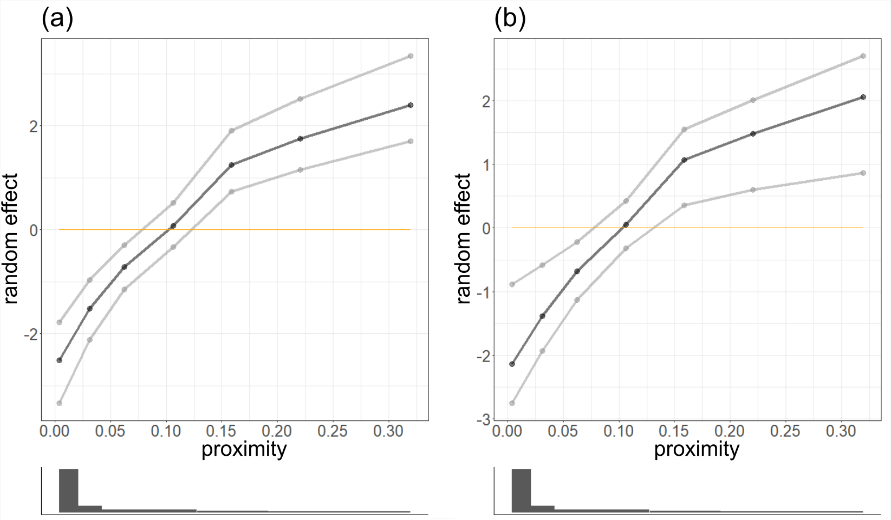

Figure S6: Fitted random effects for the proximity covariate, for (a) the mechanistic life cycle GAM model and (b) the climate and population GAM model. Black lines indicate the random effect, and gray lines indicate the lower and upper limits of 95% credible intervals.

###### Priors

For univariate random effects with a connected graph, we used penalized complexity priors^1^ for the ‘rw1’ model with INLA recommended parameters in the case of a binary response variable (P$\left( \text{log}\left( 1/\sigma^{2} \right)>0.5 \right)=0.01$, $\sigma=$ standard deviation of the random walk increments). For random effects defined on a disconnected graph, i.e., for the proximity covariate and the minimum temperature conditioned on high maximum temperature, we used the ‘besag’ model with the same penalized complexity prior as for ‘rw1’ models. The standard prior for INLAs was used for the spatio-temporal SPDE. The model with a short-range SPDE was implemented using the ‘generic’ model, which is defined by the precision matrix Q corresponding to the SPDE model evaluated at region locations. In this case, the standard prior in INLA for the ‘generic’ model was used, which is the penalized complexity prior with the same parameters used for the ‘rw1’ model, which here is P$\left( \text{log}\left( \tau\right)>0.5 \right)=0.01$, $\tau=$ scaling factor for the precision matrix. Alternatively, when using the free range-SPDE, the standard priors for the SPDE model inla.spde2.matern were used. These are i.i.d. normal priors on log-scale for two parameters related to the range and variance of the SPDE.

###### Model subsets

Simpler models using subsets of the covariates in the main models, i.e., in the mechanistic life cycle GAM model and the climate and population GAM model, are shown in Table S8. The main models perform similar on the cross-validation test sets, as is also shown in Table 1 (main text). It is clear that the predictive performance of the simpler models are reduced compared to the main models. The main contributors to the predictive performance seem to be the mechanistic life cycle-, the median temperature-, and the proximity-covariate.

|  | Deviance, training set | Deviance, test set | AUC,  test set |
| --- | --- | --- | --- |
| Mechanistic life cycle GAM model:  * mechanistic life cycle covariate (RE) * proximity covariate (RE)  * *regular fixed effects* | 1798 | **190** | 0.79 |
| Climate and population GAM model:  * all climate covariates (RE)  (median temp.; min temp. conditioned on high median temp; median rel. humidity)  * log human population (RE) * proximity covariate (RE)  * *regular fixed effects* | **1596** | 191 | **0.80** |
| Simple GAM model:  * proximity covariate (RE)  * *regular fixed effects* | 1899 | 385 | 0.75 |
| Simple GAM model:  * mechanistic life cycle covariate (RE)  * *regular fixed effects* | 1772 | 337 | 0.68 |
| Simple GAM model:  * climate covariate (RE) (median temp.)  * *regular fixed effects* | 1805 | 339 | 0.74 |
| Simple GAM model:  * climate covariate (RE) (median temp.)  * log human population (RE)  * *regular fixed effects* | 1787 | 333 | 0.74 |
| Simple GAM model:  * all climate covariates (RE)  (median temp.; min temp. conditioned on high median temp; median rel. humidity)  * *regular fixed effects* | 1740 | 325 | 0.74 |

Table S8: Comparison of model fit for the mechanistic life cycle GAM model, the climate and human population GAM model, and five simpler models with subset of the covariates of the two main models. All GAM models include the regular fixed effects ‘presence previous year’ and ‘first year’. The model fit summaries are given as averages over the 7 cross-validation fits (withholding one year ahead for each of the years 2017-2023). The best value in each column is indicated with bold font.

**Controlling the SPDE range to improve model predictions**

We found that manually setting the range-parameter $r$ of the SPDE improve the test set-deviance in a consistent way across both test sets and choice of GAM model (Figure S7 (a),(d); Table S9-S10). Test set-deviance was minimized when a fixed $r=3$ was used. The posterior mean of $r$ was around $0.5$, however, when including $r$ as a free model parameter, with the fitted model having substantially higher test set-deviance.

Comparing the model fit for the climate and population GAM model with a shorter range- and a free range-SPDE (Figure S-S9), the fitted SPDE has a considerably smaller range for the model with fixed $r=3$ (shorter range-SPDE,

Figure S (a)-(d)) compared to when $r$ is included as a free model parameter, in which case the posterior estimate is $r=0.47$ (free range-SPDE,

Figure S (e)-(h)). This is consistent with the SPDE prior, for which higher values of $r$ correspond to a shorter range SPDE field.

The fitted SPDE is less correlated with the data for higher values of $r$ (Figure S7 (b),(e)). The posterior mean of the precision for the median temperature (Figure S7 (c)) and for the mechanistic life cycle covariate (Figure S7 (f)) decrease with increasing values of $r$, indicating that the fitted variance of the random effects for these covariates increase with increasing values of $r$. This leads to fitted random effects with larger values f(w), and thus the contribution to the log-odds coming from the covariates w will be larger.

The fitted random effect f(w), for the median temperature-covariate w, is shown in

Figure S ((c)-(d), (g)-(h)). It is considerably larger for the model with a shorter range-SPDE ($r=3$) compared to the standard free range SPDE. The sum of the fitted random effect of the SPDE and median temperature-covariate show that the contribution from the median temperature-covariate is barely visible in the geographical plot for the free range-SPDE model (Figure S (b)), whereas it can clearly be seen in the corresponding plot for the short range-SPDE model (Figure S (a)).

This is not only true for the median temperature covariate: fitted random effects for all covariates are consistently larger when using a shorter range-SPDE. This trend likely explains the improved fit on the test set with shorter range-SPDE. The posterior of the SPDE is essentially zero for the test set in the region / year-setup, and so the SPDE does not help model fit on the test set. Thus, the larger portion of the total variation in the data that is explained by the covariates, the better for the fit on the test set.

Setting aside the point that the model fit on test sets is considerably improved when using a short range-SPDE, there is also a decrease in correlation between the SPDE and the data with decreasing range. The fitted SPDE seem to be more conservative when the SPDE prior has a shorter range, which could explain the difference in correlation with the data.

As seen in the histograms in

Figure S ((b),(f)), the fitted SPDE values are more clustered around zero for the model with a shorter range-SPDE prior compared to the model fitted with a free range-SPDE prior. This is also clearly seen in the geographical plots of the fitted SPDE (

Figure S (a),(e)). Another aspect of the more conservative nature of the short range-SPDE is that the values of the fitted SPDE are higher in regions where the covariates do not explain *Ae. albopictus* presence well compared to in regions where the covariates do a better job (

Figure S (a)). This effect is not as clear for the free range-SPDE model (

Figure S (e)). The regions indicated by the black arrow around the northern part of Italy (

Figure S (a),(c)) are such regions where covariates do not explain the *Ae. albopictus* presence well. These regions are deemed to be outlier regions, and the fitted values of the short range-SPDE in these outlier regions are considerably higher than in nearby regions with *Ae. albopictus* presence. Thus it is clear that the short range-SPDE model compensates for the lack of fit by covariates in these outlier regions. The outlier regions cannot however be distinguished by fitted SPDE-values for the model with a free range-SPDE (

Figure S (e)). One interpretation of these results is that the free range-SPDE compensates for the lack of fit in outlier regions just as the short range-SPDE does, but the fitted free range-SPDE also compensates in regions where the covariates explain the data well. This leads to a better fit for the model with a free range-SPDE on training data, however, more of the variation in the data is being explained by the SPDE-component and less of the variation is being explained by the covariates-component with the free range-SPDE. The SPDE-component, however, is not transferred to new data whereas the covariate-components are, and thus the model with the free range-SPDE has a worse performance on test data.

| Range parameter $r$ | DIC | Deviance, training | Deviance,  test | Correlation,  SPDE with data | Precision,  median temperature covariate |
| --- | --- | --- | --- | --- | --- |
| Free (0.45) | 2639 | 1347 | 271 | 0.31 | 6.1 |
| 0.5 | 2624 | 1331 | 265 | 0.29 | 9.9 |
| 1.0 | 3386 | 1440 | 285 | 0.25 | 5.6 |
| 1.5 | 3396 | 1549 | 253 | 0.09 | 3.5 |
| 2.0 | 3995 | 1659 | 226 | -0.01 | 3.5 |
| 2.5 | 4074 | 1565 | 220 | -0.09 | 4.0 |
| **3.0** | **3529** | **1525** | **184** | **-0.13** | **3.7** |
| 3.5 | 3385 | 1453 | 184 | -0.11 | 3.7 |
| 4.0 | 3193 | 1408 | 184 | -0.10 | 3.0 |

*Table S9: Results from fitting the climate covariate GAM model, for each of the range parameters shown in Figure S7. Results in this table are averaged over each of the cross validation-sets. Results in the first row, for models fitted with a free range-parameter, are not shown in Figure S7. We here present the range parameter; the DIC; the deviance on training set; the deviance on test set; the correlation of fitted SPDE and the data; and the fitted precision of the median temperature covariate. The row in bold indicates the setup used for the model fits presented throughout the paper.*

| Range parameter $r$ | DIC | Deviance, training | Deviance,  test | Correlation,  SPDE with data | Precision,  mechanistic life cycle covariate |
| --- | --- | --- | --- | --- | --- |
| Free (0.44) | 2727 | 1401 | 291 | 0.34 | 3.4 |
| 0.5 | 2719 | 1361 | 280 | 0.30 | 3.8 |
| 1.0 | 3672 | 1526 | 260 | 0.14 | 2.5 |
| 1.5 | 4645 | 1732 | 239 | 0.004 | 2.2 |
| 2.0 | 4929 | 1841 | 211 | -0.11 | 1.9 |
| 2.5 | 4505 | 1808 | 195 | -0.15 | 1.9 |
| **3.0** | **3869** | **1757** | **187** | **-0.16** | **1.8** |
| 3.5 | 3678 | 1616 | 190 | -0.14 | 1.7 |
| 4.0 | 3431 | 1502 | 191 | -0.13 | 1.7 |

Table S10: Results from fitting the mechanistic life cycle GAM model, with the same structure as in Table S9. The last column shows the fitted precision of the mechanistic life cycle covariate. The row in bold indicates the setup used for the model fits presented throughout the paper.

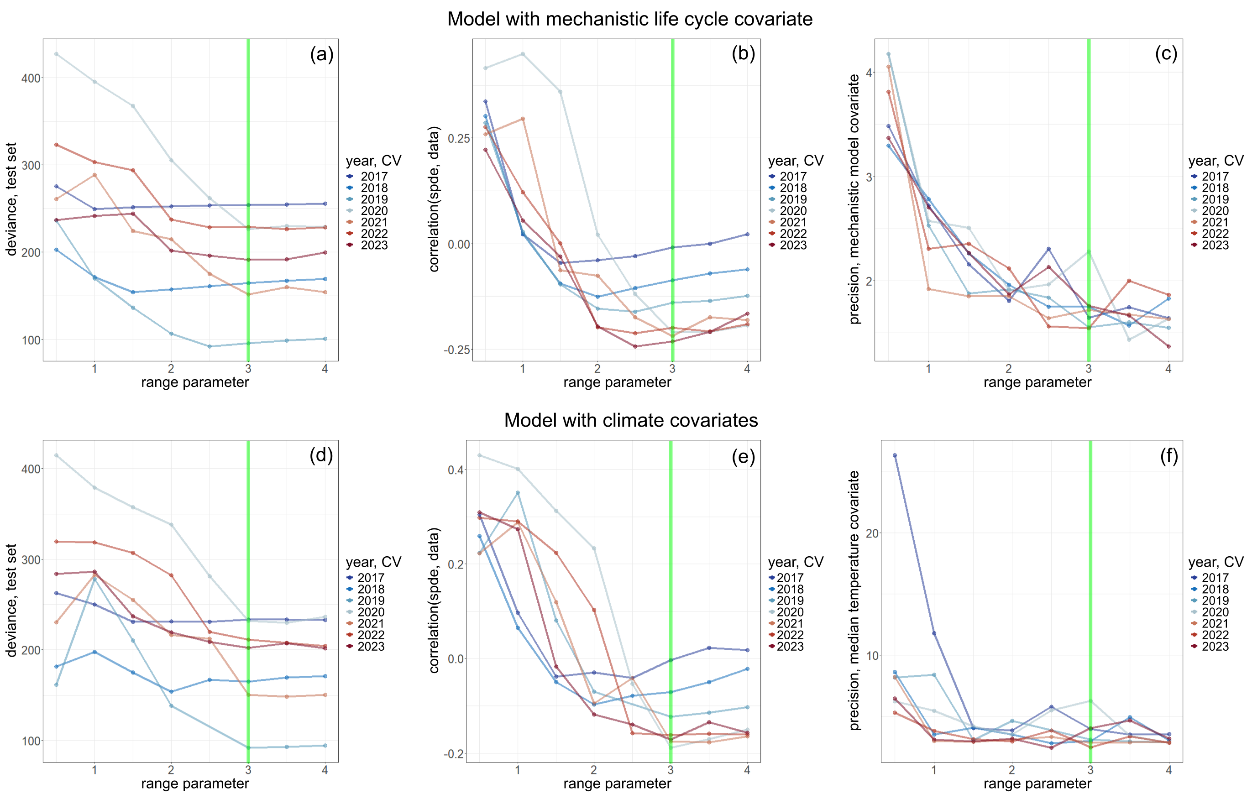

Figure S7: Results from fitting (top) the mechanistic life cycle GAM model and (bottom) the climate and population GAM model. Each model has been fitted for range parameters $r$ from 0.5 up to 4, with a rolling window-cross validation starting from leaving 2017 up to 2023. The vertical green line at range parameter $r=3$ indicates the range parameter that is used throughout the paper. For each model fit is shown the deviance on the test set (left column); the correlation between the fitted SPDE and the data (middle column); and fitted precision-hyper parameters of the mechanistic life cycle covariate (c) and of median temperature (f).

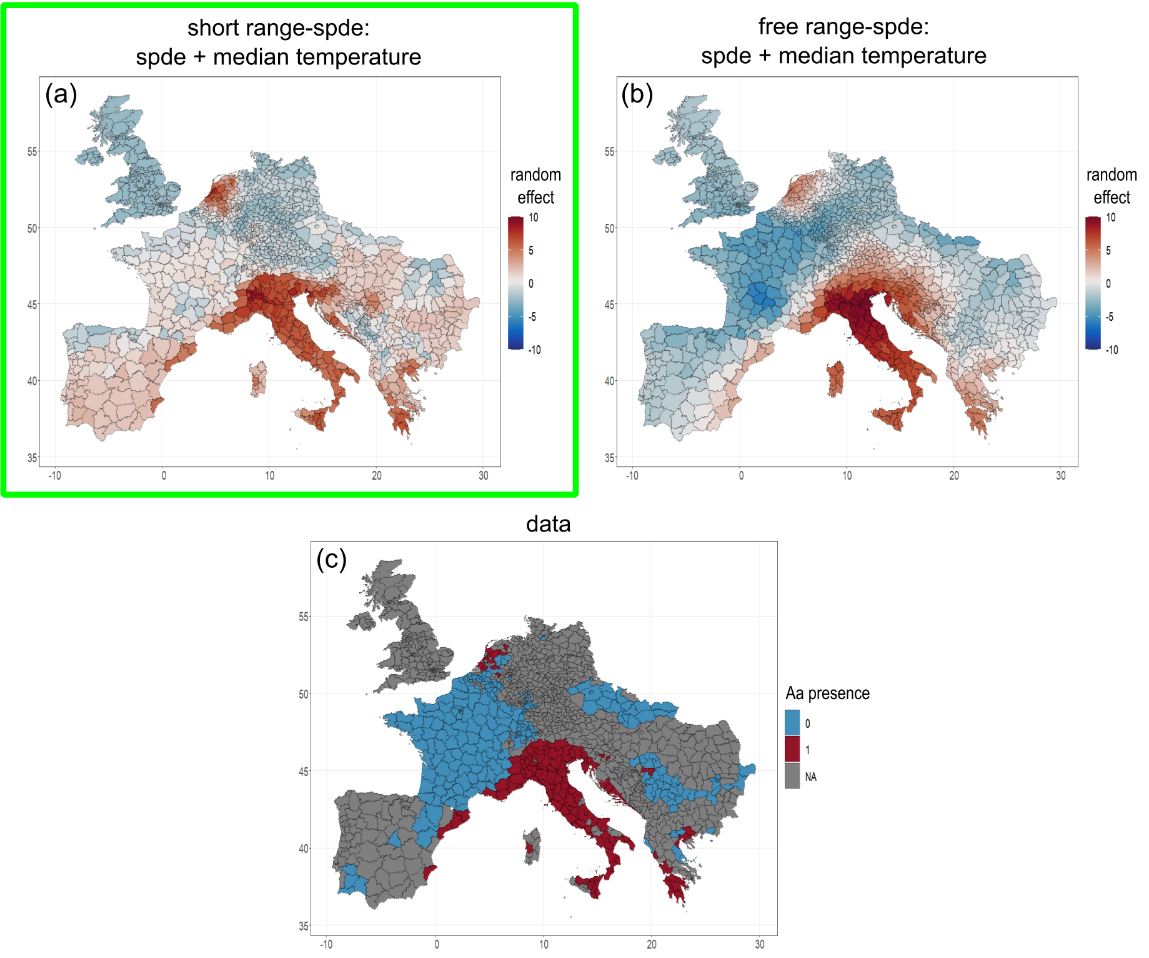

Figure S8: Comparison of model fits for the climate and population GAM model, shown for year 2010, with (a) a fixed $r=3$, so a short range-SPDE, and (b) $r$ as a free parameter, which gives an SPDE with long range. The posterior mean with a free parameter is $r=0.47$. The green frame indicates the setup ($r=3)$used throughout the paper. Images (a) and (b) show the sum of the fitted SPDE- and median temperature random effect. Figure (c) shows the data $y_{it}$, for all regions (regions in gray are regions with status “no data”). The green frame indicates the fit with range parameter $r=3$, i.e., the range parameter used throughout the paper.

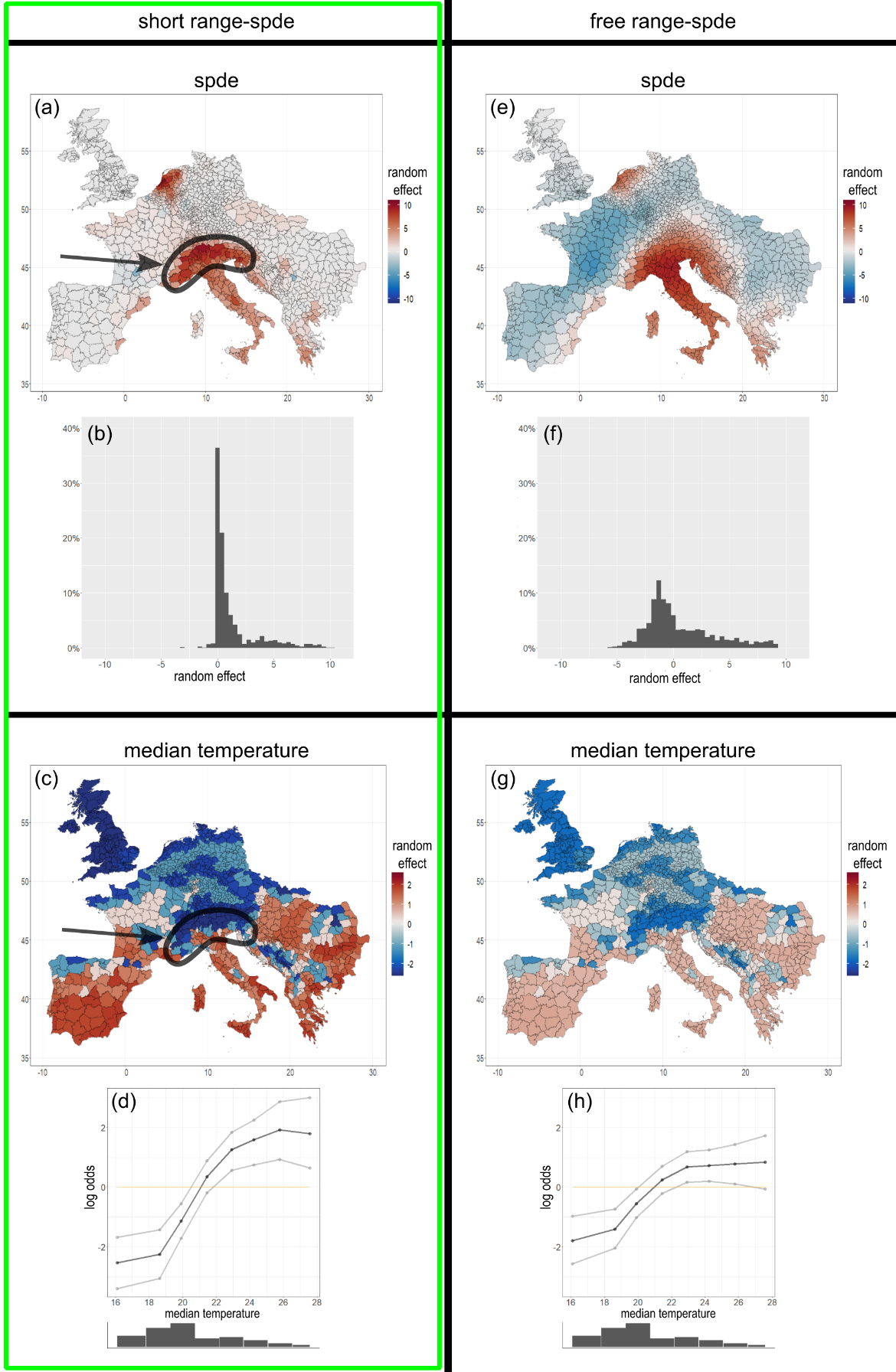

Figure S9: The same two model fits for the climate and population GAM model as shown in

Figure S, for year 2010. The difference between the two columns is that the model presented in the left column was fitted with the short range-SPDE ($r=3$) and the model presented in the right column was fitted with the free range-SPDE. The range parameter $r=3$ is used throughout the paper and is indicated with the green frame. The fitted SPDE-random effect for 2010 is shown geographically (top row) and in a histogram (second row). The fitted median temperature-random effect is shown geographically (third row) and on functional form (bottom row).

**Mechanistic life cycle model details**

***Ae. albopictus* life cycle**

The life cycle of *Ae. albopictus* mosquito show a similar structure in comparison to other mosquito species^2,3^. Adult female mosquitoes deposit eggs predominantly in artificial containers^3^ within water bodies, facilitated by favourable climatic conditions and suitable breeding sites. Subsequently, the eggs undergo hatching induced by rainfall events, and transforms into larvae following a desiccation period, or they endure adverse climatic conditions, such as winter in temperate regions, by entering a state of diapause (dormancy). Larvae then feed on nutrients from their aquatic breeding sites and transform into pupa after undergoing four stages of development^2^. Upon emergence, the sex ratio of newly eclosed females on surface of water is nearly 50:50 and rather than immediately searching for a host, these newly emerged females spent some time in resting stage. The matured adult female’s mosquitoes actively look for suitable mating partner and subsequent to mating, the females disperse to seek a host. This dispersal involves long-distance movements and entails the inherent risk of encountering host defence responses^4^. After a blood meal, the females seek for a suitable aquatic breeding site to oviposit and lay eggs in clusters within few days. The egg laying process is contingent upon the availability of daylight or photoperiod and temperature and if requisite thresholds are not met, female initiate the laying of diapausing eggs which diapause throughout the winter stage until temperature and photoperiod reaches the critical threshold again in spring. The transitions between various stages in *Ae. albopictus* population dynamics system is affected by temperature, precipitation, daylight length and human population density.

**Population dynamics model of *Ae. albopictus***

The population dynamics *Ae. albopictus* is characterized by a stage – structured, climate - data driven dynamical model based on the life cycle of *Ae. albopictus.* Building upon methodologies employed in prior studies^2-8^, we proposes a six stage differential equation model. The model comprises 3 aquatic stage as Egg ($E$), Diapausing egg ($E_{dia}$), Juvenile stage (Larval stage + Pupal stage), and 3 aerial stages as Emerging adult ($A_{em}),$ Blood fed adults ($A_{b})$and Ovipositing adults ($A_{o})$of mosquito life cycle. For simplification we have combined resting and mating stage of emerging mosquito into stage of emerging adults. The juvenile stage also integrates the Larval and Pupal stages into a single class due to constraints in data availability for parameterization of these stages. Moreover, the split of these stages did not yield an improvement in the model fit to presence data^5^. The resultant system of differential equations with parameter definition given in Table S11, where dot notations is for time derivatives can be written as:

$$\dot{E}=\beta\left( T \right) \left( 1 -\omega\left( \bar{T},\bar{S} \right) \right)M-{Q\left( W,P \right)\delta_{E}\left( T \right)E\text{ }- m}_{E}\left( T \right)E$$

$\dot{E_{dia}}=\delta_{E}\left( T \right)\omega\left( \bar{T},\bar{S} \right)E_{dia}-\sigma\left( \bar{T},S \right) Q\left( W,P \right){\delta_{E}\left( T \right)E}_{dia}- m_{E_{d}}\left( T \right)$ $E_{dia}$

$$\dot{J} = Q\left( W,P \right)\delta_{E}\left( T \right)E+\sigma\left( \bar{T},S \right) Q\left( W,P \right) {\delta_{E}\left( T \right)E}_{dia}-\delta_{J}\left( T \right)J-\left( \frac{J}{K_{J}\left( W,P \right)}+ m_{J}\left( T \right) \right) J$$

$$\dot{A_{em}}={\frac{1}{2}\delta}_{J}\left( T \right) J-\delta_{A_{em}}\left( T \right)A_{em}- m_{A}\left( T \right)A_{em}$$

$$\dot{A_{b}}=\delta_{A_{em}}\left( T \right)A_{em}+{\delta_{A_{o}}A}_{o} -{(m}_{A}\left( T \right)+r+ \delta_{A_{b}}\left( T \right)) A_{b}$$

$$\dot{A_{o}}=\delta_{A_{b}}\left( T \right)A_{b}-{(m}_{A}\left( T \right)+r + \delta_{A_{o}})A_{o}$$

where $T, S,W \mathrm{and} P$represents temperature in $℃$, photoperiod (day-light length) in hours, precipitation (mm) and human population density (people/$km^{2})$ respectively. Diurnal temperature variation throughout the day is leveraged to simulate development and mortality rates and the bar over data variables ( $\bar{T},\bar{S})$represents the mean taken over previous seven days. Precipitation in other studies^9,10^ was also included as accumulation over past seven days but the rate of evaporation reduces the contribution of previous precipitation to negligible amount^11^. Therefore, we decided to use the present-day total precipitation. The daylight model proposed by Forsythe^12^ is used to calculate the photoperiod ($S$) assuming that center of sun is even with the horizon. The model uses the earth’s revolution angle, latitude, day of year and Schoolfield’s coefficient to give daylength in hours. Diurnal temperature variation throughout the day is computed using the model by Dewit^13^, where time of sunrise is approximated using the sun declination angle^12^ which itself is contingent on the Earth's revolution angle from the day of the year. Human density also affect the egg hatching rate in a manner such that hatching is increased in areas where human density is greater^5^ than 500 people/$km^{2}$. The carrying capacity of juvenile stage ($K_{L}\left( W,P \right))$ also increase linearly with human density and rainfall accumulation but multiplied with a scaling factor ($\lambda$) to ensure that maximum mosquito abundance in a hectare area do not exceed the order ${10}^{6}$^2,4,5^. For a comprehensive understanding of model parameters and functions, please refer to Table S11.

**Data and model simulation**

Historical climate data for the year 1995 – 2019 were obtained from the Copernicus Climate Data Store (<https://cds.climate.copernicus.eu/>). Near-surface air temperature ($℃$) and total precipitation (mm) are obtained from the fifth generation of the European Centre for Medium-Range Weather Forecasts atmospheric reanalyses (ERA5). The grided ERA5-land hourly climate dataset (<https://cds.climate.copernicus.eu/cdsapp#!/dataset/reanalysis-era5-land?tab=overview>) is converted to daily estimates at a resolution 0.25 × 0.25 degree which approximately covers an area of 25 $km^{2}.$ Daily mean, minimum, maximum temperature and daily total rainfall were extracted from hourly ERA5-land dataset. All temporal and spatial data aggregation process was conducted by utilizing the Climate Data Operators (CDO) software^14^. The human population density data and population count is obtained from GPWv4 data set^15^ for the year 2000, 2005, 2010, 2015 and 2020 and GPWv3^16^ data set is used to estimate human population density of year 1995. The population density of rest of the years in between 1995 – 2019 is retrieved using linear interpolation. The models compute the population-density of mosquitoes independently in each grid cell with the assumption of well-mixed mosquito populations within 0.25° × 0.25° latitude longitude grid-cells. Daily time steps simulations are used for solving the model and then we aggregated the output to monthly time steps and then further aggregate it to yearly values. To account for diurnal temperature variation, simulation for a single day is divided into 100 time steps^5^ according to numerical solver (deSolve^17^ package in R) as 0.14, 0.19 ------ 24.00 and temperature at each of these hours is used to simulate the development and mortality rates for one day. For more details on simulation of model and an simple example model simulation code (in octave v4.2.1), one can refer to Metelmann et al. ^5,18^. Lastly, spatial aggregation at the NUTS 3 (Europe) level was executed in R version 4.1 utilizing the rasterR^19^ package.

Table S11 - Climate sensitive parameter description and functions with references where applicable

| **Description (parameter)** | **Function or parameter value** | **unit** | **Ref.** |
| --- | --- | --- | --- |
| Egg per female per day ($\beta\left( T \right)$) | $\beta\left( T \right)=max\{-0.0163+1.2897T-15.837T^{2},0\}$ | $\frac{1}{day}$ | ^13^ |
| Diapausing egg proportion ($\omega\left( \underline{T},\underline{S} \right))$ | $\omega\left( \underline{T},\underline{S} \right)=0.5 \times f\left( \underline{S}-{Š}_{a} \right) f\left( -\underline{T}- {Ť}_{D} \right)$ | NA | ^3,13^ |
| Spring hatching rate ($\sigma\left( \underline{T}, S \right)$) | $\sigma\left( \underline{T},S \right)=0.1 \times f\left( Ť-\underline{T} \right)f\left( -{Š}_{s}-S \right)$ | $\frac{1}{day}$ | ^3,12^ |
| Egg development rate ($\delta_{E}(T)$) | $\delta_{E}(T)=0.5070\left( -\left( \frac{\left( T-30.85 \right)}{12.82} \right)^{2} \right)$ | $\frac{1}{day}$ | ^12,13^ |
| Juvenile development rate ($\delta_{J}(T)$) | $\delta_{J}(T)= \frac{1}{(0.08T^{2}-4.89T+83.85)}$ | $\frac{1}{day}$ | ^3^ |
| Emerging adult development rate ($\delta_{Aem}(T)$) | $\delta_{Aem}(T)=\frac{1}{(0.069T^{2}-3.574T+50.1)}$ | $\frac{1}{day}$ | ^3^ |
| Blood fed adult development rate ($\delta_{Ab}(T)$) | $\delta_{Ab}\left( T \right)=\frac{T-10}{77}+0.2$ | $\frac{1}{day}$ | ^12^ |
| Subsequent blood meal rates after oviposition ($\delta_{o}$) | $\delta_{o}=0.2$ | $\frac{1}{day}$ | ^12^ |
| Egg mortaliy rate ($m_{E}(T))$  Fraction of hatched eggs depending on precipitation | $m_{E}\left( T \right)=-ln\left( 0.955\left( -{0.5\left( \frac{\left( T-18.8 \right)}{21.53} \right)}^{6} \right) \right)$ | $\frac{1}{day}$ | ^3^ |
| Juvenile mortality rate ($m_{J}(T))$  Diapause condition | $m_{J}\left( T \right)=-ln\left( 0.977\left( -{0.5\left( \frac{\left( T-21.8 \right)}{16.6} \right)}^{6} \right) \right)$ | $\frac{1}{day}$ | ^3^ |
| Adult mortality rate ($m_{A}(T))$ | $m_{A}\left( T_{mean} \right)=-ln\left( 0.677\left( -{0.5\left( \frac{\left( {(T}_{mean}-20.9 \right)}{13.2} \right)}^{6} \right){{(T}_{mean})}^{0.1} \right)$ | $\frac{1}{day}$ | ^3^ |
| Diapausing egg mortality rate ($m_{Ed}\left( T \right))$ | $m_{Ed}\left( T \right)=m_{E}\left( T \right)=-ln\left( 0.955\left( -{0.5\left( \frac{\left( T-18.8 \right)}{21.53} \right)}^{6} \right) \right)$ | $\frac{1}{day}$ | ^3^ |
| Hatching fraction depending in human density and rainfall ($Q\left( W,P \right))$ | $Q\left( W,P \right)=0.8\times\frac{\left( 2.5expexp \left( -0.05\left( {W(t)-8)}^{2} \right) \right) \right)}{expexp \left( -0.05\left( {W(t)-8)}^{2} \right) \right) +1.5}$ + $\frac{0.01}{0.01+ expexp \left( -0.01P(t) \right)}$ | NA | ^3^ |
| Juvenile carrying capacity ${(K}_{L}\left( W,P \right))$ | ${(K}_{L}\left( W,P \right)=\lambda\frac{0.1}{1-{0.9}^{t}}\sum_{x=1}^{t} {0.9}^{\left( t-x \right)}$  $(\alpha_{rain}W\left( x \right)+\alpha_{dens}P\left( x \right))$ | NA | ^3^ |
| Mortality rate associated with long distance travel and search behaviour ($r)$ | $r=0.8$ | $\frac{1}{day}$ | ^12^ |
| Critical day-light length in autumn | ${Š}_{a}=10.058+0.08965 \times Latitude(degrees)$ | hours | ^3^ |
| Critical day-light length in spring | ${Š}_{s}=11.25$ | hours | ^3^ |
| Critical weakly avg. temperature in spring | $Ť=$11.0 | $℃$ | ^3^ |
| Critical diapause temperature | ${Ť}_{D}=21$ | $℃$ | ^13^ |
| carrying capacity parameters | $\left[ a_{rain},a_{dens} \right]=[{10}^{-5}, {10}^{-3}]$ | ${[mm}^{-1},$  ${km}^{2}]$ | ^3^ |
| Scaling factor of carrying capacity | ${10}^{6}$ |  | ^3,7,9^ |
| Sigmoidal “step-function” | $f\left( X \right)=\left( 1+expexp \left( 20X \right) \right)^{-1}$ |  |  |

*NA = Not applicable
