## Supplementary data and code for "A climate and population dependent diffusion model forecasts the spread of *Aedes Albopictus* mosquitoes in Europe": AUC_from_prediction.R

```
# Run order:

# run_GAM_SPDE_model__fixed_window_cross_validation__mechanistic_GAM.R 
#       cutoff_year=2017 and cutoff_year=2023
# run_GAM_SPDE_model__fixed_window_cross_validation__climate_GAM.R 
#       cutoff_year=2017 and cutoff_year=2023
# AUC_fixed_window_cross_validation.R
#       case_year=1:2, case_GAM_model=1:2

# case_year==1 -- test set year 4:8: training set 2010-2012, test set 2013-2017
# case_year==2 -- test set year 10:14: training set 2010-2018, test set 2019-2023
# case_GAM_model==1 -- mechanistic life cycle GAM model
# case_GAM_model==2 -- climate and population GAM model


##########################################################################
# Load ###################################################################
##########################################################################

#---Load functions and packages
source("functions.R")

#---Load data
folder_data <- "Data"
load(paste0(folder_data, "/data.RData"))

case_year <- 1 # value: 1 or 2
case_GAM_model <- 1 # value: 1 or 2
if (case_year==1 && case_GAM_model==1) {
  run_id <- "4_8_mechanistic_GAM" # Test set: year 4:8, GAM model: mechanistic life cycle 
} else if (case_year==2 && case_GAM_model==1) {
  run_id <- "10_14_mechanistic_GAM" # Test set: year 10:14, GAM model: mechanistic life cycle
} else if (case_year==1 && case_GAM_model==2) {
  run_id <- "4_8_climate_GAM" # Test set: year 4:8, GAM model: climate and population 
} else if (case_year==2 && case_GAM_model==2) {
  run_id <- "10_14_climate_GAM" # Test set: year 10:14, GAM model: climate and population
}

load(paste0("Output/pred_", run_id, ".RData"))
 
load(paste0("Output/pred_", run_id, "_map.RData"))

data_spde_pred$sum_pred <- sum_pred / j_rep


if (case_year==1) {
  
  year_to_predict_vec <- 4:8
  
} else if (case_year==2) {
  
  year_to_predict_vec <- 10:14
  
} 


##########################################################################
# Plot prediction on map #################################################
##########################################################################

year1 <- year_to_predict_vec[1]
year2 <- year1+1
year3 <- year1+2
year4 <- year1+3
year5 <- year1+4

exponent <- 0.5
regions_at_start <- data_spde %>% filter(year_idx==!!(year1-1), presence==1) %>% 
  pull(geo)
df_to_plot <- right_join(data_spde_pred %>% as_tibble() %>% select(-geometry) %>% 
                           filter(year_idx==!!year1, !geo %in% regions_at_start), # NA values for regions with had Ae. albopictus presence at the last year in the training set
                         map_spde)
p1 <- plot_sf(df_to_plot, 
              "sum_pred", 
              legend_title = "pred. presence", 
              title = sprintf("predicted presence, year %d", year1),
              color_transformation = power_transformation(exponent=exponent))
df_to_plot <- right_join(data_spde_pred %>% as_tibble() %>% select(-geometry) %>% 
                           filter(year_idx==!!year2, !geo %in% regions_at_start), 
                         map_spde)
p2 <- plot_sf(df_to_plot, 
              "sum_pred", 
              legend_title = "pred. presence", 
              title = sprintf("predicted presence, year %d", year2),
              color_transformation = power_transformation(exponent=exponent))
df_to_plot <- right_join(data_spde_pred %>% as_tibble() %>% select(-geometry) %>% 
                           filter(year_idx==!!year3, !geo %in% regions_at_start), 
                         map_spde)
p3 <- plot_sf(df_to_plot, 
              "sum_pred", 
              legend_title = "pred. presence", 
              title = sprintf("predicted presence, year %d", year3),
              color_transformation = power_transformation(exponent=exponent))
df_to_plot <- right_join(data_spde_pred %>% as_tibble() %>% select(-geometry) %>% 
                           filter(year_idx==!!year4, !geo %in% regions_at_start), 
                         map_spde)
p4 <- plot_sf(df_to_plot, 
              "sum_pred", 
              legend_title = "pred. presence", 
              title = sprintf("predicted presence, year %d", year4),
              color_transformation = power_transformation(exponent=exponent))
df_to_plot <- right_join(data_spde_pred %>% as_tibble() %>% select(-geometry) %>% 
                           filter(year_idx==!!year5, !geo %in% regions_at_start), 
                         map_spde)
p5 <- plot_sf(df_to_plot, 
              "sum_pred", 
              legend_title = "pred. presence", 
              title = sprintf("predicted presence, year %d", year5),
              color_transformation = power_transformation(exponent=exponent))
p <- ggarrange(p1,p2,p3,p4,p5,ncol=2,nrow=3)
print(p)


##########################################################################
# AUC, fixed window cross-validation #####################################
##########################################################################

#---Compute ROC and AUC
data_new_observations <- data_spde %>% filter(presence_previous_year==0)

tb_for_AUC <- data_spde_pred %>% 
  filter(year_idx %in% year_to_predict_vec, !is.na(presence)) %>% # select test set, regions with observations
  select(geo,year_idx,presence,sum_pred) %>% 
  rename(y_orig = presence, p.mean = sum_pred) %>% 
  right_join(data_new_observations %>% select(geo, year_idx)) # select new observations (observations where Ae. albopictus was not present the previous year)

prob <- tb_for_AUC$p.mean
y <- tb_for_AUC$y_orig

output <- roc_and_auc(prob, y)

cat(sprintf("AUC on test set: %.3g\n", output$auc))


#---Plot ROC
p <- plot_roc_curve(output$tb_roc_curve)
print(p)
save_to_png(p, filename=paste0("Plots/ROC_curve_", model_str_main, ".png"), 
            resolution_scaling_width=1, resolution_scaling_height=1.5)
```
