## Supplementary data and code for "A climate and population dependent diffusion model forecasts the spread of *Aedes Albopictus* mosquitoes in Europe": functions.R

```
#---Load packages
packages <- c("tidyverse", "stringr", "caret", "sf", "spdep", "sp", 
              "INLA", "inlabru", "pROC", "png", "scales", 
              "ggplot2", "RColorBrewer", "oompaBase", "pals", "paletteer",
              "ggpubr", "ggiraph", "gtable", "grid", "gridExtra", "plotly",
              "lattice")

#uncomment if installing packages
# #---install.packages
# lapply(packages, install.packages, character.only = TRUE)

#---load packages
lapply(packages, library, character.only = TRUE)


#---Load functions
create_mesh <- function(map_spde, mesh_precision="mid") {
  
  n_spde <- list()
  n_spde$geo <- nrow(map_spde)
  
  
  # Create mesh -----------------------------------------------------------
  
  shapes <- as_Spatial(map_spde$geometry)
  region_coords <- coordinates(shapes)
  
  map_spde$region_id <- 1:nrow(map_spde)
  
  
  
  # # #---Convex hull
  # # # spde_mesh = inla.mesh.2d(loc=region_coords, cutoff=0.015, max.edge=c(0.1,0.5))
  # # # spde_mesh = inla.mesh.2d(loc=region_coords, cutoff=0.025, max.edge=c(0.2,0.5))
  # # spde_mesh = inla.mesh.2d(loc=region_coords, cutoff=0.025, max.edge=c(0.5,0.5))
  # spde_mesh = inla.mesh.2d(loc=region_coords, cutoff=0.3, max.edge=c(0.35,1))
  # spde_mesh = inla.mesh.2d(loc=region_coords, cutoff=0.3, max.edge=c(0.35,1),
  #                          offset = c(0.5, 7.5))
  # 
  # # 
  # # spde_mesh = inla.mesh.2d(loc=region_coords, cutoff=0.025, max.edge=c(0.5,1), offset=1)
  # # 
  # # cat("Mesh, size: ", spde_mesh$n, "\n")
  # # plot(spde_mesh)
  # # points(region_coords, pch=20, col=2)
  
  
  # #---Non-convex hull
  boundary <- inla.nonconvex.hull(region_coords, convex = 1.2)
  lines(boundary, add = FALSE)
  points(region_coords)
  
  # spde_mesh <- inla.mesh.2d(loc=region_coords, max.n.strict=nrow(map_spde)*5, boundary=boundary)
  
  if (mesh_precision=="high") {
    spde_mesh <- inla.mesh.2d(loc=region_coords, cutoff=0.3, max.edge=c(0.35,1), boundary=boundary)
  } else if (mesh_precision=="mid") {
    spde_mesh <- inla.mesh.2d(loc=region_coords, cutoff=0.45, max.edge=c(0.45,2), boundary=boundary)
  } else if (mesh_precision=="low") {
    spde_mesh <- inla.mesh.2d(loc=region_coords, cutoff=0.6, max.edge=c(0.6,3), boundary=boundary)
  }  
  
  
  # mesh_coords <- spde_mesh$loc[,1:2]
  # points(mesh_coords, pch=20, col=2)
  # points(matrix(mesh_coords[1050,],nrow=1,ncol=2), pch=20, col=2)
  
  # cat("Mesh, size: ", spde_mesh$n, "\n")
  # par(mfrow=c(1,1))
  # plot(spde_mesh)
  # plot_geometry(map_inla, add=TRUE)
  # points(region_coords, pch=20, col=2)
  
  n_spde$full_mesh <- nrow(spde_mesh$loc)
  
  
  return(list(spde_mesh=spde_mesh, region_coords=region_coords, n_spde=n_spde))
}


setup_for_inla <- function(map_spde, data_spde) {
  #---Projection matrix
  shapes <- as_Spatial(map_spde$geometry)
  region_coords <- coordinates(shapes)
  
  tb_coords <- left_join(data_spde %>% select(data_idx, geo),
                         tibble(geo=map_spde$geo, coords=region_coords))
  
  A <- inla.spde.make.A(mesh=spde_mesh, loc=as.matrix(tb_coords$coords),
                        group = data_spde$year_idx)
  
  #---SPDE indices
  spde_indices <- inla.spde.make.index("spde_idx", n.spde = n_spde$full_mesh, 
                                       n.group=length(unique(data_spde$year_idx)))
  
  #---Stack linking SPDE with data
  stack = inla.stack(
    data = data_spde['presence'],
    A = list(A, 1),
    effects = list(spde_indices,
                   list(Intercept = 1,
                        year_idx = data_spde$year_idx,
                        bin_idx_median_temp = data_spde$bin_idx_median_temp,
                        bin_idx_min_temp_CROSS=data_spde$bin_idx_min_temp_CROSS,
                        bin_idx_median_relative_humidity = data_spde$bin_idx_median_relative_humidity,
                        bin_idx_log_pop_merged = data_spde$bin_idx_log_pop_merged,
                        bin_idx_mean_adults_pow_merged = data_spde$bin_idx_mean_adults_pow_merged,
                        bin_idx_proximity050 = data_spde$bin_idx_proximity050,
                        presence_previous_year = data_spde$presence_previous_year,
                        year1 = data_spde$year1)),
    tag = "spde_data"
  )
  
  #---Link function for NA values
  link <- rep(NA, length(stack$data$data$presence))
  ind_NA <- which(is.na(stack$data$data$presence))
  link[ind_NA] <- 1
  
  return(list(spde_indices=spde_indices, stack=stack, link=link))
}


specify_network_by_nodes_and_links <- function(n_nodes, 
                                               nodes_to_disconnect=NULL, 
                                               links_to_remove=NULL, 
                                               links_to_add=NULL,
                                               save_to_file=NULL) {
  
  nb_grid <- cell2nb(nrow=1, ncol=n_nodes)
  if (!is.null(nodes_to_disconnect)) {
    for (j in 1:n_nodes) {
      if (j %in% nodes_to_disconnect) {
        nb_grid[[j]] <- as.integer(0)
      } else {
        nb_grid[[j]] <- setdiff(nb_grid[[j]], nodes_to_disconnect)
      }
    }
  }
  
  if (!is.null(links_to_remove)) {
    for (j in 1:length(links_to_remove)) {
      link_to_disconnect <- links_to_remove[[j]]
      
      node1 <- link_to_disconnect[1]
      node2 <- link_to_disconnect[2]
      nb_grid[[node1]] <- setdiff(nb_grid[[node1]], node2)
      nb_grid[[node2]] <- setdiff(nb_grid[[node2]], node1)
    }
  }
  
  if (!is.null(links_to_add)) {
    for (j in 1:length(links_to_add)) {
      nodes_j <- links_to_add[[j]]
      
      for (k in nodes_j) {
        nb_grid[[k]] <- c(nb_grid[[k]], as.integer(setdiff(nodes_j, k)))
      }
    }
  }
  
  coords <- tibble(x=1, y=1:n_nodes)
  coords$x[coords$y %in% nodes_to_disconnect] <- 2
  for (j in 1:length(links_to_add)) {
    nodes_j <- links_to_add[[j]]
    coords$x[coords$y %in% nodes_j] <- coords$x[coords$y %in% nodes_j] - j*0.25
  }
  plot(nb_grid, coords %>% as.matrix())
  
  if (!is.null(save_to_file))
    nb2INLA(save_to_file, nb_grid)
  
  return(nb_grid)
}


process_fitted_model <- function(data_spde, fitted_model, covariate_set, 
                                 spde_indices, stack, mesh_proj) {
  
  #---Process SPDE fit
  year_idx_vec <- sort(unique(data_spde$year_idx))
  
  tb_regions <- data_spde %>% select(geo, region_id) %>% distinct()
  data_spde_with_na <- 
    tidyr::crossing(tb_regions, tibble(year_idx=unique(data_spde$year_idx))) %>% 
    arrange(year_idx, region_id) %>% left_join(data_spde)
  
  map_full_model <- NULL
  
  for (year_idx__ in year_idx_vec) {
    
    spde_projected <- 
      inla.mesh.project(
        mesh_proj, 
        fitted_model$summary.random$spde_idx$mean[spde_indices$spde_idx.group==year_idx__]
      )
    
    map_full_model <- bind_rows(map_full_model, 
                                bind_cols(data_spde_with_na %>% filter(year_idx == year_idx__) %>% 
                                            select(presence, geo, region_id, year_idx),
                                          spde_projected=spde_projected))
  }
  
  map_full_model <- rename(map_full_model, y = presence)
  
  
  
  #---Subset results
  idx_linear_predictor <- inla.stack.index(stack, "spde_data")$data
  idx_subset <- idx_linear_predictor[1:nrow(data_spde)]
  
  map_full_model$linear_predictor.mean <- fitted_model$summary.linear.predictor[idx_subset, "mean"]
  map_full_model$linear_predictor.sd <- fitted_model$summary.linear.predictor[idx_subset, "sd"]
  map_full_model$p.mean <- fitted_model$summary.fitted.values[idx_subset, "mean"]
  map_full_model$p.sd <- fitted_model$summary.fitted.values[idx_subset, "sd"]
  
  
  
  #---Process min. temp. fit
  if (covariate_set==2) {
    fitted_model <- process_min_temp(data_spde, fitted_model)
  }
  
  
  
  #---Main summaries
  results_summary <- tibble(DIC = round(fitted_model$dic$dic,2), #goodness of fit: DIC
                            logscore = round(-mean(log(fitted_model$cpo$cpo), na.rm = T), 3), #goodness of fit: logscore
                            WAIC = round(fitted_model$waic$waic, 2),
                            fixed_effects = list(fitted_model$summary.fixed), #all fixed effect parameters
                            intercept = fitted_model$summary.fixed["Intercept","mean"], #intercept
                            hyperpar = list(fitted_model$summary.hyperpar)) #fitted hyper parameters
  
  
  
  #---Random effects, constant in time
  #---Initialize
  functions_random_effects <- list()
  map_random_effects <- data_spde %>% filter(year_idx==1) %>% 
    select(data_idx, geo, region_id) #these random effects are constant in time so enough to select first year
  
  #---Add random effects
  #---Random effects for covariates constant in time
  if (covariate_set==1) {
    
    out <- add_random_effect(map_random_effects, functions_random_effects, 
                             random_effect_col="bin_idx_mean_adults_pow_merged")
    map_random_effects <- out$map_random_effects
    functions_random_effects <- out$functions_random_effects
  } else if (covariate_set==2) {
    
    random_effect_vec <- c("bin_idx_median_temp", "bin_idx_min_temp_CROSS",
                           "bin_idx_median_relative_humidity", "bin_idx_log_pop_merged")
    
    for (random_effect_col in random_effect_vec) {
      out <- add_random_effect(map_random_effects, functions_random_effects, 
                               random_effect_col=random_effect_col)
      map_random_effects <- out$map_random_effects
      functions_random_effects <- out$functions_random_effects
    }
  }
  
  
  
  #---Random effect for time-varying covariate
  random_effect_col <- "bin_idx_proximity050"
  
  functions_random_effects[[random_effect_col]] <-
    tibble(!!random_effect_col := fitted_model$summary.random[[random_effect_col]]$ID,
           mean = fitted_model$summary.random[[random_effect_col]]$mean,
           sd = fitted_model$summary.random[[random_effect_col]]$sd,
           lower_bound = fitted_model$summary.random[[random_effect_col]]$`0.025quant`,
           upper_bound = fitted_model$summary.random[[random_effect_col]]$`0.975quant`)
  
  random_effect_str <- paste0("reffect_", random_effect_col)
  
  map_random_effects_curr <-
    left_join(data_spde %>% select(data_idx, region_id, year_idx, !!sym(random_effect_col)),
              functions_random_effects[[random_effect_col]] %>%
                rename(!!paste0(random_effect_str, ".mean") := mean,
                       !!paste0(random_effect_str, ".sd") := sd,
                       !!paste0(random_effect_str, ".lower_bound") := lower_bound,
                       !!paste0(random_effect_str, ".upper_bound") := upper_bound))
  map_random_effects_with_time <- left_join(data_spde %>% select(data_idx, geo, region_id, year_idx), 
                                            map_random_effects_curr)
  
  
  
  #---Map full model
  map_full_model$y_orig <- data_spde$y_orig
  map_full_model$training_set <- data_spde$training_set
  map_full_model$test_set <- data_spde$test_set
  
  #---Abs error and deviance
  map_full_model$error <- compute_error(map_full_model$p.mean, map_full_model$y, n=1)
  results_summary$total_abs_error <- sum(map_full_model$error, na.rm=TRUE)
  
  map_full_model$deviance <- compute_bernoulli_deviance(map_full_model$p.mean, map_full_model$y)
  results_summary$total_deviance <- sum(map_full_model$deviance, na.rm=TRUE)
  
  if (sum(map_full_model$test_set)==0) {
    
    warning("test set is empty")
    
    map_full_model$error_cv <- NA
    map_full_model$deviance_cv <- NA
    results_summary$total_abs_error_cv <- NA
    results_summary$total_deviance_cv <- NA
  } else {
    
    map_full_model$error_cv <- 
      compute_error(map_full_model$p.mean, map_full_model$y_orig)
    results_summary$total_abs_error_cv <- 
      sum(map_full_model %>% filter(test_set==TRUE) %>% pull(error_cv), na.rm=TRUE)
    
    map_full_model$deviance_cv <- 
      compute_bernoulli_deviance(map_full_model$p.mean, map_full_model$y_orig)
    results_summary$total_deviance_cv <- 
      sum(map_full_model %>% filter(test_set==TRUE) %>% pull(deviance_cv), na.rm=TRUE)
  }
  
  
  
  return(list(results_summary=results_summary,
              map_full_model=map_full_model,
              map_random_effects=map_random_effects,
              map_random_effects_with_time=map_random_effects_with_time,
              functions_random_effects=functions_random_effects))
}


add_random_effect <- function(map_random_effects, functions_random_effects, random_effect_col) {
  
  functions_random_effects[[random_effect_col]] <- 
    tibble(!!random_effect_col := fitted_model$summary.random[[random_effect_col]]$ID,
           mean = fitted_model$summary.random[[random_effect_col]]$mean,
           sd = fitted_model$summary.random[[random_effect_col]]$sd,
           lower_bound = fitted_model$summary.random[[random_effect_col]]$`0.025quant`,
           upper_bound = fitted_model$summary.random[[random_effect_col]]$`0.975quant`)
  
  random_effect_str <- paste0("reffect_", random_effect_col)
  
  map_random_effects_curr <- 
    left_join(data_spde %>% select(data_idx, region_id, !!sym(random_effect_col)),
              functions_random_effects[[random_effect_col]] %>% 
                rename(!!paste0(random_effect_str, ".mean") := mean,
                       !!paste0(random_effect_str, ".sd") := sd,
                       !!paste0(random_effect_str, ".lower_bound") := lower_bound,
                       !!paste0(random_effect_str, ".upper_bound") := upper_bound))
  map_random_effects <- left_join(map_random_effects, map_random_effects_curr)
  
  return(list(map_random_effects=map_random_effects,
              functions_random_effects=functions_random_effects))
}


compute_error <- function(p, y, n=1, abs=TRUE) {
  err <- rep(NA, length(p))
  
  idx_non_na <- which(!is.na(y))
  if (any(is.na(p[idx_non_na])))
    stop("na error")
  
  p_num <- p[idx_non_na]
  p_num[p_num > 1-1e-10] <- 1-1e-10
  p_num[p_num < 1e-10] <- 1e-10
  
  if (abs) {
    err[idx_non_na] <- abs(p_num*n - y[idx_non_na])
  } else {
    err[idx_non_na] <- p_num*n - y[idx_non_na]
  }
  
  return(err)
}


compute_bernoulli_deviance <- function(p, y) {
  deviance <- rep(NA, length(y))
  
  idx_non_na <- which(!is.na(y))
  
  if (any(is.na(p[idx_non_na])))
    stop("na error")
  
  p_num <- p[idx_non_na]
  y_num <- y[idx_non_na]
  
  p_num[p_num > 1-1e-10] <- 1-1e-10
  p_num[p_num < 1e-10] <- 1e-10
  
  # deviance = -2 * log likelihood
  deviance[idx_non_na] <- -2 * ((y_num * log(p_num) + (1-y_num) * log(1-p_num)))
  
  return(deviance)
}


process_min_temp <- function(data_spde, fitted_model) {
  
  # transfer random effect from min_temp_CROSS to median_temp (for low median_temp)
  target_for_transfer <- data_spde %>% filter(bin_idx_min_temp_CROSS==1) %>% 
    pull(bin_idx_median_temp) %>% unique()
  
  random_effect_to_transfer <- fitted_model$summary.random$bin_idx_min_temp_CROSS[1,]
  
  fitted_model$summary.random$bin_idx_median_temp$mean[1:2] <- 
    fitted_model$summary.random$bin_idx_median_temp$mean[1:2] + 
    random_effect_to_transfer$mean
  fitted_model$summary.random$bin_idx_median_temp$`0.025quant`[1:2] <-
    fitted_model$summary.random$bin_idx_median_temp$`0.025quant`[1:2] +
    random_effect_to_transfer$mean
  fitted_model$summary.random$bin_idx_median_temp$`0.975quant`[1:2] <-
    fitted_model$summary.random$bin_idx_median_temp$`0.975quant`[1:2] +
    random_effect_to_transfer$mean
  
  return(fitted_model)
  
}


plot_on_map_selected_model_parts <- function(map_random_effects, 
                                             map_random_effects_with_time,
                                             map_full_model, 
                                             map_spde,
                                             year_to_plot=9,
                                             plot_larger=FALSE) {
  
  ps_map <- list()
  
  if (!is.null(map_random_effects) && nrow(map_random_effects)>0) {
    
    if ("reffect_bin_idx_median_temp.mean" %in% names(map_random_effects)) {
      #---Plot median temp random effect (on map)
      ps_map[[length(ps_map)+1]] <- 
        plot_sf(map_random_effects %>% left_join(map_spde), 
                "reffect_bin_idx_median_temp.mean",
                title="RE, median temp.",
                legend_title="median temp.",
                plot_larger=plot_larger)
    }
    
    if ("reffect_bin_idx_min_temp_CROSS.mean" %in% names(map_random_effects)) {
      map_random_effects <- map_random_effects %>% 
        mutate(reffect_bin_idx_min_temp_CROSS.meanNA=
                 if_else(bin_idx_min_temp_CROSS==1, NA,
                         reffect_bin_idx_min_temp_CROSS.mean))
      #---Plot min temp CROSS random effect (on map)
      ps_map[[length(ps_map)+1]] <- 
        plot_sf(map_random_effects %>% left_join(map_spde), 
                "reffect_bin_idx_min_temp_CROSS.meanNA",
                title="RE, min. temp. CROSS",
                legend_title="min. temp. CROSS",
                plot_larger=plot_larger)
    }
    
    if ("reffect_bin_idx_median_relative_humidity.mean" %in% names(map_random_effects)) {
      #---Plot median relative humidity random effect (on map)
      ps_map[[length(ps_map)+1]] <- 
        plot_sf(map_random_effects %>% left_join(map_spde), 
                "reffect_bin_idx_median_relative_humidity.mean",
                title="RE, median relative humidity",
                legend_title="median rel. humidity",
                plot_larger=plot_larger)
    }
    
    if ("reffect_bin_idx_log_pop_merged.mean" %in% names(map_random_effects)) {
      #---Plot log population random effect (on map)
      ps_map[[length(ps_map)+1]] <- 
        plot_sf(map_random_effects %>% left_join(map_spde),
                "reffect_bin_idx_log_pop_merged.mean",
                title="RE, log population",
                legend_title="log pop.",
                plot_larger=plot_larger)
    }
    
    if ("reffect_bin_idx_mean_adults_pow_merged.mean" %in% names(map_random_effects)) {
      #---Plot mechanistic model random effect (on map)
      ps_map[[length(ps_map)+1]] <- 
        plot_sf(map_random_effects %>% left_join(map_spde), 
                "reffect_bin_idx_mean_adults_pow_merged.mean",
                title="RE, mechanistic model",
                legend_title="mech. model",
                plot_larger=plot_larger)
    }
    
  }
  
  
  
  if (!is.null(map_full_model$spde_projected)) {
    
    #---Plot spatio-temporal field
    ps_map[[length(ps_map)+1]] <- 
      plot_sf(map_full_model %>% left_join(map_spde) %>% 
                filter(year_idx==year_to_plot), "spde_projected",
              title=sprintf("RE, spatio-temporal field, year %d", year_to_plot),
              legend_title="SPDE",
              plot_larger=plot_larger)
  }
  
  
  
  #---Plot proximity random effect
  non_proximity_bin_idx <- 1:2 #exclude bin indices corresponding to presence previous year and year 1 from the plot
  ps_map[[length(ps_map)+1]] <- 
    plot_on_map_proximity_random_effect(map_random_effects_with_time, map_spde, 
                                        non_proximity_bin_idx,
                                        year_to_plot=year_to_plot,
                                        plot_larger=plot_larger)
  
  
  
  #---Plot fitted model probability (p.mean) and the data (y)
  #---Fitted probability
  ps_map[[length(ps_map)+1]] <- 
    plot_sf(map_full_model %>% filter(year_idx==year_to_plot) %>% 
              left_join(map_spde), 
            "p.mean",
            title=sprintf("prob., year %d", year_to_plot),
            color_transformation=power_transformation(0.1),
            legend_title="prob.",
            plot_larger=plot_larger)
  #---Fitted probability, only showing new observations
  ps_map[[length(ps_map)+1]] <- 
    plot_sf(map_full_model %>% filter(year_idx==year_to_plot) %>% left_join(data_spde) %>%  
              mutate(p.mean.new_obs=ifelse(presence_previous_year==1,NA,p.mean)) %>% # set to NA if Ae. albopictus was observed in the region the previous year
              left_join(map_spde), 
            "p.mean.new_obs",
            title=sprintf("prob., new obs., year %d", year_to_plot),
            legend_title="prob., new obs.",
            color_transformation=power_transformation(0.1),
            plot_larger=plot_larger)
  #---Data, current year
  ps_map[[length(ps_map)+1]] <- 
    plot_sf(map_full_model %>% filter(year_idx==year_to_plot) %>% 
              left_join(map_spde), 
            "y",
            title=sprintf("simulated data, year %d", year_to_plot),
            plot_larger=plot_larger,
            palette_idx=1)
  
  if (year_to_plot > 1) {
    #---Data, previous year
    ps_map[[length(ps_map)+1]] <- 
      plot_sf(map_full_model %>% filter(year_idx==(year_to_plot-1)) %>% 
                left_join(map_spde), 
              "y",
              title=sprintf("simulated data, year %d", year_to_plot-1),
              plot_larger=plot_larger,
              palette_idx=1)
  }
  
  #---Collect figures into one
  n_figures <- length(ps_map)
  ncol <- floor(sqrt(n_figures))
  nrow <- ceiling(n_figures / ncol)
  p <- do.call(ggarrange, c(ps_map, nrow=nrow, ncol=ncol))
  
  return(list(p=p, ps_map=ps_map))
}


plot_on_map_proximity_random_effect <- function(map_random_effects_with_time, 
                                                map_spde, 
                                                special_group_bin_idx, 
                                                year_to_plot=9,
                                                plot_larger=FALSE) {
  
  proximity_col <- names(map_random_effects_with_time) %>% str_subset("reffect_bin_idx_proximity.*mean")
  
  bin_idx_col <- names(map_random_effects_with_time) %>% str_subset("bin_idx_proximity[0-9]*$")
  row_idx_special_groups <- map_random_effects_with_time[[bin_idx_col]] %in% special_group_bin_idx
  
  map_random_effects_with_time$proximity_excl_special_groups <- 
    map_random_effects_with_time[[proximity_col]]
  map_random_effects_with_time$proximity_excl_special_groups[row_idx_special_groups] <- NA #exclude bin indices corresponding to presence previous year and year 1 from the plot
  
  p <- plot_sf(left_join(map_spde, map_random_effects_with_time %>% filter(year_idx==year_to_plot)), 
               "proximity_excl_special_groups",
               title=sprintf("RE, proximity, year %d", year_to_plot),
               legend_title="proximity",
               plot_larger=plot_larger)
  
  return(p)
}


plot_random_effects_as_functions <- function(functions_random_effects, keys,
                                             data_spde, model_str, 
                                             plot_rug_vector=FALSE,
                                             plot_larger=FALSE,
                                             ylim=NULL,
                                             resolution_scaling_width=1) {
  
  random_effects_col <- names(functions_random_effects)
  
  ps <- list()
  
  if ("bin_idx_median_temp" %in% random_effects_col) {
    if (plot_rug_vector) {
      rug_vector <- data_spde$median_temp
    } else {
      rug_vector <- NULL
    }
    
    keys$median_temp$n_in_bin <- sapply(keys$median_temp$bin_idx, 
                                        function(i) {sum(data_spde$bin_idx_median_temp == i)})
    p_median_temp <- plot_random_effect(functions_random_effects, keys=keys$median_temp, 
                                        col_subscript="median_temp",
                                        vector_to_plot_with_rug=rug_vector, 
                                        plot_larger=plot_larger,
                                        ylim=ylim,
                                        resolution_scaling_width=resolution_scaling_width)
    
    ps$p_median_temp <- p_median_temp
  }
  
  if ("bin_idx_min_temp_CROSS" %in% random_effects_col) {
    if (plot_rug_vector) {
      rug_vector <- data_spde$min_temp
    } else {
      rug_vector <- NULL
    }
    
    keys$min_temp_CROSS$n_in_bin <- sapply(keys$min_temp_CROSS$bin_idx, 
                                           function(i) {sum(data_spde$bin_idx_min_temp_CROSS == i, na.rm=TRUE)})
    p_min_temp_CROSS <- plot_random_effect(functions_random_effects, keys=keys$min_temp_CROSS, 
                                           col_subscript="min_temp_CROSS",
                                           vector_to_plot_with_rug=rug_vector, 
                                           plot_larger=plot_larger,
                                           ylim=ylim,
                                           resolution_scaling_width=resolution_scaling_width)
    
    ps$p_min_temp_CROSS <- p_min_temp_CROSS
  }
  
  if ("bin_idx_median_relative_humidity" %in% random_effects_col) {
    if (plot_rug_vector) {
      rug_vector <- data_spde$median_relative_humidity
    } else {
      rug_vector <- NULL
    }
    
    keys$median_relative_humidity$n_in_bin <- 
      sapply(keys$median_relative_humidity$bin_idx, 
             function(i) {sum(data_spde$bin_idx_median_relative_humidity == i, na.rm=TRUE)})
    p_median_relative_humidity <- 
      plot_random_effect(functions_random_effects, keys=keys$median_relative_humidity, 
                         col_subscript="median_relative_humidity",
                         vector_to_plot_with_rug=rug_vector, 
                         plot_larger=plot_larger,
                         ylim=ylim,
                         resolution_scaling_width=resolution_scaling_width)
    
    ps$p_median_relative_humidity <- p_median_relative_humidity
  }
  
  if ("bin_idx_log_pop_merged" %in% random_effects_col) {
    if (plot_rug_vector) {
      rug_vector <- data_spde$log_population
    } else {
      rug_vector <- NULL
    }
    
    keys$log_pop_merged$n_in_bin <- sapply(keys$log_pop_merged$bin_idx, 
                                           function(i) {sum(data_spde$bin_idx_log_pop_merged == i, na.rm=TRUE)})
    p_log_pop_merged <- plot_random_effect(functions_random_effects, keys=keys$log_pop_merged, 
                                           col_subscript="log_pop_merged",
                                           vector_to_plot_with_rug=rug_vector,
                                           plot_larger=plot_larger,
                                           ylim=ylim,
                                           resolution_scaling_width=resolution_scaling_width)
    
    ps$p_log_pop_merged <- p_log_pop_merged
  }
  
  if ("bin_idx_mean_adults_pow_merged" %in% random_effects_col) {
    if (plot_rug_vector) {
      rug_vector <- data_spde$mean_adults_pow_merged
    } else {
      rug_vector <- NULL
    }
    
    keys$mean_adults_pow_merged$n_in_bin <- 
      sapply(keys$mean_adults_pow_merged$bin_idx, 
             function(i) {sum(data_spde$bin_idx_mean_adults_pow_merged == i, na.rm=TRUE)})
    p_mean_ODE <- plot_random_effect(functions_random_effects, keys=keys$mean_adults_pow_merged, 
                                     col_subscript="mean_adults_pow_merged",
                                     vector_to_plot_with_rug=rug_vector, 
                                     plot_larger=plot_larger,
                                     ylim=ylim,
                                     resolution_scaling_width=resolution_scaling_width)
    
    ps$p_mean_ODE <- p_mean_ODE
  }
  
  if (any(str_detect(random_effects_col, "proximity"))) {
    keys$y_proximity_exp050$n_in_bin <- sapply(keys$y_proximity_exp050$bin_idx, 
                                               function(i) {sum(data_spde$bin_idx_proximity050 == i, na.rm=TRUE)})
    p_proximity <- plot_proximity_random_effect(functions_random_effects, keys=keys, data=data_spde, 
                                                plot_rug_vector=plot_rug_vector,
                                                plot_larger=plot_larger,
                                                ylim=ylim,
                                                resolution_scaling_width=resolution_scaling_width)
    
    ps$p_proximity <- p_proximity
  }
  
  return(list(ps))
}


plot_random_effect <- function(functions_random_effects, keys, 
                               col_subscript, vector_to_plot_with_rug=NULL, 
                               filename=NULL,
                               plot_larger=FALSE,
                               ylim=NULL,
                               resolution_scaling_width=1,
                               resolution_scaling_height=1) {
  
  bin_idx_col <- paste0("bin_idx_", col_subscript)
  bin_range_col <- paste0("bin_range_", col_subscript)
  bin_mean_col <- paste0("bin_mean_", col_subscript)
  
  #---Plot random effect as a function
  #---Prepare for plotting
  tb_reffect <- functions_random_effects[[bin_idx_col]]
  #---Add bin range and mean of original column
  tb_reffect <- left_join(tb_reffect, keys,
                          by = join_by(!!sym(bin_idx_col) == bin_idx)) %>% 
    relocate(sym(bin_idx_col), sym(bin_range_col), sym(bin_mean_col))
  
  #---Plot with x_column="bin_mean", and make bin position in bottom histogram correspond to the range of values within each bin 
  re_plots <- 
    linked_random_effect_and_histogram_plot(tb_reffect, 
                                            x_column_type="bin_mean", 
                                            bin_width_constant=FALSE,
                                            vector_to_plot_with_rug=vector_to_plot_with_rug,
                                            plot_larger=plot_larger,
                                            ylim=ylim)
  print(tb_reffect)
  if (!is.null(filename)) {
    #---Save plot
    save_to_png(re_plots$g, filename=filename, 
                resolution_scaling_width=resolution_scaling_width,
                resolution_scaling_height=resolution_scaling_height)
  }
  
  return(re_plots$g)
}


plot_proximity_random_effect <- function(functions_random_effects, keys, data, 
                                         filename=NULL,
                                         plot_rug_vector=FALSE,
                                         plot_larger=FALSE,
                                         ylim=NULL,
                                         resolution_scaling_width=1) {
  
  #---Plot proximity random effect (as a function)
  bin_idx_proximity_col <- names(functions_random_effects) %>% str_subset("proximity")
  proximity_exp <- str_extract(bin_idx_proximity_col, "(?<=proximity)[0-9].*")
  keys_col <- str_subset(names(keys), paste0("exp", proximity_exp))
  
  #---Prepare for plotting
  reffect_proximity <- functions_random_effects[[bin_idx_proximity_col]]
  
  reffect_proximity <- reffect_proximity %>% filter(bin_idx_proximity050>=3) #exclude bin indices corresponding to presence previous year and year 1 from the plot
  
  #---Add bin range and mean of original proximity
  reffect_proximity <- left_join(reffect_proximity, keys[[keys_col]],
                                 by = join_by(!!sym(bin_idx_proximity_col) == bin_idx))
  
  #---Plot with x_column="bin_mean", and make bin position in bottom histogram correspond to the range of values within each bin 
  if (plot_rug_vector) {
    rug_vector <- data$log_population
  } else {
    rug_vector <- NULL
  }
  
  re_plots_proximity <- 
    linked_random_effect_and_histogram_plot(reffect_proximity, 
                                            x_column_type="bin_mean", 
                                            bin_width_constant=FALSE,
                                            vector_to_plot_with_rug=rug_vector,
                                            plot_larger=plot_larger,
                                            ylim=ylim)
  # grid.draw(re_plots_max_temp$g)
  print(reffect_proximity)
  
  if (!is.null(filename)) {
    #---Save plot
    save_to_png(re_plots_proximity$g, filename=filename, resolution_scaling_width=resolution_scaling_width)
  }
  
  return(re_plots_proximity$g)
}


linked_random_effect_and_histogram_plot <- function(df_random_effect, 
                                                    x_column_type="bin_mean", #"bin_mean", "bin_idx" or "interaction_bin_mean"
                                                    interaction_x_column=NULL,
                                                    random_effect_title="",
                                                    bin_width_constant=TRUE,
                                                    vector_to_plot_with_rug=NULL,
                                                    plot_larger=FALSE,
                                                    ylim=NULL,
                                                    reduce_window_size=TRUE) {
  
  
  n_column <- names(df_random_effect) %>% str_subset("n_in_bin")
  
  if (x_column_type=="bin_mean") {
    
    x_column <- names(df_random_effect) %>% str_subset("bin_mean")
  } else if (x_column_type=="bin_idx") {
    
    x_column <- names(df_random_effect) %>% str_subset("bin_idx")
  } else if (x_column_type=="interaction_bin_idx") {
    
    x_column <- names(df_random_effect) %>% str_subset("bin_idx.*inter")
    
    if (!bin_width_constant) {
      warning("setting bin_width_constant=TRUE, which is required when x_column_type=\"interaction_bin_idx\"")
      bin_width_constant <- TRUE
    }
  } else {
    
    stop("x_column_type should equal either \"bin_mean\", \"bin_idx\", or \"interaction_bin_idx\"")
  }
  
  if (length(x_column) != 1 || length(n_column) != 1)
    stop("interaction_x_column does not match any of the interaction_bin_mean-columns")
  
  df_random_effect_tall <- bind_rows(tibble(!!sym(x_column) := df_random_effect[[x_column]],
                                            random_effect = df_random_effect$lower_bound,
                                            type = "lower_bound"),
                                     tibble(!!sym(x_column) := df_random_effect[[x_column]],
                                            random_effect = df_random_effect$mean,
                                            type = "mean"),
                                     tibble(!!sym(x_column) := df_random_effect[[x_column]],
                                            random_effect = df_random_effect$upper_bound,
                                            type = "upper_bound"))
  
  
  
  #---Manual color palette
  col_rgb <- col2rgb("gray60")
  col_hex <- rgb(col_rgb[1], col_rgb[2], col_rgb[3], maxColorValue=255)
  manual_palette <- c(col_hex, NA, col_hex)
  col_rgb <- col2rgb("gray5")
  col_hex <- rgb(col_rgb[1], col_rgb[2], col_rgb[3], maxColorValue=255)
  manual_palette[2] <- col_hex
  
  
  
  #---Plot random effect
  p_random_effect <- 
    plot_scatter_lines(df_random_effect_tall, x_column = x_column, y_column = "random_effect",
                       color_column = "type", plot_constant_lines_y = 0,
                       palette_idx = 4, 
                       title=random_effect_title,
                       plot_larger=plot_larger,
                       ylim=ylim)
  
  xlimits <- layer_scales(p_random_effect)$x$range$range
  
  
  
  #---Adjust the end-point of first and last bin in histogram to fit within x-range of
  #---the random effect plot
  first_bin <- list()
  first_bin <- df_random_effect %>% head(1) %>% select(bin_min, bin_max) %>% 
    rename(original_bin_min=bin_min)
  first_bin$bin_min <- xlimits[1]
  first_bin <- first_bin %>% mutate(original_length=bin_max-original_bin_min,
                                    new_length=bin_max-bin_min)
  
  df_random_effect[1,"bin_min"] <- first_bin$bin_min
  
  last_bin <- list()
  last_bin <- df_random_effect %>% tail(1) %>% select(bin_min, bin_max) %>% 
    rename(original_bin_max=bin_max) 
  last_bin$bin_max <- xlimits[2]
  last_bin <- last_bin %>% mutate(
    original_length=original_bin_max-bin_min,
    new_length=bin_max-bin_min)
  
  df_random_effect[nrow(df_random_effect),"bin_max"] <- last_bin$bin_max
  
  print(first_bin)
  print(last_bin)
  
  #---Define height so that height*width=number of data points, with width being the bin range
  df_random_effect <- df_random_effect %>% 
    mutate(bin_height=n_in_bin/(bin_max-bin_min))
  
  
  
  #---Plot histogram with bin width corresponding to the bin range, 
  #---and volume corresponding to number of data points within each bin
  #---with the same x-range as the random effect plot.
  #---The x-range here does not correspond to the range of values in the data!
  p_hist <- ggplot(df_random_effect, aes(x = !!sym(x_column), y = bin_height)) +
    geom_rect(aes(xmin = bin_min, xmax = bin_max, 
                  ymin = 0, ymax = bin_height)) +
    theme_classic() + 
    theme(axis.text.x=element_blank(), #remove x-axis labels since they are redundant (same as for p_random_effect)
          axis.ticks.x=element_blank(),
          axis.title.x=element_blank(),
          axis.text.y=element_blank(), #remove y-axis labels
          axis.ticks.y=element_blank(),
          axis.title.y=element_blank()
    )  
  
  if (!is.null(vector_to_plot_with_rug)) {
    
    p_random_effect <- p_random_effect + 
      geom_rug(data=data.frame(x=vector_to_plot_with_rug), aes(x=x, y=0), 
               inherit.aes = FALSE, alpha = 1/2, position = "jitter", sides="b")
  }
  
  g <- align_plots(p_random_effect, p_hist) 
  grid.draw(g)
  
  
  return(list(g=g, p_random_effect=p_random_effect, p_hist=p_hist))
}


roc_and_auc <- function(prob, y) {
  
  roc_curve <- roc(y, prob)
  
  auc_value <- auc(roc_curve)
  
  return(list(auc=auc_value, roc=roc_curve,
              tb_roc_curve=tibble(sensitivity=roc_curve$sensitivities, 
                                  false_negative_rate=1-roc_curve$specificities)))
}


plot_roc_curve <- function(roc_curve, color_column=NULL, plot_larger=TRUE, title="") {
  
  p <- 
    plot_scatter_lines(roc_curve,
                       "false_negative_rate", "sensitivity", color_column=color_column,
                       title=title, label_x="false negative rate", label_y="sensitivity",
                       plot_larger=plot_larger)
  
  return(p)
}


compute_roc_for_cv_set <- function(map_all, 
                                   roc_curve_long=NULL, 
                                   model_prob_long=NULL,
                                   model_name,
                                   plot_curve=TRUE) {
  
  map_sub <- map_all %>% 
    filter(training_set==0, !is.na(y_orig), presence_previous_year==0)
  
  prob <- map_sub$p.mean
  y <- map_sub$y_orig
  
  model_prob_long <- bind_rows(
    model_prob_long,
    tibble(prob=prob, y=y, model_name=model_name)
  )
  
  if (!identical(sort(unique(map_sub$y_orig)), c(0,1))) {
    return(list(model_prob_long=model_prob_long))
  }
  
  output <- roc_and_auc(prob, y)
  
  if (plot_curve) 
    plot_roc_curve(output$tb_roc_curve, title=model_name) %>% print()
  
  roc_curve_long <- bind_rows(
    roc_curve_long,
    tibble(
      false_negative_rate=output$tb_roc_curve$false_negative_rate,
      sensitivity=output$tb_roc_curve$sensitivity,
      model_name=model_name
    )
  )
  
  return(list(roc_curve_long=roc_curve_long, auc=output$auc,
              model_prob_long=model_prob_long))
}


compute_roc_and_auc <- function(tb_results_cumul) {
  
  n_fits <- nrow(tb_results_cumul)
  
  roc_curve_long <- NULL
  model_prob_long <- NULL
  auc_vec <- rep(NA, n_fits)
  for (j in 1:n_fits) {
    
    map_for_ROC <- tb_results_cumul$map_for_ROC[[j]]
    
    output <- compute_roc_for_cv_set(map_for_ROC, 
                                     roc_curve_long=roc_curve_long, 
                                     model_prob_long=model_prob_long,
                                     model_name=tb_results_cumul$model_str[j],
                                     plot_curve=FALSE)
    
    model_prob_long <- output$model_prob_long
    
    if (!is.null(output$roc_curve_long)) {
      roc_curve_long <- output$roc_curve_long
      auc_vec[j] <- output$auc
    }
  }
  
  output <- roc_and_auc(model_prob_long$prob, model_prob_long$y)
  
  return(output)
}


compute_Q_besag <- function(nb_map, n_geo, proper_version = FALSE,
                            lambda = NULL) {
  if (proper_version) {
    if (is.null(lambda)) 
      stop("lambda parameter should be supplied for proper version of Besag")
    if (!(lambda > 0 && lambda < 1))
      stop("invalid lambda-value")
  }
  if (!proper_version && !is.null(lambda)) 
    stop("lambda parameter is not required when proper_version == FALSE")
  
  if (n_geo != length(nb_map))
    stop("n_geo error")
  neighs <- vector(mode = "list", length = n_geo)
  n_neighs <- vector(length = n_geo)
  
  Q = matrix(0, nrow = n_geo, ncol = n_geo)
  for (j in seq_along(nb_map)) {
    ns <- nb_map[[j]]
    if (!all(ns == 0)) {
      neighs[[j]] <- ns
      n_neighs[j] <- length(ns)
      
      Q[neighs[[j]],j] = -1
      Q[j,neighs[[j]]] = -1
      Q[j,j] = n_neighs[j]
    } 
  }
  
  if (proper_version) {
    Q <- lambda * Q
    for (i in 1:n_geo) 
      Q[i,i] <- Q[i,i] + (1-lambda)
  }
  
  return(Q)
}


collect_Q_and_eig <- function(Q) {
  
  rank <- rankMatrix(Q)
  E <- eigen(Q)
  determinant_log <- sum(log(E$values[1:rank]))
  
  return(list(rank = rank, 
              determinant_log = determinant_log,
              E = E,
              Q = Q))
}


Q_convolution <- function(Q_with_eig, exponent, x, NAval = NA, use_noise = FALSE) {
  
  n <- nrow(Q_with_eig$Q)
  
  if (!is.na(NAval)) {
    ind <- which(x==NAval)
  } else {
    ind <- which(is.na(x))
  }
  
  Q_rank <- Q_with_eig$rank
  if (Q_rank < nrow(Q_with_eig$Q)) {
    Q_with_eig$E$values[(Q_with_eig$rank[1]+1):n] <- 0 #numerical instabilities with the 0 eigenvalues
  }
  if (use_noise) {
    d <- median(Q_with_eig$E$values) / 5
  } else {
    d <- 0
  }
  Q <- Q_with_eig$E$vectors %*% diag(Q_with_eig$E$values^exponent + d) %*% t(Q_with_eig$E$vectors)
  
  Q_AA <- Q[ind, ind]
  Q_AB <- Q[ind, -ind]
  
  if (rankMatrix(Q_AA) != nrow(Q_AA)) 
    stop("rank error, Q_AA")
  
  
  if (length(x[-ind]) > 1) {
    x[ind] <- - solve(Q_AA, Q_AB %*% x[-ind])
  } else {
    x[ind] <- - solve(Q_AA, Q_AB * x[-ind])
  }
  
  
  return(x)
}


proximity_from_Q <- function(Q_with_eig, tb, y_column, map, exponent_vec=1,
                             add_year_one=TRUE) {
  #---Convolve matrix Q^exponent (in Q_with_eig <- collect_Q_and_eig(Q)) with the column tb[[y_column]] 
  #---on map.
  #---tb and map are linked through the column geo. 
  #---The convolution is done on a yearly basis by convolving with the known mosq. distribution tb[[y_column]] 
  #---for the previous year. 
  #---The convolution is set to zero the first year since there is no information about the mosq. distribution 
  #---of the previous year. 
  #---The exponent is implemented in Q_convolution as Q^exponent = E %*% L^exponent %*% E^T, where 
  #---Q = E %*% L %*% E^T is the eigen-value decomposition of Q.
  
  year_vec <- sort(unique(tb$year_idx))
  # year_vec <- setdiff(year_vec, max(year_vec))
  df_proximity <- tb %>% select(geo, year_idx)
  df_proximity_last_year <- tibble(geo=unique(df_proximity$geo), year_idx=max(tb$year_idx)+1)
  df_proximity <- bind_rows(df_proximity, df_proximity_last_year)
  for (exponent in exponent_vec) {
    
    # cat(sprintf("Computing convolution of y and Q^exponent, with exponent: %.2g\n", exponent))
    
    y_proximity_str <- sprintf("y_proximity_exp%03d", exponent*100)
    
    #---Compute proximity by convolving y with Q^exponent with current exponent, for each year separately (convolution from previous year as proximity for the next year)
    df_proximity_curr <- NULL
    for (year in year_vec) {
      
      map_year <- left_join(map, tb %>% filter(year_idx == year))
      
      y <- map_year[[y_column]]
      y[is.na(y)] <- 0
      
      #---Convolve y with Q^exponent for the current year
      y_convolved <- Q_convolution(Q_with_eig, exponent=exponent, y, 
                                   NAval=0, use_noise=TRUE)
      
      #---Add convolution with current year as proximity for the next year
      df_proximity_curr <-  df_proximity_curr %>% 
        bind_rows(tibble(!!sym(y_proximity_str) := y_convolved,
                         y = y, 
                         year_idx = year + 1,
                         geo = map_year$geo))
    }
    
    if (add_year_one) {
      #---Set year 1 to zero (as we don't have any data from the previous year to convolve with)
      df_proximity_curr <- df_proximity_curr %>% 
        bind_rows(tibble(!!sym(y_proximity_str) := 0,
                         y = 0, 
                         year_idx = min(df_proximity$year_idx),
                         geo = map_year$geo))
    } 
    
    #---Add convolution with current exponent
    df_proximity <- left_join(df_proximity, df_proximity_curr)
  }
  
  proximity_cols <- str_subset(names(df_proximity_curr), "y_proximity")
  for (col in proximity_cols) {
    df_proximity <- df_proximity[!is.na(df_proximity[[col]]),]
  }
  
  # p_conv <- plot_sf(left_join(map, df_proximity %>% filter(year_idx==2)), "y_conv_exp100")
  # p_conv
  
  return(df_proximity)
}


compute_proximity_for_prediction <- function(df, map, Q_with_eig) {
  
  df <- df %>% 
    mutate(y_to_convolve = 1*(y_predict==1)) #y==1 if presence==1, y==0 if presence %in% c(1, NA)
  
  y_column <- "y_to_convolve"
  
  exponent_vec <- 0.5
  
  df_proximity <- proximity_from_Q(Q_with_eig, df, y_column, map, exponent_vec,
                                   add_year_one=FALSE)
  df_proximity <- df_proximity %>% rename(presence_previous_year=y)
  
  df_surrounded <- get_surrounded_regions()
  
  proximity_col_vec <- names(df_proximity) %>% str_subset("y_proximity")
  
  for (j in 1:nrow(df_proximity)) {
    df_curr <- df_proximity[j,]
    if (df_curr$geo %in% df_surrounded$inner) {
      geo_outer <- df_surrounded %>% filter(inner == df_curr$geo) %>% pull(outer)
      df_curr_match <- df_proximity %>% filter(geo == geo_outer, year_idx == df_curr$year_idx)
      if (df_curr_match$presence_previous_year != 1) {
        for (proximity_col in proximity_col_vec) {
          df_proximity[[proximity_col]][j] <- df_curr_match[[proximity_col]]
        }
      }
    }
  }
  
  return(df_proximity)
}


geo_to_neighs <- function(map, use_snap=TRUE, use_queen=TRUE) {
  #---Assumes that geometry features are stored in a column called geometry
  
  if ("sfc" %in% class(map)) {
    #---Class of input map is valid, but the geometry features are not stored correctly
    #---Create a valid data.frame
    
    warning("creating data.frame from sfc input")
    
    map_geometry <- map
    map <- tibble(geo_idx=1:length(map_geometry), geometry=map_geometry)
  } else if (!"sfc" %in% class(map$geometry)) {
    #---Class of geometry column is not valid
    
    stop("no valid geometry column")
  }
  
  if (!"geo_idx" %in% colnames(map)) {
    #---Create missing geometry index column
    
    map <- map %>% select(geometry) %>% mutate(geo_idx=1:nrow(map))
  }
  
  
  #---Check that there are no duplicate geometries in map
  n_geo <- map$geometry %>% unique() %>% length()
  if (n_geo != nrow(map))
    stop('duplicate geometries in map')
  
  
  #---Create SpatialPolygons object from the geometry column (sfc object) 
  shapes <- as_Spatial(map$geometry)
  # nb_overlap <- st_overlaps(map$geometry, sparse = FALSE) #check if any geometries overlap
  
  
  #---Build neighbor list based on region boundaries
  if (use_snap) {
    
    nb_map <- poly2nb(shapes, snap=sqrt(.Machine$double.eps)*5e5, #snap: boundary points less than snap distance apart are considered neighbors
                      queen = use_queen) #queen: if true, a single shared boundary point meets the neighbor criteria, if false, more than one shared point is required
  } else {
    
    nb_map <- poly2nb(shapes, queen = use_queen) 
  }
  #---Store central coordinates of each geometry
  coords <- coordinates(shapes)
  
  
  
  return(list(geo_idx = map$geo_idx,
              shapes = shapes,
              nb_map = nb_map,
              n_geo = n_geo,
              coords = coords))
}


get_surrounded_regions <- function() {
  tb_surrounded3 <- tibble(outer=c("CZ020", "HU120", "RO322", "EL306", "EL305",
                                   "EL305", "EL305", "BG412", "AT126"),
                           inner=c("CZ010", "HU110", "RO321", "EL302",
                                   "EL304", "EL303", "EL301", "BG411",
                                   "AT130"))
  
  tb_surrounded <- tibble(outer=c("DEG0L", "DEE0B", "DE40G", "DEA2C", "DE734",
                                  "DE716", "DE94E", "DE238", "DE145", "DED2F",
                                  "DE276", "DE219", "DE40E", "DE40E", "DEE07",
                                  "DEE05"),
                          inner=c("DEG02", "DEE02", "DE402", "DEA22", "DE731",
                                  "DE711", "DE944", "DE232", "DE144", "DED21",
                                  "DE271", "DE211", "DE404", "DE401", "DEE03",
                                  "DEE01"))
  
  tb_surrounded2 <- 
    tibble(outer=c("DEB3K", "DEB3H", "DEB3F", "DE124", "DE12B", "DE118",
                   "DEB25", "DE124", "DE27E", "DE27B", "DE256", "DE26C",
                   "DE26B", "DE245", "DE247", "DE246", "DE249", "DE237",
                   "DE234", "DE21K", "DE227", "DE22B", "DEG0G", "DEG0P",
                   "DE734", "DEG0J"), 
           inner=c("DEB37", "DEB33", "DEB32", "DE121", "DE129", "DE117",
                   "DEB21", "DE121", "DE273", "DE272", "DE251", "DE263",
                   "DE262", "DE241", "DE243", "DE242", "DE244", "DE233",
                   "DE231", "DE213", "DE221", "DE223", "DEG05", "DEG0N",
                   "DE731", "DEG03"))
  
  tb_surrounded <- bind_rows(tb_surrounded, tb_surrounded2) %>% 
    bind_rows(tb_surrounded3) %>% distinct()
  
  return(tb_surrounded)
}


compute_proximity_one_year_ahead <- function(df, map, Q_with_eig, y_column) {
  
  df <- df %>% 
    mutate(y_to_convolve = 1*(!!sym(y_column)==1)) #y==1 if presence==1, y==0 if presence %in% c(1, NA)
  
  y_column <- "y_to_convolve"
  
  exponent_vec <- 0.5
  
  df_proximity <- proximity_from_Q(Q_with_eig, df, y_column, map, exponent_vec,
                                   add_year_one=FALSE)
  df_proximity <- df_proximity %>% rename(presence_previous_year=y)
  
  df_surrounded <- get_surrounded_regions()
  
  proximity_col_vec <- names(df_proximity) %>% str_subset("y_proximity")
  
  for (j in 1:nrow(df_proximity)) {
    df_curr <- df_proximity[j,]
    if (df_curr$geo %in% df_surrounded$inner) {
      geo_outer <- df_surrounded %>% filter(inner == df_curr$geo) %>% pull(outer)
      df_curr_match <- df_proximity %>% filter(geo == geo_outer, year_idx == df_curr$year_idx)
      if (df_curr_match$presence_previous_year != 1) {
        for (proximity_col in proximity_col_vec) {
          df_proximity[[proximity_col]][j] <- df_curr_match[[proximity_col]]
        }
      }
    }
  }
  
  return(df_proximity)
}


compute_proximity <- function(data, map_spde) {
  #---Compute precision matrix for the proximity diffusion process
  geo_neighs <- geo_to_neighs(map_spde)
  # plot_geo_neighs(geo_neighs)
  
  Q_besag <- compute_Q_besag(geo_neighs$nb_map, geo_neighs$n_geo, proper_version=TRUE, lambda=0.5)
  
  Q_with_eig <- collect_Q_and_eig(Q_besag)
  
  year_vec <- sort(unique(data$year_idx))
  
  df_proximity <- data %>% filter(year_idx==1) %>% select(geo, year_idx) %>%  
    mutate(y_proximity_exp050=0, presence_previous_year=0)
  
  for (curr_year in year_vec) {
    
    #---Compute proximity based on data_simulated$y for the current year
    df_proximity_curr <- compute_proximity_one_year_ahead(data %>% 
                                                            filter(year_idx==curr_year), 
                                                          map_spde, Q_with_eig, y_column="y")
    
    df_proximity <- bind_rows(df_proximity, df_proximity_curr)
    
    cat(sprintf("compute_proximity: progress %d of %d\n", curr_year, length(year_vec)))
  }
  
  df_proximity <- df_proximity %>% filter(year_idx %in% data$year_idx)
  
  return(df_proximity)
}


process_covariates <- function(data_spde, df_proximity, keys) {
  
  #---Bin the proximity covariate (covariates in data_original_covariates have already been binned)
  breaks <- c(-Inf,
              quantile(df_proximity %>% filter(year_idx>1, presence_previous_year==0) %>% 
                         pull(y_proximity_exp050), 
                       c(seq(0.5,0.9,by=0.1), 0.95)),
              Inf)
  df_proximity <- df_proximity %>% mutate(
    bin_idx_proximity050 = as.numeric(cut(y_proximity_exp050,breaks=breaks)),
    bin_idx_proximity050 = case_when(
      presence_previous_year==1 ~ 1,
      year_idx==1 ~ 2, 
      TRUE ~ bin_idx_proximity050+2
    )
  )
  
  #---Compute range of original y_proximity_exp050 for each bin (will be used for plotting)
  keys$y_proximity_exp050 <- compute_bin_keys(df_proximity, 
                                              bin_column="bin_idx_proximity050", 
                                              original_column="y_proximity_exp050",
                                              print_format="[%.2f,%.2f]")
  
  levels(keys$y_proximity_exp050[[2]])[1] <- "albo_prev_year"
  levels(keys$y_proximity_exp050[[2]])[2] <- "year1"
  
  
  
  #---Join data 
  data_spde <- data_spde %>% left_join(df_proximity) %>% 
    relocate(data_idx, geo, region_id, year_idx, date_start, CountryISO,
             presence, presence_previous_year, y_proximity_exp050, bin_idx_proximity050)
  
  
  
  #---Add year1 covariate
  data_spde <- data_spde %>% mutate(year1 = 1*(year_idx==1))
  
  
  
  return(list(data_spde=data_spde, keys=keys))
}


compute_bin_keys <- function(data, bin_column, original_column, 
                             print_format="[%.1f,%.1f]") {
  
  #---Prepare data.frame
  bin_range_column <- paste0("bin_range_", original_column)
  bin_mean_column <- paste0("bin_mean_", original_column)
  
  keys_to_bins <- tibble(bin_idx = sort(unique(data[[bin_column]])),
                         !!sym(bin_range_column) := NA,
                         bin_min = NA, bin_max = NA,
                         !!sym(bin_mean_column) := NA,
                         n_in_bin = NA)
  
  for (j in 1:nrow(keys_to_bins)) {
    
    #---Select rows in data with current bin_idx
    curr_bin_idx <- keys_to_bins$bin_idx[j]
    data_curr_bin <- data %>% filter(!!sym(bin_column) == curr_bin_idx) #filter on bin_column == curr_bin_idx
    # print(data_curr_bin %>% relocate(all_of(c(original_column, bin_column))))
    
    #---Compute range of values within current bin
    bin_range <- range(data_curr_bin[[original_column]]) #range of values in original_column for this bin
    bin_range <- round(bin_range, digits=13)
    keys_to_bins[[bin_range_column]][j] <- sprintf(print_format, bin_range[1], bin_range[2]) #make string of range
    keys_to_bins$bin_min[j] <- bin_range[1]
    keys_to_bins$bin_max[j] <- bin_range[2]
    
    #---Mean of values which lie within current bin
    keys_to_bins[[bin_mean_column]][j] <- mean(data_curr_bin[[original_column]])
    
    #---Number of rows with current bin_idx
    keys_to_bins$n_in_bin[j] <- nrow(data_curr_bin) 
    
  }
  
  keys_to_bins <- keys_to_bins %>% mutate(!!sym(bin_range_column) := 
                                            factor(!!sym(bin_range_column), 
                                                   levels=!!sym(bin_range_column), ordered=TRUE))  #make factor to keep the bin range strings sorted by numerical value when plotting  
  
  return(keys_to_bins)
}


update_f <- function(f, str_to_add) {
  f_updated <- f %>% as.character()
  
  f_updated[[3]] <- paste0(f_updated[[3]], " + ", str_to_add)
  
  f_updated <- sprintf("%s ~ %s", 
                       as.character(f_updated[[2]]), 
                       f_updated[[3]]) %>% 
    as.formula()
  
  return(f_updated)
}


process_fitted_model_bru <- function(data_spde, fitted_model, covariate_set,
                                     spde_model, spde_mesh) {
  
  #---Process SPDE fit
  year_idx_vec <- sort(unique(data_spde$year_idx))
  
  tb_regions <- data_spde %>% select(geo, region_id) %>% distinct()
  data_spde_with_na <- 
    tidyr::crossing(tb_regions, tibble(year_idx=unique(data_spde$year_idx))) %>% 
    arrange(year_idx, region_id) %>% left_join(data_spde)
  
  map_full_model <- NULL
  
  s_post <- predict(fitted_model, as_Spatial(data_spde), ~ s)
  
  for (year_idx__ in year_idx_vec) {
    
    map_full_model <- bind_rows(map_full_model, s_post %>% as_tibble() %>% filter(year_idx == year_idx__) %>% 
                                  select(y=presence, geo, region_id, year_idx, spde_projected=mean))
  }
  
  
  
  #---Subset results
  idx_subset <- which(data_spde$year_idx %in% year_idx_vec)
  
  map_full_model$linear_predictor.mean <- fitted_model$summary.linear.predictor[idx_subset, "mean"]
  map_full_model$linear_predictor.sd <- fitted_model$summary.linear.predictor[idx_subset, "sd"]
  map_full_model$p.mean <- fitted_model$summary.fitted.values[idx_subset, "mean"]
  map_full_model$p.sd <- fitted_model$summary.fitted.values[idx_subset, "sd"]
  
  
  #---Main summaries
  results_summary <- tibble(DIC = round(fitted_model$dic$dic,2), #goodness of fit: DIC
                            logscore = round(-mean(log(fitted_model$cpo$cpo), na.rm = T), 3), #goodness of fit: logscore
                            WAIC = round(fitted_model$waic$waic, 2),
                            fixed_effects = list(fitted_model$summary.fixed), #all fixed effect parameters
                            intercept = fitted_model$summary.fixed["Intercept","mean"], #intercept
                            hyperpar = list(fitted_model$summary.hyperpar)) #fitted hyper parameters
  
  
  
  #---Random effects, constant in time
  #---Initialize
  functions_random_effects <- list()
  map_random_effects <- data_spde %>% as_tibble() %>% filter(year_idx==1) %>% 
    select(data_idx, geo, region_id) #these random effects are constant in time so enough to select first year
  
  #---Add random effects
  #---Random effects for covariates constant in time
  if (covariate_set==1) {
    
    out <- add_random_effect_bru(map_random_effects, functions_random_effects,
                                 random_effect_col="re_mechanistic",
                                 covariate_col="bin_idx_mean_adults_pow_merged")
    map_random_effects <- out$map_random_effects
    functions_random_effects <- out$functions_random_effects
  } else if (covariate_set==2) {
    
    covariate_vec <- c("bin_idx_median_temp", "bin_idx_min_temp_CROSS",
                       "bin_idx_median_relative_humidity", "bin_idx_log_pop_merged")
    random_effect_vec <- c("re_median_temp", "re_min_temp_CROSS",
                           "re_median_relative_humidity", "re_log_population")
    
    for (j in 1:length(random_effect_vec)) {
      out <- add_random_effect_bru(map_random_effects, functions_random_effects, 
                                   random_effect_col=random_effect_vec[j],
                                   covariate_col=covariate_vec[j])
      map_random_effects <- out$map_random_effects
      functions_random_effects <- out$functions_random_effects
    }
  }
  
  
  
  #---Random effect for time-varying covariate
  random_effect_col <- "re_proximity"
  covariate_col <- "bin_idx_proximity050"
  
  functions_random_effects[[covariate_col]] <-
    tibble(!!covariate_col := fitted_model$summary.random[[random_effect_col]]$ID,
           mean = fitted_model$summary.random[[random_effect_col]]$mean,
           sd = fitted_model$summary.random[[random_effect_col]]$sd,
           lower_bound = fitted_model$summary.random[[random_effect_col]]$`0.025quant`,
           upper_bound = fitted_model$summary.random[[random_effect_col]]$`0.975quant`)
  
  random_effect_str <- paste0("reffect_", covariate_col)
  
  map_random_effects_curr <-
    left_join(data_spde %>% as_tibble() %>% select(data_idx, region_id, year_idx, !!sym(covariate_col)),
              functions_random_effects[[covariate_col]] %>%
                rename(!!paste0(random_effect_str, ".mean") := mean,
                       !!paste0(random_effect_str, ".sd") := sd,
                       !!paste0(random_effect_str, ".lower_bound") := lower_bound,
                       !!paste0(random_effect_str, ".upper_bound") := upper_bound))
  map_random_effects_with_time <- left_join(data_spde %>% as_tibble() %>% select(data_idx, geo, region_id, year_idx), 
                                            map_random_effects_curr)
  
  
  
  #---Map full model
  map_full_model$y_orig <- data_spde$y_orig
  map_full_model$training_set <- data_spde$training_set
  map_full_model$test_set <- data_spde$test_set
  
  #---Abs error and deviance
  map_full_model$error <- compute_error(map_full_model$p.mean, map_full_model$y, n=1)
  results_summary$total_abs_error <- sum(map_full_model$error, na.rm=TRUE)
  
  map_full_model$deviance <- compute_bernoulli_deviance(map_full_model$p.mean, map_full_model$y)
  results_summary$total_deviance <- sum(map_full_model$deviance, na.rm=TRUE)
  
  if (sum(map_full_model$test_set)==0) {
    
    warning("test set is empty")
    
    map_full_model$error_cv <- NA
    map_full_model$deviance_cv <- NA
    results_summary$total_abs_error_cv <- NA
    results_summary$total_deviance_cv <- NA
  } else {
    
    map_full_model$error_cv <- 
      compute_error(map_full_model$p.mean, map_full_model$y_orig)
    results_summary$total_abs_error_cv <- 
      sum(map_full_model %>% filter(test_set==TRUE) %>% pull(error_cv), na.rm=TRUE)
    
    map_full_model$deviance_cv <- 
      compute_bernoulli_deviance(map_full_model$p.mean, map_full_model$y_orig)
    results_summary$total_deviance_cv <- 
      sum(map_full_model %>% filter(test_set==TRUE) %>% pull(deviance_cv), na.rm=TRUE)
  }
  
  
  
  return(list(results_summary=results_summary,
              map_full_model=map_full_model,
              map_random_effects=map_random_effects,
              map_random_effects_with_time=map_random_effects_with_time,
              functions_random_effects=functions_random_effects))
}


add_random_effect_bru <- function(map_random_effects, functions_random_effects, random_effect_col,
                                  covariate_col) {
  
  functions_random_effects[[covariate_col]] <- 
    tibble(!!covariate_col := fitted_model$summary.random[[random_effect_col]]$ID,
           mean = fitted_model$summary.random[[random_effect_col]]$mean,
           sd = fitted_model$summary.random[[random_effect_col]]$sd,
           lower_bound = fitted_model$summary.random[[random_effect_col]]$`0.025quant`,
           upper_bound = fitted_model$summary.random[[random_effect_col]]$`0.975quant`)
  
  random_effect_str <- paste0("reffect_", covariate_col)
  
  map_random_effects_curr <- 
    left_join(data_spde %>% as_tibble() %>% select(data_idx, region_id, !!sym(covariate_col)),
              functions_random_effects[[covariate_col]] %>% 
                rename(!!paste0(random_effect_str, ".mean") := mean,
                       !!paste0(random_effect_str, ".sd") := sd,
                       !!paste0(random_effect_str, ".lower_bound") := lower_bound,
                       !!paste0(random_effect_str, ".upper_bound") := upper_bound))
  map_random_effects <- left_join(map_random_effects, map_random_effects_curr)
  
  return(list(map_random_effects=map_random_effects,
              functions_random_effects=functions_random_effects))
}


plot_on_map_selected_model_parts_bru <- function(map_random_effects, 
                                                 map_random_effects_with_time,
                                                 map_full_model, 
                                                 map_spde,
                                                 year_to_plot=9,
                                                 plot_larger=FALSE) {
  
  ps_map <- list()
  
  if (!is.null(map_random_effects) && nrow(map_random_effects)>0) {
    
    if ("reffect_bin_idx_median_temp.mean" %in% names(map_random_effects)) {
      #---Plot median temp random effect (on map)
      ps_map[[length(ps_map)+1]] <- 
        plot_sf(map_random_effects %>% left_join(map_spde), 
                "reffect_bin_idx_median_temp.mean",
                title="RE, median temp.",
                legend_title="median temp.",
                plot_larger=plot_larger)
    }
    
    if ("reffect_bin_idx_min_temp_CROSS.mean" %in% names(map_random_effects)) {
      map_random_effects <- map_random_effects %>% 
        mutate(reffect_bin_idx_min_temp_CROSS.meanNA=
                 if_else(bin_idx_min_temp_CROSS==1, NA,
                         reffect_bin_idx_min_temp_CROSS.mean))
      #---Plot min temp CROSS random effect (on map)
      ps_map[[length(ps_map)+1]] <- 
        plot_sf(map_random_effects %>% left_join(map_spde), 
                "reffect_bin_idx_min_temp_CROSS.meanNA",
                title="RE, min. temp. CROSS",
                legend_title="min. temp. CROSS",
                plot_larger=plot_larger)
    }
    
    if ("reffect_bin_idx_median_relative_humidity.mean" %in% names(map_random_effects)) {
      #---Plot median relative humidity random effect (on map)
      ps_map[[length(ps_map)+1]] <- 
        plot_sf(map_random_effects %>% left_join(map_spde), 
                "reffect_bin_idx_median_relative_humidity.mean",
                title="RE, median relative humidity",
                legend_title="median rel. humidity",
                plot_larger=plot_larger)
    }
    
    if ("reffect_bin_idx_log_pop_merged.mean" %in% names(map_random_effects)) {
      #---Plot log population random effect (on map)
      ps_map[[length(ps_map)+1]] <- 
        plot_sf(map_random_effects %>% left_join(map_spde),
                "reffect_bin_idx_log_pop_merged.mean",
                title="RE, log population",
                legend_title="log pop.",
                plot_larger=plot_larger)
    }
    
    if ("reffect_bin_idx_mean_adults_pow_merged.mean" %in% names(map_random_effects)) {
      #---Plot mechanistic model random effect (on map)
      ps_map[[length(ps_map)+1]] <- 
        plot_sf(map_random_effects %>% left_join(map_spde), 
                "reffect_bin_idx_mean_adults_pow_merged.mean",
                title="RE, mechanistic model",
                legend_title="mech. model",
                plot_larger=plot_larger)
    }
    
  }
  
  
  
  if (!is.null(map_full_model$spde_projected)) {
    
    #---Plot spatio-temporal field
    ps_map[[length(ps_map)+1]] <- 
      plot_sf(map_full_model %>% left_join(map_spde) %>% 
                filter(year_idx==year_to_plot), "spde_projected",
              title=sprintf("RE, spatio-temporal field, year %d", year_to_plot),
              legend_title="SPDE",
              plot_larger=plot_larger)
  }
  
  
  
  #---Plot proximity random effect
  ps_map[[length(ps_map)+1]] <- 
    plot_on_map_proximity_random_effect(map_random_effects_with_time, map_spde, 
                                        special_group_bin_idx=NULL,
                                        year_to_plot=year_to_plot,
                                        plot_larger=plot_larger)
  
  
  
  #---Plot fitted model probability (p.mean) and the data (y)
  #---Fitted probability
  ps_map[[length(ps_map)+1]] <- 
    plot_sf(map_full_model %>% filter(year_idx==year_to_plot) %>% 
              left_join(map_spde), 
            "p.mean",
            title=sprintf("prob., year %d", year_to_plot),
            color_transformation=power_transformation(0.1),
            legend_title="prob.",
            plot_larger=plot_larger)
  #---Fitted probability, only showing new observations
  ps_map[[length(ps_map)+1]] <- 
    plot_sf(map_full_model %>% filter(year_idx==year_to_plot) %>% 
              left_join(data_spde %>% as_tibble() %>% select(-geometry)) %>%  
              mutate(p.mean.new_obs=ifelse(presence_previous_year==1,NA,p.mean)) %>% # set to NA if Ae. albopictus was observed in the region the previous year
              left_join(map_spde), 
            "p.mean.new_obs",
            title=sprintf("prob., new obs., year %d", year_to_plot),
            legend_title="prob., new obs.",
            color_transformation=power_transformation(0.1),
            plot_larger=plot_larger)
  #---Data, current year
  ps_map[[length(ps_map)+1]] <- 
    plot_sf(map_full_model %>% filter(year_idx==year_to_plot) %>% 
              left_join(map_spde), 
            "y",
            title=sprintf("simulated data, year %d", year_to_plot),
            plot_larger=plot_larger,
            palette_idx=1)
  
  if (year_to_plot > 1) {
    #---Data, previous year
    ps_map[[length(ps_map)+1]] <- 
      plot_sf(map_full_model %>% filter(year_idx==(year_to_plot-1)) %>% 
                left_join(map_spde), 
              "y",
              title=sprintf("simulated data, year %d", year_to_plot-1),
              plot_larger=plot_larger,
              palette_idx=1)
  }
  
  #---Collect figures into one
  n_figures <- length(ps_map)
  ncol <- floor(sqrt(n_figures))
  nrow <- ceiling(n_figures / ncol)
  p <- do.call(ggarrange, c(ps_map, nrow=nrow, ncol=ncol)) #ggarrange(ps_map, nrow=nrow, ncol=ncol)
  
  return(list(p=p, ps_map=ps_map))
}


plot_proximity_random_effect_bru <- function(functions_random_effects, keys, data, 
                                             filename=NULL,
                                             plot_rug_vector=FALSE,
                                             plot_larger=FALSE,
                                             ylim=NULL,
                                             resolution_scaling_width=1) {
  
  #---Plot proximity random effect (as a function)
  bin_idx_proximity_col <- names(functions_random_effects) %>% str_subset("proximity")
  proximity_exp <- str_extract(bin_idx_proximity_col, "(?<=proximity)[0-9].*")
  keys_col <- str_subset(names(keys), paste0("exp", proximity_exp))
  
  #---Prepare for plotting
  reffect_proximity <- functions_random_effects[[bin_idx_proximity_col]]
  
  #---Add bin range and mean of original proximity
  reffect_proximity <- left_join(reffect_proximity, keys[[keys_col]],
                                 by = join_by(!!sym(bin_idx_proximity_col) == bin_idx))
  
  #---Plot with x_column="bin_mean", and make bin position in bottom histogram correspond to the range of values within each bin 
  if (plot_rug_vector) {
    rug_vector <- data$log_population
  } else {
    rug_vector <- NULL
  }
  
  re_plots_proximity <- 
    linked_random_effect_and_histogram_plot(reffect_proximity, 
                                            x_column_type="bin_mean", 
                                            bin_width_constant=FALSE,
                                            vector_to_plot_with_rug=rug_vector,
                                            plot_larger=plot_larger,
                                            ylim=ylim)
  # grid.draw(re_plots_max_temp$g)
  print(reffect_proximity)
  
  if (!is.null(filename)) {
    #---Save plot
    save_to_png(re_plots_proximity$g, filename=filename, resolution_scaling_width=resolution_scaling_width)
  }
  
  return(re_plots_proximity$g)
}


plot_geometry <- function(map, color="yellow", opacity=0.5, add=FALSE) {
  plot(st_geometry(map %>% pull(geometry)), add=add, col=scales::alpha(color, opacity))
}


plot_sf <- function(map, feature, title = "", subtitle = "",
                    legend_title = feature,
                    opacity = 0.8,
                    xlim = NA, ylim = NA,
                    palette_idx = 5,
                    manual_palette = NULL,
                    background_color = "white",
                    edge_color = "black",
                    invert_colors = FALSE,
                    color_limits = NULL,
                    color_transformation = NULL,
                    interactive = FALSE,
                    crop_plot_to_central_Europe = FALSE,
                    strict_cropping = FALSE,
                    title_size = 12,
                    legend_title_length = 25,
                    theme_paper = TRUE,
                    plot_larger = FALSE) {
  
  if (plot_larger) {
    legend_title_length <- 9
    opacity <- 1
  }
  
  cat(sprintf("plotting feature: %s\n", feature))
  #---Compute x- and y-limits of the map
  if (!crop_plot_to_central_Europe) {
    
    if (any(is.na(xlim))) {
      
      lims <- compute_boundary_of_sf(map, compute_x = TRUE, compute_y = FALSE)
      xlim <- lims$xlim
    }
    if (any(is.na(ylim))) {
      
      lims <- compute_boundary_of_sf(map, compute_x = FALSE, compute_y = TRUE)
      ylim <- lims$ylim
    }
  } else {
    
    cat("cropping plot to central Europe (approx. longitude: [-10, 30], latitude: [35, 60])\n") #this is the regions where I have ODE simulations
    # xlim <- c(-10, 30)
    # ylim <- c(35, 60)
    xlim <- c(-10, 27.5)
    ylim <- c(35.5, 58)
    
    if (strict_cropping) {
      warning("using strict cropping, will take some time")
      crop_factor <- st_bbox(c(xmin = xlim[1], 
                               xmax = xlim[2], 
                               ymax = ylim[1], 
                               ymin = ylim[2]),
                             crs = st_crs(map$geometry[1]))
      
      geometry_cropped_lst <- rep(NA, nrow(map))
      for (j in 1:length(map$geometry)) {
        
        geometry_cropped <- st_crop(map$geometry[j], crop_factor)
        geometry_cropped_lst[j] <- ifelse(length(geometry_cropped)==1,
                                          geometry_cropped, NA) 
      }
      
      idx_has_geometry <- which(!is.na(geometry_cropped_lst))
      geometry_cropped_lst <- geometry_cropped_lst[idx_has_geometry]
      map <- map[idx_has_geometry,]
      map$geometry <- st_as_sfc(geometry_cropped_lst)
      
    }
  }
  
  #---Plot labels
  len <- nchar(legend_title)
  if (len > legend_title_length) {
    legend_title <- substr(legend_title, 1, legend_title_length)
  } else if (len > legend_title_length) {
    legend_title <- paste0(legend_title, strrep(" ", legend_title_length-len))
  }
  
  plot_labs <- labs(title = title, fill = legend_title)
  
  
  
  #---Number of colors to plot
  n_cols <- length(setdiff(unique(map[[feature]]), NA))
  n_discrete <- n_cols
  if (n_cols <= 3) {
    
    map[[feature]] <- map[[feature]] %>% as.factor()
  }
  if (is.factor(map[[feature]]) || is.logical(map[[feature]]) ||
      is.character(map[[feature]])) {
    
    is_continuous <- FALSE
    map[[feature]] <- map[[feature]] %>% as.factor()
    
    if (n_cols > 25) {
      
      warning("plotting a discrete factor with many unique elements")
    }
  } else {
    
    is_continuous <-TRUE
  }
  if (!is.na(n_discrete) && n_discrete==0) 
    n_discrete <- 1
  
  
  #---Colormap
  if (!is.null(manual_palette)) {
    cols <- color_palette(manual_palette = manual_palette,
                          invert_colors = invert_colors,
                          n_discrete = n_discrete,
                          is_continuous = is_continuous,
                          color_limits = color_limits,
                          color_transformation = color_transformation)
  } else {
    cols <- color_palette(palette_idx, 
                          n_discrete = n_discrete,
                          is_continuous = is_continuous,
                          invert_colors = invert_colors,
                          color_limits = color_limits,
                          color_transformation = color_transformation)
  }
  col <- cols$scale$fill
  
  
  #---Plotting
  if (!interactive) {
    #---Non-interactive version
    g <- ggplot(map, aes(fill = !!sym(feature), geometry = geometry)) + 
      geom_sf(color = alpha(edge_color,1/3), #edges
              alpha = opacity) #fill
  } else {
    #---Interactive version, use with make_interactive
    g <- ggplot(map, aes(fill = !!sym(feature), geometry = geometry)) + 
      geom_sf_interactive(color = alpha(edge_color,1/3), #edges
                          alpha = opacity,
                          aes(tooltip = !!sym(feature), data_id = !!sym(feature))) #fill
  }
  
  p <- g +
    col +
    coord_sf(xlim=xlim, ylim=ylim) +
    plot_labs +
    theme(panel.background = element_rect(fill = background_color)) 
  
  p <- p + theme(plot.title = element_text(size=title_size))
  
  if (theme_paper) {
    p <- p + theme_bw()
  }
  
  if (plot_larger) {
    p <- p + theme(legend.key.width = unit(1.25, "cm"),
                   legend.key.height = unit(1.5, "cm")) + 
      theme(text = element_text(size = 24))
  }
  
  if (n_discrete > 50 && !is_continuous) {
    warning("not showing legend, too many unique elements")
    p <- p + theme(legend.position = "none")
  }
  
  
  return(p)
}


compute_boundary_of_sf <- function(map, compute_x = TRUE, compute_y = TRUE) {
  if (compute_x) {
    xmin <- Inf; xmax <- -Inf
    for (j in seq_along(map$geometry)) {
      b <- st_bbox(map[[j,"geometry"]])
      xmin <- min(xmin, b$xmin)
      xmax <- max(xmax, b$xmax)
    }
    xlim <- c(xmin, xmax)
  } else {
    xlim <- c(NaN, NaN)
  }
  
  if (compute_y) {
    ymin <- Inf; ymax <- -Inf
    for (j in seq_along(map$geometry)) {
      b <- st_bbox(map[[j,"geometry"]])
      ymin <- min(ymin, b$ymin)
      ymax <- max(ymax, b$ymax)
    }
    ylim <- c(ymin, ymax)
  } else {
    ylim <- c(NaN, NaN)
  }
  return(list(xlim = xlim, ylim = ylim))
}


#---Plot graph saved as the output from the function geo_to_neighs
#---use as: geo_neighs <- geo_to_neighs(map); plot_geo_neighs(geo_neighs)
plot_geo_neighs <- function(geo_neighs, filename = "", selection_nodes = NULL,
                            node_col = "red", resolution_scaling = 1) {
  
  #---Change coordinate class for plotting
  coords_sp <- SpatialPoints(geo_neighs$coords)
  
  #---Save to png if filename was provided
  if (filename != "") {
    cat(sprintf("saving plot to file: %s\n", filename))
    
    png(filename, 
        width=1200 * resolution_scaling, 
        height=800 * resolution_scaling)
  }
  
  #---Plot geometries if geo_neighs contains shapes
  if ("shapes" %in% names(geo_neighs)) {
    plot(geo_neighs$shapes, border = gray(.5)) 
    add <- TRUE
  } else {
    add <- FALSE
  }
  
  #---Plot nodes and edges of graph in nb_map
  plot(geo_neighs$nb_map, geo_neighs$coords, add = add) 
  plot(coords_sp, add = TRUE, col = node_col, pch = 20, cex = 0.95) 
  
  #---Highlight nodes in selection_nodes
  if (!is.null(selection_nodes))
    plot(coords_sp[selection_nodes], add = TRUE, col = "green", pch = 20, cex = 0.75)
  
  #---Close connection to png file
  if (filename != "") 
    dev.off()
}


plot_network <- function(links, n_nodes, node_coords,
                         node_weights=NULL, link_weights=NULL,
                         link_color="tomato",
                         sc_link_width=10, sc_node_size=6,
                         title="") {
  
  nodes <- 1:n_nodes
  
  net <- graph.data.frame(links, nodes, directed=T)
  
  if (!is.null(node_weights)) {
    V(net)$size <- node_weights / max(node_weights) * sc_node_size
  } else {
    V(net)$size <- sc_node_size
  }
  if (!is.null(link_weights)) {
    E(net)$width <- link_weights / max(link_weights) * sc_link_width
  } else {
    E(net)$width <- sc_link_width
  }
  
  plot(net, layout = node_coords, 
       edge.arrow.size = 0.05, edge.curved = 0.1,
       edge.color = link_color,
       vertex.label = NA,
       main = title)
}


# Use as: g <- align_plots(p1, p2); grid.draw(g)
align_plots <- function(p_above, p_below, ratio_above=8) {
  g_above <- ggplotGrob(p_above)
  g_below <- ggplotGrob(p_below)
  
  g_below <- gtable_add_cols(g_below, unit(0,"mm"))
  g_above$widths <- unit.pmax(g_above$widths, g_below$widths)
  g_below$widths <- unit.pmax(g_above$widths, g_below$widths)
  
  g <- rbind(g_above, g_below, size = "first")
  g$widths <- unit.pmax(g_above$widths, g_below$widths)
  g <- resize_grob_heights(g, c(ratio_above,1))
  
  return(g)
}

resize_grob_heights <- function(g, heights = rep(1, length(idpanels))){
  idpanels <- unique(g$layout[grepl("panel",g$layout$name), "t"])
  g$heights[idpanels] <- unit.c(do.call(unit, list(heights, 'null')))
  g
}


plot_random_effects_out <- function(tb_reffects_all) {
  
  title_size <- 12
  
  plots3 <- list()
  ps_re <- list()
  y_range <- c(Inf, -Inf)
  for (plot_round in 1:2) {
    for (k in 1:nrow(tb_reffects_all)) {
      reffect_name <- tb_reffects_all$reffect[k]
      tb_long <- tb_reffects_all$tb_long[[k]]
      
      df_hist_presence01 <- tb_reffects_all$df_hist_presence01[[k]]
      
      if (plot_round==2) {
        if (!grepl("x_cond", reffect_name)) {
          epsilon <- (y_range[2]-y_range[1]) * 0.025
          y_range_curr <- c(y_range[1]-epsilon, y_range[2]+epsilon)
        } else {
          n_bins <- max(tb_long$x)
          tb_long <- tb_long %>% filter(x != n_bins)
          y_range_curr <- range(tb_long$y)
          epsilon <- (y_range_curr[2]-y_range_curr[1]) * 0.025
          y_range_curr <- c(y_range_curr[1]-epsilon, y_range_curr[2]+epsilon)
          df_hist_presence01 <- df_hist_presence01 %>% filter(!!sym(reffect_name) != n_bins)
        }
        
      } else if (plot_round==1) {
        
        y_range_curr <- range(tb_long$y)
        if (!grepl("x_cond", reffect_name)) {
          y_range[1] <- min(y_range[1], y_range_curr[1])
          y_range[2] <- max(y_range[2], y_range_curr[2])
        }
        
      }
      
      main_plot <- 
        plot_scatter_lines(tb_long,
                           x_column = "x", y_column = "y",
                           color_column = "type",
                           title = paste0("re: ", reffect_name),
                           title_size = title_size,
                           plot_lines = TRUE,
                           n_cutoff = 0,
                           is_continuous = FALSE,
                           ylim = y_range_curr) 
      xlimits <- layer_scales(main_plot)$x$range$range
      hist <- ggplot(df_hist_presence01, aes_string(x = reffect_name, y = "count")) +
        geom_bar(stat="identity") + coord_cartesian(xlim = xlimits) + theme_classic() +
        theme(axis.text.x=element_blank(),
              axis.ticks.x=element_blank(),
              axis.title.x=element_blank(),
        )
      
      g_top <- ggplotGrob(main_plot)
      g_bottom <- ggplotGrob(hist)
      
      g_bottom <- gtable_add_cols(g_bottom, unit(0,"mm"))
      g_top$widths <- unit.pmax(g_top$widths, g_bottom$widths)
      g_bottom$widths <- unit.pmax(g_top$widths, g_bottom$widths)
      
      g <- rbind(g_top, g_bottom, size = "first")
      g$widths <- unit.pmax(g_top$widths, g_bottom$widths)
      g <- resize_grob_heights(g, c(8,1))
      
      if (plot_round==2) {
        plots3[[length(plots3)+1]] <- g
      } else if (plot_round==1) {
        ps_re[[length(ps_re)+1]] <- g
      }
      
    }
    
    # if (!is.null(file_pdf) && plot_round==1) {
    #   options <- save_to_pdf_via_png0(step = 2, options) 
    #   n_col <- floor(sqrt(length(plots3)))
    #   figure3 <- do.call(grid.arrange, c(plots3, ncol = n_col)) %>% print()
    # }
  }
  
  return(list(plots3=plots3, ps_re=ps_re))
}


color_palette <- function(palette_idx = 1, 
                          manual_palette = NULL,
                          is_continuous = TRUE,
                          n_discrete = 10,
                          invert_colors = FALSE,
                          color_limits = NULL,
                          color_transformation = NULL) {
  
  if (!is.null(manual_palette)) {
    
    col_palette <- manual_palette
  } else {
    
    if (!palette_idx %in% 1:6)
      stop("palette_idx should be an integer between 1 and 6")
    
    #---Extract color palette
    if (palette_idx == 1) {
      
      col_palette <- brewer.pal(11, "RdYlBu")
      col_palette <- col_palette[seq(length(col_palette),1,-1)]
    } else if (palette_idx == 2) {
      
      if (is_continuous) {
        
        col_palette <- jetColors(500)
      } else {
        
        col_palette <- jetColors(n_discrete)
      }
    } else if (palette_idx == 3) {
      
      col_palette <- paletteer_c("grDevices::Blues 3", 30)
    } else if (palette_idx == 4) {
      
      col_palette <- brewer.pal(12, "Paired")
      if (!is_continuous) {
        
        col_palette <- rep(col_palette, length.out = n_discrete)
      }
    } else if (palette_idx == 5) {
      
      if (!is_continuous) {
        
        n <- n_discrete
        if (n>7) {
          
          n <- n+2
          col_palette <- pals::ocean.balance(n)
          col_palette <- col_palette[2:(n-1)]
        } else if (n>3) {
          
          n <- n*4
          i <- seq(1,n-1,by=4)
          i[length(i)] <- n-2
          col_palette <- pals::ocean.balance(n)
          col_palette <- col_palette[i]
        } else {
          
          col_palette <- pals::ocean.balance(n)
        }
      } else {
        
        n <- 34
        col_palette <- pals::ocean.balance(n)
        col_palette <- col_palette[3:(n-3)]
      }
    } else if (palette_idx == 6) {
      
      col_palette <- brewer.pal(9, "Greys")
    }
    
    
    #---Invert colors
    if (invert_colors) {
      col_palette <- col_palette[seq(length(col_palette),1,-1)]
    }
  }
  
  
  #---Compute fill and col
  if (is_continuous) {
    
    cat("Continuous color scale\n")
    
    #Continuous color scale
    if (!is.null(color_transformation) && !is.null(color_limits))
      stop("not implemented")
    
    if (!is.null(color_transformation)) {
      scale_fill <- scale_fill_gradientn(colours = col_palette, trans = color_transformation)
      scale_col <- scale_color_gradientn(colours = col_palette, trans = color_transformation)
    } else {
      if (is.null(color_limits)) {
        scale_fill <- scale_fill_gradientn(colours = col_palette)
        scale_col <- scale_color_gradientn(colours = col_palette)
      } else {
        scale_fill <- scale_fill_gradientn(colours = col_palette, 
                                           limits = color_limits)
        scale_col <- scale_color_gradientn(colours = col_palette, 
                                           limits = color_limits)
      }
    }
  } else {
    
    cat("Discrete color scale\n")
    
    if (!is.null(color_transformation)) {
      cat("Color transformation is only implemented for continuous color scales\n")
    }
    
    #Discrete color scale
    col_palette <- colorRampPalette(col_palette)(n_discrete)
    
    scale_fill <- scale_fill_manual(values = col_palette)
    scale_col <- scale_color_manual(values = col_palette)
  }
  
  
  return(list(scale = list(fill = scale_fill, 
                           col = scale_col), 
              col_palette = col_palette))
}


make_interactive <- function(p) {
  
  #---Plot p interactively, fill with cyan on hovering
  girafe(ggobj = p) %>%
    girafe_options(opts_hover(css = "fill:cyan;"))
}


# manual palette example 1: 
# manual_palette <- color_palette(palette_idx=6)
# manual_palette <- manual_palette$col_palette
# manual palette example 2:
# col_rgb <- col2rgb("gray40")
# col_hex <- rgb(col_rgb[1], col_rgb[2], col_rgb[3], maxColorValue=255)
# manual_palette <- c(col_hex, NA, col_hex)
# col_rgb <- col2rgb("gray5")
# col_hex <- rgb(col_rgb[1], col_rgb[2], col_rgb[3], maxColorValue=255)
# manual_palette[2] <- col_hex
plot_scatter_lines <- function(tb, x_column, y_column,
                               color_column = NULL,
                               facet_column = NULL,
                               size_column = NULL,
                               point_label_column = NULL,
                               label_nudge = 0,
                               x_segment_columns = NULL,
                               y_segment_columns = NULL,
                               interactive_column = NULL,
                               plot_scatter = TRUE,
                               plot_lines = TRUE,
                               plot_smooth = FALSE,
                               plot_ribbon = FALSE,
                               plot_contours = FALSE,
                               plot_segments = FALSE,
                               plot_marginals = FALSE,
                               plot_diagonal = FALSE,
                               plot_constant_lines_y = NULL,
                               plot_constant_lines_x = NULL,
                               n_cutoff = 0,
                               n_contours = 10,
                               n_marginal_bins = 100,
                               smooth_gam_param = 2,
                               smooth_span = NULL, #0.1
                               smooth_standard_span = NULL,
                               smoothing_method = NULL, # NULL or "gam"
                               display_smooth_se = TRUE,
                               add_segment_arrows = FALSE,
                               point_size = 2,
                               size_scaling_offset = 0.5,
                               line_size = 0.8,
                               palette_idx = 14, #discrete 18, continuous 9, diverging 14
                               manual_palette = NULL,
                               opacity = 1, #0.8
                               invert_colors = FALSE,
                               cap_colors = NULL,
                               facet_free_x = FALSE, facet_free_y = FALSE,
                               logx = FALSE, logy = FALSE,
                               log_transform = FALSE,
                               color_transformation = NULL,
                               axis_transformation_x = NULL,
                               axis_transformation_y = NULL,
                               xlim = NULL, ylim = NULL,
                               title = "",
                               title_size = 15,
                               label_legend = NULL,
                               label_x = NULL,
                               label_y = NULL,
                               nchar_legend_title = 10,
                               n_legend_cutoff = 30,
                               is_continuous = NULL,
                               coord_flip = FALSE,
                               aspect_ratio = NULL,
                               plot_larger = FALSE,
                               remove_alpha_from_legend = TRUE) {
  # if (plot_larger) {
  #   nchar_legend_title <- 9
  # }
  #---Setup
  if (nrow(tb) > n_cutoff && n_cutoff != 0)
    warning("nrow(tb) > n_cutoff")
  
  if (!is.null(plot_constant_lines_y) || !is.null(plot_constant_lines_x)) {
    plot_constant_line <- TRUE
  }
  
  if (log_transform) { 
    if (logx && min(tb[[x_column]]) <= 0) {
      tb[[x_column]] <- apply(cbind(tb[[x_column]],0), 1, max) + 0.5
      title <- paste0(title, ", log transf. x")
      transformed_x <- TRUE
      cat("some x-values are <= 0, transforming for log scale\n")
    } else {
      transformed_x <- FALSE
    }
    if (logy && min(tb[[y_column]]) <= 0) {
      tb[[y_column]] <- apply(cbind(tb[[y_column]],0), 1, max) + 0.5
      title <- paste0(title, ", log transf. y")
      transformed_y <- TRUE
      cat("some y-values are <= 0, transforming for log scale\n")
    } else {
      transformed_y <- FALSE
    }
    if (transformed_x || transformed_y) {
      if (plot_diagonal) {
        cat(paste0("plot_diagonal is incompatible with log_transform", 
                   ", setting plot_diagonal to FALSE\n"))
        plot_diagonal <- FALSE
      }
      if (!is.null(plot_constant_line)) {
        cat(paste0("plot_constant_line is incompatible with log_transform", 
                   ", setting plot_constant_line to FALSE\n"))
        plot_constant_line <- NULL
      }
      
    }
  }
  
  if (!is.null(color_column)) {
    color <- color_column
    if (length(unique(tb[[color_column]])) == 1) {
      tb[[color_column]] <- tb[[color_column]] %>% as.factor()
    }
    
    if (is.null(is_continuous)) {
      if (is.factor(tb[[color_column]]) || is.logical(tb[[color_column]]) ||
          is.character(tb[[color_column]])) {
        is_continuous = FALSE
      } else {
        is_continuous = TRUE
      }
    } else if (!is_continuous && 
               !(is.factor(tb[[color_column]]) || is.logical(tb[[color_column]]) ||
                 is.character(tb[[color_column]]))) {
      warning("turning column into factor column")
      
      tb[[color_column]] <- as.factor(tb[[color_column]])
    }
    
    if (is.null(label_legend)) {
      title_legend <- color_column
    } else {
      title_legend <- label_legend
    }
    n <- nchar(title_legend)
    if (n < nchar_legend_title) {
      str_spaces <- strrep(" ", nchar_legend_title - n)
      title_legend <- paste0(title_legend, str_spaces)
    } else if (n > nchar_legend_title) {
      title_legend <- substr(title_legend, 1, nchar_legend_title)
    }
  }
  
  if (plot_contours) {
    if (is.factor(tb[[x_column]])) {
      stop("x is a factor, not compatible with contours")
      plot_contours <- FALSE
    }
    if (is.factor(tb[[y_column]])) {
      stop("y is a factor, not compatible with contours")
      plot_contours <- FALSE
    }
    
    #geom_density_2d doesn't work if quantiles are not unique
    if (is.null(color_column)) {
      xq <- quantile(tb[[x_column]], c(0.25,0.5,0.75), na.rm = TRUE)
      if (length(unique(xq)) < 3) {
        warning("x-quantiles are not unique, not plotting contours")
        plot_contours <- FALSE
        title <- paste0(title, "__non-unique quantiles of x")
      }
      
      yq <- quantile(tb[[y_column]], c(0.25,0.5,0.75), na.rm = TRUE)
      if (length(unique(yq)) < 3) {
        warning("y-quantiles are not unique, not plotting contours")
        plot_contours <- FALSE
        title <- paste0(title, "__non-unique quantiles of y")
      }
      
    } else {
      color_vals <- unique(tb[[color_column]])
      for (color_val0 in color_vals) {
        tb_group <- tb %>% filter(!!sym(color_column) == color_val0)
        xq <- quantile(tb_group[[x_column]], c(0.25,0.5,0.75), na.rm = TRUE)
        if (length(unique(xq)) < 3) {
          str_color_val <- sprintf("color %s: %s", color_column, color_val0)
          warning(paste0("x-quantiles are not unique, ",
                         str_color_val, 
                         ", not plotting contours\n"))
          plot_contours <- FALSE
          title <- paste0(title, "__non-unique quantiles of x",
                          str_color_val)
        }
        
        yq <- quantile(tb_group[[y_column]], c(0.25,0.5,0.75), na.rm = TRUE) 
        if (length(unique(yq)) < 3) {
          str_color_val <- sprintf("color %s: %s", color_column, color_val0)
          warning(paste0("y-quantiles are not unique, ",
                         str_color_val, 
                         ", not plotting contours\n"))
          plot_contours <- FALSE
          title <- paste0(title, "__non-unique quantiles of y",
                          str_color_val)
        }
      }
    }
  }
  
  if (plot_segments) {
    
    if (is.null(x_segment_columns)) {
      
      x_segment_columns$start <- x_column
      x_segment_columns$end <- paste0(x_column, "_end")
      tb[[x_segment_columns$end]] <- c(unlist(tb[2:nrow(tb),x_column]), NA)
    } 
    if (is.null(y_segment_columns)) {
      
      y_segment_columns$start <- y_column
      y_segment_columns$end <- paste0(y_column, "_end")
      tb[[y_segment_columns$end]] <- c(unlist(tb[2:nrow(tb),y_column]), NA)
    } 
    
  }
  
  
  #---Plotting
  if (is.null(color_column)) {
    p <- ggplot(tb, aes(x = !!sym(x_column), y = !!sym(y_column),
                        alpha = opacity))
  } else {
    p <- ggplot(tb, aes(x = !!sym(x_column), y = !!sym(y_column), 
                        color = !!sym(color),
                        alpha = opacity))
  }
  
  if (coord_flip) {
    p <- p + coord_flip()
  }
  
  if (plot_scatter && (nrow(tb) <= n_cutoff || n_cutoff == 0)) {
    if (!is.null(size_column)) {
      size_scaling <- tb[[size_column]]
      
      size_scaling <- size_scaling - min(size_scaling)
      size_scaling <- size_scaling / max(size_scaling)
      
      size_scaling <- size_scaling + size_scaling_offset
    } else {
      size_scaling <- 1
    }
    
    if (is.null(interactive_column)) {
      #---Non-interactive version
      p <- p + geom_point(size = point_size * size_scaling)
    } else {
      #---Interactive version, use with make_interactive
      p <- p + geom_point_interactive(size = point_size * size_scaling,
                                      aes(tooltip = !!sym(interactive_column), 
                                          data_id = !!sym(interactive_column)))
    }
  }
  
  if (plot_lines && (nrow(tb) <= n_cutoff || n_cutoff == 0)) {
    p <- p + geom_line(linewidth = line_size)
  }
  
  if (plot_contours) {  
    p <- p + geom_density_2d(bins = n_contours)
  }
  
  if (plot_segments) {
    if (add_segment_arrows) {
      
      p <- p + geom_segment(data = tibble(!!sym(x_segment_columns$start) := tb[[x_segment_columns$start]],
                                          !!sym(y_segment_columns$start) := tb[[y_segment_columns$start]],
                                          !!sym(x_segment_columns$end) := tb[[x_segment_columns$end]],
                                          !!sym(y_segment_columns$end) := tb[[y_segment_columns$end]],
                                          !!sym(color_column) := tb[[color_column]]),
                            aes(x = !!sym(x_segment_columns$start),
                                xend = !!sym(x_segment_columns$end), 
                                y = !!sym(y_segment_columns$start), 
                                yend = !!sym(y_segment_columns$end)),
                            arrow=arrow(length=unit(0.4,"cm")))
    } else {
      
      segment_df <- tibble(!!sym(x_segment_columns$start) := tb[[x_segment_columns$start]],
                           !!sym(y_segment_columns$start) := tb[[y_segment_columns$start]],
                           !!sym(x_segment_columns$end) := tb[[x_segment_columns$end]],
                           !!sym(y_segment_columns$end) := tb[[y_segment_columns$end]],
                           !!sym(color_column) := tb[[color_column]]) %>% mutate(local_idx=1:n())
      idx <- segment_df %>% filter(!!sym(x_segment_columns$start) > !!sym(x_segment_columns$end)) %>% 
        pull(local_idx)
      segment_df <- segment_df[-idx,]
      p <- p + geom_segment(data = segment_df,
                            aes(x = !!sym(x_segment_columns$start),
                                xend = !!sym(x_segment_columns$end), 
                                y = !!sym(y_segment_columns$start), 
                                yend = !!sym(y_segment_columns$end)),
                            alpha = 1,
                            size = 1.25)
    }
  }
  
  if (plot_marginals) {
    p <- ggMarginal(p, type = "histogram", bins = n_marginal_bins)
  }
  
  
  
  #---Facets
  if (is.character(facet_column)) {
    if (facet_free_y && facet_free_x) {
      scales = "free"
    } else if (facet_free_x) {
      scales = "free_x"
    } else if (facet_free_y) {
      scales = "free_y"
    } else {
      scales = "fixed"
    }
    p <- p + facet_wrap(facet_column, labeller = label_both, 
                        scales = scales)
  }
  
  
  
  #---Colors 
  if (!is.null(color_column)) {
    n_groups <- tb[,color_column] %>% unlist() %>% unique() %>% length()
    cat(sprintf("color groups: %d\n", n_groups))
    
    if (!is_continuous) {
      n_discrete <- length(unique(tb[[color_column]]))
      if (n_discrete == 1) { 
        palette_idx <- 1
      }
    } else {
      n_discrete <- NULL
    }
    
    if (is.null(manual_palette)) {
      if (palette_idx==14 && !is_continuous && n_discrete==3) {
        cols <- color_palette_manual_by_idx(manual_palette_idx=14, n_discrete=3,
                                            reorder=c(1,3,2))
      } else {
        cols <- color_palette(palette_idx, 
                              invert_colors = invert_colors,
                              n_discrete = n_discrete,
                              is_continuous = is_continuous,
                              color_transformation = color_transformation)
      }
    } else {
      cols <- manual_palette
    }
    
    p <- p + cols$scale$col
  }
  
  
  #---axes
  if (!is.null(xlim) && !is.null(ylim)) {
    p <- p + coord_cartesian(xlim = xlim, ylim = ylim)
  } else if (!is.null(xlim)) {
    p <- p + coord_cartesian(xlim = xlim)
  } else if (!is.null(ylim)) {
    p <- p + coord_cartesian(ylim = ylim)
  }
  
  
  if (!logx && is.numeric(unlist(tb[,x_column]))) {
    p <- p + scale_x_continuous(breaks = pretty_breaks())
  }
  
  
  if (logx && logy) {
    p <- p + coord_trans(x = "log10", y = "log10")
  } else if (logx) {
    p <- p + coord_trans(x = "log10")
  } else if (logy) {
    p <- p + coord_trans(y = "log10")
  }
  
  if (!is.null(axis_transformation_x) & !is.null(axis_transformation_y)) {
    p <- p + coord_trans(x=axis_transformation_x, y=axis_transformation_y,
                         xlim = xlim, ylim = ylim)
  } else if (!is.null(axis_transformation_x)) {
    p <- p + coord_trans(x=axis_transformation_x,
                         xlim = xlim)
  } else if (!is.null(axis_transformation_y)) {
    p <- p + coord_trans(y=axis_transformation_y,
                         ylim = ylim)
  } 
  
  if (plot_smooth) {
    
    if (logy) {
      p <- p + ylim(min(tb[[y_column]]), max(tb[[y_column]]))
    }
    
    if (identical(smoothing_method, "gam")) {
      
      p <- p + geom_smooth(method="gam", se=display_smooth_se,
                           method.args=list(gamma=smooth_gam_param)) #smooth_gam_param=2
    } else if (is.null(smoothing_method)) {
      
      if (is.null(smooth_standard_span)) {
        p <- p + geom_smooth(se=display_smooth_se)
      } else {
        # p <- p + geom_smooth(method="loess", span = smooth_standard_span) #doesn't work, actually makes geom_smooth not use method loess...
        p <- p + geom_smooth(span = smooth_standard_span, se=display_smooth_se) #if it doesn't work with axis_transformation, set ylim and xlim so that no infinite values are introduced
      }
      if (nrow(tb) > n_cutoff && n_cutoff !=0 && nrow(tb) <= 1e5) {
        p <- p + geom_smooth(span = 0.1, se=display_smooth_se)
      }
      if (!is.null(smooth_span)) {
        p <- p + geom_smooth(span = smooth_span, se=display_smooth_se)
      }
    } else {
      stop("smoothing_method error")
    }
  }
  # if (plot_ribbon) {
  #   p <- p + geom_ribbon() #needs min_y, max_y
  # }
  if (plot_diagonal) {
    x_range <- range(tb[[x_column]])
    y_range <- range(tb[[y_column]])
    start_val <- max(x_range[1], y_range[1])
    end_val <- min(x_range[2], y_range[2])
    p <- p + geom_segment(aes(x = start_val, y = start_val, 
                              xend = end_val, yend = end_val),
                          color = "orange")
  }
  if (!is.null(plot_constant_lines_y)) {
    x_range <- range(tb[[x_column]])
    for (j in 1:length(plot_constant_lines_y)) {
      p <- p + geom_segment(aes(x = x_range[1], y = plot_constant_lines_y[j], 
                                xend = x_range[2], yend = plot_constant_lines_y[j]),
                            color = "orange")
    }
  }
  if (!is.null(plot_constant_lines_x)) {
    y_range <- range(tb[[y_column]])
    for (j in 1:length(plot_constant_lines_x)) {
      p <- p + geom_segment(aes(y = y_range[1], x = plot_constant_lines_x[j], 
                                yend = y_range[2], xend = plot_constant_lines_x[j]),
                            color = "orange")
    }
  }
  
  
  #---Aspect ratio
  if (!is.null(aspect_ratio)) {
    p <- p + coord_fixed(ratio=aspect_ratio)
  }
  
  
  
  #---Title and labels
  if (!is.null(size_column)) {
    if (title != "") {
      title <- paste0(title, ", ")
    }
    title <- paste0(title, "size: ", size_column)
  }
  if (!is.null(color_column)) {
    p_labs <- labs(title = title, color = title_legend)
  } else {
    p_labs <- labs(title = title)
  }
  p <- p + p_labs
  
  
  if (!is.null(label_x)) {
    p <- p + xlab(label_x)
  }  
  if (!is.null(label_y)) {
    p <- p + ylab(label_y)
  }
  
  #---Point label
  if (!is.null(point_label_column)) {
    p <- p + geom_label(aes(label=!!sym(point_label_column)), 
                        nudge_x = label_nudge, nudge_y = label_nudge, 
                        check_overlap = T)
  }
  
  
  #---Theme
  if (plot_larger) {
    p <- p + theme(legend.key.width = unit(1.25, "cm"),
                   legend.key.height = unit(1.5, "cm")) +
      theme_bw() +
      theme(text = element_text(size = 24))
  } else {
    p <- p + theme(legend.key.width = unit(1.25, "cm"),
                   legend.key.height = unit(1.5, "cm")) +
      theme_bw() +
      theme(plot.title = element_text(size = title_size)) +
      theme(text = element_text(size = 15))
  }
  
  if (!is.null(color_column)) {
    if (n_groups > n_legend_cutoff && n_legend_cutoff != 0 && !is_continuous) {
      p <- p + theme(legend.position = "none")
    }
  }
  
  if (remove_alpha_from_legend) {
    p <- p + scale_alpha(guide = 'none')
  }
  
  return(p)
}


plot_histogram <- function(tb, x_column, 
                           facet_column = NULL,
                           xlim = NULL, ylim = NULL,
                           logx = FALSE,
                           count = FALSE,
                           n_bins = 100,
                           title = "", title_size = 8,
                           plot_larger = TRUE, 
                           aspect_ratio = NULL) {
  
  if (logx) {
    title <- paste0(title, ", LOG")
    
    x_column_prim <- paste0(x_column, "_LOG")
    tb <- tb %>% rename(!!x_column_prim := !!sym(x_column))
    x_column <- x_column_prim
    
    tb[,x_column] <- log(tb[,x_column])
  }
  p <- ggplot(tb, aes(x = !!sym(x_column))) +
    labs(title = title) +
    theme(plot.title = element_text(size = title_size))
  
  if (count) {
    p <- p + geom_histogram(stat = "count") 
  } else {
    # p <- p + geom_histogram(bins = n_bins) 
    p <- p + geom_histogram(aes(y = after_stat(count / sum(count))), bins = n_bins) +
      scale_y_continuous(labels = scales::percent)
  }
  
  if (is.character(facet_column)) {
    p <- p + facet_wrap(facet_column, labeller = label_both)
  }
  if (plot_larger) {
    p <- p + theme(legend.key.width = unit(1.25, "cm"),
                   legend.key.heigh = unit(1.5, "cm")) + 
      theme(text = element_text(size = 32)) #24
  }
  if (!is.null(aspect_ratio)) {
    p <- p + coord_fixed(ratio=aspect_ratio)
  }
  
  if (!is.null(xlim) && !is.null(ylim)) {
    p <- p + coord_cartesian(xlim = xlim, ylim = ylim)
  } else if (!is.null(xlim)) {
    p <- p + coord_cartesian(xlim = xlim)
  } else if (!is.null(ylim)) {
    p <- p + coord_cartesian(ylim = ylim)
  }
  
  
  return(p)
}


plot_3d_scatter_and_surface <- function(tb_data, x_column, y_column, z_column, 
                                        color_column=NULL, size_column=NULL, 
                                        size_scaling=50, #size=size_vec, size_scaling=c(25,75), sets range of sizes, scaled proportionally according to values in size_vec
                                        x_surface=NULL, y_surface=NULL, z_surface=NULL,
                                        opacity_surface=0.6,
                                        title=NULL, x_label=NULL, y_label=NULL, z_label=NULL) {
  
  #---Parse input
  surface_input <- c(!is.null(x_surface), !is.null(y_surface), !is.null(z_surface))
  if (!(all(surface_input) || all(!surface_input)))
    stop("supply all surface inputs or none")
  
  if (is.null(size_column)) {
    size <- 1
  } else {
    size <- tb_data[[size_column]]
  }
  
  if (!is.null(color_column)) {
    color <- tb_data[[color_column]]
  }
  
  if (is.null(title)) {
    title <- ""
  }
  if (is.null(x_label)) {
    x_label <- x_column
  }
  if (is.null(y_label)) {
    y_label <- y_column
  }
  if (is.null(z_label)) {
    z_label <- z_column
  }
  
  
  
  #---Plot scatter points
  p_scatter3d <- plot_ly(data=tb_data, 
                         x=as.formula(paste0("~", x_column)), 
                         y=as.formula(paste0("~", y_column)), 
                         z=as.formula(paste0("~", z_column)),
                         color=color, size=size, sizes=size_scaling,
                         type = "scatter3d", mode="markers",
                         showlegend=TRUE)
  
  #---Plot surface
  if (all(surface_input)) {
    p_scatter3d <- add_trace(p=p_scatter3d, 
                             x=x_surface, y=y_surface, z=z_surface,
                             opacity=opacity_surface,
                             type = "surface",
                             surfacecolor=color)
  }
  
  
  
  #---Labels
  p_scatter3d <- p_scatter3d %>% layout(title=title,
                                        scene=list(xaxis = list(title=x_label),
                                                   yaxis = list(title=y_label),
                                                   zaxis = list(title=z_label)))
  
  
  
  return(p_scatter3d)
}


power_transformation <- function(exponent=0.5) {
  
  #---Use as input argument color_transformation to plot_scatter_lines() and plot_sf()
  transformation = trans_new(
    "power",
    trans=function(x) {x^exponent},
    inverse=function(x) {x^(1/exponent)},
    domain = c(0,Inf)
  )
  
  return(transformation)
}

# NOTE: Other possible transformations are transformation=="sqrt", "log", "exp" ...


save_to_png <- function(p, filename, resolution_scaling=1,
                        resolution_scaling_width=1, 
                        resolution_scaling_height=1) {
  
  cat(sprintf("saving plot to file: %s\n", filename))
  
  png(filename, 
      width=1200 * resolution_scaling * resolution_scaling_width, 
      height=800 * resolution_scaling * resolution_scaling_height)
  
  if ("gtable" %in% class(p)) {
    grid.draw(p)
  } else {
    print(p)
  }
  
  dev.off()
  
  cat("saved plot\n")
}
```
