## Supplementary data and code for "A climate and population dependent diffusion model forecasts the spread of *Aedes Albopictus* mosquitoes in Europe": run_GAM_SPDE_model__fixed_window_cross_validation__climate_GAM.R

```
# Order to run scripts:
# run_GAM_SPDE_model__fixed_window_cross_validation__mechanistic_GAM.R (either cutoff_year=2023 or cutoff_year=2017)
# AUC_for_prediction.R

# Run GAM SPDE model with fixed window cross-validation, for 
#     the climate and pop. GAM model, with
#     ** test set year 4:8: training set 2010-2012, test set 2013-2017 (cutoff_year==2017), or
#        test set year 10:14: training set 2010-2018, test set 2019-2023 (cutoff_year==2017)
#     ** short range SPDE (short_range_spde==TRUE) or free range SPDE (short_range_spde==FALSE)
# NOTE: we can't share data on Ae. albopictus presence / absence. We can share
#     covariates. The code therefore runs on simulated data for Ae. albopictus 
#     presence / absence, but with real covariates.


##########################################################################
# Settings ###############################################################
##########################################################################

# options(error = recover)

# cutoff_year <- 2023
cutoff_year <- 2017

short_range_spde <- TRUE


##########################################################################
# Loading ################################################################
##########################################################################

#---Load functions and packages
source("functions.R")

#---Load data
folder_data <- "Data"

# #---Process data
# #---Data that can be shared
# out <- load(paste0(folder_data, "/data_original.RData"))
# #---Simulated data that replaces the data that cannot be shared
# out <- load(paste0(folder_data, "/data_simulated.RData"))
# 
# #---Compute proximity from data_simulated
# df_proximity <- compute_proximity(data_simulated %>% mutate(y=presence), map_spde)
# 
# #---Bin and add proximity-covariate + add year1-covariate
# out <- process_covariates(data_simulated, df_proximity, keys)
# data_spde <- out$data_spde
# keys <- out$keys
# 
# #---Join the data that can be shared with simulated data
# data_spde <- left_join(data_spde, data_original_covariates)
# 
# save("data_spde", "map_spde", "keys", file=paste0(folder_data, "/data_full.RData"))

#---Load already processed data
out <- load(paste0(folder_data, "/data_full.RData"))

# # VectorNet data
# out <- load(paste0(folder_data, "/data_vectornet_original.RData"))

data_spde <- data_spde %>% filter(date_start <= cutoff_year)


#---Set bins that are not to be used to NA
# data_spde___ <- data_spde
data_spde$bin_idx_proximity050 <- data_spde$bin_idx_proximity050 - 2
data_spde <- data_spde %>% mutate(bin_idx_proximity050 = ifelse(bin_idx_proximity050 <= 0, NA, 
                                                                bin_idx_proximity050))

data_spde$bin_idx_min_temp_CROSS <- data_spde$bin_idx_min_temp_CROSS - 1
data_spde <- data_spde %>% mutate(bin_idx_min_temp_CROSS = ifelse(bin_idx_min_temp_CROSS <= 0, NA, 
                                                                  bin_idx_min_temp_CROSS))

# keys___ <- keys
keys$y_proximity_exp050 <- keys$y_proximity_exp050[3:nrow(keys$y_proximity_exp050),]
keys$y_proximity_exp050[,1] <- keys$y_proximity_exp050[,1]-2

keys$min_temp_CROSS <- keys$min_temp_CROSS[2:nrow(keys$min_temp_CROSS),]
keys$min_temp_CROSS[,1] <- keys$min_temp_CROSS[,1]-1


##########################################################################
# Prepare for SPDE model #################################################
##########################################################################

#---SPDE mesh
out <- create_mesh(map_spde, mesh_precision="low")
spde_mesh <- out$spde_mesh
region_coords <- out$region_coords
n_spde <- out$n_spde

cat("Mesh, size: ", spde_mesh$n, "\n")
par(mfrow=c(1,1))
plot(spde_mesh)
plot_geometry(map_spde, add=TRUE)
points(region_coords, pch=20, col=2)

#---SPDE model
spde_model <- inla.spde2.matern(spde_mesh, alpha=2)

#---Precision matrix from SPDE model, used when short_range_spde==TRUE
Q_params <- c(log(1e-1),log(3)) # tuning parameters for the SPDE, controls SPDE range
Q_spde <- inla.spde2.precision(spde_model,theta=Q_params)


##########################################################################
# Cross-validation #######################################################
##########################################################################

data_spde__full <- data_spde

tb_results_cumul <- NULL


##########################################################################
# Settings ###############################################################
##########################################################################

n_years_cv <- 5 # number of years for fixed window cross-validation

covariate_set <- 2 # 1: mechanistic model covariate, 2: climate and log population covariates

model_str_main <- sprintf("covariate_set%d", covariate_set)

if (short_range_spde) {
  model_str_main <- paste0(model_str_main, "_short_range_SPDE")
} else {
  model_str_main <- paste0(model_str_main, "_free_range_SPDE")
}

j_cv <- n_years_cv

##########################################################################
# Prepare for INLA with SPDE model #######################################
##########################################################################

#---Subset data for cross-validation
unique_years <- sort(unique(data_spde__full$year_idx))
n_max <- max(unique_years)
cv_set <- list(years_training=1:(n_max - j_cv), years_test=n_max - j_cv + 1) # subset to years 1:(n_years-j), predict for year (n_years-j+1)
if (cv_set$years_test > n_max)
  cv_set$years_test <- NA # if cross-validation year is larger than the last year in the data, then use the full dataset as training and no data as test
if (length(cv_set$years_test) > 1)
  stop("not implemented") # the code is made to run cross-validation for one year ahead

ind_year <- data_spde__full$year_idx %in% c(cv_set$years_training, cv_set$years_test)

data_spde <- data_spde__full[ind_year,] %>% 
  mutate(training_set = 1*(year_idx %in% cv_set$years_training & !is.na(presence)), # inla model will be fit on training set
         test_set = 1*(year_idx %in% cv_set$years_test & !is.na(presence)), # model will be used to predict on test set
         y_orig = presence) # keep original values

data_spde$presence[data_spde$test_set] <- NA # inla gives prediction where presence==NA

model_str <- sprintf("%s_cv%d", model_str_main, j_cv)


#---Specific for inlabru
year_mapper <- bru_mapper(fm_mesh_1d(sort(unique(data_spde$year_idx))), indexed = TRUE)

data_spde___ <- data_spde
data_spde <- st_as_sf(data_spde %>% left_join(map_spde %>% select(geo, longitude, latitude)) %>% 
                        select(longitude, latitude), 
                      coords=c("longitude", "latitude"))
data_spde <- bind_cols(data_spde, data_spde___)
rm(data_spde___)


###########################################################################################
# Fit model up until first cross-validation year, predict for year 1 in the test set ######
###########################################################################################

#---Define formula
#---Basic part
f_basic <- "presence ~ Intercept(1) + presence_previous_year + year1 " %>% as.formula()

#---SPDE part
if (!short_range_spde) {
  f_spde <- paste0("Intercept(1) + s(main=geometry, model=spde_model, ",
                   "group=year_idx, group_mapper=year_mapper, ",
                   "control.group = list(model = \"iid\"))")
  
} else {
  f_spde <- paste0("Intercept(1) + s(geometry, model = \"generic0\", Cmatrix = Q_spde, ",
                   "mapper = bru_mapper(spde_mesh), group = year_idx, ",
                   "control.group = list(model = \"iid\"))")
}

#---Random effects part
if (covariate_set==1) {
  
  f_re <- 
    paste0(
      "re_proximity(main=bin_idx_proximity050, model=\"rw1\", ", # proximity random effect
      "n=length(unique(data_spde$bin_idx_proximity050)), scale.model=FALSE, ",
      "hyper=list(theta = list(prior = \"pc.prec\", param = c(0.5, 0.01)))) + ",
      "re_mechanistic(main=bin_idx_mean_adults_pow_merged, model=\"rw1\", ", # mechanistic random effect
      "n=length(unique(data_spde$bin_idx_mean_adults_pow_merged)), scale.model=TRUE, ",
      "hyper=list(theta = list(prior = \"pc.prec\", param = c(0.5, 0.01))))"
    )
  
  f_for_prediction <- "Intercept + presence_previous_year + year1 + re_proximity +
          re_mechanistic" # not used explicitly in prediction, but use to check that the prediction formula fits with the formula used for the prediction
} else {
  
  f_re <- 
    paste0(
      "presence_previous_year + year1 + ",
      "re_proximity(main=bin_idx_proximity050, model=\"rw1\", ", # proximity random effect
      "n=length(unique(data_spde$bin_idx_proximity050)), constr=TRUE, scale.model=TRUE, ",
      "hyper=list(theta = list(prior = \"pc.prec\", param = c(0.5, 0.01)))) + ",
      "re_median_temp(main=bin_idx_median_temp, model=\"rw1\", ", # median temperature random effect
      "n=length(unique(data_spde$bin_idx_median_temp)), constr=TRUE, scale.model=TRUE, ",
      "hyper=list(theta = list(prior = \"pc.prec\", param = c(0.5, 0.01)))) + ",
      "re_min_temp_CROSS(main=bin_idx_min_temp_CROSS, model=\"rw1\", ", # min temperature (cross high median temperature) random effect
      "n=length(unique(data_spde$bin_idx_min_temp_CROSS)), constr=TRUE, scale.model=TRUE, ",
      "hyper=list(theta = list(prior = \"pc.prec\", param = c(0.5, 0.01)))) + ",
      "re_median_relative_humidity(main=bin_idx_median_relative_humidity, model=\"rw1\", ", # median relative humidity random effect
      "n=length(unique(data_spde$bin_idx_median_relative_humidity)), constr=TRUE, scale.model=TRUE, ",
      "hyper=list(theta = list(prior = \"pc.prec\", param = c(0.5, 0.01)))) + ",
      "re_log_population(main=bin_idx_log_pop_merged, model=\"rw1\", ", # log population random effect
      "n=length(unique(data_spde$bin_idx_log_pop_merged)), constr=TRUE, scale.model=TRUE, ",
      "hyper=list(theta = list(prior = \"pc.prec\", param = c(0.5, 0.01))))"
    )
  
  f_for_prediction <- "Intercept + presence_previous_year + year1 + re_proximity +
          re_median_temp + re_min_temp_CROSS + re_median_relative_humidity + re_log_population"
}

#---Combine parts
f <- f_basic %>% update_f(f_spde) %>% update_f(f_re)


#---Run INLA BRU
fitted_model <- bru(f, data = data_spde, family = "binomial", Ntrials=1,
                    options = list(control.compute = list(dic=TRUE, cpo=TRUE, waic=TRUE)))

summary(fitted_model)


##########################################################################
# Summarize fit ##########################################################
##########################################################################

processed_model <- process_fitted_model_bru(data_spde, fitted_model, covariate_set,
                                            spde_model, spde_mesh)

map_full_model <- processed_model$map_full_model
map_random_effects <- processed_model$map_random_effects
map_random_effects_with_time <- processed_model$map_random_effects_with_time
functions_random_effects <- processed_model$functions_random_effects
results_summary <- processed_model$results_summary


################################################################################
# Prepare for prediction for cross-validation for year 2-5 in the test set #####
################################################################################

#---Settings
plot_prediction <- FALSE

#---Select years for prediction
first_year <- max(cv_set$years_training) + 1
n_years_prediction <- max(data_spde__full$year_idx) - first_year
year_to_predict_vec <- first_year:(first_year + n_years_prediction)

#---Select data with covariates used for prediction
data_spde_pred <- left_join(data_spde__full, data_spde %>% select(geo, geometry) %>% distinct())

#---Set covariates that depend on prediction to NA for prediction years
data_spde_pred <- data_spde_pred %>% mutate(
  presence_previous_year = ifelse(year_idx < first_year+1, presence_previous_year, NA),
  y_proximity_exp050 = ifelse(year_idx < first_year+1, y_proximity_exp050, NA)
)

#---Print fitted random effects
functions_random_effects$bin_idx_median_temp

functions_random_effects$bin_idx_min_temp_CROSS

functions_random_effects$bin_idx_median_relative_humidity

functions_random_effects$bin_idx_log_pop_merged

functions_random_effects$bin_idx_proximity050


#---inverse logit-function is used to go from the linear predictor to the probability
# logit <- function(p) log(p / (1-p))
inverse_logit <- function(x) 1 / (1 + exp(-x))


#---Compute precision matrix used to compute proximity based on prediction (need only do once)
geo_neighs <- geo_to_neighs(map_spde)
# plot_geo_neighs(geo_neighs)

Q_besag <- compute_Q_besag(geo_neighs$nb_map, geo_neighs$n_geo, proper_version=TRUE, lambda=0.5)

Q_with_eig <- collect_Q_and_eig(Q_besag)


#---Set up for iterating predicting one year ahead of time, for each year in the
#---test set
run_id <- paste0(year_to_predict_vec[1], "_", 
                 year_to_predict_vec[length(year_to_predict_vec)],  
                 "_climate_GAM")

sum_pred <- NULL

n_rep <- 200

save("data_spde_pred", file=paste0("Output/pred_", run_id, "_map.RData"))

for (j_rep in 1:n_rep) {
  
  print(sprintf('Iteration %d of %d.', j_rep, n_rep))
  
  data_spde_pred <- data_spde_pred %>% mutate(p_predict=NA, y_predict=NA) %>% 
    relocate(data_idx, geo, region_id, year_idx, p_predict, y_predict, 
             presence_previous_year, y_proximity_exp050)
  
  ##########################################################################
  # Predict ################################################################
  ##########################################################################
  
  for (year_to_predict in year_to_predict_vec) {
    
    #---Select data for 'year_to_predict'
    #---Uses presence_previous_year computed from last prediction for current prediction
    tb_for_pred <- data_spde_pred %>% filter(year_idx %in% year_to_predict)
    
    #---Predict probability for 'year_to_predict' 
    tb_pred <- predict(fitted_model, tb_for_pred, ~ data.frame(
      inverse_logit(Intercept + presence_previous_year + year1 + 
                      re_proximity + re_median_temp + re_min_temp_CROSS +
                      re_median_relative_humidity + re_log_population)
    ))
    
    
    
    #---Use probability to simulate Ae. albopictus presence for 
    #---year 'year_to_predict'
    tb_pred <- as_tibble(tb_pred) %>% select(geo, year_idx, p=mean) #tb_pred$mean contains the predicted probabilities
    tb_pred$y_predict <- rbinom(length(tb_pred$p), 1, tb_pred$p) #use predicted probabilities to simulate presence
    
    #---Store predicted probabilites and simulated presence in 'data_spde_pred'
    for (j in 1:nrow(tb_pred)) {
      idx <- which(data_spde_pred$geo==tb_pred$geo[j] & 
                     data_spde_pred$year_idx==tb_pred$year_idx[j])
      
      data_spde_pred$p_predict[idx] <- tb_pred$p[j]
      data_spde_pred$y_predict[idx] <- tb_pred$y_predict[j]
    }
    
    
    
    #---Compute proximity 'presence_previous_year' for next year ('year_to_predict' + 1)
    df_proximity <- compute_proximity_for_prediction(tb_pred, map_spde, Q_with_eig)
    
    #---Store computed proximity and 'presence_previous_year' in 'data_spde_pred'
    for (j in 1:nrow(df_proximity)) {
      idx <- which(data_spde_pred$geo==df_proximity$geo[j] & 
                     data_spde_pred$year_idx==df_proximity$year_idx[j])
      
      data_spde_pred$y_proximity_exp050[idx] <- df_proximity$y_proximity_exp050[j]
      data_spde_pred$presence_previous_year[idx] <- df_proximity$presence_previous_year[j]
    }
    
    
    
    #---Plot prediction
    if (plot_prediction) {
      p1 <- plot_sf(left_join(data_spde_pred %>% as_tibble() %>% select(-geometry) %>% # both data_spde and map_spde contain a column named geometry, here we only need the one in map_spde
                                filter(year_idx==year_to_predict), map_spde), 
                    "y_predict", 
                    legend_title = "pred. presence", title = "predicted presence, current year", 
                    palette_idx=1)
      p2 <- plot_sf(right_join(data_spde_pred %>% as_tibble() %>% select(-geometry) %>% 
                                 filter(year_idx==year_to_predict+1), map_spde), 
                    "y_proximity_exp050", 
                    legend_title = "pred. proximity", title = "predicted proximity, next year")
      p <- ggarrange(p1,p2,ncol=1)
      print(p)
    }
    
  }
  
  add_to_prediction <- ifelse(is.na(data_spde_pred$y_predict), 0, 
                              data_spde_pred$y_predict)
  if (is.null(sum_pred)) {
    sum_pred <- add_to_prediction
  } else {
    sum_pred <- sum_pred + add_to_prediction
  }
  
  save("sum_pred", "j_rep", "n_rep", file=paste0("Output/pred_", run_id, ".RData"))
  
}

data_spde_pred$sum_pred <- sum_pred / j_rep


#---Plot predicted presence averaged over all iterations, for each year in the test set
year1 <- year_to_predict_vec[1]
year2 <- year1+1
year3 <- year1+2
year4 <- year1+3
year5 <- year1+4

p1 <- plot_sf(right_join(data_spde_pred %>% as_tibble() %>% select(-geometry) %>% 
                           filter(year_idx==!!year1), map_spde), 
              "sum_pred", 
              legend_title = "pred. presence", 
              title = sprintf("predicted presence, year %d", year1))
p2 <- plot_sf(right_join(data_spde_pred %>% as_tibble() %>% select(-geometry) %>% 
                           filter(year_idx==!!year2), map_spde), 
              "sum_pred", 
              legend_title = "pred. presence", 
              title = sprintf("predicted presence, year %d", year2))
p3 <- plot_sf(right_join(data_spde_pred %>% as_tibble() %>% select(-geometry) %>% 
                           filter(year_idx==!!year3), map_spde), 
              "sum_pred", 
              legend_title = "pred. presence", 
              title = sprintf("predicted presence, year %d", year3))
p4 <- plot_sf(right_join(data_spde_pred %>% as_tibble() %>% select(-geometry) %>% 
                           filter(year_idx==!!year4), map_spde), 
              "sum_pred", 
              legend_title = "pred. presence", 
              title = sprintf("predicted presence, year %d", year4))
p5 <- plot_sf(right_join(data_spde_pred %>% as_tibble() %>% select(-geometry) %>% 
                           filter(year_idx==!!year5), map_spde), 
              "sum_pred", 
              legend_title = "pred. presence", 
              title = sprintf("predicted presence, year %d", year5))
p <- ggarrange(p1,p2,p3,p4,p5,ncol=2,nrow=3)
print(p)
```
