## Supplementary data and code for "A climate and population dependent diffusion model forecasts the spread of *Aedes Albopictus* mosquitoes in Europe": functions.R

```
create_mesh <- function(map_spde, mesh_precision="mid") {
  
  n_spde <- list()
  n_spde$geo <- nrow(map_spde)
  
  
  # Create mesh -----------------------------------------------------------
  
  shapes <- as_Spatial(map_spde$geometry)
  region_coords <- coordinates(shapes)
  
  map_spde$region_id <- 1:nrow(map_spde)
  
  
  
  # # #---Convex hull
  # # # spde_mesh = inla.mesh.2d(loc=region_coords, cutoff=0.015, max.edge=c(0.1,0.5))
  # # # spde_mesh = inla.mesh.2d(loc=region_coords, cutoff=0.025, max.edge=c(0.2,0.5))
  # # spde_mesh = inla.mesh.2d(loc=region_coords, cutoff=0.025, max.edge=c(0.5,0.5))
  # spde_mesh = inla.mesh.2d(loc=region_coords, cutoff=0.3, max.edge=c(0.35,1))
  # spde_mesh = inla.mesh.2d(loc=region_coords, cutoff=0.3, max.edge=c(0.35,1),
  #                          offset = c(0.5, 7.5))
  # 
  # # 
  # # spde_mesh = inla.mesh.2d(loc=region_coords, cutoff=0.025, max.edge=c(0.5,1), offset=1)
  # # 
  # # cat("Mesh, size: ", spde_mesh$n, "\n")
  # # plot(spde_mesh)
  # # points(region_coords, pch=20, col=2)
  
  
  # #---Non-convex hull
  boundary <- inla.nonconvex.hull(region_coords, convex = 1.2)
  lines(boundary, add = FALSE)
  points(region_coords)
  
  # spde_mesh <- inla.mesh.2d(loc=region_coords, max.n.strict=nrow(map_spde)*5, boundary=boundary)
  
  if (mesh_precision=="high") {
    spde_mesh <- inla.mesh.2d(loc=region_coords, cutoff=0.3, max.edge=c(0.35,1), boundary=boundary)
  } else if (mesh_precision=="mid") {
    spde_mesh <- inla.mesh.2d(loc=region_coords, cutoff=0.45, max.edge=c(0.45,2), boundary=boundary)
  } else if (mesh_precision=="low") {
    spde_mesh <- inla.mesh.2d(loc=region_coords, cutoff=0.6, max.edge=c(0.6,3), boundary=boundary)
  }  
  
  
  # mesh_coords <- spde_mesh$loc[,1:2]
  # points(mesh_coords, pch=20, col=2)
  # points(matrix(mesh_coords[1050,],nrow=1,ncol=2), pch=20, col=2)
  
  # cat("Mesh, size: ", spde_mesh$n, "\n")
  # par(mfrow=c(1,1))
  # plot(spde_mesh)
  # plot_geometry(map_inla, add=TRUE)
  # points(region_coords, pch=20, col=2)
  
  n_spde$full_mesh <- nrow(spde_mesh$loc)
  
  
  return(list(spde_mesh=spde_mesh, region_coords=region_coords, n_spde=n_spde))
}


setup_for_inla <- function(map_spde, data_spde) {
  #---Projection matrix
  shapes <- as_Spatial(map_spde$geometry)
  region_coords <- coordinates(shapes)
  
  tb_coords <- left_join(data_spde %>% select(data_idx, geo),
                         tibble(geo=map_spde$geo, coords=region_coords))
  
  A <- inla.spde.make.A(mesh=spde_mesh, loc=as.matrix(tb_coords$coords),
                        group = data_spde$year_idx)
  
  #---SPDE indices
  spde_indices <- inla.spde.make.index("spde_idx", n.spde = n_spde$full_mesh, 
                                       n.group=length(unique(data_spde$year_idx)))
  
  #---Stack linking SPDE with data
  stack = inla.stack(
    data = data_spde['presence'],
    A = list(A, 1),
    effects = list(spde_indices,
                   list(Intercept = 1,
                        year_idx = data_spde$year_idx,
                        bin_idx_median_temp = data_spde$bin_idx_median_temp,
                        bin_idx_min_temp_CROSS=data_spde$bin_idx_min_temp_CROSS,
                        bin_idx_median_relative_humidity = data_spde$bin_idx_median_relative_humidity,
                        bin_idx_log_pop_merged = data_spde$bin_idx_log_pop_merged,
                        bin_idx_mean_adults_pow_merged = data_spde$bin_idx_mean_adults_pow_merged,
                        bin_idx_proximity050 = data_spde$bin_idx_proximity050,
                        presence_previous_year = data_spde$presence_previous_year,
                        year1 = data_spde$year1)),
    tag = "spde_data"
  )
  
  #---Link function for NA values
  link <- rep(NA, length(stack$data$data$presence))
  ind_NA <- which(is.na(stack$data$data$presence))
  link[ind_NA] <- 1
  
  return(list(spde_indices=spde_indices, stack=stack, link=link))
}


specify_network_by_nodes_and_links <- function(n_nodes, 
                                               nodes_to_disconnect=NULL, 
                                               links_to_remove=NULL, 
                                               links_to_add=NULL,
                                               save_to_file=NULL) {
  
  nb_grid <- cell2nb(nrow=1, ncol=n_nodes)
  if (!is.null(nodes_to_disconnect)) {
    for (j in 1:n_nodes) {
      if (j %in% nodes_to_disconnect) {
        nb_grid[[j]] <- as.integer(0)
      } else {
        nb_grid[[j]] <- setdiff(nb_grid[[j]], nodes_to_disconnect)
      }
    }
  }
  
  if (!is.null(links_to_remove)) {
    for (j in 1:length(links_to_remove)) {
      link_to_disconnect <- links_to_remove[[j]]
      
      node1 <- link_to_disconnect[1]
      node2 <- link_to_disconnect[2]
      nb_grid[[node1]] <- setdiff(nb_grid[[node1]], node2)
      nb_grid[[node2]] <- setdiff(nb_grid[[node2]], node1)
    }
  }
  
  if (!is.null(links_to_add)) {
    for (j in 1:length(links_to_add)) {
      nodes_j <- links_to_add[[j]]
      
      for (k in nodes_j) {
        nb_grid[[k]] <- c(nb_grid[[k]], as.integer(setdiff(nodes_j, k)))
      }
    }
  }
  
  coords <- tibble(x=1, y=1:n_nodes)
  coords$x[coords$y %in% nodes_to_disconnect] <- 2
  for (j in 1:length(links_to_add)) {
    nodes_j <- links_to_add[[j]]
    coords$x[coords$y %in% nodes_j] <- coords$x[coords$y %in% nodes_j] - j*0.25
  }
  plot(nb_grid, coords %>% as.matrix())
  
  if (!is.null(save_to_file))
    nb2INLA(save_to_file, nb_grid)
  
  return(nb_grid)
}


process_fitted_model <- function(data_spde, fitted_model, covariate_set, 
                                 spde_indices, stack, mesh_proj) {
  
  #---Process SPDE fit
  year_idx_vec <- sort(unique(data_spde$year_idx))
  
  tb_regions <- data_spde %>% select(geo, region_id) %>% distinct()
  data_spde_with_na <- 
    tidyr::crossing(tb_regions, tibble(year_idx=unique(data_spde$year_idx))) %>% 
    arrange(year_idx, region_id) %>% left_join(data_spde)
  
  map_full_model <- NULL
  
  for (year_idx__ in year_idx_vec) {
    
    spde_projected <- 
      inla.mesh.project(
        mesh_proj, 
        fitted_model$summary.random$spde_idx$mean[spde_indices$spde_idx.group==year_idx__]
      )
    
    map_full_model <- bind_rows(map_full_model, 
                                bind_cols(data_spde_with_na %>% filter(year_idx == year_idx__) %>% 
                                            select(presence, geo, region_id, year_idx),
                                          spde_projected=spde_projected))
  }
  
  map_full_model <- rename(map_full_model, y = presence)
  
  
  
  #---Subset results
  idx_linear_predictor <- inla.stack.index(stack, "spde_data")$data
  idx_subset <- idx_linear_predictor[1:nrow(data_spde)]
  
  map_full_model$linear_predictor.mean <- fitted_model$summary.linear.predictor[idx_subset, "mean"]
  map_full_model$linear_predictor.sd <- fitted_model$summary.linear.predictor[idx_subset, "sd"]
  map_full_model$p.mean <- fitted_model$summary.fitted.values[idx_subset, "mean"]
  map_full_model$p.sd <- fitted_model$summary.fitted.values[idx_subset, "sd"]
  
  
  
  #---Process min. temp. fit
  if (covariate_set==2) {
    fitted_model <- process_min_temp(data_spde, fitted_model)
  }
  
  
  
  #---Main summaries
  results_summary <- tibble(DIC = round(fitted_model$dic$dic,2), #goodness of fit: DIC
                            logscore = round(-mean(log(fitted_model$cpo$cpo), na.rm = T), 3), #goodness of fit: logscore
                            WAIC = round(fitted_model$waic$waic, 2),
                            fixed_effects = list(fitted_model$summary.fixed), #all fixed effect parameters
                            intercept = fitted_model$summary.fixed["Intercept","mean"], #intercept
                            hyperpar = list(fitted_model$summary.hyperpar)) #fitted hyper parameters
  
  
  
  #---Random effects, constant in time
  #---Initialize
  functions_random_effects <- list()
  map_random_effects <- data_spde %>% filter(year_idx==1) %>% 
    select(data_idx, geo, region_id) #these random effects are constant in time so enough to select first year
  
  #---Add random effects
  #---Random effects for covariates constant in time
  if (covariate_set==1) {
    
    random_effect_vec <- c("bin_idx_mean_adults_pow_merged")
  } else if (covariate_set==2) {
    
    random_effect_vec <- c("bin_idx_median_temp", "bin_idx_min_temp_CROSS",
                           "bin_idx_median_relative_humidity", "bin_idx_log_pop_merged")
  } else if (covariate_set==3) {
    
    random_effect_vec <- NULL
  } else if (covariate_set==4) {
    
    random_effect_vec <- c("bin_idx_median_temp")
  } else if (covariate_set==5) {
    
    random_effect_vec <- c("bin_idx_median_temp", "bin_idx_min_temp_CROSS",
                           "bin_idx_median_relative_humidity")
  }else if (covariate_set==6) {
    
    random_effect_vec <- c("bin_idx_median_temp", "bin_idx_log_pop_merged")
  }else if (covariate_set==7) {
    
    random_effect_vec <- c("bin_idx_mean_adults_pow_merged")
  }
  
  if (!is.null(random_effect_vec)) {
    
    #---Process covariates constant in time
    for (random_effect_col in random_effect_vec) {
      out <- add_random_effect(map_random_effects, functions_random_effects, 
                               random_effect_col=random_effect_col)
      map_random_effects <- out$map_random_effects
      functions_random_effects <- out$functions_random_effects
    }
  } else {
    
    #---If no covariates constant in time
    map_random_effects <- NULL
    functions_random_effects <- list()
  }
  
  
  
  #---Random effect for time-varying covariate
  random_effect_col <- "bin_idx_proximity050"
  
  if (random_effect_col %in% names(fitted_model$summary.random)) {
    
    functions_random_effects[[random_effect_col]] <-
      tibble(!!random_effect_col := fitted_model$summary.random[[random_effect_col]]$ID,
             mean = fitted_model$summary.random[[random_effect_col]]$mean,
             sd = fitted_model$summary.random[[random_effect_col]]$sd,
             lower_bound = fitted_model$summary.random[[random_effect_col]]$`0.025quant`,
             upper_bound = fitted_model$summary.random[[random_effect_col]]$`0.975quant`)
    
    random_effect_str <- paste0("reffect_", random_effect_col)
    
    map_random_effects_curr <-
      left_join(data_spde %>% select(data_idx, region_id, year_idx, !!sym(random_effect_col)),
                functions_random_effects[[random_effect_col]] %>%
                  rename(!!paste0(random_effect_str, ".mean") := mean,
                         !!paste0(random_effect_str, ".sd") := sd,
                         !!paste0(random_effect_str, ".lower_bound") := lower_bound,
                         !!paste0(random_effect_str, ".upper_bound") := upper_bound))
    map_random_effects_with_time <- left_join(data_spde %>% select(data_idx, geo, region_id, year_idx), 
                                              map_random_effects_curr)
  } else {
    
    map_random_effects_with_time <- NULL
  }
  
  
  
  #---Map full model
  map_full_model$y_orig <- data_spde$y_orig
  map_full_model$training_set <- data_spde$training_set
  map_full_model$test_set <- data_spde$test_set
  
  #---Abs error and deviance
  map_full_model$error <- compute_error(map_full_model$p.mean, map_full_model$y, n=1)
  results_summary$total_abs_error <- sum(map_full_model$error, na.rm=TRUE)
  
  map_full_model$deviance <- compute_bernoulli_deviance(map_full_model$p.mean, map_full_model$y)
  results_summary$total_deviance <- sum(map_full_model$deviance, na.rm=TRUE)
  
  if (sum(map_full_model$test_set)==0) {
    
    warning("test set is empty")
    
    map_full_model$error_cv <- NA
    map_full_model$deviance_cv <- NA
    results_summary$total_abs_error_cv <- NA
    results_summary$total_deviance_cv <- NA
  } else {
    
    map_full_model$error_cv <- 
      compute_error(map_full_model$p.mean, map_full_model$y_orig)
    results_summary$total_abs_error_cv <- 
      sum(map_full_model %>% filter(test_set==TRUE) %>% pull(error_cv), na.rm=TRUE)
    
    map_full_model$deviance_cv <- 
      compute_bernoulli_deviance(map_full_model$p.mean, map_full_model$y_orig)
    results_summary$total_deviance_cv <- 
      sum(map_full_model %>% filter(test_set==TRUE) %>% pull(deviance_cv), na.rm=TRUE)
  }
  
  
  
  return(list(results_summary=results_summary,
              map_full_model=map_full_model,
              map_random_effects=map_random_effects,
              map_random_effects_with_time=map_random_effects_with_time,
              functions_random_effects=functions_random_effects))
}


add_random_effect <- function(map_random_effects, functions_random_effects, random_effect_col) {
  
  functions_random_effects[[random_effect_col]] <- 
    tibble(!!random_effect_col := fitted_model$summary.random[[random_effect_col]]$ID,
           mean = fitted_model$summary.random[[random_effect_col]]$mean,
           sd = fitted_model$summary.random[[random_effect_col]]$sd,
           lower_bound = fitted_model$summary.random[[random_effect_col]]$`0.025quant`,
           upper_bound = fitted_model$summary.random[[random_effect_col]]$`0.975quant`)
  
  random_effect_str <- paste0("reffect_", random_effect_col)
  
  map_random_effects_curr <- 
    left_join(data_spde %>% select(data_idx, region_id, !!sym(random_effect_col)),
              functions_random_effects[[random_effect_col]] %>% 
                rename(!!paste0(random_effect_str, ".mean") := mean,
                       !!paste0(random_effect_str, ".sd") := sd,
                       !!paste0(random_effect_str, ".lower_bound") := lower_bound,
                       !!paste0(random_effect_str, ".upper_bound") := upper_bound))
  map_random_effects <- left_join(map_random_effects, map_random_effects_curr)
  
  return(list(map_random_effects=map_random_effects,
              functions_random_effects=functions_random_effects))
}


compute_error <- function(p, y, n=1, abs=TRUE) {
  err <- rep(NA, length(p))
  
  idx_non_na <- which(!is.na(y))
  if (any(is.na(p[idx_non_na])))
    stop("na error")
  
  p_num <- p[idx_non_na]
  p_num[p_num > 1-1e-10] <- 1-1e-10
  p_num[p_num < 1e-10] <- 1e-10
  
  if (abs) {
    err[idx_non_na] <- abs(p_num*n - y[idx_non_na])
  } else {
    err[idx_non_na] <- p_num*n - y[idx_non_na]
  }
  
  return(err)
}


compute_bernoulli_deviance <- function(p, y) {
  deviance <- rep(NA, length(y))
  
  idx_non_na <- which(!is.na(y))
  
  if (any(is.na(p[idx_non_na])))
    stop("na error")
  
  p_num <- p[idx_non_na]
  y_num <- y[idx_non_na]
  
  p_num[p_num > 1-1e-10] <- 1-1e-10
  p_num[p_num < 1e-10] <- 1e-10
  
  # deviance = -2 * log likelihood
  deviance[idx_non_na] <- -2 * ((y_num * log(p_num) + (1-y_num) * log(1-p_num)))
  
  return(deviance)
}


process_min_temp <- function(data_spde, fitted_model) {
  
  # transfer random effect from min_temp_CROSS to median_temp (for low median_temp)
  target_for_transfer <- data_spde %>% filter(bin_idx_min_temp_CROSS==1) %>% 
    pull(bin_idx_median_temp) %>% unique()
  
  random_effect_to_transfer <- fitted_model$summary.random$bin_idx_min_temp_CROSS[1,]
  
  fitted_model$summary.random$bin_idx_median_temp$mean[1:2] <- 
    fitted_model$summary.random$bin_idx_median_temp$mean[1:2] + 
    random_effect_to_transfer$mean
  fitted_model$summary.random$bin_idx_median_temp$`0.025quant`[1:2] <-
    fitted_model$summary.random$bin_idx_median_temp$`0.025quant`[1:2] +
    random_effect_to_transfer$mean
  fitted_model$summary.random$bin_idx_median_temp$`0.975quant`[1:2] <-
    fitted_model$summary.random$bin_idx_median_temp$`0.975quant`[1:2] +
    random_effect_to_transfer$mean
  
  return(fitted_model)
  
}


plot_on_map_selected_model_parts <- function(map_random_effects, 
                                             map_random_effects_with_time,
                                             map_full_model, 
                                             map_spde,
                                             year_to_plot=9,
                                             plot_larger=FALSE) {
  
  ps_map <- list()
  
  if (!is.null(map_random_effects) && nrow(map_random_effects)>0) {
    
    if ("reffect_bin_idx_median_temp.mean" %in% names(map_random_effects)) {
      #---Plot median temp random effect (on map)
      ps_map[[length(ps_map)+1]] <- 
        plot_sf(map_random_effects %>% left_join(map_spde), 
                "reffect_bin_idx_median_temp.mean",
                title="RE, median temp.",
                legend_title="median temp.",
                plot_larger=plot_larger)
    }
    
    if ("reffect_bin_idx_min_temp_CROSS.mean" %in% names(map_random_effects)) {
      map_random_effects <- map_random_effects %>% 
        mutate(reffect_bin_idx_min_temp_CROSS.meanNA=
                 if_else(bin_idx_min_temp_CROSS==1, NA,
                         reffect_bin_idx_min_temp_CROSS.mean))
      #---Plot min temp CROSS random effect (on map)
      ps_map[[length(ps_map)+1]] <- 
        plot_sf(map_random_effects %>% left_join(map_spde), 
                "reffect_bin_idx_min_temp_CROSS.meanNA",
                title="RE, min. temp. CROSS",
                legend_title="min. temp. CROSS",
                plot_larger=plot_larger)
    }
    
    if ("reffect_bin_idx_median_relative_humidity.mean" %in% names(map_random_effects)) {
      #---Plot median relative humidity random effect (on map)
      ps_map[[length(ps_map)+1]] <- 
        plot_sf(map_random_effects %>% left_join(map_spde), 
                "reffect_bin_idx_median_relative_humidity.mean",
                title="RE, median relative humidity",
                legend_title="median rel. humidity",
                plot_larger=plot_larger)
    }
    
    if ("reffect_bin_idx_log_pop_merged.mean" %in% names(map_random_effects)) {
      #---Plot log population random effect (on map)
      ps_map[[length(ps_map)+1]] <- 
        plot_sf(map_random_effects %>% left_join(map_spde),
                "reffect_bin_idx_log_pop_merged.mean",
                title="RE, log population",
                legend_title="log pop.",
                plot_larger=plot_larger)
    }
    
    if ("reffect_bin_idx_mean_adults_pow_merged.mean" %in% names(map_random_effects)) {
      #---Plot mechanistic model random effect (on map)
      ps_map[[length(ps_map)+1]] <- 
        plot_sf(map_random_effects %>% left_join(map_spde), 
                "reffect_bin_idx_mean_adults_pow_merged.mean",
                title="RE, mechanistic model",
                legend_title="mech. model",
                plot_larger=plot_larger)
    }
    
  }
  
  
  
  if (!is.null(map_full_model$spde_projected)) {
    
    #---Plot spatio-temporal field
    ps_map[[length(ps_map)+1]] <- 
      plot_sf(map_full_model %>% left_join(map_spde) %>% 
                filter(year_idx==year_to_plot), "spde_projected",
              title=sprintf("RE, spatio-temporal field, year %d", year_to_plot),
              legend_title="SPDE",
              plot_larger=plot_larger)
  }
  
  
  
  if (!is.null(map_random_effects_with_time)) {
    #---Plot proximity random effect
    non_proximity_bin_idx <- 1:2 #exclude bin indices corresponding to presence previous year and year 1 from the plot
    ps_map[[length(ps_map)+1]] <- 
      plot_on_map_proximity_random_effect(map_random_effects_with_time, map_spde, 
                                          non_proximity_bin_idx,
                                          year_to_plot=year_to_plot,
                                          plot_larger=plot_larger)
  }
  
  
  
  #---Plot fitted model probability (p.mean) and the data (y)
  #---Fitted probability
  ps_map[[length(ps_map)+1]] <- 
    plot_sf(map_full_model %>% filter(year_idx==year_to_plot) %>% 
              left_join(map_spde), 
            "p.mean",
            title=sprintf("prob., year %d", year_to_plot),
            color_transformation=power_transformation(0.1),
            legend_title="prob.",
            plot_larger=plot_larger)
  #---Fitted probability, only showing new observations
  ps_map[[length(ps_map)+1]] <- 
    plot_sf(map_full_model %>% filter(year_idx==year_to_plot) %>% left_join(data_spde) %>%  
              mutate(p.mean.new_obs=ifelse(presence_previous_year==1,NA,p.mean)) %>% # set to NA if Ae. albopictus was observed in the region the previous year
              left_join(map_spde), 
            "p.mean.new_obs",
            title=sprintf("prob., new obs., year %d", year_to_plot),
            legend_title="prob., new obs.",
            color_transformation=power_transformation(0.1),
            plot_larger=plot_larger)
  #---Data, current year
  ps_map[[length(ps_map)+1]] <- 
    plot_sf(map_full_model %>% filter(year_idx==year_to_plot) %>% 
              left_join(map_spde), 
            "y",
            title=sprintf("simulated data, year %d", year_to_plot),
            plot_larger=plot_larger,
            palette_idx=1)
  
  if (year_to_plot > 1) {
    #---Data, previous year
    ps_map[[length(ps_map)+1]] <- 
      plot_sf(map_full_model %>% filter(year_idx==(year_to_plot-1)) %>% 
                left_join(map_spde), 
              "y",
              title=sprintf("simulated data, year %d", year_to_plot-1),
              plot_larger=plot_larger,
              palette_idx=1)
  }
  
  #---Collect figures into one
  n_figures <- length(ps_map)
  ncol <- floor(sqrt(n_figures))
  nrow <- ceiling(n_figures / ncol)
  p <- do.call(ggarrange, c(ps_map, nrow=nrow, ncol=ncol))
  
  return(list(p=p, ps_map=ps_map))
}


plot_on_map_proximity_random_effect <- function(map_random_effects_with_time, 
                                                map_spde, 
                                                special_group_bin_idx, 
                                                year_to_plot=9,
                                                plot_larger=FALSE) {
  
  proximity_col <- names(map_random_effects_with_time) %>% str_subset("reffect_bin_idx_proximity.*mean")
  
  bin_idx_col <- names(map_random_effects_with_time) %>% str_subset("bin_idx_proximity[0-9]*$")
  row_idx_special_groups <- map_random_effects_with_time[[bin_idx_col]] %in% special_group_bin_idx
  
  map_random_effects_with_time$proximity_excl_special_groups <- 
    map_random_effects_with_time[[proximity_col]]
  map_random_effects_with_time$proximity_excl_special_groups[row_idx_special_groups] <- NA #exclude bin indices corresponding to presence previous year and year 1 from the plot
  
  p <- plot_sf(left_join(map_spde, map_random_effects_with_time %>% filter(year_idx==year_to_plot)), 
               "proximity_excl_special_groups",
               title=sprintf("RE, proximity, year %d", year_to_plot),
               legend_title="proximity",
               plot_larger=plot_larger)
  
  return(p)
}


# Version with categories
plot_random_effects_as_functions__ <- function(functions_random_effects, keys,
                                               model_str, 
                                               categories=NULL,
                                               plot_rug_vector=FALSE,
                                               plot_larger=FALSE,
                                               ylim=NULL,
                                               resolution_scaling_width=1,
                                               folder_plots=".") {
  
  random_effects_col <- names(functions_random_effects)
  
  ps <- list()
  
  if ("bin_idx_median_temp" %in% random_effects_col) {
    rug_vector <- NULL
    
    p_median_temp <- plot_random_effect(functions_random_effects, keys=keys$median_temp, 
                                        col_subscript="median_temp",
                                        categories=categories,
                                        vector_to_plot_with_rug=rug_vector, 
                                        filename=paste0(folder_plots, "/", model_str, 
                                                        "_median_temp_reffect"),
                                        plot_larger=plot_larger,
                                        ylim=ylim,
                                        resolution_scaling_width=resolution_scaling_width)
    
    ps$p_median_temp <- p_median_temp
  }
  
  if ("bin_idx_min_temp_CROSS" %in% random_effects_col) {
    rug_vector <- NULL
    
    functions_random_effects[["bin_idx_min_temp_CROSS"]] <- 
      functions_random_effects[["bin_idx_min_temp_CROSS"]] %>% 
      filter(bin_idx_min_temp_CROSS >= 2)
    
    p_min_temp_CROSS <- plot_random_effect(functions_random_effects, keys=keys$min_temp_CROSS, 
                                           col_subscript="min_temp_CROSS",
                                           categories=categories,
                                           vector_to_plot_with_rug=rug_vector, 
                                           filename=paste0(folder_plots, "/", model_str, 
                                                           "_min_temp_CROSS_reffect"),
                                           plot_larger=plot_larger,
                                           ylim=ylim,
                                           resolution_scaling_width=resolution_scaling_width)
    
    ps$p_min_temp_CROSS <- p_min_temp_CROSS
  }
  
  if ("bin_idx_median_relative_humidity" %in% random_effects_col) {
    rug_vector <- NULL
    
    p_median_relative_humidity <- 
      plot_random_effect(functions_random_effects, keys=keys$median_relative_humidity, 
                         col_subscript="median_relative_humidity",
                         categories=categories,
                         vector_to_plot_with_rug=rug_vector, 
                         filename=paste0(folder_plots, "/", model_str, 
                                         "_median_relative_humidity_reffect"),
                         plot_larger=plot_larger,
                         ylim=ylim,
                         resolution_scaling_width=resolution_scaling_width)
    
    ps$p_median_relative_humidity <- p_median_relative_humidity
  }
  
  if ("bin_idx_log_population" %in% random_effects_col) {
    rug_vector <- NULL
    
    p_log_pop <- plot_random_effect(functions_random_effects, keys=keys$log_pop, 
                                    col_subscript="log_population",
                                    vector_to_plot_with_rug=rug_vector, 
                                    filename=paste0(folder_plots, "/", model_str, 
                                                    "_log_pop"),
                                    plot_larger=plot_larger,
                                    ylim=ylim,
                                    resolution_scaling_width=resolution_scaling_width)
    
    ps$p_log_pop <- p_log_pop
  }
  
  if ("bin_idx_log_pop_merged" %in% random_effects_col) {
    rug_vector <- NULL
    
    p_log_pop_merged <- plot_random_effect(functions_random_effects, keys=keys$log_pop_merged, 
                                           col_subscript="log_pop_merged",
                                           categories=categories,
                                           vector_to_plot_with_rug=rug_vector, 
                                           filename=paste0(folder_plots, "/", model_str, 
                                                           "_log_pop_merged"),
                                           plot_larger=plot_larger,
                                           ylim=ylim,
                                           resolution_scaling_width=resolution_scaling_width)
    
    ps$p_log_pop_merged <- p_log_pop_merged
  }
  
  if ("bin_idx_mean_adults_pow_merged" %in% random_effects_col) {
    rug_vector <- NULL
    
    p_mean_ODE <- plot_random_effect(functions_random_effects, keys=keys$mean_adults_pow_merged, 
                                     col_subscript="mean_adults_pow_merged",
                                     categories=categories,
                                     vector_to_plot_with_rug=rug_vector, 
                                     filename=paste0(folder_plots, "/", model_str, 
                                                     "_mean_ODE_merged"),
                                     plot_larger=plot_larger,
                                     ylim=ylim,
                                     resolution_scaling_width=resolution_scaling_width)
    
    ps$p_mean_ODE <- p_mean_ODE
  }
  
  if (any(str_detect(random_effects_col, "proximity"))) {
    p_proximity <- plot_proximity_random_effect(functions_random_effects, keys=keys, 
                                                categories=categories,
                                                filename=paste0(folder_plots, "/", model_str, 
                                                                "_proximity"),
                                                plot_rug_vector=plot_rug_vector,
                                                plot_larger=plot_larger,
                                                ylim=ylim,
                                                resolution_scaling_width=resolution_scaling_width)
    
    ps$p_proximity <- p_proximity
  }
  
  return(list(ps))
}
# Version without categories
plot_random_effects_as_functions <- function(functions_random_effects, keys,
                                             data_spde, model_str,
                                             plot_rug_vector=FALSE,
                                             plot_larger=FALSE,
                                             ylim=NULL,
                                             resolution_scaling_width=1) {

  random_effects_col <- names(functions_random_effects)

  ps <- list()

  if ("bin_idx_median_temp" %in% random_effects_col) {
    if (plot_rug_vector) {
      rug_vector <- data_spde$median_temp
    } else {
      rug_vector <- NULL
    }

    keys$median_temp$n_in_bin <- sapply(keys$median_temp$bin_idx,
                                        function(i) {sum(data_spde$bin_idx_median_temp == i)})
    p_median_temp <- plot_random_effect(functions_random_effects, keys=keys$median_temp,
                                        col_subscript="median_temp",
                                        vector_to_plot_with_rug=rug_vector,
                                        plot_larger=plot_larger,
                                        ylim=ylim,
                                        resolution_scaling_width=resolution_scaling_width)

    ps$p_median_temp <- p_median_temp
  }

  if ("bin_idx_min_temp_CROSS" %in% random_effects_col) {
    rug_vector <- NULL

    keys$min_temp_CROSS$n_in_bin <- sapply(keys$min_temp_CROSS$bin_idx,
                                           function(i) {sum(data_spde$bin_idx_min_temp_CROSS == i, na.rm=TRUE)})

    keys$min_temp_CROSS <- keys$min_temp_CROSS[2:5,] %>%
      mutate(bin_idx=1:n())
    functions_random_effects$bin_idx_min_temp_CROSS <- functions_random_effects$bin_idx_min_temp_CROSS[2:5,] %>%
      mutate(bin_idx_min_temp_CROSS=1:n())
    p_min_temp_CROSS <- plot_random_effect(functions_random_effects, keys=keys$min_temp_CROSS,
                                           col_subscript="min_temp_CROSS",
                                           vector_to_plot_with_rug=rug_vector,
                                           plot_larger=plot_larger,
                                           ylim=ylim,
                                           resolution_scaling_width=resolution_scaling_width)

    ps$p_min_temp_CROSS <- p_min_temp_CROSS
  }

  if ("bin_idx_median_relative_humidity" %in% random_effects_col) {
    if (plot_rug_vector) {
      rug_vector <- data_spde$median_relative_humidity
    } else {
      rug_vector <- NULL
    }

    keys$median_relative_humidity$n_in_bin <-
      sapply(keys$median_relative_humidity$bin_idx,
             function(i) {sum(data_spde$bin_idx_median_relative_humidity == i, na.rm=TRUE)})
    p_median_relative_humidity <-
      plot_random_effect(functions_random_effects, keys=keys$median_relative_humidity,
                         col_subscript="median_relative_humidity",
                         vector_to_plot_with_rug=rug_vector,
                         plot_larger=plot_larger,
                         ylim=ylim,
                         resolution_scaling_width=resolution_scaling_width)

    ps$p_median_relative_humidity <- p_median_relative_humidity
  }

  if ("bin_idx_log_pop_merged" %in% random_effects_col) {
    if (plot_rug_vector) {
      rug_vector <- data_spde$log_population
    } else {
      rug_vector <- NULL
    }

    keys$log_pop_merged$n_in_bin <- sapply(keys$log_pop_merged$bin_idx,
                                           function(i) {sum(data_spde$bin_idx_log_pop_merged == i, na.rm=TRUE)})
    p_log_pop_merged <- plot_random_effect(functions_random_effects, keys=keys$log_pop_merged,
                                           col_subscript="log_pop_merged",
                                           vector_to_plot_with_rug=rug_vector,
                                           plot_larger=plot_larger,
                                           ylim=ylim,
                                           resolution_scaling_width=resolution_scaling_width)

    ps$p_log_pop_merged <- p_log_pop_merged
  }

  if ("bin_idx_mean_adults_pow_merged" %in% random_effects_col) {
    if (plot_rug_vector) {
      rug_vector <- data_spde$mean_adults_pow_merged
    } else {
      rug_vector <- NULL
    }

    keys$mean_adults_pow_merged$n_in_bin <-
      sapply(keys$mean_adults_pow_merged$bin_idx,
             function(i) {sum(data_spde$bin_idx_mean_adults_pow_merged == i, na.rm=TRUE)})
    p_mean_ODE <- plot_random_effect(functions_random_effects, keys=keys$mean_adults_pow_merged,
                                     col_subscript="mean_adults_pow_merged",
                                     vector_to_plot_with_rug=rug_vector,
                                     plot_larger=plot_larger,
                                     ylim=ylim,
                                     resolution_scaling_width=resolution_scaling_width)

    ps$p_mean_ODE <- p_mean_ODE
  }

  if (any(str_detect(random_effects_col, "proximity"))) {
    keys$y_proximity_exp050$n_in_bin <- sapply(keys$y_proximity_exp050$bin_idx,
                                               function(i) {sum(data_spde$bin_idx_proximity050 == i, na.rm=TRUE)})
    p_proximity <- plot_proximity_random_effect(functions_random_effects, keys=keys, data=data_spde,
                                                plot_rug_vector=plot_rug_vector,
                                                plot_larger=plot_larger,
                                                ylim=ylim,
                                                resolution_scaling_width=resolution_scaling_width)

    ps$p_proximity <- p_proximity
  }

  return(list(ps))
}


plot_random_effect <- function(functions_random_effects, keys, 
                               col_subscript,
                               categories=NULL,
                               vector_to_plot_with_rug=NULL, 
                               filename=NULL,
                               plot_larger=FALSE,
                               ylim=NULL,
                               resolution_scaling_width=1,
                               resolution_scaling_height=1) {
  
  bin_idx_col <- paste0("bin_idx_", col_subscript)
  bin_range_col <- paste0("bin_range_", col_subscript)
  bin_mean_col <- paste0("bin_mean_", col_subscript)
  
  #---Plot random effect as a function
  #---Prepare for plotting
  tb_reffect <- functions_random_effects[[bin_idx_col]]
  #---Add bin range and mean of original column
  tb_reffect <- left_join(tb_reffect, keys,
                          by = join_by(!!sym(bin_idx_col) == bin_idx)) %>% 
    relocate(sym(bin_idx_col), sym(bin_range_col), sym(bin_mean_col))
  
  #---Plot with x_column="bin_mean", and make bin position in bottom histogram correspond to the range of values within each bin 
  re_plots <- 
    linked_random_effect_and_histogram_plot(tb_reffect, 
                                            x_column_type="bin_mean", 
                                            categories=categories,
                                            bin_width_constant=FALSE,
                                            vector_to_plot_with_rug=vector_to_plot_with_rug,
                                            plot_larger=plot_larger,
                                            ylim=ylim)
  print(tb_reffect)
  if (!is.null(filename)) {
    #---Save plot
    save_to_png(re_plots$g, filename=paste0(filename, ".png"), 
                resolution_scaling_width=resolution_scaling_width,
                resolution_scaling_height=resolution_scaling_height)
  }
  
  if (!is.null(categories)) {
    ggsave(paste0(filename, "_categories.png"), 
           re_plots$p_categories,
           bg="transparent", 
           width=1200 * resolution_scaling_width * 2, 
           height=800 * resolution_scaling_height * 2 * 0.9,
           units="px")
    
    if (!is.null(re_plots$p_categories_with_segments)) {
      
      ggsave(paste0(filename, "_categories_with_segments.png"), 
             re_plots$p_categories_with_segments,
             width=1200 * resolution_scaling_width * 2 * 0.8, 
             height=800 * resolution_scaling_height * 2 * 0.9,
             units="px")
    }
    
  }
  
  return(re_plots$g)
}


plot_proximity_random_effect <- function(functions_random_effects, keys,
                                         categories=categories,
                                         filename=NULL,
                                         plot_rug_vector=FALSE,
                                         plot_larger=FALSE,
                                         ylim=NULL,
                                         resolution_scaling_width=1,
                                         resolution_scaling_height=1) {
  
  #---Plot proximity random effect (as a function)
  bin_idx_proximity_col <- names(functions_random_effects) %>% str_subset("proximity")
  proximity_exp <- str_extract(bin_idx_proximity_col, "(?<=proximity)[0-9].*")
  keys_col <- str_subset(names(keys), paste0("exp", proximity_exp))
  
  #---Prepare for plotting
  reffect_proximity <- functions_random_effects[[bin_idx_proximity_col]]
  
  reffect_proximity <- reffect_proximity %>% filter(bin_idx_proximity050>=3)
  
  #---Add bin range and mean of original proximity
  reffect_proximity <- left_join(reffect_proximity, keys[[keys_col]],
                                 by = join_by(!!sym(bin_idx_proximity_col) == bin_idx))
  
  #---Plot with x_column="bin_mean", and make bin position in bottom histogram correspond to the range of values within each bin 
  re_plots_proximity <- 
    linked_random_effect_and_histogram_plot(reffect_proximity, 
                                            x_column_type="bin_mean", 
                                            categories=categories,
                                            bin_width_constant=FALSE,
                                            vector_to_plot_with_rug=NULL,
                                            plot_larger=plot_larger,
                                            ylim=ylim)
  # grid.draw(re_plots_max_temp$g)
  print(reffect_proximity)
  
  #---Save plot
  save_to_png(re_plots_proximity$g, filename=paste0(filename, ".png"), 
              resolution_scaling_width=resolution_scaling_width)
  
  if (!is.null(categories)) {
    ggsave(paste0(filename, "_categories_with_segments.png"), 
           re_plots_proximity$p_categories_with_segments,
           width=1200 * resolution_scaling_width * 2 * 0.8, 
           height=800 * resolution_scaling_height * 2 * 0.9,
           units="px")
    
    ggsave(paste0(filename, "_categories.png"), 
           re_plots_proximity$p_categories,
           bg="transparent", 
           width=1200 * resolution_scaling_width * 2, 
           height=800 * resolution_scaling_height * 2 * 0.9,
           units="px")
  }
  
  return(re_plots_proximity$g)
}


linked_random_effect_and_histogram_plot <- function(df_random_effect, 
                                                    x_column_type="bin_mean", #"bin_mean", "bin_idx" or "interaction_bin_mean"
                                                    categories=NULL,
                                                    interaction_x_column=NULL,
                                                    random_effect_title="",
                                                    bin_width_constant=TRUE,
                                                    vector_to_plot_with_rug=NULL,
                                                    plot_larger=FALSE,
                                                    ylim=NULL,
                                                    reduce_window_size=TRUE) {
  
  
  n_column <- names(df_random_effect) %>% str_subset("n_in_bin")
  
  if (x_column_type=="bin_mean") {
    
    x_column <- names(df_random_effect) %>% str_subset("bin_mean")
  } else if (x_column_type=="bin_idx") {
    
    x_column <- names(df_random_effect) %>% str_subset("bin_idx")
  } else if (x_column_type=="interaction_bin_idx") {
    
    x_column <- names(df_random_effect) %>% str_subset("bin_idx.*inter")
    
    if (!bin_width_constant) {
      warning("setting bin_width_constant=TRUE, which is required when x_column_type=\"interaction_bin_idx\"")
      bin_width_constant <- TRUE
    }
  } else {
    
    stop("x_column_type should equal either \"bin_mean\", \"bin_idx\", or \"interaction_bin_idx\"")
  }
  
  if (length(x_column) != 1 || length(n_column) != 1)
    stop("interaction_x_column does not match any of the interaction_bin_mean-columns")
  
  df_random_effect_tall <- bind_rows(tibble(!!sym(x_column) := df_random_effect[[x_column]],
                                            random_effect = df_random_effect$lower_bound,
                                            type = "lower_bound"),
                                     tibble(!!sym(x_column) := df_random_effect[[x_column]],
                                            random_effect = df_random_effect$mean,
                                            type = "mean"),
                                     tibble(!!sym(x_column) := df_random_effect[[x_column]],
                                            random_effect = df_random_effect$upper_bound,
                                            type = "upper_bound"))
  
  
  
  #---Manual color palette
  col_rgb <- col2rgb("gray60")
  col_hex <- rgb(col_rgb[1], col_rgb[2], col_rgb[3], maxColorValue=255)
  manual_palette <- c(col_hex, NA, col_hex)
  col_rgb <- col2rgb("gray5")
  col_hex <- rgb(col_rgb[1], col_rgb[2], col_rgb[3], maxColorValue=255)
  manual_palette[2] <- col_hex
  
  
  
  #---Plot random effect
  p_random_effect <- 
    plot_scatter_lines(df_random_effect_tall, x_column = x_column, y_column = "random_effect",
                       color_column = "type", plot_constant_lines_y = 0,
                       palette_idx = 4, 
                       title=random_effect_title,
                       plot_larger=plot_larger,
                       ylim=ylim)
  
  xlimits <- layer_scales(p_random_effect)$x$range$range
  
  if (!is.null(categories)) {
    
    plot_segments <- TRUE
    
    # nudge_x <- 0.0075
    # nudge_y <- -0.0075
    nudge_x <- 0.015
    nudge_y <- -0.015
    
    if (x_column=="bin_mean_median_temp") {
      
      df_categories <- categories %>% mutate(!!sym(x_column) := median_temp) %>% 
        relocate(sym(x_column), shape_column)
      
      x_segment_columns <- list()
      x_segment_columns$start <- "x_median_temp_start"
      x_segment_columns$end <- "x_median_temp_end"
      
      y_segment_columns <- list()
      y_segment_columns$start <- "y_median_temp_start"
      y_segment_columns$end <- "y_median_temp_end"
      
    } else if (x_column=="bin_mean_median_relative_humidity") {
      
      df_categories <- categories %>% mutate(!!sym(x_column) := median_relative_humidity) %>% 
        relocate(sym(x_column), shape_column)
      
      x_segment_columns <- list()
      x_segment_columns$start <- "x_humidity_start"
      x_segment_columns$end <- "x_humidity_end"
      
      y_segment_columns <- list()
      y_segment_columns$start <- "y_humidity_start"
      y_segment_columns$end <- "y_humidity_end"
      
    } else if (x_column=="bin_mean_log_pop_merged") {
      
      df_categories <- categories %>% mutate(!!sym(x_column) := log_population) %>% 
        relocate(sym(x_column), shape_column)
      
      x_segment_columns <- list()
      x_segment_columns$start <- "x_log_pop_start"
      x_segment_columns$end <- "x_log_pop_end"
      
      y_segment_columns <- list()
      y_segment_columns$start <- "y_log_pop_start"
      y_segment_columns$end <- "y_log_pop_end"
      
    } else if (x_column=="bin_mean_min_temp_CROSS") {
      
      df_categories <- categories %>% mutate(!!sym(x_column) := min_temp) %>% 
        relocate(sym(x_column), shape_column)
      
      x_segment_columns <- list()
      x_segment_columns$start <- "x_min_temp_start"
      x_segment_columns$end <- "x_min_temp_end"
      
      y_segment_columns <- list()
      y_segment_columns$start <- "y_min_temp_start"
      y_segment_columns$end <- "y_min_temp_end"
      
    } else if (x_column=="bin_mean_y_proximity_exp050") {
      
      df_categories <- categories %>% mutate(!!sym(x_column) := proximity_new_establishments) %>% 
        relocate(sym(x_column), shape_column)
      
      x_segment_columns <- list()
      x_segment_columns$start <- "x_proximity_new_start"
      x_segment_columns$end <- "x_proximity_new_end"
      
      y_segment_columns <- list()
      y_segment_columns$start <- "y_proximity_new_start"
      y_segment_columns$end <- "y_proximity_new_end"
      
    } else if (x_column=="bin_mean_mean_adults_pow_merged") {
      
      df_categories <- categories %>% mutate(!!sym(x_column) := mean_adults_pow) %>% 
        relocate(sym(x_column), shape_column)
      
      plot_segments <- FALSE
      
    } else {
      
      browser()
      stop("x_column error")
    }
    
    
    
    if (plot_segments) {
      
      p_categories_with_segments <- 
        plot_scatter_lines(df_categories,
                           x_column = x_column, y_column = "presence",
                           color_column = "median_temp", size_column = "n", 
                           shape_column = "shape_column", 
                           plot_lines = FALSE, plot_segments = TRUE,
                           x_segment_columns = x_segment_columns,
                           y_segment_columns = y_segment_columns,
                           transparent_background = TRUE)
      
      p_categories_with_segments <- p_categories_with_segments + 
        geom_text(aes(label=text_label), alpha=1, size=4, 
                  position=position_nudge(x=nudge_x,y=nudge_y)) +
        theme(text = element_text(size = 18)) +
        theme(legend.position = "none")
      
      
      print(p_categories_with_segments)
    } else {
      
      p_categories_with_segments <- NULL
    }
    
    p_categories <- 
      plot_scatter_lines(df_categories, 
                         x_column = x_column, y_column = "presence",
                         color_column = "median_temp", size_column = "n", 
                         shape_column = "shape_column", 
                         plot_lines = FALSE,
                         ylim = c(0,1),
                         transparent_background = TRUE)
    print(p_categories)
    
    
    
    
    xlimits_categories <- layer_scales(p_categories)$x$range$range
    
    xlimits <- c(min(xlimits[1], xlimits_categories[1]),
                 max(xlimits[2], xlimits_categories[2]))
    
    p_categories <- p_categories + xlim(xlimits[1], xlimits[2])
    p_categories_with_segments <- p_categories_with_segments + 
      xlim(xlimits[1], xlimits[2])
    p_random_effect <- p_random_effect + xlim(xlimits[1], xlimits[2])
    
  } else {
    p_categories_with_segments <- NULL
    p_categories <- NULL
  }
  
  
  
  #---Adjust the end-point of first and last bin in histogram to fit within x-range of
  #---the random effect plot
  first_bin <- list()
  first_bin <- df_random_effect %>% head(1) %>% select(bin_min, bin_max) %>% 
    rename(original_bin_min=bin_min)
  first_bin$bin_min <- xlimits[1]
  first_bin <- first_bin %>% mutate(original_length=bin_max-original_bin_min,
                                    new_length=bin_max-bin_min)
  
  df_random_effect[1,"bin_min"] <- first_bin$bin_min
  
  last_bin <- list()
  last_bin <- df_random_effect %>% tail(1) %>% select(bin_min, bin_max) %>% 
    rename(original_bin_max=bin_max) 
  last_bin$bin_max <- xlimits[2]
  last_bin <- last_bin %>% mutate(
    original_length=original_bin_max-bin_min,
    new_length=bin_max-bin_min)
  
  df_random_effect[nrow(df_random_effect),"bin_max"] <- last_bin$bin_max
  
  print(first_bin)
  print(last_bin)
  
  #---Define height so that height*width=number of data points, with width being the bin range
  df_random_effect <- df_random_effect %>% 
    mutate(bin_height=n_in_bin/(bin_max-bin_min))
  
  
  
  #---Plot histogram with bin width corresponding to the bin range, 
  #---and volume corresponding to number of data points within each bin
  #---with the same x-range as the random effect plot.
  #---The x-range here does not correspond to the range of values in the data!
  p_hist <- ggplot(df_random_effect, aes(x = !!sym(x_column), y = bin_height)) +
    geom_rect(aes(xmin = bin_min, xmax = bin_max, 
                  ymin = 0, ymax = bin_height)) +
    theme_classic() + 
    theme(axis.text.x=element_blank(), #remove x-axis labels since they are redundant (same as for p_random_effect)
          axis.ticks.x=element_blank(),
          axis.title.x=element_blank(),
          axis.text.y=element_blank(), #remove y-axis labels
          axis.ticks.y=element_blank(),
          axis.title.y=element_blank()
    )  
  
  if (!is.null(vector_to_plot_with_rug)) {
    
    p_random_effect <- p_random_effect + 
      geom_rug(data=data.frame(x=vector_to_plot_with_rug), aes(x=x, y=0), 
               inherit.aes = FALSE, alpha = 1/2, position = "jitter", sides="b")
  }
  
  g <- align_plots(p_random_effect, p_hist) 
  grid.draw(g)
  
  
  return(list(g=g, p_random_effect=p_random_effect, p_hist=p_hist, 
        p_categories_with_segments=p_categories_with_segments,
        p_categories=p_categories))
}


compute_roc_and_auc_inner <- function(prob, y) {
  
  roc_curve <- roc(y, prob)
  
  auc_value <- auc(roc_curve)
  
  roc_coords <- coords(roc_curve, "local maximas", ret=c("threshold", "specificity", "sensitivity", "npv", "ppv"))
  
  roc_coords_best <- coords(roc_curve, "best", ret=c("threshold", "specificity", "sensitivity", "npv", "ppv"))
  
  return(list(auc=auc_value, roc=roc_curve,
              tb_roc_curve=tibble(sensitivity=roc_curve$sensitivities, 
                                  false_negative_rate=1-roc_curve$specificities),
              coords=roc_coords,
              coords_best=roc_coords_best))
}


plot_roc_curve <- function(roc_curve, color_column=NULL, plot_larger=TRUE, title="") {
  
  p <- 
    plot_scatter_lines(roc_curve,
                       "false_negative_rate", "sensitivity", color_column=color_column,
                       title=title, label_x="false negative rate", label_y="sensitivity",
                       plot_larger=plot_larger)
  
  return(p)
}


compute_roc_for_cv_set <- function(map_all, 
                                   roc_curve_long=NULL, 
                                   model_prob_long=NULL,
                                   model_name,
                                   plot_curve=TRUE) {
  
  map_sub <- map_all %>% 
    filter(test_set==1, !is.na(y_orig), presence_previous_year==0)
  
  prob <- map_sub$p.mean
  y <- map_sub$y_orig
  
  model_prob_long <- bind_rows(
    model_prob_long,
    tibble(prob=prob, y=y, model_name=model_name)
  )
  
  if (!identical(sort(unique(map_sub$y_orig)), c(0,1))) {
    return(list(model_prob_long=model_prob_long))
  }
  
  output <- compute_roc_and_auc_inner(prob, y)
  
  if (plot_curve) 
    plot_roc_curve(output$tb_roc_curve, title=model_name) %>% print()
  
  roc_curve_long <- bind_rows(
    roc_curve_long,
    tibble(
      false_negative_rate=output$tb_roc_curve$false_negative_rate,
      sensitivity=output$tb_roc_curve$sensitivity,
      model_name=model_name
    )
  )
  
  return(list(roc_curve_long=roc_curve_long, auc=output$auc,
              model_prob_long=model_prob_long))
}


compute_roc_and_auc <- function(tb_results_cumul, geo_subset=NULL) {
  
  n_fits <- nrow(tb_results_cumul)
  
  roc_curve_long <- NULL
  model_prob_long <- NULL
  auc_vec <- rep(NA, n_fits)
  for (j in 1:n_fits) {
    
    map_for_ROC <- tb_results_cumul$map_for_ROC[[j]]
    
    if (!is.null(geo_subset)) {
      map_for_ROC <- map_for_ROC %>% filter(geo %in% geo_subset)
    }
    
    output <- compute_roc_for_cv_set(map_for_ROC, 
                                     roc_curve_long=roc_curve_long, 
                                     model_prob_long=model_prob_long,
                                     model_name=tb_results_cumul$model_str[j],
                                     plot_curve=FALSE)
    
    model_prob_long <- output$model_prob_long
    
    if (!is.null(output$roc_curve_long)) {
      roc_curve_long <- output$roc_curve_long
      auc_vec[j] <- output$auc
    }
  }
  
  auc_simultaneous <- compute_roc_and_auc_inner(model_prob_long$prob, model_prob_long$y)
  
  return(list(auc_simultaneous=auc_simultaneous, auc_separate=auc_vec))
}


compute_Q_besag <- function(nb_map, n_geo, proper_version = FALSE,
                            lambda = NULL) {
  if (proper_version) {
    if (is.null(lambda)) 
      stop("lambda parameter should be supplied for proper version of Besag")
    if (!(lambda > 0 && lambda < 1))
      stop("invalid lambda-value")
  }
  if (!proper_version && !is.null(lambda)) 
    stop("lambda parameter is not required when proper_version == FALSE")
  
  if (n_geo != length(nb_map))
    stop("n_geo error")
  neighs <- vector(mode = "list", length = n_geo)
  n_neighs <- vector(length = n_geo)
  
  Q = matrix(0, nrow = n_geo, ncol = n_geo)
  for (j in seq_along(nb_map)) {
    ns <- nb_map[[j]]
    if (!all(ns == 0)) {
      neighs[[j]] <- ns
      n_neighs[j] <- length(ns)
      
      Q[neighs[[j]],j] = -1
      Q[j,neighs[[j]]] = -1
      Q[j,j] = n_neighs[j]
    } 
  }
  
  if (proper_version) {
    Q <- lambda * Q
    for (i in 1:n_geo) 
      Q[i,i] <- Q[i,i] + (1-lambda)
  }
  
  return(Q)
}


collect_Q_and_eig <- function(Q) {
  
  rank <- rankMatrix(Q)
  E <- eigen(Q)
  determinant_log <- sum(log(E$values[1:rank]))
  
  return(list(rank = rank, 
              determinant_log = determinant_log,
              E = E,
              Q = Q))
}


Q_convolution <- function(Q_with_eig, exponent, x, NAval = NA, use_noise = FALSE) {
  
  n <- nrow(Q_with_eig$Q)
  
  if (!is.na(NAval)) {
    ind <- which(x==NAval)
  } else {
    ind <- which(is.na(x))
  }
  
  Q_rank <- Q_with_eig$rank
  if (Q_rank < nrow(Q_with_eig$Q)) {
    Q_with_eig$E$values[(Q_with_eig$rank[1]+1):n] <- 0 #numerical instabilities with the 0 eigenvalues
  }
  if (use_noise) {
    d <- median(Q_with_eig$E$values) / 5
  } else {
    d <- 0
  }
  Q <- Q_with_eig$E$vectors %*% diag(Q_with_eig$E$values^exponent + d) %*% t(Q_with_eig$E$vectors)
  
  Q_AA <- Q[ind, ind]
  Q_AB <- Q[ind, -ind]
  
  if (rankMatrix(Q_AA) != nrow(Q_AA)) 
    stop("rank error, Q_AA")
  
  
  if (length(x[-ind]) > 1) {
    x[ind] <- - solve(Q_AA, Q_AB %*% x[-ind])
  } else {
    x[ind] <- - solve(Q_AA, Q_AB * x[-ind])
  }
  
  
  return(x)
}


proximity_from_Q <- function(Q_with_eig, tb, y_column, map, exponent_vec=1,
                             add_year_one=TRUE) {
  #---Convolve matrix Q^exponent (in Q_with_eig <- collect_Q_and_eig(Q)) with the column tb[[y_column]] 
  #---on map.
  #---tb and map are linked through the column geo. 
  #---The convolution is done on a yearly basis by convolving with the known mosq. distribution tb[[y_column]] 
  #---for the previous year. 
  #---The convolution is set to zero the first year since there is no information about the mosq. distribution 
  #---of the previous year. 
  #---The exponent is implemented in Q_convolution as Q^exponent = E %*% L^exponent %*% E^T, where 
  #---Q = E %*% L %*% E^T is the eigen-value decomposition of Q.
  
  year_vec <- sort(unique(tb$year_idx))
  # year_vec <- setdiff(year_vec, max(year_vec))
  df_proximity <- tb %>% select(geo, year_idx)
  df_proximity_last_year <- tibble(geo=unique(df_proximity$geo), year_idx=max(tb$year_idx)+1)
  df_proximity <- bind_rows(df_proximity, df_proximity_last_year)
  for (exponent in exponent_vec) {
    
    # cat(sprintf("Computing convolution of y and Q^exponent, with exponent: %.2g\n", exponent))
    
    y_proximity_str <- sprintf("y_proximity_exp%03d", exponent*100)
    
    #---Compute proximity by convolving y with Q^exponent with current exponent, for each year separately (convolution from previous year as proximity for the next year)
    df_proximity_curr <- NULL
    for (year in year_vec) {
      
      map_year <- left_join(map, tb %>% filter(year_idx == year))
      
      y <- map_year[[y_column]]
      y[is.na(y)] <- 0
      
      #---Convolve y with Q^exponent for the current year
      y_convolved <- Q_convolution(Q_with_eig, exponent=exponent, y, 
                                   NAval=0, use_noise=TRUE)
      
      #---Add convolution with current year as proximity for the next year
      df_proximity_curr <-  df_proximity_curr %>% 
        bind_rows(tibble(!!sym(y_proximity_str) := y_convolved,
                         y = y, 
                         year_idx = year + 1,
                         geo = map_year$geo))
    }
    
    if (add_year_one) {
      #---Set year 1 to zero (as we don't have any data from the previous year to convolve with)
      df_proximity_curr <- df_proximity_curr %>% 
        bind_rows(tibble(!!sym(y_proximity_str) := 0,
                         y = 0, 
                         year_idx = min(df_proximity$year_idx),
                         geo = map_year$geo))
    } 
    
    #---Add convolution with current exponent
    df_proximity <- left_join(df_proximity, df_proximity_curr)
  }
  
  proximity_cols <- str_subset(names(df_proximity_curr), "y_proximity")
  for (col in proximity_cols) {
    df_proximity <- df_proximity[!is.na(df_proximity[[col]]),]
  }
  
  # p_conv <- plot_sf(left_join(map, df_proximity %>% filter(year_idx==2)), "y_conv_exp100")
  # p_conv
  
  return(df_proximity)
}


compute_proximity_for_prediction <- function(df, map, Q_with_eig) {
  
  df <- df %>% 
    mutate(y_to_convolve = 1*(y_predict==1)) #y==1 if presence==1, y==0 if presence %in% c(1, NA)
  
  y_column <- "y_to_convolve"
  
  exponent_vec <- 0.5
  
  df_proximity <- proximity_from_Q(Q_with_eig, df, y_column, map, exponent_vec,
                                   add_year_one=FALSE)
  df_proximity <- df_proximity %>% rename(presence_previous_year=y)
  
  df_surrounded <- get_surrounded_regions()
  
  proximity_col_vec <- names(df_proximity) %>% str_subset("y_proximity")
  
  for (j in 1:nrow(df_proximity)) {
    df_curr <- df_proximity[j,]
    if (df_curr$geo %in% df_surrounded$inner) {
      geo_outer <- df_surrounded %>% filter(inner == df_curr$geo) %>% pull(outer)
      df_curr_match <- df_proximity %>% filter(geo == geo_outer, year_idx == df_curr$year_idx)
      if (df_curr_match$presence_previous_year != 1) {
        for (proximity_col in proximity_col_vec) {
          df_proximity[[proximity_col]][j] <- df_curr_match[[proximity_col]]
        }
      }
    }
  }
  
  return(df_proximity)
}


geo_to_neighs <- function(map, use_snap=TRUE, use_queen=TRUE) {
  #---Assumes that geometry features are stored in a column called geometry
  
  if ("sfc" %in% class(map)) {
    #---Class of input map is valid, but the geometry features are not stored correctly
    #---Create a valid data.frame
    
    warning("creating data.frame from sfc input")
    
    map_geometry <- map
    map <- tibble(geo_idx=1:length(map_geometry), geometry=map_geometry)
  } else if (!"sfc" %in% class(map$geometry)) {
    #---Class of geometry column is not valid
    
    stop("no valid geometry column")
  }
  
  if (!"geo_idx" %in% colnames(map)) {
    #---Create missing geometry index column
    
    map <- map %>% select(geometry) %>% mutate(geo_idx=1:nrow(map))
  }
  
  
  #---Check that there are no duplicate geometries in map
  n_geo <- map$geometry %>% unique() %>% length()
  if (n_geo != nrow(map))
    stop('duplicate geometries in map')
  
  
  #---Create SpatialPolygons object from the geometry column (sfc object) 
  shapes <- as_Spatial(map$geometry)
  # nb_overlap <- st_overlaps(map$geometry, sparse = FALSE) #check if any geometries overlap
  
  
  #---Build neighbor list based on region boundaries
  if (use_snap) {
    
    nb_map <- poly2nb(shapes, snap=sqrt(.Machine$double.eps)*5e5, #snap: boundary points less than snap distance apart are considered neighbors
                      queen = use_queen) #queen: if true, a single shared boundary point meets the neighbor criteria, if false, more than one shared point is required
  } else {
    
    nb_map <- poly2nb(shapes, queen = use_queen) 
  }
  #---Store central coordinates of each geometry
  coords <- coordinates(shapes)
  
  
  
  return(list(geo_idx = map$geo_idx,
              shapes = shapes,
              nb_map = nb_map,
              n_geo = n_geo,
              coords = coords))
}


get_surrounded_regions <- function() {
  tb_surrounded3 <- tibble(outer=c("CZ020", "HU120", "RO322", "EL306", "EL305",
                                   "EL305", "EL305", "BG412", "AT126"),
                           inner=c("CZ010", "HU110", "RO321", "EL302",
                                   "EL304", "EL303", "EL301", "BG411",
                                   "AT130"))
  
  tb_surrounded <- tibble(outer=c("DEG0L", "DEE0B", "DE40G", "DEA2C", "DE734",
                                  "DE716", "DE94E", "DE238", "DE145", "DED2F",
                                  "DE276", "DE219", "DE40E", "DE40E", "DEE07",
                                  "DEE05"),
                          inner=c("DEG02", "DEE02", "DE402", "DEA22", "DE731",
                                  "DE711", "DE944", "DE232", "DE144", "DED21",
                                  "DE271", "DE211", "DE404", "DE401", "DEE03",
                                  "DEE01"))
  
  tb_surrounded2 <- 
    tibble(outer=c("DEB3K", "DEB3H", "DEB3F", "DE124", "DE12B", "DE118",
                   "DEB25", "DE124", "DE27E", "DE27B", "DE256", "DE26C",
                   "DE26B", "DE245", "DE247", "DE246", "DE249", "DE237",
                   "DE234", "DE21K", "DE227", "DE22B", "DEG0G", "DEG0P",
                   "DE734", "DEG0J"), 
           inner=c("DEB37", "DEB33", "DEB32", "DE121", "DE129", "DE117",
                   "DEB21", "DE121", "DE273", "DE272", "DE251", "DE263",
                   "DE262", "DE241", "DE243", "DE242", "DE244", "DE233",
                   "DE231", "DE213", "DE221", "DE223", "DEG05", "DEG0N",
                   "DE731", "DEG03"))
  
  tb_surrounded <- bind_rows(tb_surrounded, tb_surrounded2) %>% 
    bind_rows(tb_surrounded3) %>% distinct()
  
  return(tb_surrounded)
}


compute_proximity_one_year_ahead <- function(df, map, Q_with_eig, y_column) {
  
  df <- df %>% 
    mutate(y_to_convolve = 1*(!!sym(y_column)==1)) #y==1 if presence==1, y==0 if presence %in% c(1, NA)
  
  y_column <- "y_to_convolve"
  
  exponent_vec <- 0.5
  
  df_proximity <- proximity_from_Q(Q_with_eig, df, y_column, map, exponent_vec,
                                   add_year_one=FALSE)
  df_proximity <- df_proximity %>% rename(presence_previous_year=y)
  
  df_surrounded <- get_surrounded_regions()
  
  proximity_col_vec <- names(df_proximity) %>% str_subset("y_proximity")
  
  for (j in 1:nrow(df_proximity)) {
    df_curr <- df_proximity[j,]
    if (df_curr$geo %in% df_surrounded$inner) {
      geo_outer <- df_surrounded %>% filter(inner == df_curr$geo) %>% pull(outer)
      df_curr_match <- df_proximity %>% filter(geo == geo_outer, year_idx == df_curr$year_idx)
      if (df_curr_match$presence_previous_year != 1) {
        for (proximity_col in proximity_col_vec) {
          df_proximity[[proximity_col]][j] <- df_curr_match[[proximity_col]]
        }
      }
    }
  }
  
  return(df_proximity)
}


compute_proximity <- function(data, map_spde) {
  #---Compute precision matrix for the proximity diffusion process
  geo_neighs <- geo_to_neighs(map_spde)
  # plot_geo_neighs(geo_neighs)
  
  Q_besag <- compute_Q_besag(geo_neighs$nb_map, geo_neighs$n_geo, proper_version=TRUE, lambda=0.5)
  
  Q_with_eig <- collect_Q_and_eig(Q_besag)
  
  year_vec <- sort(unique(data$year_idx))
  
  df_proximity <- data %>% filter(year_idx==1) %>% select(geo, year_idx) %>%  
    mutate(y_proximity_exp050=0, presence_previous_year=0)
  
  for (curr_year in year_vec) {
    
    #---Compute proximity based on data_simulated$y for the current year
    df_proximity_curr <- compute_proximity_one_year_ahead(data %>% 
                                                            filter(year_idx==curr_year), 
                                                          map_spde, Q_with_eig, y_column="y")
    
    df_proximity <- bind_rows(df_proximity, df_proximity_curr)
    
    cat(sprintf("compute_proximity: progress %d of %d\n", curr_year, length(year_vec)))
  }
  
  df_proximity <- df_proximity %>% filter(year_idx %in% data$year_idx)
  
  return(df_proximity)
}


process_covariates <- function(data_simulated, df_proximity, data_original_covariates, keys) {
  
  #---Bin the proximity covariate (covariates in data_original_covariates have already been binned)
  breaks <- c(-Inf,
              quantile(df_proximity %>% filter(year_idx>1, presence_previous_year==0) %>% 
                         pull(y_proximity_exp050), 
                       c(seq(0.5,0.9,by=0.1), 0.95)),
              Inf)
  df_proximity <- df_proximity %>% mutate(
    bin_idx_proximity050 = as.numeric(cut(y_proximity_exp050,breaks=breaks)),
    bin_idx_proximity050 = case_when(
      presence_previous_year==1 ~ 1,
      year_idx==1 ~ 2, 
      TRUE ~ bin_idx_proximity050+2
    )
  )
  
  #---Compute range of original y_proximity_exp050 for each bin (will be used for plotting)
  keys$y_proximity_exp050 <- compute_bin_keys(df_proximity, 
                                              bin_column="bin_idx_proximity050", 
                                              original_column="y_proximity_exp050",
                                              print_format="[%.2f,%.2f]")
  
  levels(keys$y_proximity_exp050[[2]])[1] <- "albo_prev_year"
  levels(keys$y_proximity_exp050[[2]])[2] <- "year1"
  
  
  
  #---Join data 
  data_simulated <- data_simulated %>% left_join(df_proximity) %>% 
    left_join(data_original_covariates) %>% 
    relocate(data_idx, geo, region_id, year_idx, date_start, CountryISO,
             presence, presence_previous_year, y_proximity_exp050, bin_idx_proximity050)
  
  
  
  #---Add year1 covariate
  data_simulated <- data_simulated %>% mutate(year1 = 1*(year_idx==1))
  
  
  
  return(list(data_simulated=data_simulated, keys=keys))
}


compute_bin_keys <- function(data, bin_column, original_column, 
                             print_format="[%.1f,%.1f]") {
  
  #---Prepare data.frame
  bin_range_column <- paste0("bin_range_", original_column)
  bin_mean_column <- paste0("bin_mean_", original_column)
  
  keys_to_bins <- tibble(bin_idx = sort(unique(data[[bin_column]])),
                         !!sym(bin_range_column) := NA,
                         bin_min = NA, bin_max = NA,
                         !!sym(bin_mean_column) := NA,
                         n_in_bin = NA)
  
  for (j in 1:nrow(keys_to_bins)) {
    
    #---Select rows in data with current bin_idx
    curr_bin_idx <- keys_to_bins$bin_idx[j]
    data_curr_bin <- data %>% filter(!!sym(bin_column) == curr_bin_idx) #filter on bin_column == curr_bin_idx
    # print(data_curr_bin %>% relocate(all_of(c(original_column, bin_column))))
    
    #---Compute range of values within current bin
    bin_range <- range(data_curr_bin[[original_column]]) #range of values in original_column for this bin
    bin_range <- round(bin_range, digits=13)
    keys_to_bins[[bin_range_column]][j] <- sprintf(print_format, bin_range[1], bin_range[2]) #make string of range
    keys_to_bins$bin_min[j] <- bin_range[1]
    keys_to_bins$bin_max[j] <- bin_range[2]
    
    #---Mean of values which lie within current bin
    keys_to_bins[[bin_mean_column]][j] <- mean(data_curr_bin[[original_column]])
    
    #---Number of rows with current bin_idx
    keys_to_bins$n_in_bin[j] <- nrow(data_curr_bin) 
    
  }
  
  keys_to_bins <- keys_to_bins %>% mutate(!!sym(bin_range_column) := 
                                            factor(!!sym(bin_range_column), 
                                                   levels=!!sym(bin_range_column), ordered=TRUE))  #make factor to keep the bin range strings sorted by numerical value when plotting  
  
  return(keys_to_bins)
}


setup_for_distance_to_presence <- function(data_spde, n_points=5000, year_idx_=5,
                                           do_plot=FALSE) {
  
  ##############################################################################
  # Setup ######################################################################
  ##############################################################################
  
  setup <- list()
  
  geo_with_presence <- data_spde %>% filter(year_idx==year_idx_, presence==1) %>% pull(geo)
  
  setup$geometry_with_presence <- map_spde %>% filter(geo %in% geo_with_presence) %>% 
    pull(geometry) %>% st_union()
  
  
  setup$map_without_presence <- map_spde %>% filter(!geo %in% geo_with_presence)
  setup$geometry_without_presence <- setup$map_without_presence %>% pull(geometry) %>% st_union()
  
  
  
  
  
  ##############################################################################
  # Regular grid of points #####################################################
  ##############################################################################
  
  points <- spsample(as_Spatial(setup$geometry_without_presence), n_points, type = "regular")
  # points <- spsample(as_Spatial(setup$geometry_without_presence), 500, type = "regular")
  
  if (do_plot) {
    plot(setup$geometry_without_presence, add=FALSE, col=scales::alpha('red', 0.5))
    plot(setup$geometry_with_presence, add=TRUE, col=scales::alpha('blue', 0.5))
    points(points, pch=20, col=1)
  }
  
  
  setup$tb_points <- tibble(points=st_as_sf(points) %>% pull(geometry))
  
  setup$tb_points$dist <- st_distance(st_as_sf(setup$tb_points$points), setup$geometry_with_presence) %>% 
    as.double()
  
  
  
  return(setup)
}


regions_close_to_presence <- function(setup, distance_threshold=1e5,
                                      do_plot=TRUE) {
  
  ##############################################################################
  # Points close to recorded presence ##########################################
  ##############################################################################
  
  points_close <- setup$tb_points %>% filter(dist < distance_threshold) %>% pull(points)
  
  if (do_plot) {
    plot(setup$geometry_without_presence, add=FALSE, col=scales::alpha('red', 0.5))
    plot(setup$geometry_with_presence, add=TRUE, col=scales::alpha('blue', 0.5))
    points(points_close %>% as_Spatial(), pch=20, col=3)
  }
  
  
  
  
  ##############################################################################
  # Match points to region #####################################################
  ##############################################################################
  
  g_points <- as_geos_geometry(points_close)
  
  g_geometry <- as_geos_geometry(setup$map_without_presence %>% pull(geometry))
  
  
  wk::wk_crs(g_points) <- NULL
  tree <- geos_strtree(g_points)
  
  wk::wk_crs(g_geometry) <- NULL
  
  keys <- geos_contains_matrix(g_geometry, tree)
  
  
  
  
  
  # ##############################################################################
  # # Plot for one region ########################################################
  # ##############################################################################
  # 
  # keys[[995]]
  # 
  # plot(setup$geometry_without_presence, add=FALSE, col=scales::alpha('red', 0.5))
  # plot(setup$geometry_with_presence, add=TRUE, col=scales::alpha('blue', 0.5))
  # points(points, pch=20, col=1)
  # 
  # plot(setup$map_without_presence$geometry[995], add=TRUE, col=scales::alpha('yellow', 0.5))
  # points(as_Spatial(points_close[keys[[995]]]), pch=20, col=3)
  
  
  
  
  
  ##############################################################################
  # Extract matching regions ###################################################
  ##############################################################################
  
  
  idx_match <- keys %>% sapply(length) %>% sapply(function(x) {x>0}) %>% which()
  
  geo_matching <- setup$map_without_presence$geo[idx_match]
  
  geometry_matching <- setup$map_without_presence %>% filter(geo %in% geo_matching) %>% 
    pull(geometry) %>% st_union()
  
  map_matching <- map_spde %>% filter(geo %in% geo_matching)
  
  
  
  if (do_plot)
    plot(geometry_matching, add=TRUE, col=scales::alpha('yellow', 0.5))
  
  
  
  return(map_matching) 
}


roc_subsetted_by_distance <- function(tb_results_cumul,
                                      distance_threshold_vec=c(0.5e5, 1e5, 1.5e5, 2e5, 5e5, NA)) {
  
  models_vec <- tb_results_cumul$model_str %>% str_extract(".*(?=_cv)") %>% unique()
  
  tb_roc <- NULL
  
  for (model_curr in models_vec) {
    
    #---Fits for current model
    tb_results_curr <- tb_results_cumul %>% filter(str_detect(model_str, model_curr))
    
    #---Setup
    n_cv <- nrow(tb_results_curr)
    
    for (j_distance in 1:length(distance_threshold_vec)) {
      
      distance_curr <- distance_threshold_vec[j_distance]
      
      tb_roc_inner <- tibble(year_cv=rep(NA, n_cv), auc=NA, roc_curve=NA, n_regions=NA)
      
      map_for_ROC_simultaneous <- NULL
      
      cat(sprintf('--------------Distance: %.2g\n', distance_curr))
      
      for (j_cv in 1:n_cv) {
        
        cat(sprintf('             year: %d\n', j_cv))
        
        #---Current map
        tb_results <- tb_results_curr[j_cv,]
        
        map_for_ROC <- tb_results$map_for_ROC[[1]]
        
        year_cv <- unique(map_for_ROC$year_idx)
        
        if (length(year_cv) != 1)
          stop("cv error")
        
        year_prev <- year_cv - 1
        
        if (year_prev < 1)
          stop("year error")
        
        
        
        if (!is.na(distance_curr)) {
          
          #---Subset to regions within distance
          setup <- setup_for_distance_to_presence(data_spde__, n_points=2.5e4, year_idx_=year_prev)
          
          map_close <- regions_close_to_presence(setup, distance_threshold=distance_curr)
          
          geo_subset <- map_close$geo
          
          map_for_ROC <- map_for_ROC %>% filter(geo %in% geo_subset)
          
          map_for_ROC_simultaneous <- bind_rows(map_for_ROC_simultaneous, map_for_ROC)
        }
        
        #---Compute ROC
        roc_curve_long <- NULL
        model_prob_long <- NULL
        
        output <- compute_roc_for_cv_set(map_for_ROC, 
                                         roc_curve_long=roc_curve_long, 
                                         model_prob_long=model_prob_long,
                                         model_name=model_curr,
                                         plot_curve=FALSE)
        
        if (!is.null(output$roc_curve_long)) {
          roc_curve_long <- output$roc_curve_long
          auc <- as.double(output$auc)
          
        } else {
          auc <- NA
        }
        
        # #---Display AUC
        # cat(sprintf("AUC on test set: %.3g\n", output$auc))
        
        
        #---Save to inner table
        tb_roc_inner[j_cv,] <- tibble(year_cv=year_cv, 
                                      auc=auc, 
                                      roc_curve=list(roc_curve_long),
                                      n_regions=nrow(map_for_ROC))
        print(tb_roc_inner)
      }
      
      
      
      #---All cv-years simultaneous
      roc_curve_long <- NULL
      model_prob_long <- NULL
      
      output <- compute_roc_for_cv_set(map_for_ROC_simultaneous, 
                                       roc_curve_long=roc_curve_long, 
                                       model_prob_long=model_prob_long,
                                       model_name=model_curr,
                                       plot_curve=FALSE)
      
      if (!is.null(output$roc_curve_long)) {
        roc_curve_simultaneous <- output$roc_curve_long
        auc_simultaneous <- as.double(output$auc)
        
      } else {
        roc_curve_simultaneous <- NA
        auc_simultaneous <- NA
      }
      
      tb_roc_simultaneous <- tibble(auc=auc_simultaneous, 
                                    roc_curve=list(roc_curve_simultaneous),
                                    n_regions=nrow(map_for_ROC_simultaneous))
      
      
      
      #---Save to outer table
      tb_roc <- bind_rows(tb_roc, tibble(model_str=model_curr, distance=distance_curr, 
                                         roc=list(tb_roc_inner), roc_simultaneous=list(tb_roc_simultaneous)))
      print(tb_roc_inner)
    }
    
  }
  
  return(tb_roc)
}


roc_subsetted_by_year <- function(tb_results_cumul) {
  
  models_vec <- tb_results_cumul$model_str %>% str_extract(".*(?=_cv)") %>% unique()
  
  tb_roc <- NULL
  
  for (model_curr in models_vec) {
    
    #---Fits for current model
    tb_results_curr <- tb_results_cumul %>% filter(str_detect(model_str, model_curr))
    
    #---Setup
    n_cv <- nrow(tb_results_curr)
    tb_roc_inner <- tibble(year_cv=rep(NA, n_cv), auc=NA, roc_curve=NA)
    map_for_ROC_simultaneous <- NULL
    
    for (j_cv in 1:n_cv) {
      
      #---Current map
      tb_results <- tb_results_curr[j_cv,]
      
      map_for_ROC <- tb_results$map_for_ROC[[1]]
      
      map_for_ROC_simultaneous <- bind_rows(map_for_ROC_simultaneous, map_for_ROC)
      
      year_cv <- unique(map_for_ROC$year_idx)
      
      if (length(year_cv) != 1)
        stop("cv error")
      
      year_prev <- year_cv - 1
      
      if (year_prev < 1)
        stop("year error")
      
      
      
      #---Compute ROC
      roc_curve_long <- NULL
      model_prob_long <- NULL
      
      output <- compute_roc_for_cv_set(map_for_ROC, 
                                       roc_curve_long=roc_curve_long, 
                                       model_prob_long=model_prob_long,
                                       model_name=model_curr,
                                       plot_curve=FALSE)
      
      model_prob_long <- output$model_prob_long
      
      if (!is.null(output$roc_curve_long)) {
        roc_curve_long <- output$roc_curve_long
        auc <- as.double(output$auc)
        
      } else {
        auc <- NA
      }
      
      # #---Display AUC
      # cat(sprintf("AUC on test set: %.3g\n", output$auc))
      
      
      #---Save to inner table
      tb_roc_inner[j_cv,] <- tibble(year_cv=year_cv, 
                                    auc=auc, 
                                    roc_curve=list(roc_curve_long))
    }
    
    
    
    #---All cv-years simultaneous
    roc_curve_long <- NULL
    model_prob_long <- NULL
    
    output <- compute_roc_for_cv_set(map_for_ROC_simultaneous, 
                                     roc_curve_long=roc_curve_long, 
                                     model_prob_long=model_prob_long,
                                     model_name=model_curr,
                                     plot_curve=FALSE)
    
    if (!is.null(output$roc_curve_long)) {
      roc_curve_simultaneous <- output$roc_curve_long
      auc_simultaneous <- as.double(output$auc)
      
    } else {
      roc_curve_simultaneous <- NA
      auc_simultaneous <- NA
    }
    
    tb_roc_simultaneous <- tibble(auc=auc_simultaneous, 
                                  roc_curve=list(roc_curve_simultaneous),
                                  n_regions=nrow(map_for_ROC_simultaneous))
    
    
    
    #---Save to outer table
    tb_roc <- bind_rows(tb_roc, tibble(model_str=model_curr, roc=list(tb_roc_inner),
                                       roc_simultaneous=list(tb_roc_simultaneous)))
  }
  
  
  return(tb_roc)
}


roc_subsetted_by_class <- function(tb_results_cumul, n_cv, tb_class, tb_class_time=NULL) {
  
  models_vec <- tb_results_cumul$model_str %>% str_extract(".*(?=_cv)") %>% unique()
  
  tb_roc <- NULL
  
  if (is.null(tb_class_time))
    tb_class_time <- tibble(class="period_full", year_idx=list(1:n_cv))
  
  for (model_curr in models_vec) {
    
    #---Fits for current model
    tb_results_curr <- tb_results_cumul %>% filter(str_detect(model_str, model_curr))
    
    for (j_class in 1:nrow(tb_class)) {
      
      for (j_class_time in 1:nrow(tb_class_time)) {
        
        #---Setup
        tb_roc_inner <- tibble(year_cv=rep(NA, n_cv), auc=NA, roc_curve=NA)
        
        roc_curve_long <- NULL
        model_prob_long <- NULL
        map_for_ROC_simultaneous <- NULL
        
        cv_years <- intersect(1:n_cv, tb_class_time$year_idx[[j_class_time]])
        
        for (j_cv in cv_years) {
          
          #---Current map
          tb_results <- tb_results_curr[j_cv,]
          
          map_for_ROC <- tb_results$map_for_ROC[[1]]
          
          year_cv <- unique(map_for_ROC$year_idx)
          
          if (length(year_cv) != 1)
            stop("cv error")
          
          year_prev <- year_cv - 1
          
          if (year_prev < 1)
            stop("year error")
          
          
          
          #---Subset to regions in current class
          map_for_ROC <- map_for_ROC %>% filter(geo %in% tb_class$geo[[j_class]])
          
          map_for_ROC_simultaneous <- bind_rows(map_for_ROC_simultaneous, map_for_ROC)
          
          
          
          #---Compute ROC
          output <- compute_roc_for_cv_set(map_for_ROC, 
                                           roc_curve_long=roc_curve_long, 
                                           model_prob_long=model_prob_long,
                                           model_name=model_curr,
                                           plot_curve=FALSE)
          
          model_prob_long <- output$model_prob_long
          
          if (!is.null(output$roc_curve_long)) {
            roc_curve_long <- output$roc_curve_long
            auc <- as.double(output$auc)
            
          } else {
            
            auc <- NA
          }
          
          # #---Display AUC
          # cat(sprintf("AUC on test set: %.3g\n", output$auc))
          
          
          #---Save to inner table
          tb_roc_inner[j_cv,] <- tibble(year_cv=year_cv, 
                                        auc=auc, 
                                        roc_curve=list(roc_curve_long))
        }
        
        
        
        
        #---All cv-years simultaneous
        roc_curve_long <- NULL
        model_prob_long <- NULL
        
        output <- compute_roc_for_cv_set(map_for_ROC_simultaneous, 
                                         roc_curve_long=roc_curve_long, 
                                         model_prob_long=model_prob_long,
                                         model_name=model_curr,
                                         plot_curve=FALSE)
        
        if (!is.null(output$roc_curve_long)) {
          roc_curve_simultaneous <- output$roc_curve_long
          auc_simultaneous <- as.double(output$auc)
          
        } else {
          roc_curve_simultaneous <- NA
          auc_simultaneous <- NA
        }
        
        tb_roc_simultaneous <- tibble(auc=auc_simultaneous, 
                                      roc_curve=list(roc_curve_simultaneous),
                                      n_regions=nrow(map_for_ROC_simultaneous))
        
        
        
        
        #---Save to outer table
        tb_roc <- bind_rows(tb_roc, tibble(model_str=model_curr, 
                                           class=tb_class$class[j_class], 
                                           class_time=tb_class_time$class[j_class_time],
                                           roc=list(tb_roc_inner),
                                           roc_simultaneous=list(tb_roc_simultaneous)))
        
      }
      
    }
  }
  
  
  return(tb_roc)
}
```
