## Supplementary data and code for "A climate and population dependent diffusion model forecasts the spread of *Aedes Albopictus* mosquitoes in Europe": functions_plot_figures.R

```
plot_figure_5 <- function(file="./Data/recreate_figure_3_to_5/figure_5.RData", 
                          folder_plots=".") {
  
  #---Load data
  out <- load(file=file)
  
  ps <- list()
  
  #---Median temperature
  print(category_median_temp)
  
  x_column <- "median_temp"
  y_column <- "presence"
  
  ps[[length(ps)+1]] <- plot_scatter_lines(category_median_temp, x_column, y_column, 
                                           color_column="median_temp", 
                                           point_size=8,
                                           plot_lines=FALSE,
                                           plot_larger=TRUE)
  print(ps[[length(ps)]])
  save_to_png(ps[[length(ps)]], 
              filename=paste0(folder_plots, "/grouping_presence_median_temp.png"),
              resolution_scaling_width=0.5)
  
  
  
  
  
  
  
  #---Median temperature and log population
  print(category_median_temp_and_log_pop)
  
  grouping_column <- "category_median_temp"
  x_column <- "log_population"
  y_column <- "weighted_presence"
  out <- create_segments(category_median_temp_and_log_pop, 
                         x_column, y_column, grouping_column)
  
  ps[[length(ps)+1]] <- plot_scatter_lines(out$df_grouped, x_column, y_column, 
                                           color_column="median_temp", 
                                           plot_segments = TRUE,
                                           x_segment_columns=out$x_segment_columns,
                                           y_segment_columns=out$y_segment_columns,
                                           plot_lines=FALSE,
                                           plot_larger=TRUE)
  print(ps[[length(ps)]])
  save_to_png(ps[[length(ps)]], 
              filename=paste0(folder_plots, "/grouping_WEIGHTEDpresence_log_population.png"),
              resolution_scaling_width=0.5)
  
  
  
  
  
  #---Median temperature and min temperature
  print(category_median_temp_and_min_temp)
  
  grouping_column <- "category_median_temp"
  x_column <- "min_temp"
  y_column <- "presence"
  out <- create_segments(category_median_temp_and_min_temp, 
                         x_column, y_column, grouping_column)
  
  ps[[length(ps)+1]] <- plot_scatter_lines(out$df_grouped, x_column, y_column, 
                                           color_column="median_temp", 
                                           plot_segments = TRUE,
                                           x_segment_columns=out$x_segment_columns,
                                           y_segment_columns=out$y_segment_columns,
                                           plot_lines=FALSE,
                                           plot_larger=TRUE)
  print(ps[[length(ps)]])
  save_to_png(ps[[length(ps)]], 
              filename=paste0(folder_plots, "/grouping_presence_min_temp.png"),
              resolution_scaling_width=0.5)
  
  
  
  
  
  
  #---Median temperature and relative humidity
  print(category_median_temp_and_rel_humidity)
  
  category_median_temp_and_rel_humidity <- 
    category_median_temp_and_rel_humidity %>% filter(n>=30)
  
  grouping_column <- "category_median_temp"
  x_column <- "median_relative_humidity"
  y_column <- "presence"
  out <- create_segments(category_median_temp_and_rel_humidity, 
                         x_column, y_column, grouping_column)
  
  ps[[length(ps)+1]] <- plot_scatter_lines(out$df_grouped, x_column, y_column,
                                           color_column="median_temp",
                                           plot_segments = TRUE,
                                           x_segment_columns=out$x_segment_columns,
                                           y_segment_columns=out$y_segment_columns,
                                           plot_lines=FALSE,
                                           plot_larger=TRUE,
                                           color_limits=c(16,27))
  print(ps[[length(ps)]])
  save_to_png(ps[[length(ps)]], 
              filename=paste0(folder_plots, "/grouping_presence_relative_humidity.png"),
              resolution_scaling_width=0.5)
  
  
  
  
  
  # Mechanistic model (ODE)
  print(category_mechanistic_model)
  
  x_column <- "mean_adults_pow"
  y_column <- "presence"
  
  ps[[length(ps)+1]] <- plot_scatter_lines(category_mechanistic_model, 
                                           x_column, y_column, 
                                           color_column="mean_adults_pow", 
                                           point_size=8,
                                           plot_lines=FALSE,
                                           plot_larger=TRUE)
  print(ps[[length(ps)]])
  save_to_png(ps[[length(ps)]], 
              filename=paste0(folder_plots, "/grouping_ODE_presence_ODE__new_establishments.png"),
              resolution_scaling_width=0.5)
  
  
  
  
  
  
  #---Mechanistic model (ODE) and proximity
  print(category_mechanistic_model_and_proximity)
  
  grouping_column <- "category_ODE"
  x_column <- "proximity"
  y_column <- "presence"
  out <- create_segments(category_mechanistic_model_and_proximity, 
                         x_column, y_column, grouping_column)
  
  ps[[length(ps)+1]] <- 
    plot_scatter_lines(out$df_grouped, x_column, y_column, 
                       color_column="mean_adults_pow", 
                       plot_segments = TRUE,
                       x_segment_columns=out$x_segment_columns,
                       y_segment_columns=out$y_segment_columns,
                       plot_lines=FALSE,
                       plot_larger=TRUE)
  print(ps[[length(ps)]])
  save_to_png(ps[[length(ps)]], 
              filename=paste0(folder_plots, "/grouping_ODE_presence_proximity", 
                              "__only_new_establishments.png"),
              resolution_scaling_width=0.5)
  
  return(ps)
}


plot_figure_3 <- function(folder_plots = ".", 
                          file_climate_and_pop_GAM = 
                            paste0("./Data/recreate_figure_3_to_5/", 
                                   "random_effects_on_map__climate_and_population_GAM_model.RData"), 
                          file_mechanistic_GAM = 
                            paste0("./Data/recreate_figure_3_to_5/", 
                                   "random_effects_on_map__mechanistic_GAM_model.RData")) {
  
  
  ps <- list()
  
  # Plot random effects for the climate and population GAM model
  out <- load(file=file_climate_and_pop_GAM)
  
  # Plot random effect for median temperature
  ps[[length(ps)+1]] <- 
    plot_sf(map_all %>% filter(year_idx==1) %>% left_join(map_spde) %>% # the fitted random effect is the same for each year, select year_idx 1
              rename(reffect_median_temp = reffect_bin_idx_median_temp.mean), 
            "reffect_median_temp",
            title="random effect, median temp",
            plot_larger=plot_larger,
            color_limits=NULL)
  print(ps[[length(ps)]])
  save_to_png(ps[[length(ps)]], 
              filename=paste0(folder_plots, "/reffect_median_temp_on_map.png"),
              resolution_scaling_width=1)
  
  # Plot random effect for min. temperature CROSS max. temperature, with NA where 
  # it is undefined (where the max. temperature is low)
  ps[[length(ps)+1]] <- 
    plot_sf(map_all %>% filter(year_idx==1) %>% left_join(map_spde) %>% # the fitted random effect is the same for each year, select year_idx 1
              rename(reffect_min_temp_CROSS = reffect_bin_idx_min_temp_CROSS.meanNA), 
            "reffect_min_temp_CROSS",
            title="random effect, min temp CROSS",
            plot_larger=plot_larger,
            color_limits=NULL)
  print(ps[[length(ps)]])
  save_to_png(ps[[length(ps)]], 
              filename=paste0(folder_plots, "/reffect_min_temp_CROSS_on_map.png"),
              resolution_scaling_width=1)
  
  # Plot random effect for relative humidity
  ps[[length(ps)+1]] <- 
    plot_sf(map_all %>% filter(year_idx==1) %>% left_join(map_spde) %>% # the fitted random effect is the same for each year, select year_idx 1
              rename(reffect_median_relative_humidity = 
                       reffect_bin_idx_median_relative_humidity.mean), 
            "reffect_median_relative_humidity",
            title="random effect, median relative humidity",
            plot_larger=plot_larger,
            color_limits=NULL)
  print(ps[[length(ps)]])
  save_to_png(ps[[length(ps)]], 
              filename=paste0(folder_plots, "/reffect_median_relative_humidity_on_map.png"),
              resolution_scaling_width=1)
  
  
  
  
  
  # Plot random effects for the mechanistic GAM model
  out <- load(file=file_mechanistic_GAM)
  
  #---Plot random effect for the mechanistic covariate
  ps[[length(ps)+1]] <- 
    plot_sf(map_all %>% filter(year_idx==1) %>% left_join(map_spde) %>% # the fitted random effect is the same for each year, select year_idx 1
              rename(reffect_mechanistic_covariate = 
                       reffect_bin_idx_mean_adults_pow_merged.mean), 
            "reffect_mechanistic_covariate",
            title="random effect, mechanistic covariate",
            plot_larger=plot_larger,
            color_limits=NULL)
  print(ps[[length(ps)]])
  save_to_png(ps[[length(ps)]], 
              filename=paste0(folder_plots, "/reffect_mechanistic_covariate_on_map.png"),
              resolution_scaling_width=1)
  
  # # CANNOT SHARE PROXIMITY COVARIATE SINCE IT SHOWS WHERE Ae. albopictus WAS PRESENT THE PREVIOUS YEARS
  # #---Plot proximity at year 2024
  # map_all$year <- map_all$year_idx + 2009 # convert year_idx to the actual year
  # proximity_year <- 2024 # year to plot
  # 
  # # set proximity to NA in regions where Ae. albopictus was present the previous year
  # map_all <- map_all %>% mutate(reffect_bin_idx_proximity050.meanNA=
  #                                 if_else(presence_previous_year==1, NA,
  #                                         reffect_bin_idx_proximity050.mean))
  # 
  # ps[[length(ps)+1]] <- 
  #   plot_sf(map_all %>% filter(year==!!proximity_year) %>% left_join(map_spde) %>%
  #             rename(reffect_proximity = reffect_bin_idx_proximity050.meanNA), 
  #           "reffect_proximity",
  #           title=paste0("random effect, proximity, year ", proximity_year),
  #           plot_larger=plot_larger,
  #           color_limits=NULL)
  # print(ps[[length(ps)]])
  # save_to_png(ps[[length(ps)]], 
  #             filename=paste0(folder_plots, "/reffect_proximity_year_", 
  #                             proximity_year, "_on_map.png"),
  #             resolution_scaling_width=1)
  
  return(ps)
}


create_segments <- function(category_all_tot, x_column, y_column, grouping_column) {
  x_segment_columns <- list(start="x_segment_start",
                            end="x_segment_end")
  y_segment_columns <- list(start="y_segment_start",
                            end="y_segment_end")
  
  df_grouped <- category_all_tot %>% group_by(!!sym(grouping_column)) %>% 
    mutate(group_idx=1:n()) %>% relocate(group_idx)
  
  df_grouped <- bind_cols(df_grouped, tibble(x_segment_start=NA,
                                             x_segment_end=NA,
                                             y_segment_start=NA,
                                             y_segment_end=NA))
  for (j in 1:nrow(df_grouped)) {
    group_idx_curr <- df_grouped$group_idx[j]
    grouping_value_curr <- df_grouped[[grouping_column]][j]
    
    next_row <- df_grouped %>% filter(group_idx==group_idx_curr+1,
                                      !!sym(grouping_column)==grouping_value_curr)
    
    if (nrow(next_row)==1) {
      df_grouped$x_segment_start[j] <- df_grouped[[x_column]][j]
      df_grouped$x_segment_end[j] <- next_row[[x_column]]
      
      df_grouped$y_segment_start[j] <- df_grouped[[y_column]][j]
      df_grouped$y_segment_end[j] <- next_row[[y_column]]
    } else if (nrow(next_row)>1) {
      stop("error, next_row should have either one or zero rows")
    }
  }
  
  return(list(df_grouped=df_grouped, 
              x_segment_columns=x_segment_columns,
              y_segment_columns=y_segment_columns))
}
```
