## Supplementary data and code for "A climate and population dependent diffusion model forecasts the spread of *Aedes Albopictus* mosquitoes in Europe": run_plots_from_paper.R

```
# options(error = recover)

out <- load(file="Data/map.RData")


source("functions.R") # general functions
source("functions_plotting.R") # general functions for plotting
source("functions_plot_figures.R") # functions specific for plotting figures for the paper

plot_larger <- TRUE


###########################################################
###########################################################
# Plot Figure 3 ###########################################

# Plots random effects on map, saves to folder 'Plots'
# NOTE: can't share the full proximity covariate since it shows where 
# Ae. albopictus was present the previous years and we cannot share 
# Ae. albopictus presence / absence data. Therefore the map showing the 
# proximity random effect can't be recreated here.
ps <- plot_figure_3(folder_plots = "./Plots", 
                    file_climate_and_pop_GAM = 
                      paste0("./Data/recreate_figure_3_to_5/", 
                             "random_effects_on_map__climate_and_population_GAM_model.RData"), 
                    file_mechanistic_GAM = 
                      paste0("./Data/recreate_figure_3_to_5/", 
                             "random_effects_on_map__mechanistic_GAM_model.RData"))

ps[[1]]
ps[[2]]
ps[[3]]
ps[[4]]


###########################################################
###########################################################
# Plot Figure 4 ###########################################

out <- load(file="./Data/recreate_figure_3_to_5/figure_4.RData")

out <- load(file=paste0("./Data/recreate_figure_3_to_5/",
                        "random_effects_functional_form__climate_and_population_GAM_model.RData"))
#---Plots random effects + empirical Ae. albopictus ratios for 4-way grouping
#---of the data (grouping represented by category_4way_grouping) and for 
#---the climate and population GAM model
#---All plots are saved to file (folder 'Plots') but only random effect-plots are 
#---returned in 'ps'. Empirical Ae. albopictus ratio-plots are saved with zero
#---opacity of the background to be able to overlay on random effect-plots.
ps <- plot_random_effects_as_functions__(functions_random_effects, keys, 
                                         model_str="climate_and_population_GAM",
                                         categories=categories_4way_grouping,
                                         plot_larger=plot_larger,
                                         ylim=ylim,
                                         resolution_scaling_width=0.75,
                                         folder_plots = "./Plots")

# Plot fitted random effects
grid.draw(ps[[1]][[1]])
grid.draw(ps[[1]][[2]])
grid.draw(ps[[1]][[3]])
grid.draw(ps[[1]][[4]])
grid.draw(ps[[1]][[5]])

out <- load(file=paste0("./Data/recreate_figure_3_to_5/",
                        "random_effects_functional_form__mechanistic_GAM_model.RData"))
#---Same as above, but here for the mechanistic GAM model
ps <- plot_random_effects_as_functions__(functions_random_effects, keys,
                                         model_str="mechanistic_GAM_model",
                                         categories=categories_4way_grouping,
                                         plot_larger=plot_larger,
                                         ylim=c(-3,4.5),
                                         resolution_scaling_width=0.75,
                                         folder_plots = "./Plots")

# Plot fitted random effects
grid.draw(ps[[1]][[1]])
grid.draw(ps[[1]][[2]])


###########################################################
###########################################################
# Plot Figure 5 ###########################################

#---Plots empirical Ae. albopictus ratios for 1- and 2-way groupings
#---of the data. Saves to folder 'Plots'. Groupings are loaded from 
#---"figure_5.Rdata".
ps <- plot_figure_5(file="./Data/recreate_figure_3_to_5/figure_5.RData", 
                    folder_plots="./Plots")

#---Plot the ratios for the six groupings
ps[[1]]
ps[[2]]
ps[[3]]
ps[[4]]
ps[[5]]
ps[[6]]
```
