## Supplementary data and code for "A climate and population dependent diffusion model forecasts the spread of *Aedes Albopictus* mosquitoes in Europe": run_GAM_SPDE_model.R

```
# Run GAM SPDE model with rolling window cross-validation, for 
#     ** mechanistic GAM model ('covariate_set'==1) or climate and pop. GAM model ('covariate set'==2)
#     ** 'n_years_cv' number of years in the rolling window cross-validation
#     ** short range SPDE (short_range_spde==TRUE) or free range SPDE (short_range_spde==FALSE)
# NOTE: we can't share data on Ae. albopictus presence / absence. We can share
#     covariates. The code therefore runs on simulated data for Ae. albopictus 
#     presence / absence, but with real covariates. 


##########################################################################
# Settings ###############################################################
##########################################################################

# options(error = recover)

covariate_set <- 1 # 1: mechanistic model covariate, 2: climate and log population covariates

n_years_cv <- 3 # number of years for rolling window cross-validation

short_range_spde <- TRUE


##########################################################################
# Load ###################################################################
##########################################################################

#---Load functions and packages
source("functions.R")

#---Load data
folder_data <- "Data"

# #---Process data
# #---Data that can be shared
# out <- load(paste0(folder_data, "/data_original.RData"))
# #---Simulated data that replaces the data that cannot be shared
# out <- load(paste0(folder_data, "/data_simulated.RData"))
# 
# #---Compute proximity from data_simulated
# df_proximity <- compute_proximity(data_simulated %>% mutate(y=presence), map_spde)
# 
# #---Bin and add proximity-covariate + add year1-covariate
# out <- process_covariates(data_simulated, df_proximity, keys)
# data_spde <- out$data_spde
# keys <- out$keys
# 
# #---Join the data that can be shared with simulated data
# data_spde <- left_join(data_spde, data_original_covariates)
# 
# save("data_spde", "map_spde", "keys", file=paste0(folder_data, "/data.RData"))

#---Load already processed data
out <- load(paste0(folder_data, "/data.RData"))


##########################################################################
# Prepare for SPDE model #################################################
##########################################################################

#---SPDE mesh
out <- create_mesh(map_spde, mesh_precision="low")
spde_mesh <- out$spde_mesh
region_coords <- out$region_coords
n_spde <- out$n_spde

mesh_proj <- inla.mesh.projector(spde_mesh, loc=region_coords)

cat("Mesh, size: ", spde_mesh$n, "\n")
par(mfrow=c(1,1))
plot(spde_mesh)
plot_geometry(map_spde, add=TRUE)
points(region_coords, pch=20, col=2)

#---SPDE model
spde_model <- inla.spde2.matern(spde_mesh, alpha=2)

if (short_range_spde) {
  #---precision matrix from SPDE model
  Q_params <- c(log(1e-1),log(2)) # tuning parameters for the SPDE, controls SPDE range
  Q_spde <- inla.spde2.precision(spde_model,theta=Q_params)
}


##########################################################################
# Cross-validation #######################################################
##########################################################################

model_str_main <- sprintf("covariate_set%d", covariate_set)

if (short_range_spde) {
  model_str_main <- paste0(model_str_main, "_short_range_SPDE")
} else {
  model_str_main <- paste0(model_str_main, "_free_range_SPDE")
}

data_spde__ <- data_spde

tb_results_cumul <- NULL

for (j_cv in n_years_cv:1) {
  
  ##########################################################################
  # Prepare for INLA with SPDE model #######################################
  ##########################################################################
  
  #---Subset data for cross-validation
  unique_years <- sort(unique(data_spde__$year_idx))
  n_max <- max(unique_years)
  cv_set <- list(years_training=1:(n_max - j_cv), years_test=n_max - j_cv + 1) # subset to years 1:(n_years-j), predict for year (n_years-j+1)
  
  ind_year <- data_spde__$year_idx %in% c(cv_set$years_training, cv_set$years_test)
  
  data_spde <- data_spde__[ind_year,] %>% 
    mutate(training_set = 1*(year_idx %in% cv_set$years_training & !is.na(presence)), # inla model will be fit on training set
           test_set = 1*(year_idx %in% cv_set$years_test & !is.na(presence)), # model will be used to predict on test set
           y_orig = presence) # keep original values
  
  data_spde$presence[data_spde$test_set] <- NA # inla gives prediction where presence==NA
  
  model_str <- sprintf("%s_cv%d", model_str_main, j_cv)
  
  
  
  #---Compute stack which connectes SPDE and observations / link which is used to handle NA values
  out <- setup_for_inla(map_spde, data_spde)
  stack <- out$stack # connects SPDE and observations
  spde_indices <- out$spde_indices # used for plotting
  link <- out$link # for handling NA values
  
  
  
  
  
  ##########################################################################
  # Run INLA ###############################################################
  ##########################################################################
  
  #---Define formula
  if (covariate_set==1) {
    
    if (short_range_spde) {
      f <- presence ~ -1 + Intercept + 
        f(spde_idx, model = "generic0", Cmatrix = Q_spde, # SPDE
          group = spde_idx.group, control.group = list(model = "iid")) + 
        presence_previous_year + year1 + # previous year and year 1 covariates
        f(bin_idx_mean_adults_pow_merged, # mechanistic model covariate
          model = "rw1", scale.model = TRUE, constr = TRUE, 
          hyper = list(theta = list(prior = "pc.prec", param = c(0.5, 0.01)))) + 
        f(bin_idx_proximity050, model = "besag", scale.model = TRUE, constr = TRUE, # proximity covariate
          graph = file_nb_proximity,
          hyper = list(theta = list(prior = "pc.prec", param = c(0.5, 0.01))))
    } else {
      f <- presence ~ -1 + Intercept + 
        f(spde_idx, model = spde_model, # SPDE 
          group = spde_idx.group, control.group=list(model="iid")) + 
        presence_previous_year + year1 + # previous year and year 1 covariates
        f(bin_idx_mean_adults_pow_merged, # mechanistic model covariate
          model = "rw1", scale.model = TRUE, constr = TRUE, 
          hyper = list(theta = list(prior = "pc.prec", param = c(0.5, 0.01)))) + 
        f(bin_idx_proximity050, model = "besag", scale.model = TRUE, constr = TRUE, # proximity covariate
          graph = file_nb_proximity,
          hyper = list(theta = list(prior = "pc.prec", param = c(0.5, 0.01))))
      
      
    }
  } else {
    
    if (short_range_spde) {
      f <- presence ~ -1 + Intercept + 
        f(spde_idx, model = "generic0", Cmatrix = Q_spde, # SPDE
          group = spde_idx.group, control.group = list(model = "iid")) + 
        presence_previous_year + year1 + # previous year and year 1 covariates
        f(bin_idx_median_temp, model = "rw1", scale.model = TRUE, constr = TRUE, # median temperature covariate 
          hyper = list(theta = list(prior = "pc.prec", param = c(0.5, 0.01)))) + 
        f(bin_idx_min_temp_CROSS, model = "besag", scale.model = TRUE, constr = TRUE, # min. temp. conditioned on high median temp. covariate
          graph = file_nb_min_temp_CROSS,
          hyper = list(theta = list(prior = "pc.prec", param = c(0.5, 0.01)))) + 
        f(bin_idx_median_relative_humidity, model = "rw1", scale.model = TRUE, constr = TRUE, # median rel. humidity covariate
          hyper = list(theta = list(prior = "pc.prec", param = c(0.5, 0.01)))) + 
        f(bin_idx_log_pop_merged, model = "rw1", scale.model = TRUE, constr = TRUE, # log population covariate
          hyper = list(theta = list(prior = "pc.prec", param = c(0.5, 0.01)))) + 
        f(bin_idx_proximity050, model = "besag", scale.model = TRUE, constr = TRUE, # proximity covariate
          graph = file_nb_proximity,
          hyper = list(theta = list(prior = "pc.prec", param = c(0.5, 0.01))))
    } else {
      
      f <- presence ~ -1 + Intercept + 
        f(spde_idx, model = spde_model, # SPDE 
          group = spde_idx.group, control.group=list(model="iid")) + 
        presence_previous_year + year1 + # previous year and year 1 covariates
        f(bin_idx_median_temp, model = "rw1", scale.model = TRUE, constr = TRUE, # median temperature covariate 
          hyper = list(theta = list(prior = "pc.prec", param = c(0.5, 0.01)))) + 
        f(bin_idx_min_temp_CROSS, model = "besag", scale.model = TRUE, constr = TRUE, # min. temp. conditioned on high median temp. covariate
          graph = file_nb_min_temp_CROSS,
          hyper = list(theta = list(prior = "pc.prec", param = c(0.5, 0.01)))) + 
        f(bin_idx_median_relative_humidity, model = "rw1", scale.model = TRUE, constr = TRUE, # median rel. humidity covariate
          hyper = list(theta = list(prior = "pc.prec", param = c(0.5, 0.01)))) + 
        f(bin_idx_log_pop_merged, model = "rw1", scale.model = TRUE, constr = TRUE, # log population covariate
          hyper = list(theta = list(prior = "pc.prec", param = c(0.5, 0.01)))) + 
        f(bin_idx_proximity050, model = "besag", scale.model = TRUE, constr = TRUE, # proximity covariate
          graph = file_nb_proximity,
          hyper = list(theta = list(prior = "pc.prec", param = c(0.5, 0.01))))
    }
    
    #---Graph for bin_idx_min_temp_CROSS
    nb_grid_min_temp_CROSS <- 
      specify_network_by_nodes_and_links(n_nodes=length(unique(data_spde__[["bin_idx_min_temp_CROSS"]])), 
                                         nodes_to_disconnect=1) # min_temp_CROSS has one node (1) that is not connected to the other nodes in the graph, i.e., to nodes 2:n
    
    file_nb_min_temp_CROSS <- paste0(folder_data, "/graph_min_temp_CROSS.adj")
    nb2INLA(file_nb_min_temp_CROSS, nb_grid_min_temp_CROSS)
  } 
  
  #---Graph for bin_idx_proximity050
  proximity_col <- "bin_idx_proximity050"
  nb_grid_proximity <- 
    specify_network_by_nodes_and_links(n_nodes=length(unique(data_spde__[[proximity_col]])), 
                                       nodes_to_disconnect=1:2) # proximity has two nodes (1:2) that are not connected to the other nodes in the graph, i.e., to nodes 3:n 
  file_nb_proximity <- paste0(folder_data, "/graph_proxmity.adj")
  nb2INLA(file_nb_proximity, nb_grid_proximity)
  
  #---Run INLA
  fitted_model <- inla(formula = f, data = inla.stack.data(stack),
                       family = "binomial", Ntrials=1,
                       control.family = list(link = "logit"),
                       control.compute = list(dic = TRUE, cpo = TRUE, waic = TRUE),
                       control.predictor = list(A = inla.stack.A(stack),
                                                link = link,
                                                compute = TRUE),
                       control.inla = list(strategy = "simplified.laplace", int.strategy = "auto"),
                       verbose = F)
  
  summary(fitted_model)
  
  
  
  
  
  ##########################################################################
  # Summarize fit ##########################################################
  ##########################################################################
  
  processed_model <- process_fitted_model(data_spde, fitted_model, covariate_set,
                                          spde_indices, stack, mesh_proj)
  
  map_full_model <- processed_model$map_full_model
  map_random_effects <- processed_model$map_random_effects
  map_random_effects_with_time <- processed_model$map_random_effects_with_time
  functions_random_effects <- processed_model$functions_random_effects
  results_summary <- processed_model$results_summary
  
  
  
  
  
  ##########################################################################
  # Plot fit ###############################################################
  ##########################################################################
  
  #---Plot random effects + selected model components on map
  year_to_plot_vec <- c(2,max(data_spde$year_idx)) # select years to plot
  
  for (year_to_plot in year_to_plot_vec) {
    #---Plot on map
    out <- plot_on_map_selected_model_parts(map_random_effects,
                                            map_random_effects_with_time,
                                            map_full_model,
                                            map_spde,
                                            year_to_plot=year_to_plot)
    
    #---Save joint plot to file
    save_to_png(out$p,
                filename=paste0("Plots/INLA_fitted_model_", model_str,
                                "_map_random_effects_and_prob_year_", year_to_plot, ".png"))
    
    #---Display joint plot
    print(out$p) # look at zoomed plot to see the subfigures
    
    #---Display separate plots
    print(out$ps_map[[1]])
    print(out$ps_map[[2]])
    #...
  }
  
  
  
  #---Plot random effects in functional form
  if (covariate_set==1) {
    ylim=c(-4,4.5)
  } else {
    ylim=c(-3,3)
  }
  
  ps <- plot_random_effects_as_functions(functions_random_effects, keys, data_spde,
                                         model_str=model_str, ylim=ylim)
  
  #---Collect in one joint plot
  if (covariate_set==1) {
    
    p <- grid.arrange(
      grobs = ps[[1]],
      widths = c(1, 1),
      layout_matrix = rbind(c(1, 2))
    )
  } else {
    
    p <- grid.arrange(
      grobs = ps[[1]],
      widths = c(1, 1, 1),
      layout_matrix = rbind(c(1, 2, 3),
                            c(4, 5, NA))
    ) # look at zoomed plot to see the subfigures
  }
  
  #---Save joint plot
  save_to_png(p, filename=paste0("Plots/INLA_fitted_model_", model_str,
                                 "_function_random_effects.png"))
  
  #---Display separate plots
  grid.draw(ps[[1]][[1]])
  grid.draw(ps[[1]][[2]])
  #...
  
  
  
  
  
  ##########################################################################
  # Save to table ##########################################################
  ##########################################################################
  
  #---Select new observations in test set, will be used to compute ROC curve
  map_for_ROC <- 
    map_full_model %>% filter(test_set==TRUE, !is.na(y_orig)) %>% # select test set, regions with observations
    select(geo,year_idx,training_set,test_set,y_orig,y,p.mean) %>% 
    left_join(data_spde %>% select(geo,year_idx,presence_previous_year)) %>% 
    filter(presence_previous_year==0) # select new observations (observations where Ae. albopictus was not present the previous year)
  
  #---Save main results to table
  tb_results_cumul <- bind_rows(
    tb_results_cumul,
    bind_cols(tibble(model_str=model_str, 
                     cv_set=list(cv_set), 
                     map_for_ROC=list(map_for_ROC)), 
              results_summary %>% relocate(DIC, WAIC, logscore, 
                                           matches("deviance"), matches("error")))
  )
  
  save(tb_results_cumul, file=paste0("Output/results_", model_str_main, ".RData"))
}


##########################################################################
# Summarize for all fitted models ########################################
##########################################################################

#---Summarise deviance
if (covariate_set==1) {
  cat("Fitted model on mechanistic model covariate-set")
} else {
  cat("Fitted model on climate and population covariate-set")
}
if (short_range_spde) {
  cat(" with short range-SPDE\n")
} else {
  cat(" with free range-SPDE\n")
}

mean_deviance <- mean(tb_results_cumul$total_deviance)
cat(sprintf("Mean deviance on full set: %.4g\n", mean_deviance))

mean_deviance_test <- mean(tb_results_cumul$total_deviance_cv)
cat(sprintf("Mean deviance on test set: %.4g\n", mean_deviance_test))


#---Compute ROC and AUC
output <- compute_roc_and_auc(tb_results_cumul)

#---Display AUC
cat(sprintf("AUC on test set: %.3g\n", output$auc))

#---Plot ROC
p <- plot_roc_curve(output$tb_roc_curve) 
print(p)
save_to_png(p, filename=paste0("Plots/ROC_curve_", model_str_main, ".png"), 
            resolution_scaling_width=1, resolution_scaling_height=1.5)
```
