## Supplementary figures and images for "A climate and population dependent diffusion model forecasts the spread of *Aedes Albopictus* mosquitoes in Europe"

### climate_and_population_GAM_log_pop_merged.png

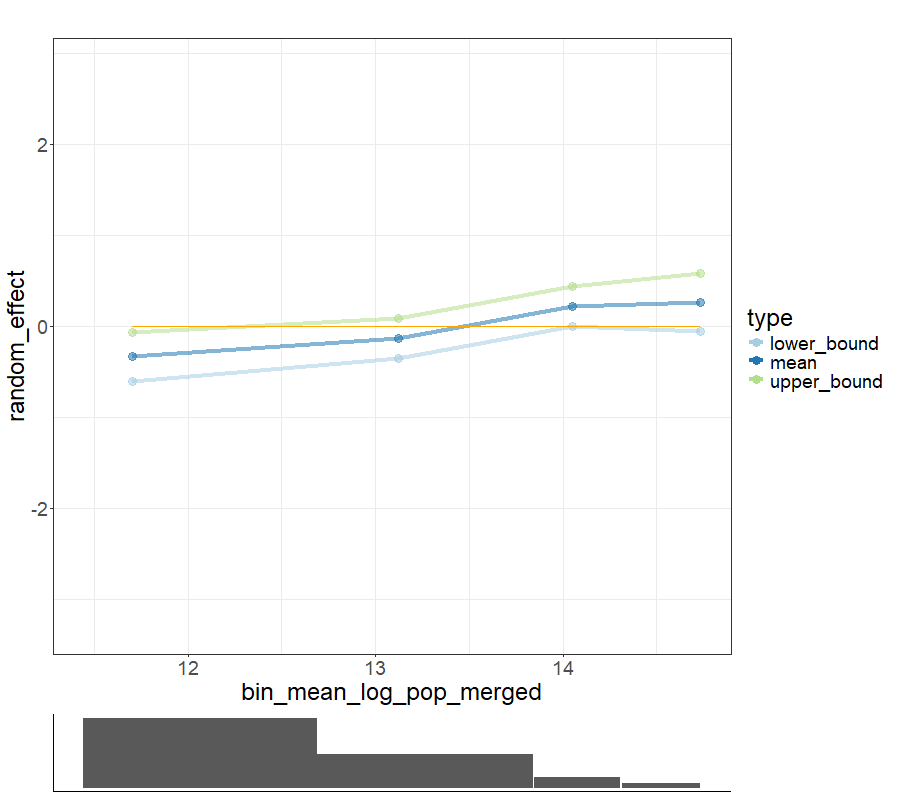

### climate_and_population_GAM_log_pop_merged_categories.png

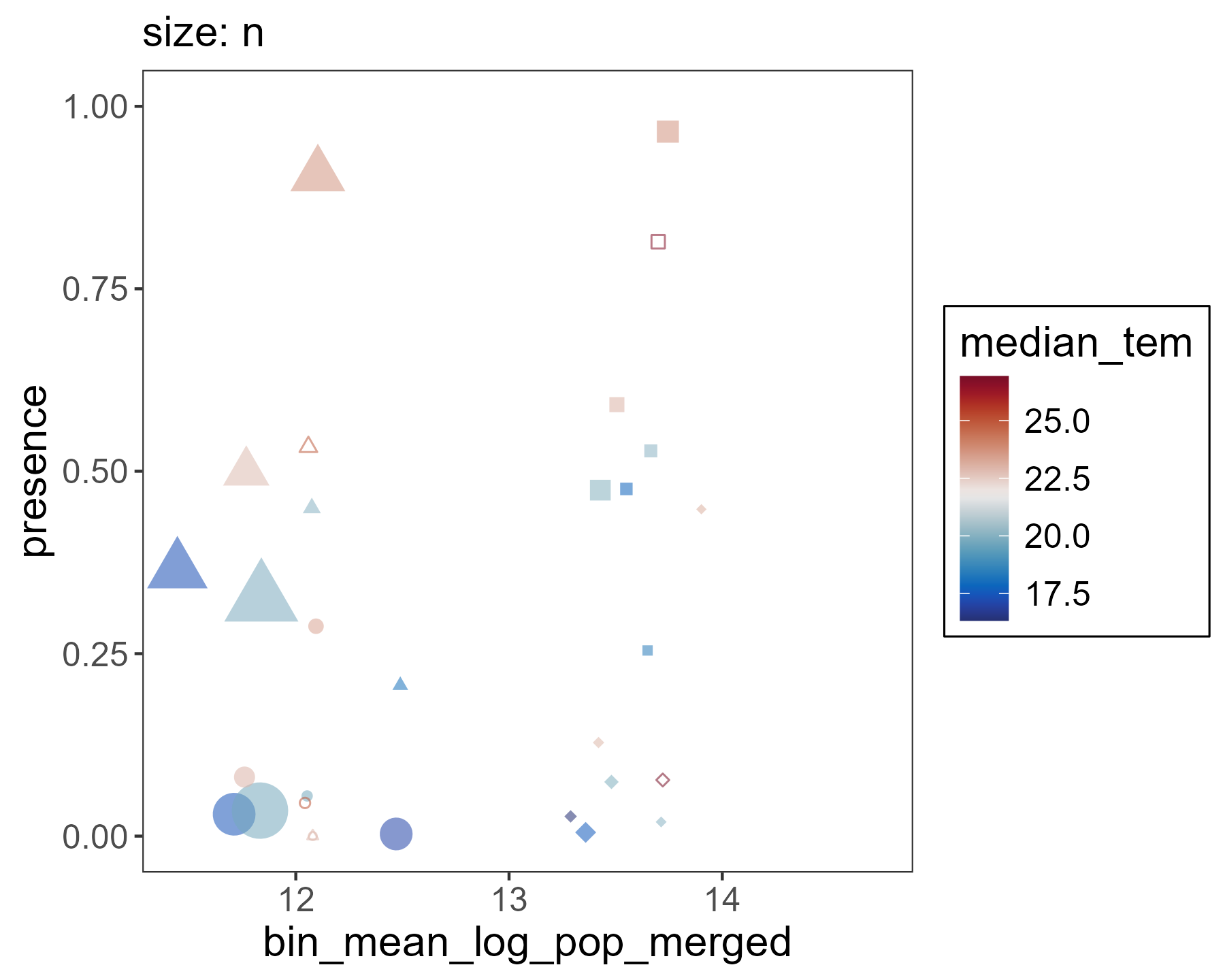

### climate_and_population_GAM_log_pop_merged_categories_with_segments.png

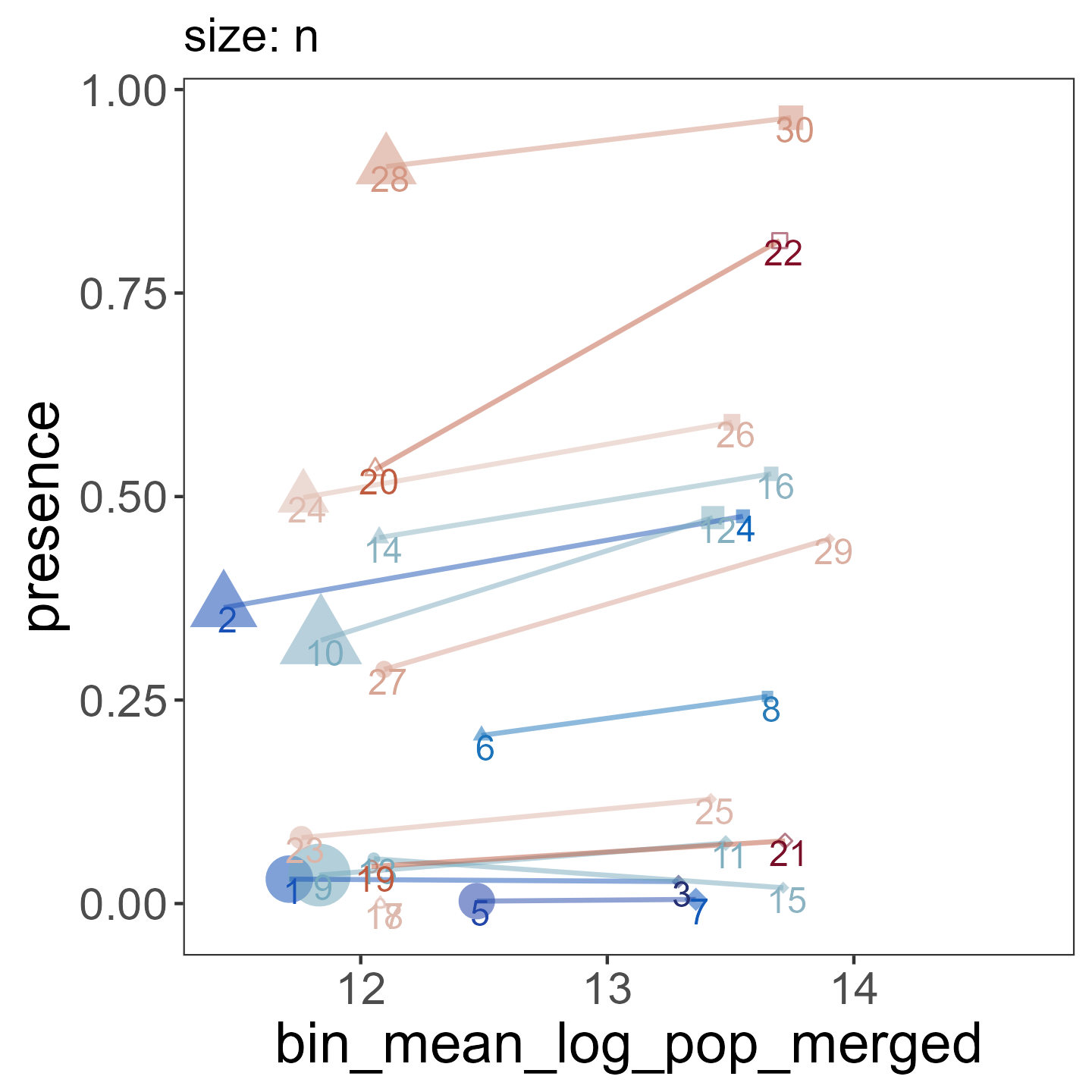

### climate_and_population_GAM_median_relative_humidity_reffect.png

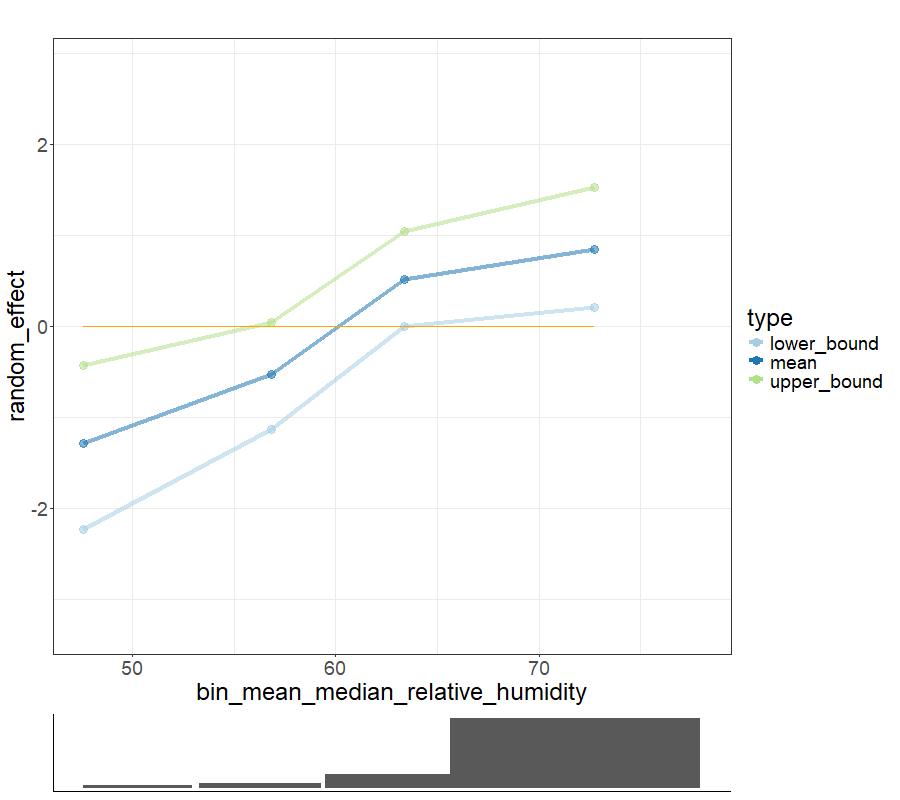

### climate_and_population_GAM_median_relative_humidity_reffect_categories.png

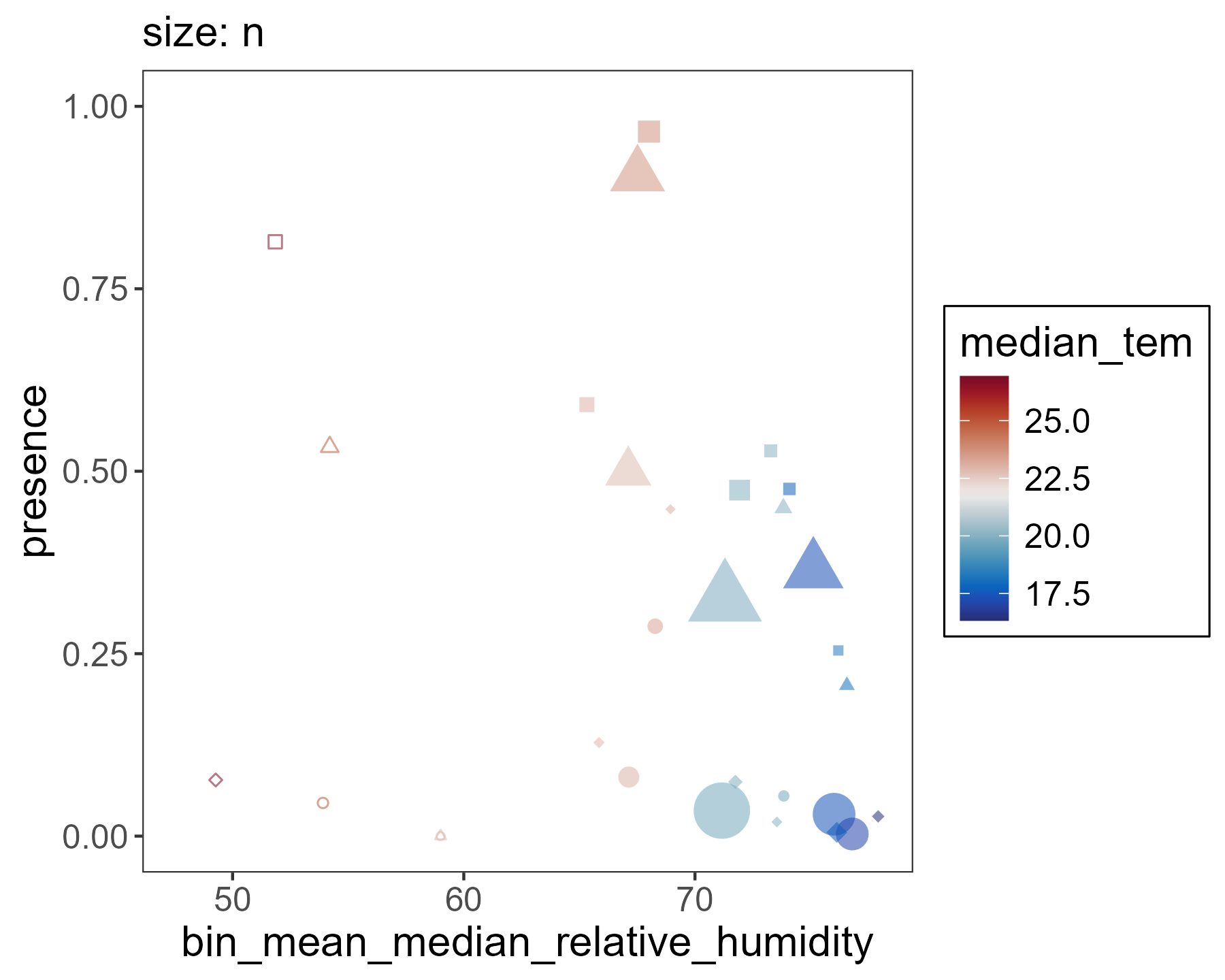

### climate_and_population_GAM_median_relative_humidity_reffect_categories_with_segments.png

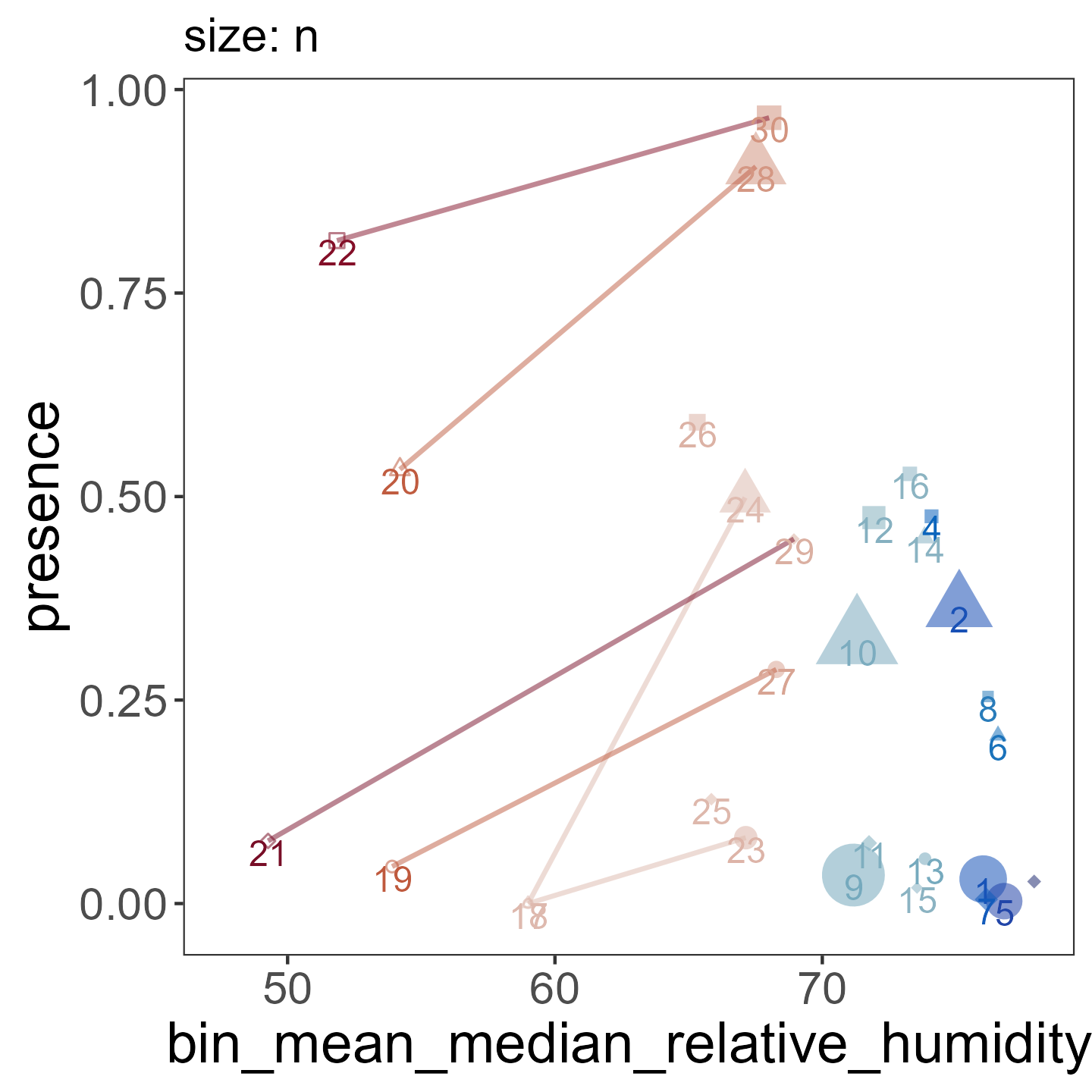

### climate_and_population_GAM_median_temp_reffect.png

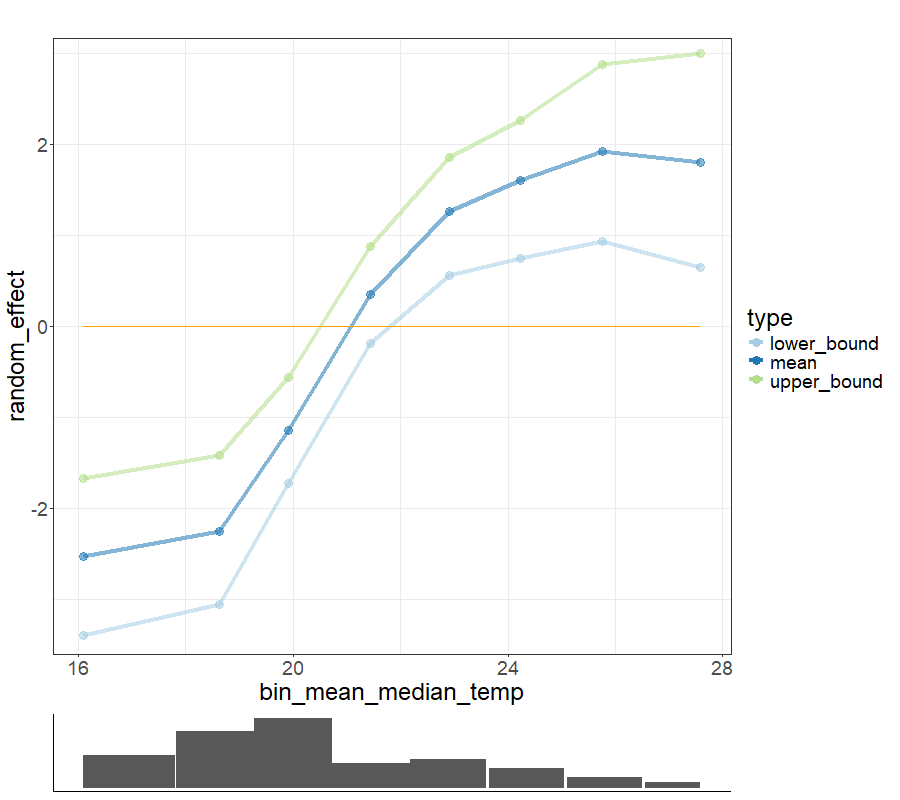

### climate_and_population_GAM_median_temp_reffect_categories.png

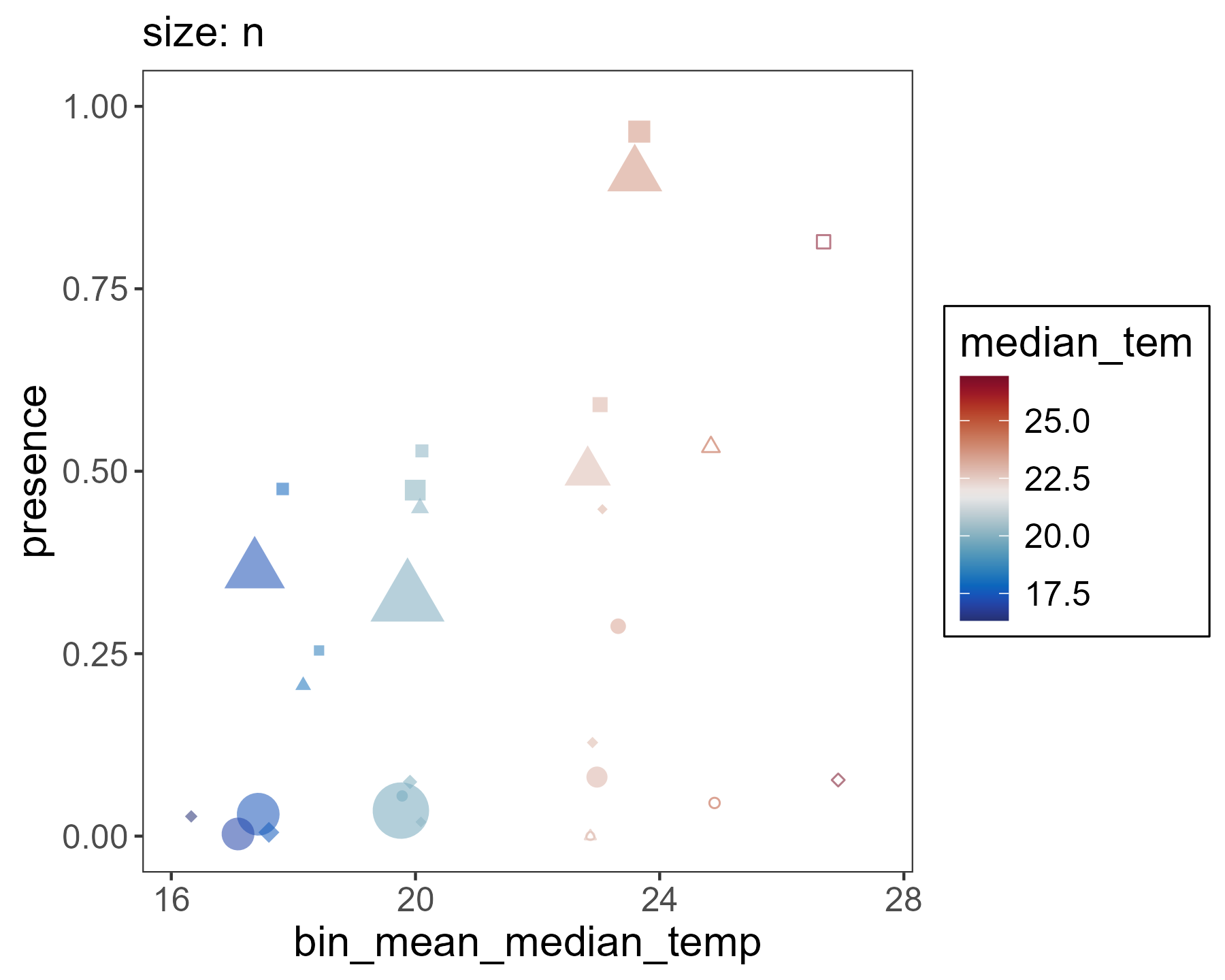

### climate_and_population_GAM_median_temp_reffect_categories_with_segments.png

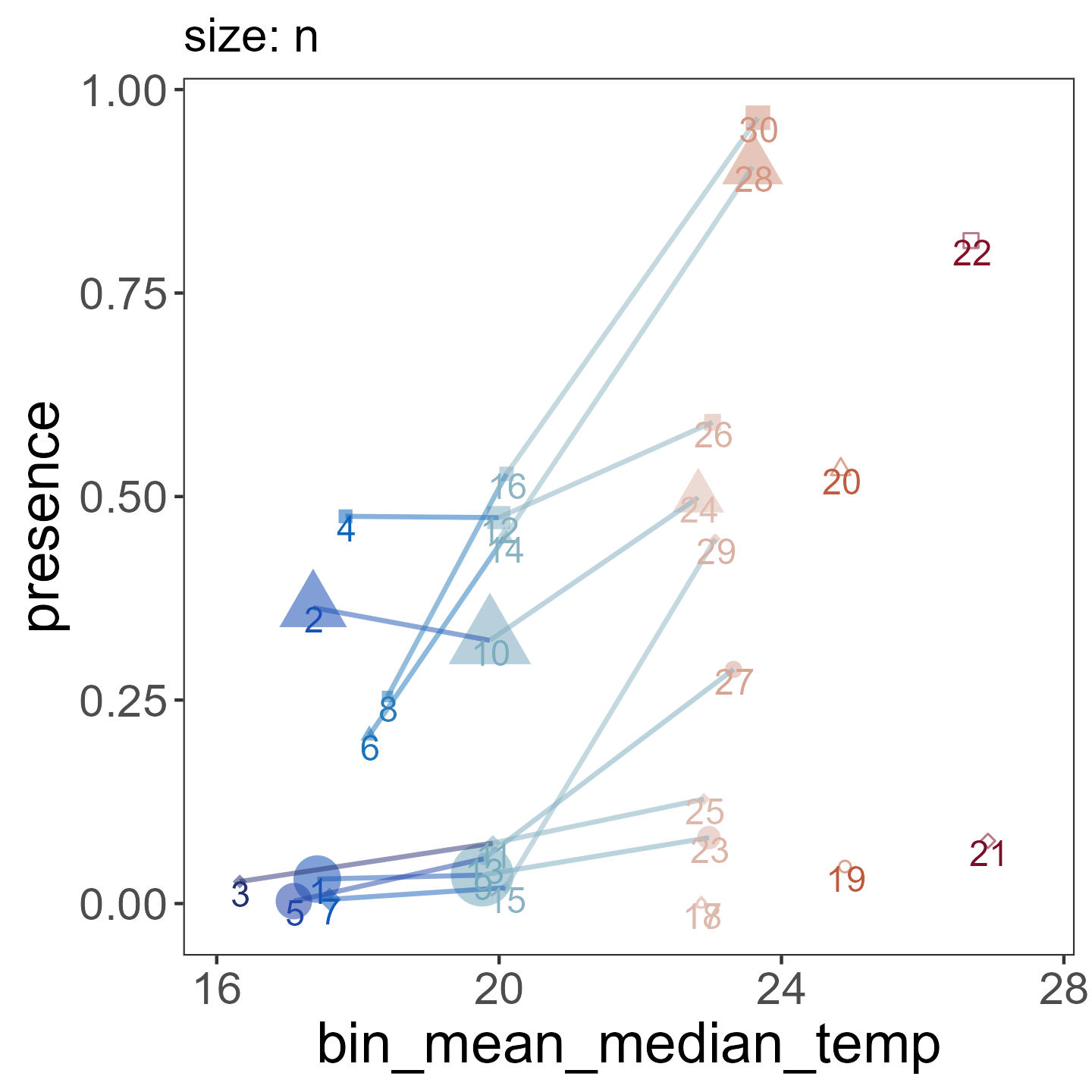

### climate_and_population_GAM_min_temp_CROSS_reffect.png

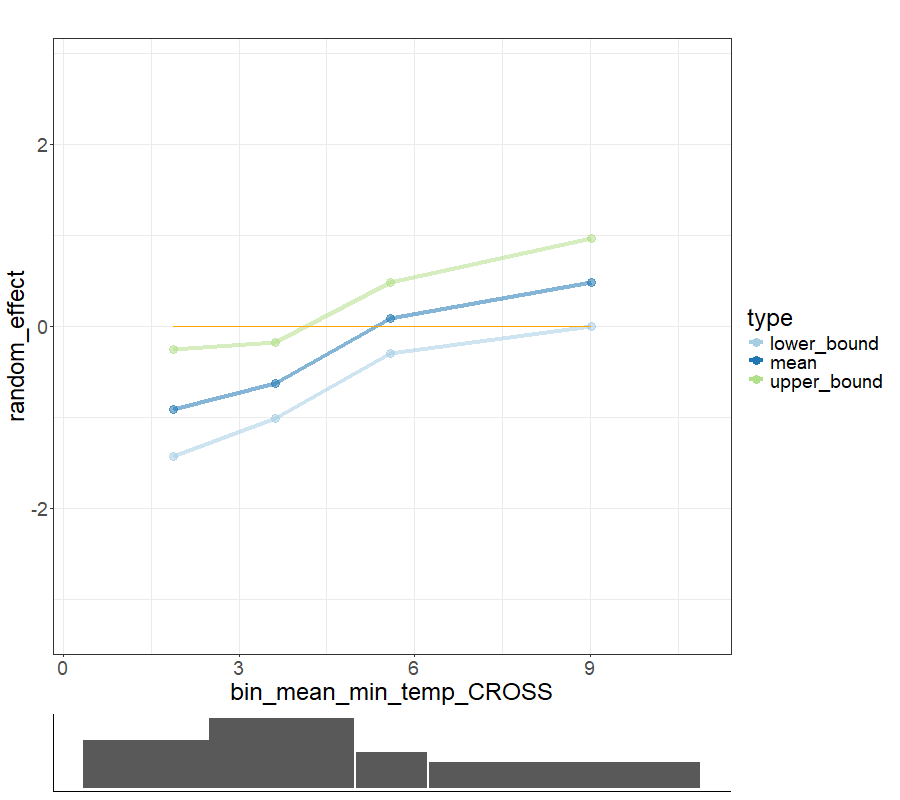

### climate_and_population_GAM_min_temp_CROSS_reffect_categories.png

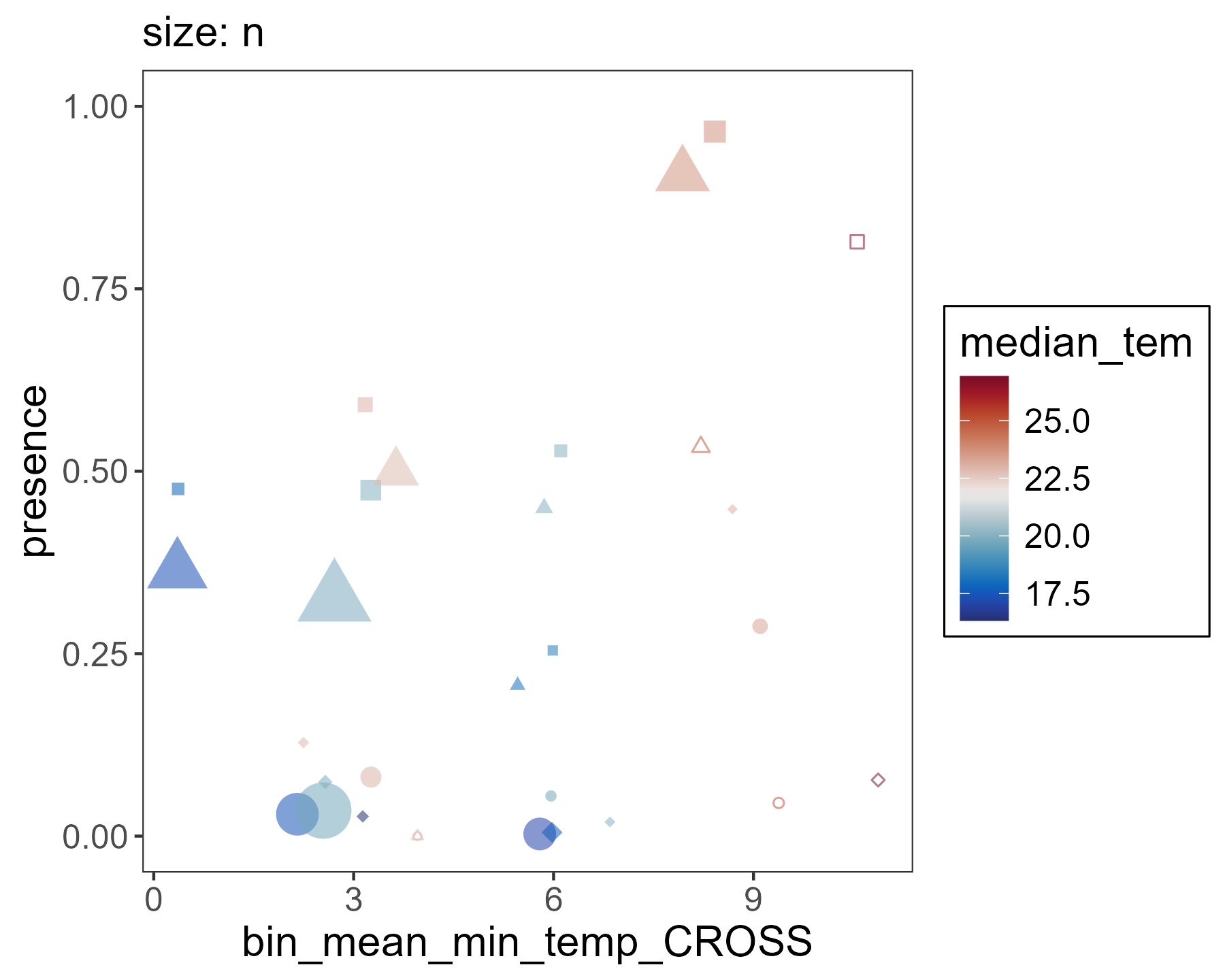

### climate_and_population_GAM_min_temp_CROSS_reffect_categories_with_segments.png

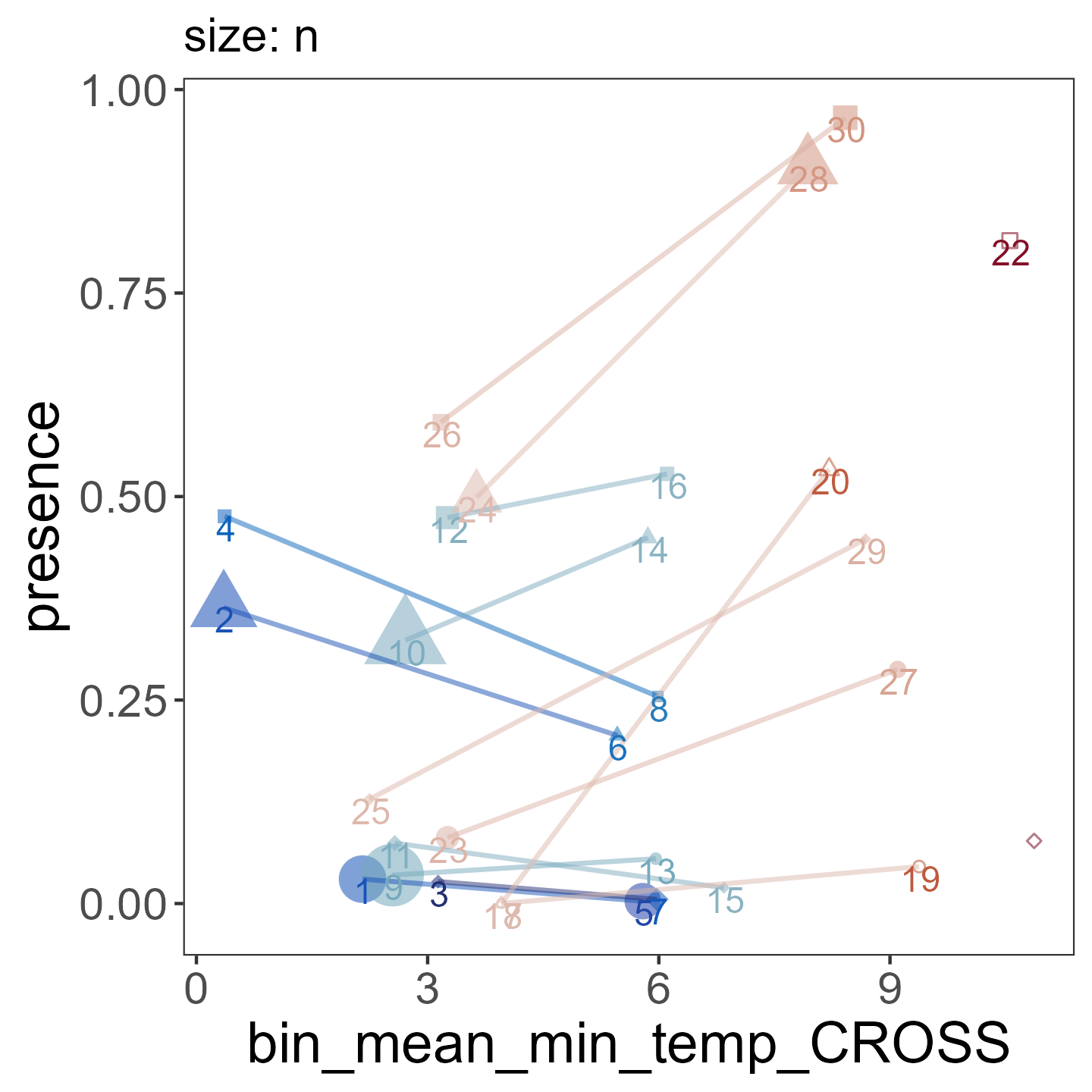

### climate_and_population_GAM_proximity.png

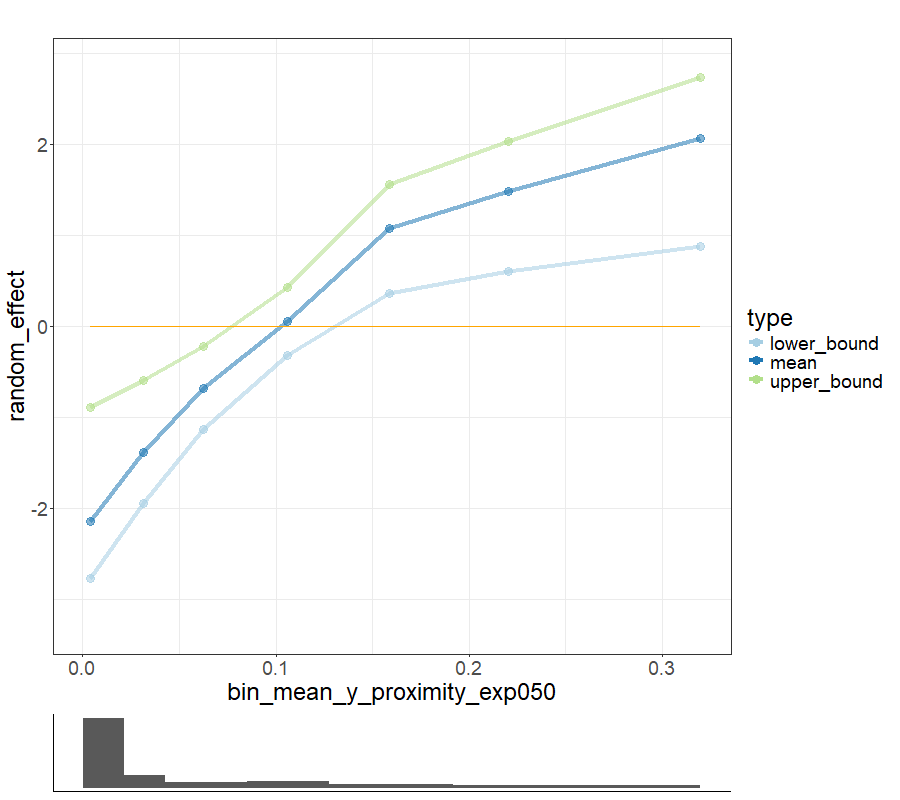

### climate_and_population_GAM_proximity_categories.png

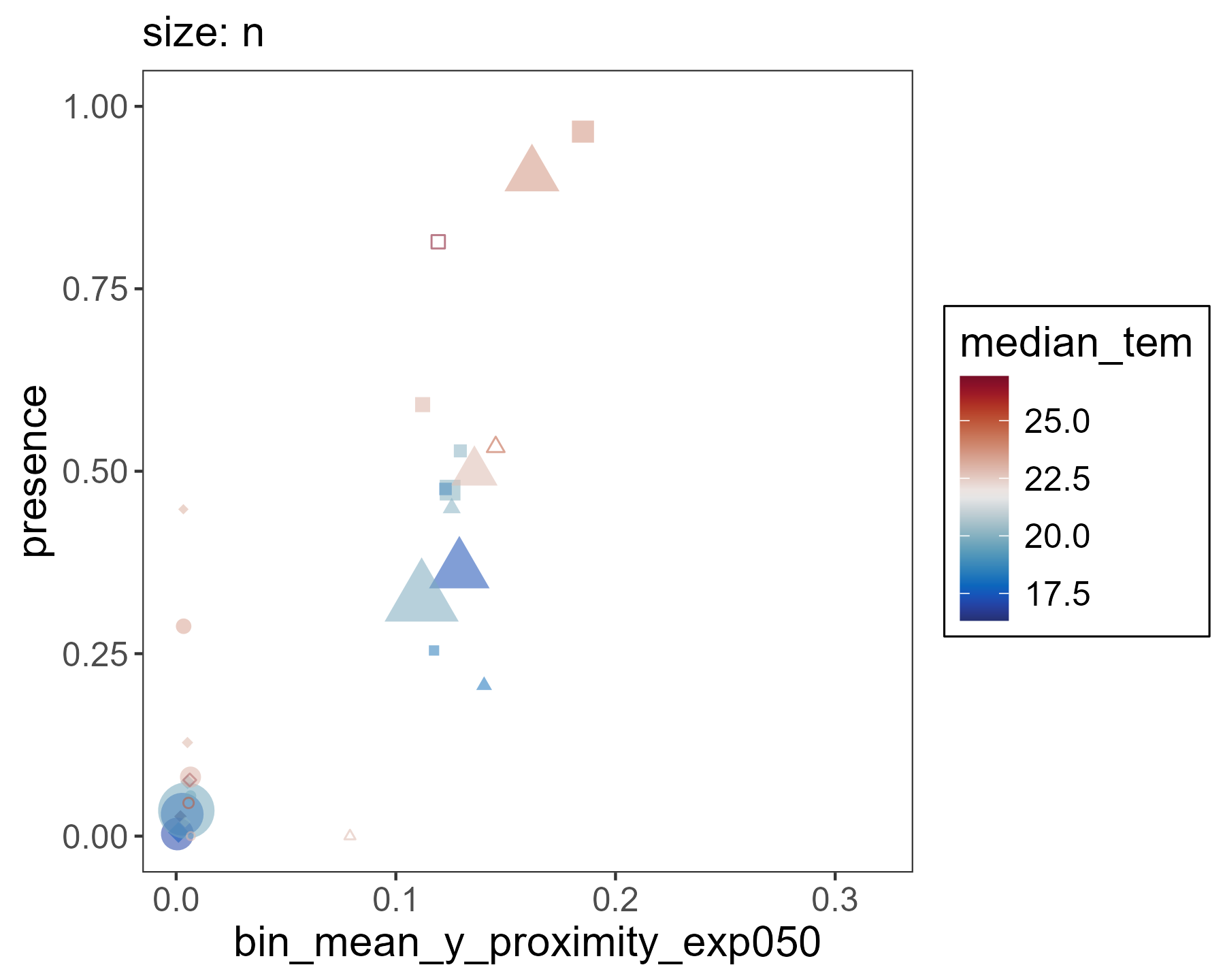

### climate_and_population_GAM_proximity_categories_with_segments.png

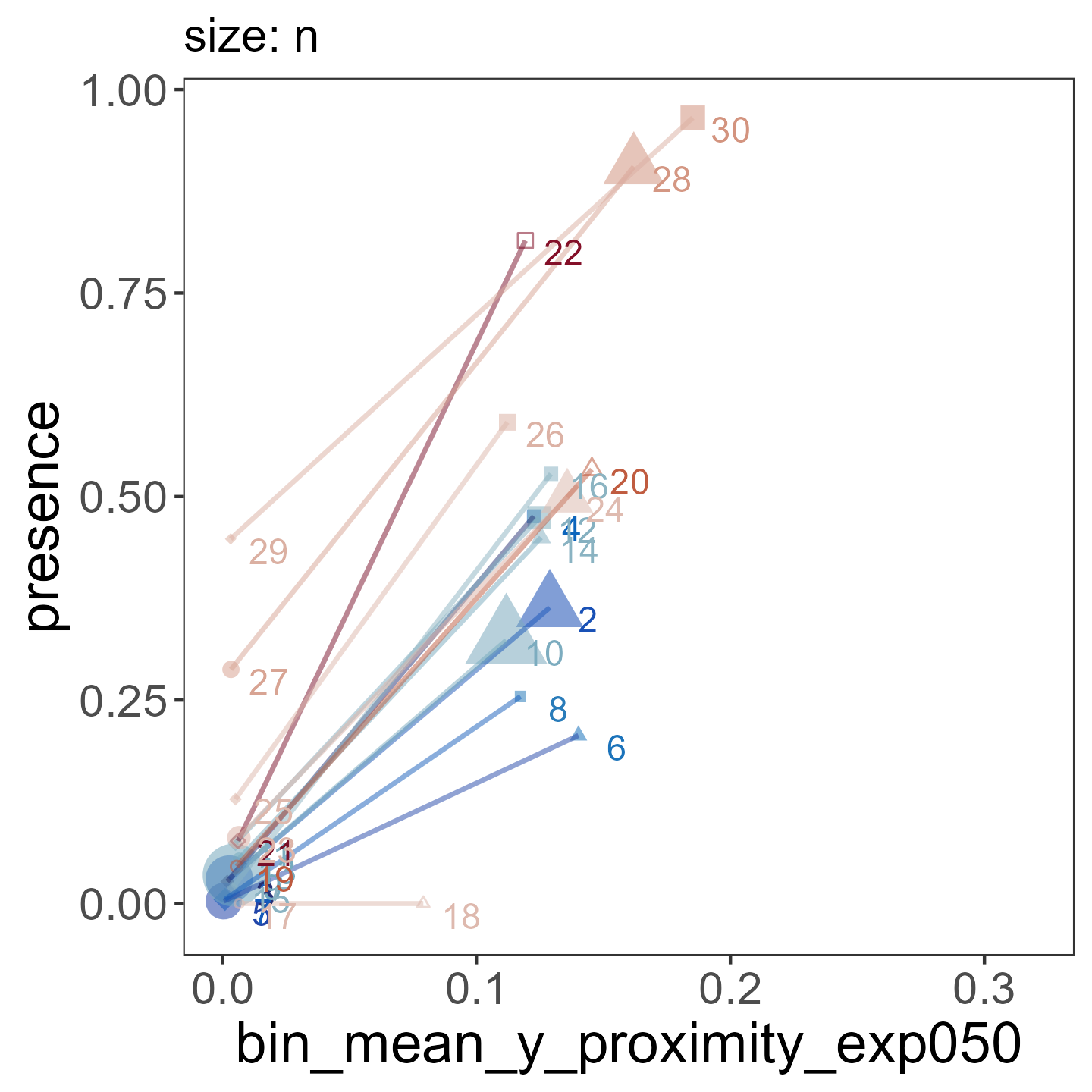

### grouping_ODE_presence_ODE__new_establishments.png

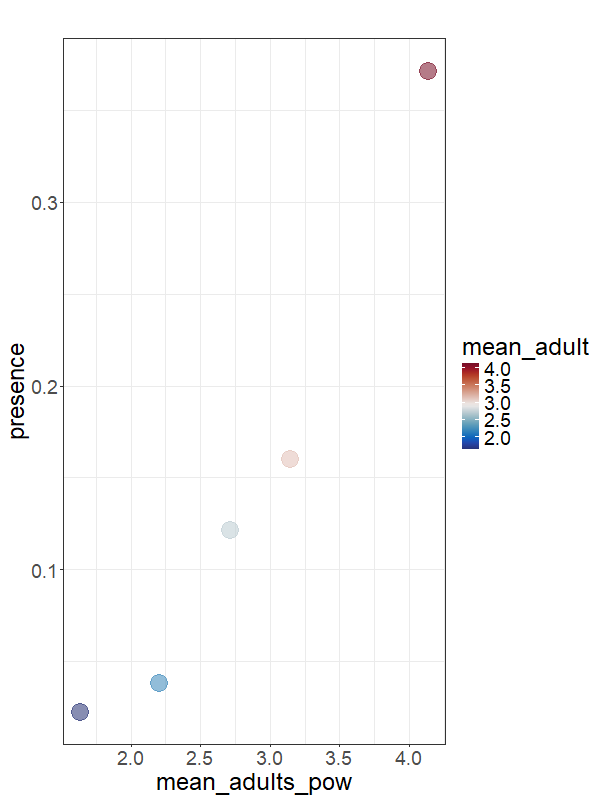

### grouping_ODE_presence_proximity__only_new_establishments.png

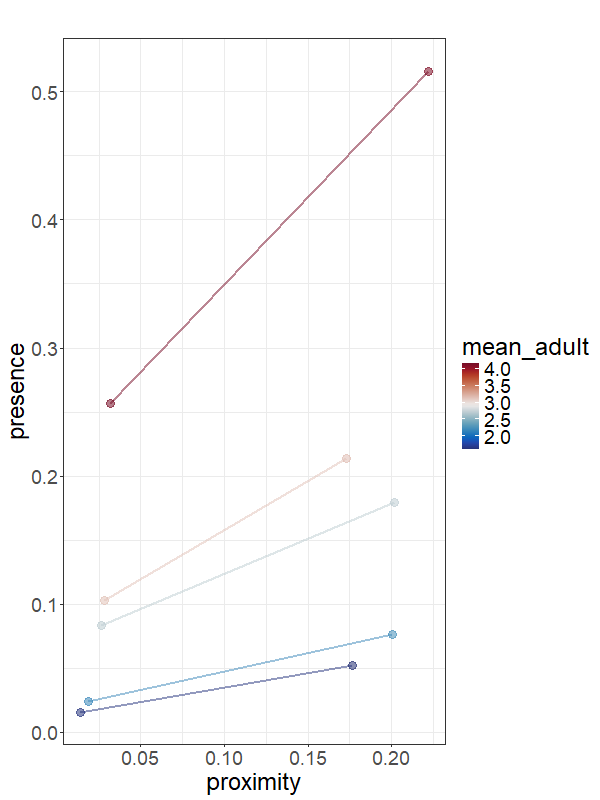

### grouping_presence_median_temp.png

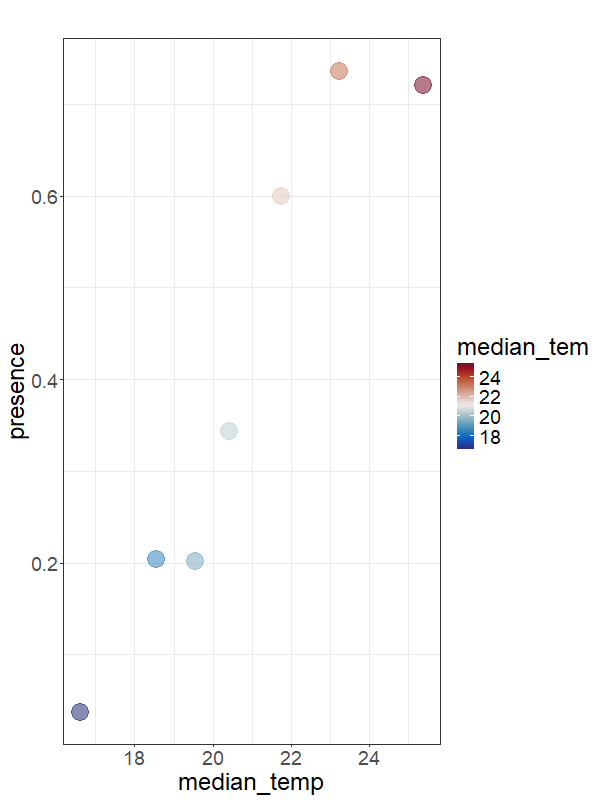

### grouping_presence_min_temp.png

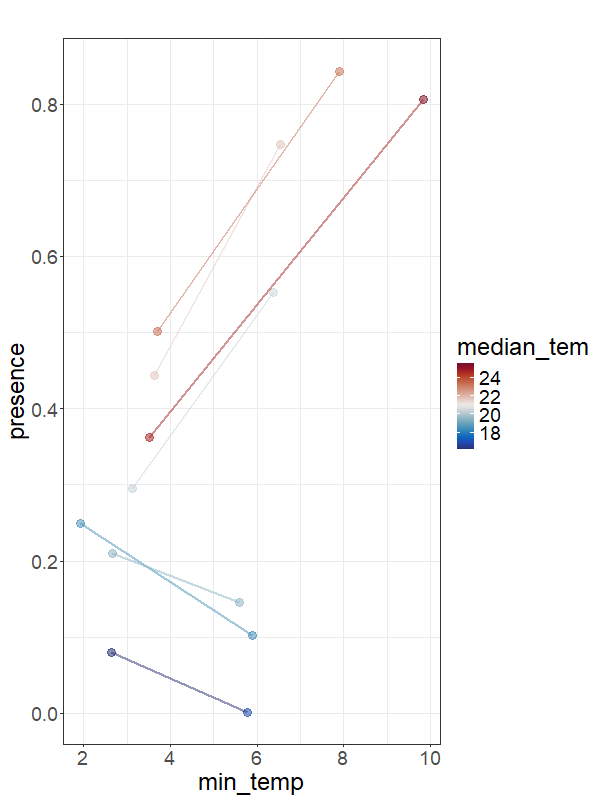

### grouping_presence_relative_humidity.png

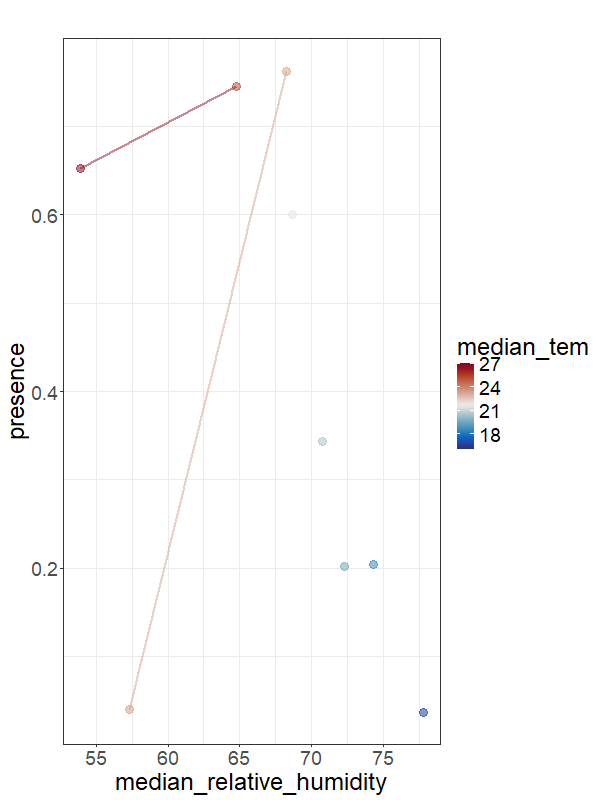

### grouping_WEIGHTEDpresence_log_population.png

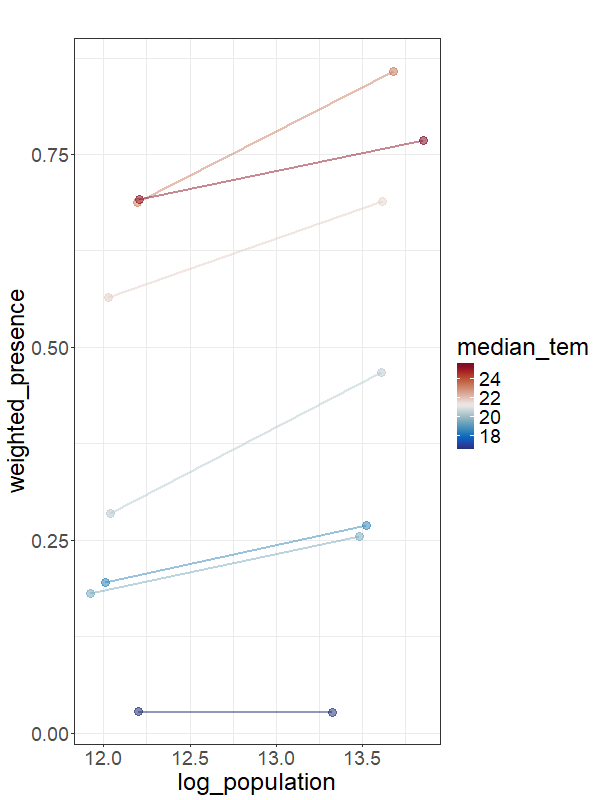

### mechanistic_GAM_model_mean_ODE_merged.png

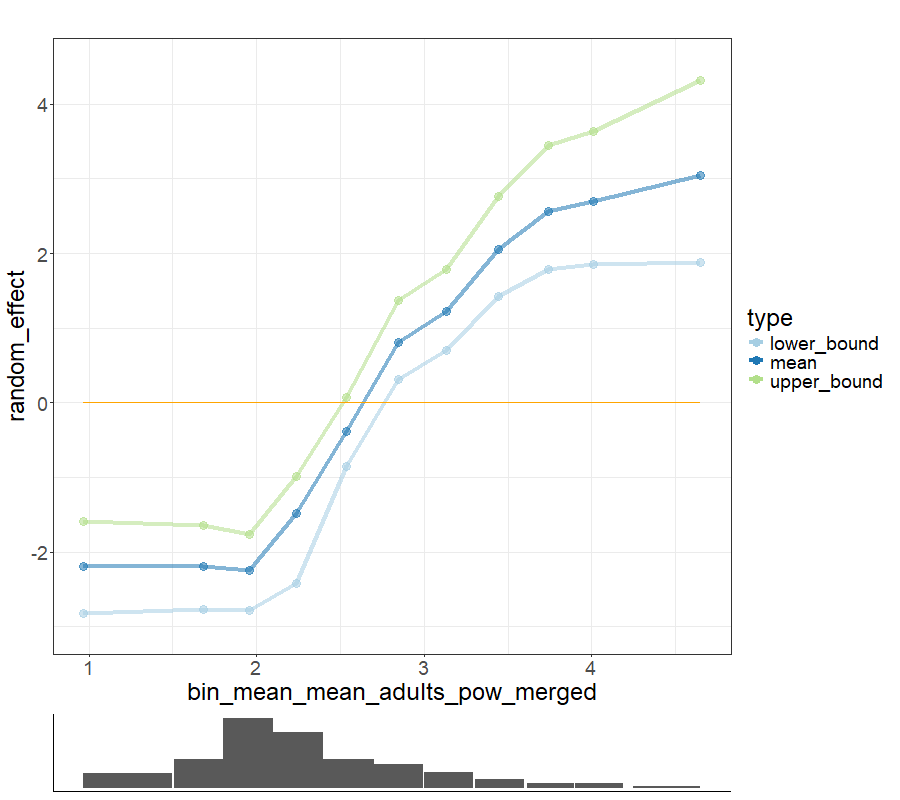

### mechanistic_GAM_model_mean_ODE_merged_categories.png

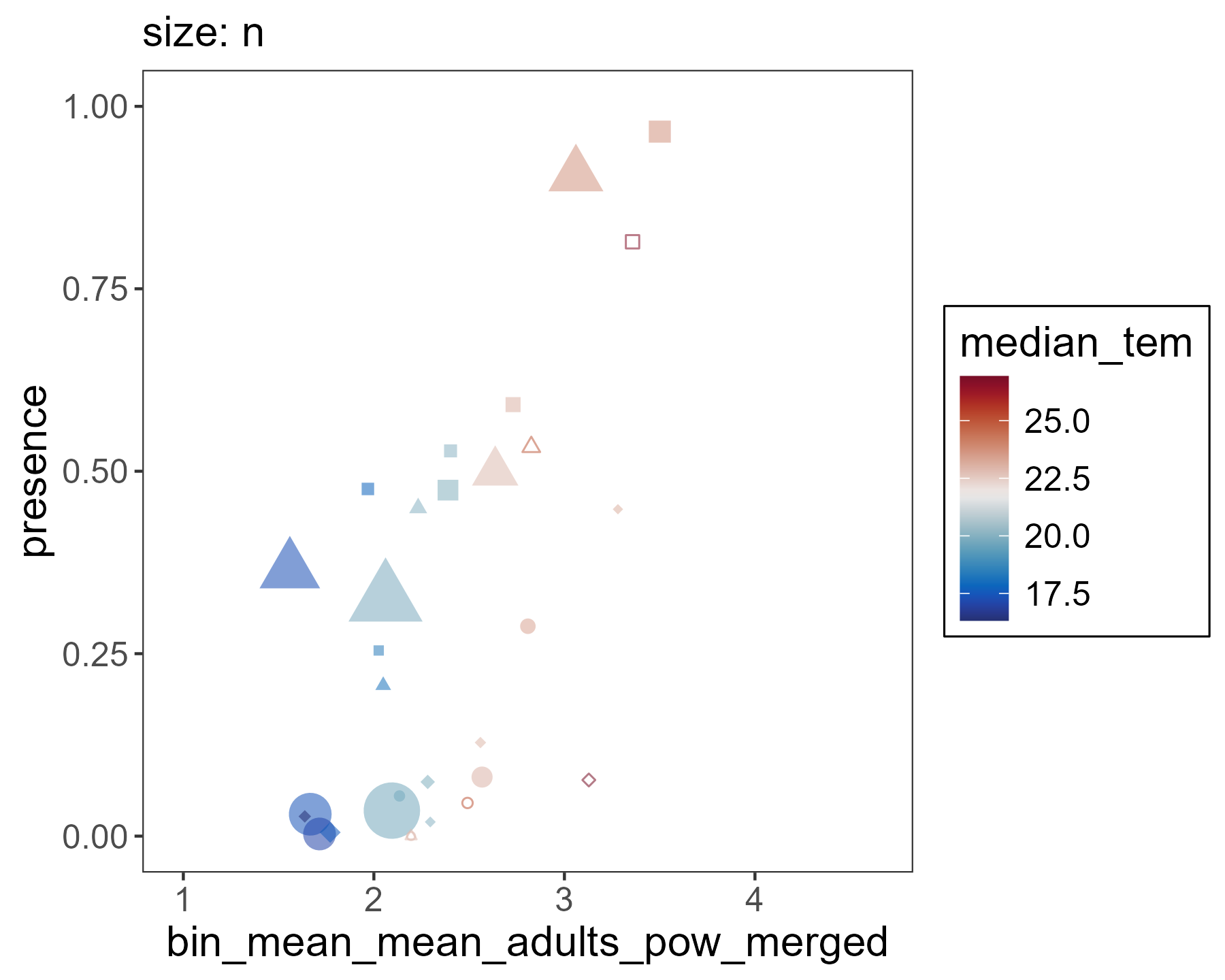

### mechanistic_GAM_model_proximity.png

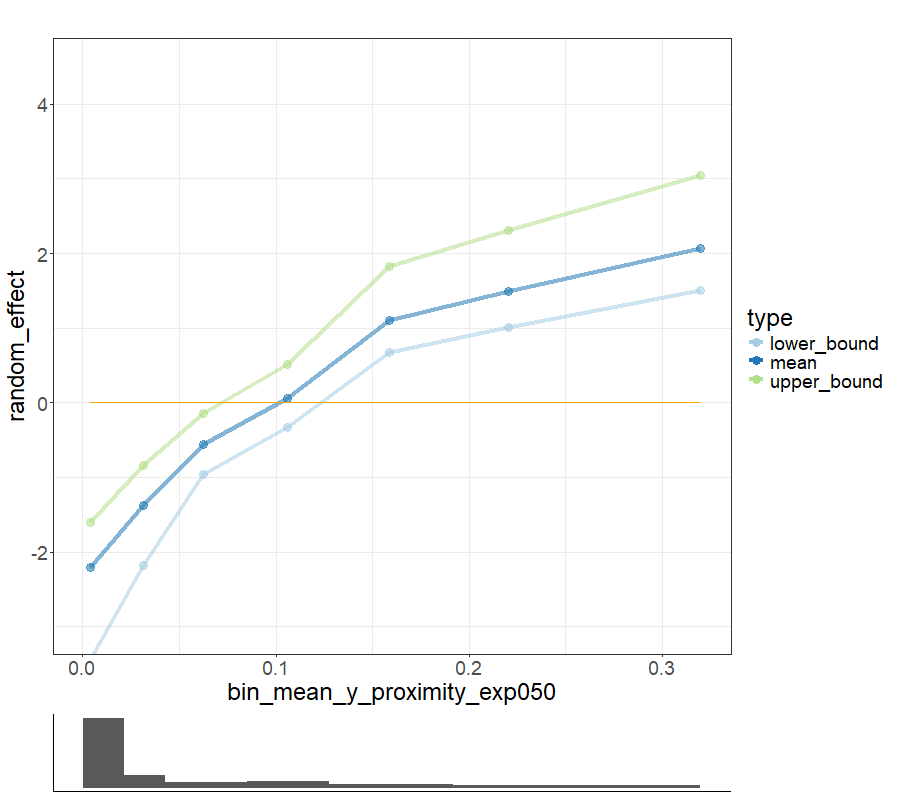

### reffect_mechanistic_covariate_on_map.png

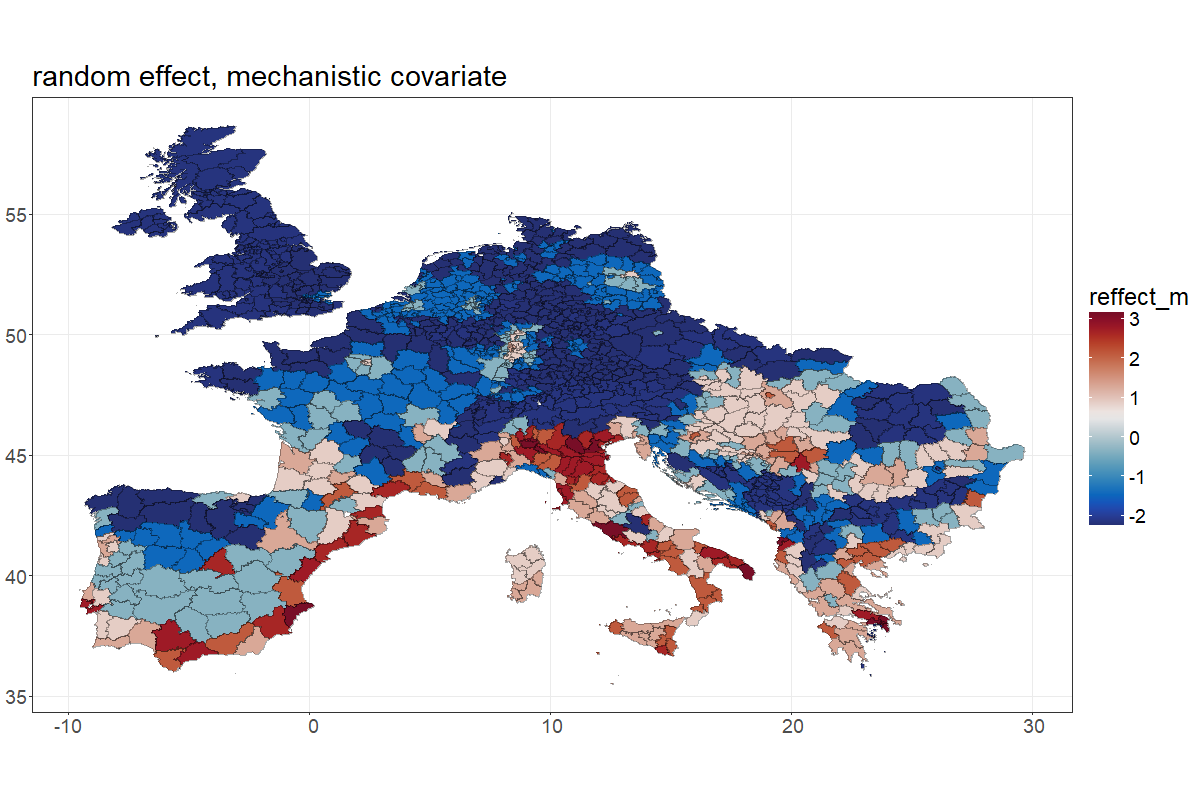

### reffect_median_relative_humidity_on_map.png

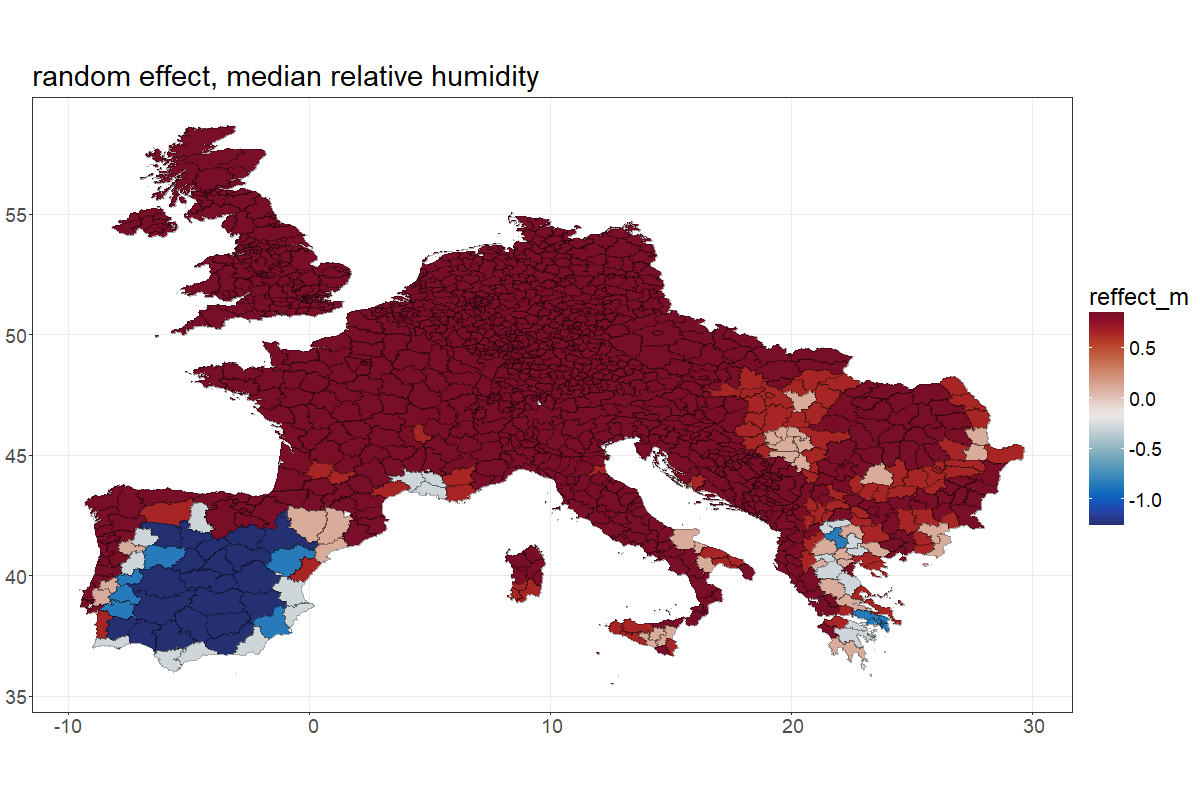

### reffect_median_temp_on_map.png

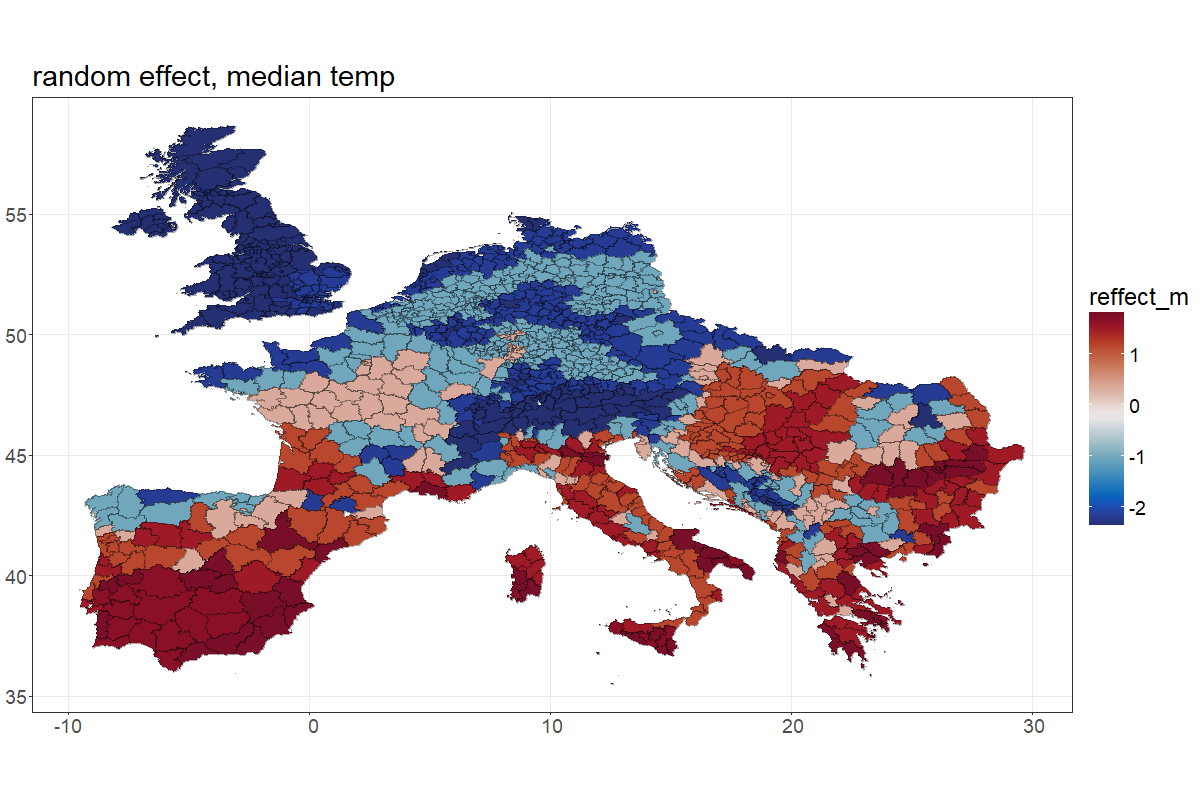

### reffect_min_temp_CROSS_on_map.png

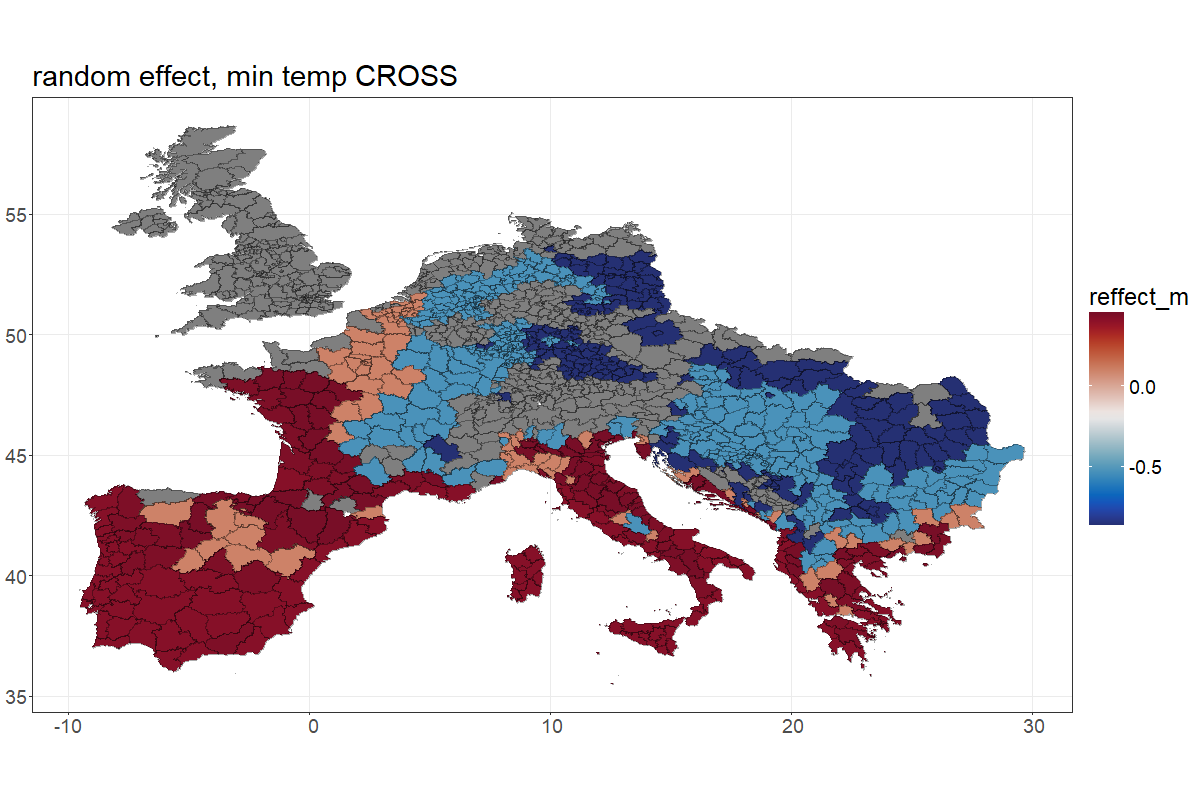
